# Identification of non-conventional small molecule degraders and stabilizers of squalene synthase

**DOI:** 10.1101/2023.06.02.543387

**Authors:** Joseph Hoock, Cecilia Rossetti, Mesut Bilgin, Laura Depta, Kasper Enemark-Rasmussen, John C. Christianson, Luca Laraia

## Abstract

**Graphical abstract:** 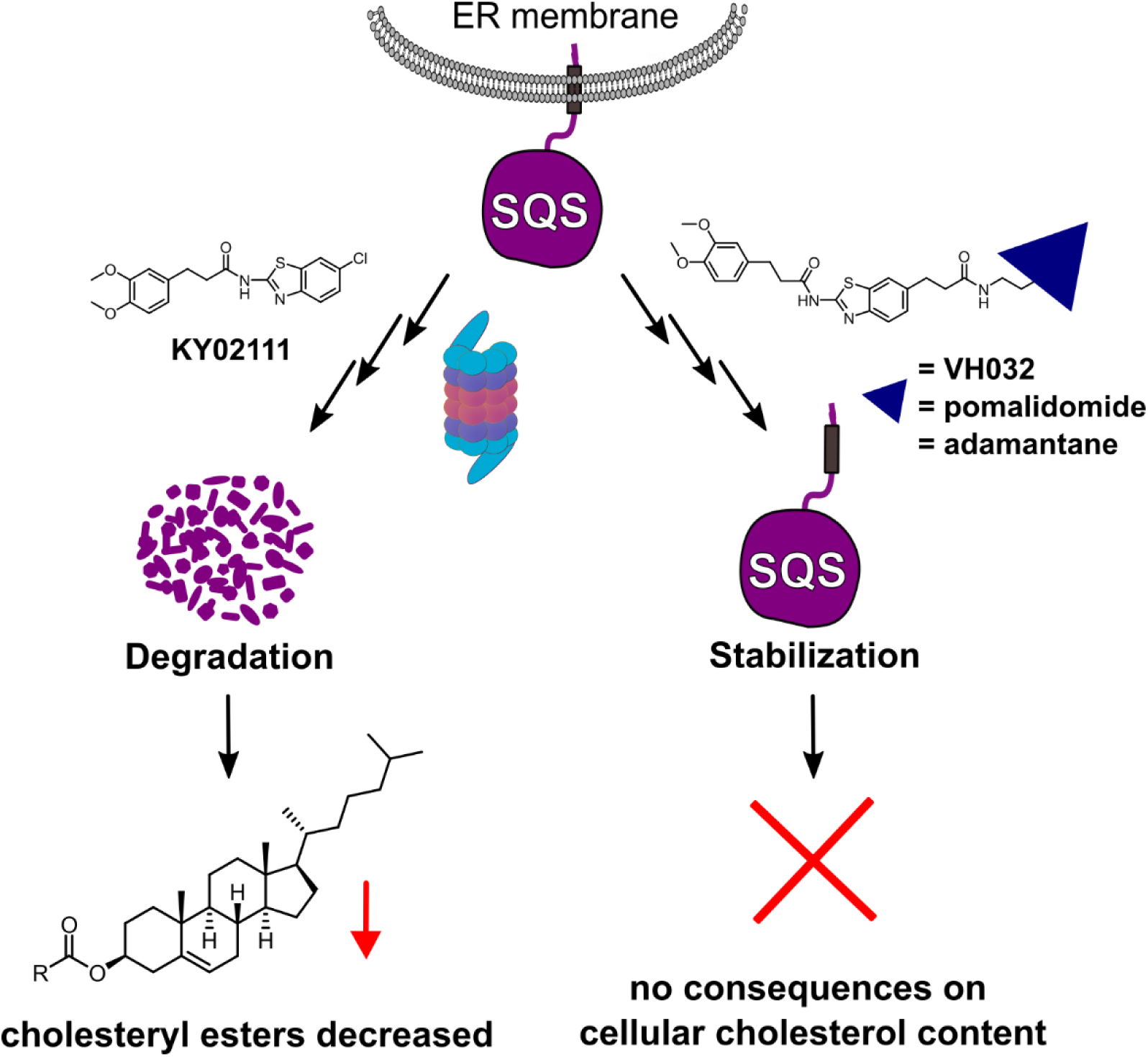

Squalene synthase (SQS) is an essential enzyme in the mevalonate pathway whose abundance and activity control cholesterol biosynthesis and homeostasis. Although catalytic inhibitors of SQS have been developed to attenuate cholesterol, none so far have been approved for therapeutic use. Herein we sought to develop SQS degraders using targeted protein degradation (TPD) as an approach to lower overall cellular cholesterol content. We found that KY02111, a small molecule ligand of SQS, could selectively cause SQS to degrade in a proteasome-dependent manner. In contrast, compounds based on the same scaffold linked to E3 ligase recruiting ligands led to SQS stabilization. Whole cell proteomic analysis found KY02111 to reduce only the levels of SQS, while lipidomic analysis determined that KY02111 treatment concomitantly reduced cellular cholesteryl ester content. SQS stabilizers were shown to shield SQS from its natural turnover without recruiting their matching E3 ligase. Our work shows that degradation of SQS is possible despite a challenging biological setting and lays the groundwork for future development of either SQS degrading or stabilizing probes.

## Introduction

Squalene synthase (SQS), also known as farnesyl-diphosphate farnesyltransferase 1 (FDFT1), is an endoplasmic reticulum (ER) resident membrane protein positioned at a unique branch-point between the sterol– and non-sterol arms of the mevalonate pathway. By catalyzing the condensation of two farnesylpyrophosphate (FPP) molecules into squalene, SQS plays a crucial role in the biosynthesis of cholesterol.[1] Since SQS commits FPP into the cholesterol branch, it can be regarded as a “switch” which is utilized by cells to directly dictate the flow of FPP.[2] Furthermore, additional non-catalytic functions of SQS in the TGFβ pathway have recently been discovered and roles in early embryonic development have been indicated.[3–5] Its unique position in cholesterol biosynthesis has led to the investigation of SQS as a therapeutic target to lower cholesterol levels.[6–8] SQS activity has been linked to hypercholesterolemia-associated diseases, as well as lung– and or prostate cancer[9] or neurodegenerative disorders[10], making the enzyme a desirable drug target. A number of known ligands exists, with zaragozic acid A (ZAA, also known as squalestatin 1)[11] or TAK-475 (also known as lapaquistat acetate)[12] being potent examples of active site binding inhibitors with activity in the nanomolar (nM) range. Despite this, no SQS inhibitor has been successfully brought to the market to date. TAK-475 was the most advanced molecule investigated for the treatment of hypercholesteremia, but development was stopped in phase II and III clinical trials after hepatotoxicity accompanied by elevated bilirubin levels were detected.[13]

With the recent rise of targeted protein degradation (TPD) as a promising therapeutic approach[14], we wanted to revisit SQS as an attractive drug target and develop a probe that could selectively degrade SQS. We hypothesized that a compound able to reduce SQS levels rather than just inhibiting its activity could be an alternative approach to attenuate cholesterol biosynthesis and lower overall cholesterol levels, with possible applications in cancer therapy. Furthermore, SQS degraders could aid the discovery of additional non-catalytic functions of SQS unrelated to the cholesterol biosynthetic pathway.[15]

Herein we report the identification of the small molecule SQS degrader KY02111, a recently reported SQS ligand, as well as the serendipitous discovery of SQS stabilizers.[3] While compounds designed to function as proteolysis-targeting chimeras (PROTACs) based on ligands with diverse SQS binding sites[3,16] led to increased target levels, KY02111 treatment lowered SQS levels in HeLa and U2OS cells in a concentration-, time– and proteasome-dependent manner with excellent selectivity across the proteome. KY02111 did not accelerate the natural degradation of SQS, nor did it affect its insertion in the ER or cause its aggregation. However, partial SQS knockdown using KY02111 led to an overall decrease of cholesterol levels in the form of cholesteryl esters (CE), further supporting its applicability as a tool to study SQS function in a wide range of contexts. Interestingly, envisioned SQS degraders based on the KY02111 scaffold linked to E3 ligase recruiting ligands shielded SQS from its natural degradation by engaging in strong binary interactions with the protein, which might be a relevant strategy for treatment of rare genetic diseases where SQS expression is low.[5]

## Results

### Synthesis of an SQS degrader compound library based on two distinct ligands

We initiated our efforts to identify a small molecule degrader of SQS by synthesizing a small yet diverse degrader library based on two reported SQS-ligands: an active site inhibitor, herein referred to as SQSI, and an unknown-binding site ligand known as KY02111 (Fig. 1A). To cover as large a chemical space as possible, each ligand was diversified by varying linker length and composition as well as by generating different protein of interest (POI) ligand-degrader modality pairings.

**Figure 1:**
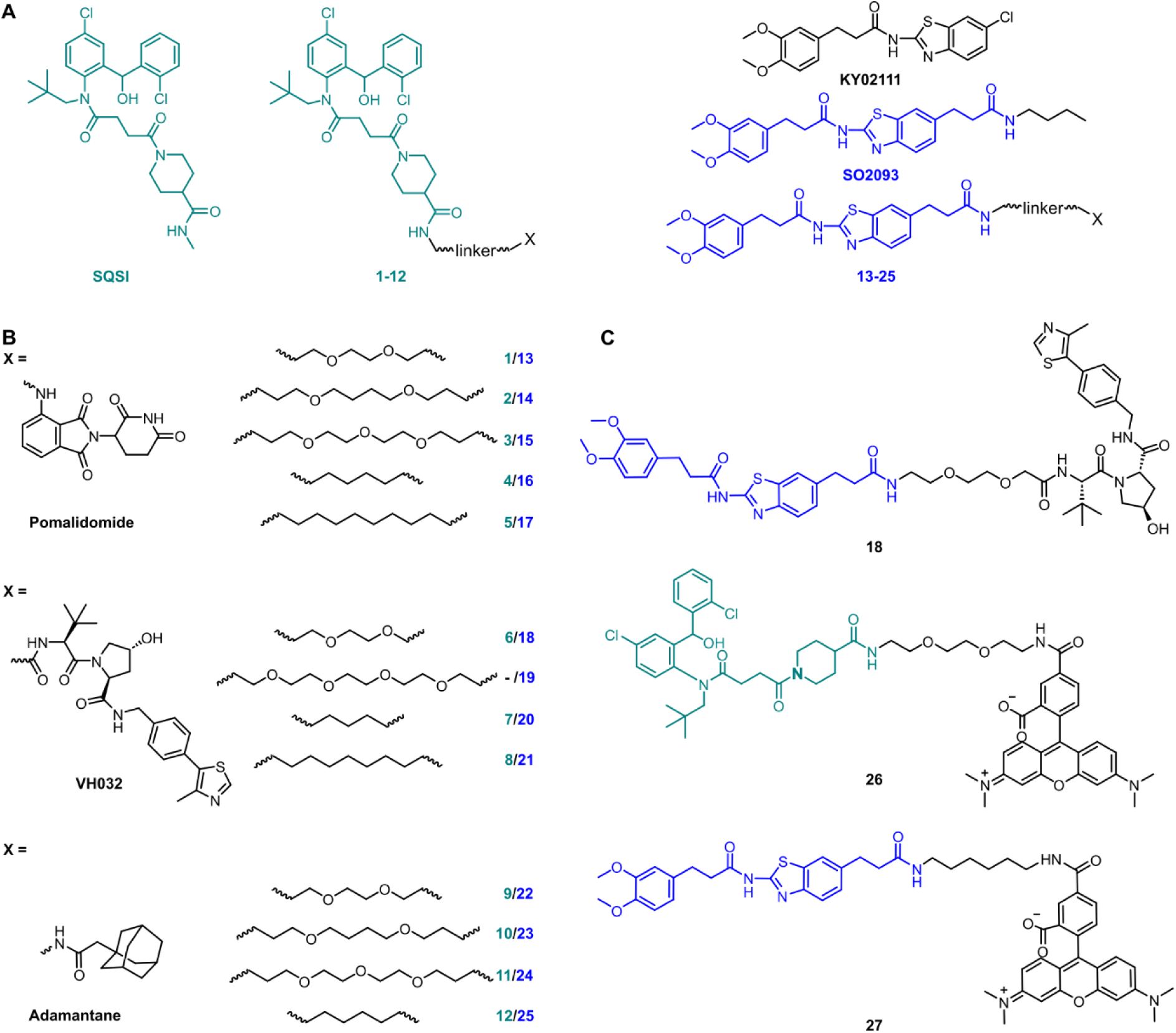
Full SQS-degrader library. **A**) Structure of SQS ligands used to design PROTACs: one active site inhibitor (SQSI, turquoise) and one unknown binding site ligand (KY02111, blue). The final envisioned SQS degraders connect the SQS ligands at positions which are accepted for modification without interfering with target engagement. **B)** The final compounds contain one of three different degrader modalities (X): pomalidomide, VH032 and an adamantane tag. The SQS-ligands and the degrader modalities were linked by PEG or alkyl-based linkers of varying lengths. **C)** Full structure of KY02111-based PROTAC **18** and fluorescent probes based on the parent SQS ligands linked to 5-carboxytetramethylrhodamine (TAMRA) via linkers.

SQSI was part of a 2-aminobenzhydrol compound series developed by Daiichi Sankyo, which inhibited SQS activity in rat hepatocytes with an IC_50_ of 1.3 nM.[16] Additional interest in this compound was sparked by the fact that the crystal structure of a close analogue revealed exposure of the piperidine-4-carboxamide to the cytosol (PDB: 3ASX), making it a desirable linker attachment position (Fig. 1A).[16] Together with a readily accessible synthetic route, SQSI was an ideal compound for our efforts. On the other hand, KY02111 was only recently identified as a SQS-binder.[3] KY02111 showed no catalytic inhibition in an *in vitro* activity assay as well as in an *in cellulo* SREBP reporter gene assay. The authors found that KY02111 likely interferes with a so far unknown protein-protein interaction (PPI) between SQS and TMEM43, which resulted in downregulation of TGFβ-induced signaling. A reduction of SQS levels upon compound treatment was not observed by the authors. Overall, the work suggested that KY02111 likely does not bind SQS at the same site as SQSI. To increase potential ternary complex diversity and thereby the probability of success in identifying an SQS degrader, we based the second half of our library on a KY02111 analogue (SO2093, Fig. 1A).

We implemented additional structural diversity by attaching the two SQS-ligands to three different degrader modalities via a variety of linkers (Fig. 1B). For the formation of PROTACs, we utilized the well described[17,18] *Cereblon (CRBN)* and *von Hippel Lindau (VHL)* ligands pomalidomide[19] and VH032[20], respectively. As a less frequently used degrader modality of TPD, we generated hydrophobic tags by attaching an adamantane moiety.[21] We linked both ligands via readily available flexible first generation linkers varying in length (8-15 total atoms for pomalidomide and adamantane, 7-16 total atoms for VH032) and composition (PEG and alkyl).

Synthesis of the full SQS-degrader library was conducted using procedures reported previously (SI Fig. 1/2). While preparing compounds based on SQSI we observed the presence of four different isomers when the samples were dissolved in DMSO. These were fully assigned using variable temperature NMR as two sets of interconvertible atropisomers (SI Fig. 3).

### *In vitro* binding assays to recombinant SQS

We initially screened the SQS-degrader library for binary target engagement. We first expressed a catalytically active SQS construct (31-370) lacking the C-terminal transmembrane domain in *E. coli*.[22] With the recombinant protein in hand, we set up two individual *in vitro* screens to evaluate SQS-binding by our compounds – differential scanning fluorimetry (DSF) and fluorescence polarization (FP). DSF measurements (SI Fig. 4) showed dose-dependent stabilization with an increase in melting temperature (ΔTm) of up to 6.3 °C for the active site-based compounds (SQSI, **6**, **7**, **8**). The non-active site compounds inspired by KY02111 also stabilized SQS, yet not to the same extent (ΔTm ≤ 3.6 °C) and with diminished dose-dependency compared to the first half of the compound series.

To confirm the observed target binding, we also developed a fluorescence polarization (FP) assay by synthesizing two 5-TAMRA conjugated probes based on SQSI (**26**) and KY02111 (**27**) (Fig. 1C). Protein titration with both fluorescent probes showed strong binding of SQS with similar affinities reflected by *K_d_* values around 100 nM (Fig. 2A). Competition experiments of the SQS-degrader library versus the corresponding FP probe (**1-12** vs **26**, **13-25** vs **27**) yielded the IC_50_ and *k_i_* values shown (Fig. 2C). All compounds (except **5**) were able to outcompete their parent probe. A general trend can be observed where increasing affinity is correlated with attachment of a linker-degrader modality in comparison to the unmodified ligands (e.g. **3**, **4**, **7**, **11** compared to SQSI; **14**, **18**, **19**, **22**, **23** compared to KY02111). Three out of four PROTACs containing the C9/10 alkyl chain (**5**, **8**, **17**) make an exception to this rule due to decreased molecular flexibility and solubility. Importantly, both of the active site (SQSI) and unknown binding site ligands (KY02111, **18**), were able to outcompete both active site (**26**) and unknown site (**27)** probes (Fig. 2B). For example, SQSI showed slightly stronger displacement of fluorescent probes when competed versus **27** (*k_i_* = 0.3 µM) than versus **26** (*k_i_* = 0.7 µM). This was surprising since data by Takemoto *et al.* showed that KY02111 did not inhibit the catalytic activity of SQS, leading to the hypothesis that the interaction occurs outside of the active site. Furthermore, the data suggested a binding site on the external surface since KY02111 inhibited the PPI between SQS and TMEM43.[3] We hypothesized that our FP results could be explained by either a close proximity for the two binding sites or a conformational shift induced upon compound binding.

**Figure 2:**
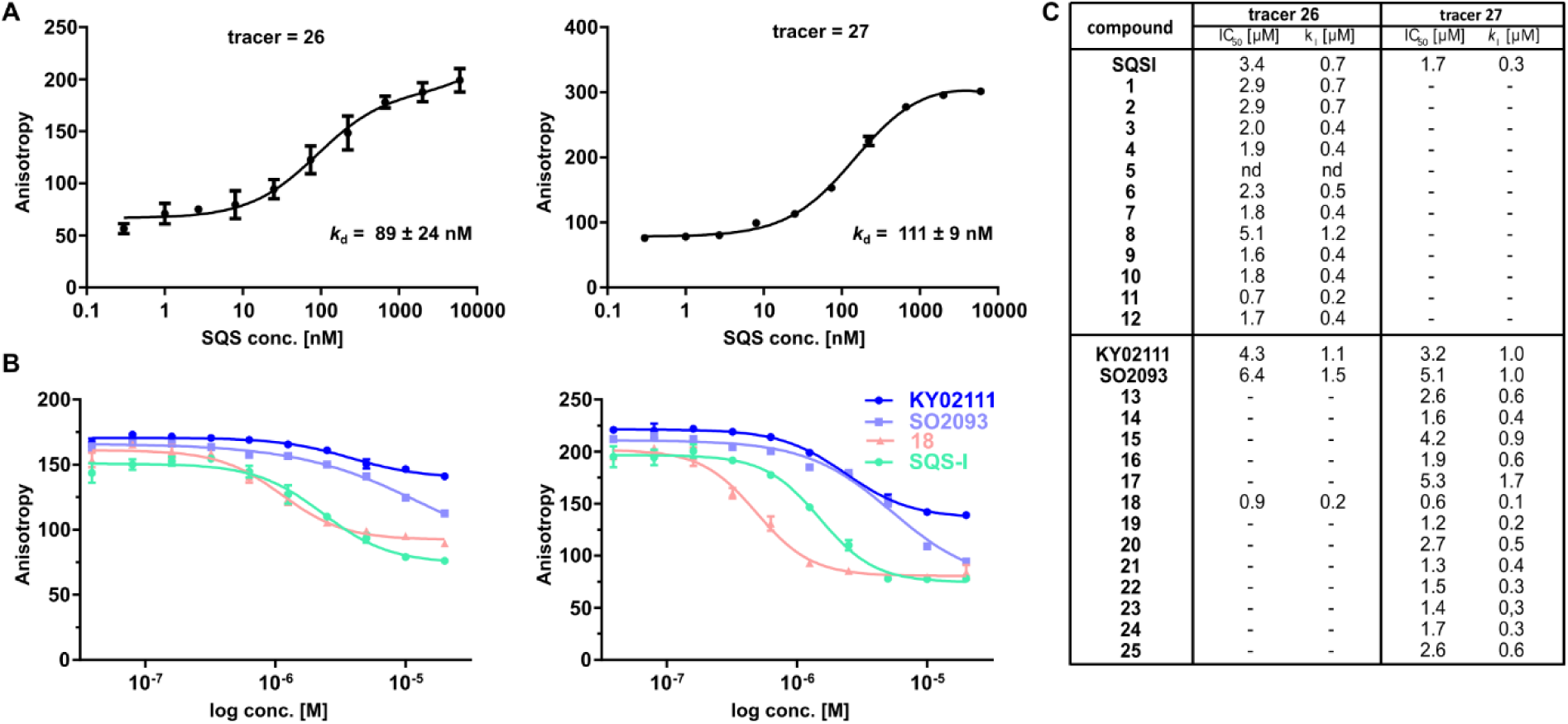
FP assay confirms binding of degrader library to recombinant SQS *in vitro*. **A**) Titration of recombinant SQS vs active site tracer **26** (c = 20 nM) and unknown site tracer **27** (c = 5 nM) (n = 2; conducted in technical duplicates; mean ± SEM shown.). **B)** Competition of selected compounds vs **26** and vs **27**. **26** can be outcompeted by compounds assumed not to bind the active site (**KY02111**, **SO2093**, **18**). Vice versa, **27** can be outcompeted by the active-site binding **SQSI**. (n = 4, conducted in technical duplicates, mean ± SEM shown). **C)** Competition experiments of the full library vs either active site FP probe **26** or unknown binding site probe **27** showed that all compounds (except **5**) are able to bind SQS *in vitro* (n = 1-4; mean shown).

### Identification of SQS degraders and stabilizers

While *in vitro* assays showed that linker attachment to both ligands would not interfere with SQS engagement, we probed our library for degradation ability in HeLa cancer cells. Since endogenous SQS is constitutively turned over with a modest half-life (*t_1/2_* ∼ 5 h)[23] we incubated the compounds for 18 h to ensure clear correlation between lower SQS abundance and compound treatment. Intensive screening via western blotting identified KY02111 as the most potent SQS-degrader, whereas the designated degrader molecules almost exclusively lead to an increase in SQS levels (Fig. 3A/B). PROTACs (**1-8**, 30 nM to 3 µM) and HyTs (**9-12**, 2 and 20 µM) derived from SQSI universally lead to an increase in SQS levels at all tested concentrations. Remarkably, KY02111 derived compounds **13** (at 3 µM), **18** (at 3 µM), **21** (at 3 µM) and **25** (2 and 20 µM) all showed a similar trend (Fig. 3A/B).

**Figure 3:**
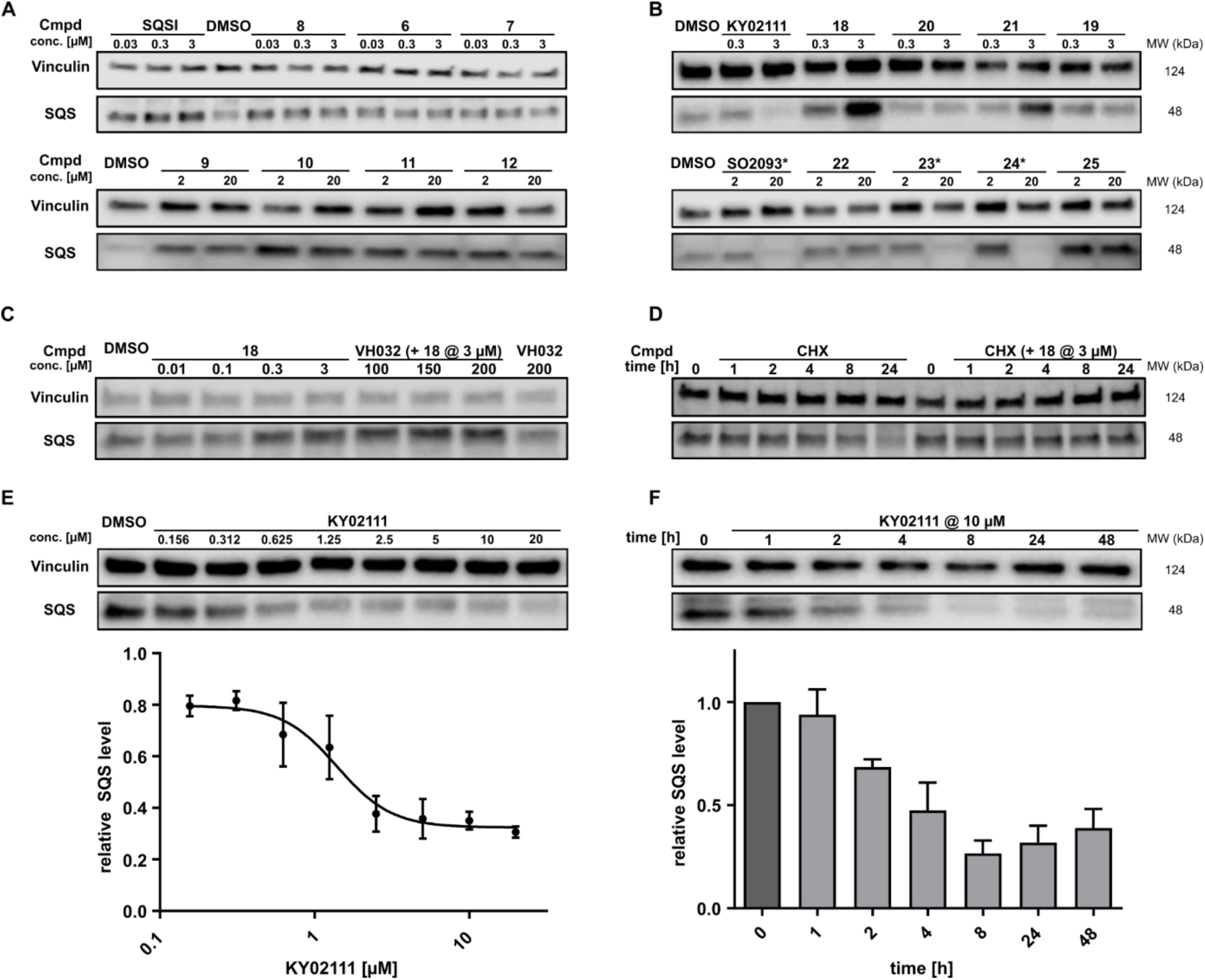
*In cellulo* SQS-degrader screen reveals KY02111 as only SQS-degrader. **A/B**) Western blots showing the changes in SQS protein levels after 18 h treatment of HeLa cells with compounds at indicated concentrations (n = 2 or 3). * compounds precipitated at 20 µM and caused cell death. **C)** SQS accumulation by PROTAC **18** is independent of VHL. HeLa cells were treated with **18** and/or VH032 for 18 h (n = 3). **D)** CHX co-incubation shows that PROTAC **18** stabilizes SQS rather than upregulating it. HeLa cells were treated with **18** (3 µM) and/or CHX (100 mg/mL) for the indicated time points (n = 3). **E)** KY02111 reduces SQS levels in a dose-dependent manner. The concentration to reduce SQS levels by half is reached at DC_50_ = 1.4 µM and overall SQS levels can be reduced to D_max_ = 68%. HeLa cells were treated with the indicated concentrations of KY02111 for 18h (n = 4; mean ± SEM shown). **F)** KY02111 reduces SQS levels in a time-dependent manner. HeLa cells were treated with KY02111 (10 µM) for the indicated time points (n = 3; mean ± SD shown).

DSF and FP measurements showed target binding by most of our library molecules (SI Fig. 4 and Fig. 2, respectively). Additionally, we observed a clear change in SQS band intensity for compound treated samples compared to DMSO controls, which we correlated to SQS binding *in cellulo*. Together with the fact that compounds **1-12** are based on an active site inhibitor, we hypothesized that these molecules still act as inhibitors. Instead of forming a productive ternary complex only a binary complex with SQS is formed, thereby blocking the catalytic activity, and leading to compensatory upregulation. We confirmed this by monitoring expression of the sterol-regulated transcription factor SREBP using a reporter assay, which showed increased activity for cells treated with SQSI (SI Fig. 6). Alternatively, another reason for observation of increased SQS levels, for example for VHL-PROTAC compound **18** which is based on KY02111, could be a stabilizing ligand-SQS interaction interfering with the natural degradation of SQS (Fig. 3B, right). For compound **18**, we detected elevated POI levels for incubation at 3 µM compared to 0.3 µM and DMSO. To exclude the possibility of an early Hook-effect[24,25] we incubated HeLa cells with even lower concentrations of compound **18**. Yet, we could not detect any degradation (Fig. 3C). A recent study by Poirson *et al.*[26] showed that VHL can also act as a protein stabilizer instead of a degrader, possibly due to the formation of a non-productive ternary complex. We co-incubated compound **18** with a large excess of VH032, the parent VHL-ligand to compete with a potentially stabilizing ternary complex (Fig. 3C). Since SQS levels are still increased in the presence of VH032, the observed SQS stabilization is likely based on a binary interaction between compound **18** and SQS rather than a non-productive ternary complex. This is supported by our finding that co-incubation of compound **18** with the protein biosynthesis inhibitor cycloheximide (CHX)[27,28] leads to a stabilization of SQS levels after 8 and 24 h compared to CHX-treated controls (Fig. 3D). Of note, **18** showed the highest degree of stabilization (T_m_ = 3.6°C, SI Fig. 4C) for the compound series based on KY02111 and the overall highest affinity (*k*_I_ = 0.1 µM) for SQS in our FP assay (Fig. 2B/C) which is in agreement with the above observations. Based on this, we believe that bifunctional compounds based on KY02111 linked to an E3 ligase ligand cause an increase in SQS band intensity by stabilizing SQS through tight binary binding interactions.

In addition to KY02111, we observed a dose-dependent decrease in SQS abundance for three additional hydrophobic molecules: SO0293 and the hydrophobic tag-containing compounds **23** and **24** (Fig. 3B, right). However, during sample preparation, a clear increase in cell death in samples treated with 20 µM of these compounds was seen, which was accompanied by compound precipitation. We confirmed this cytotoxicity for SO2093, and compounds **23** and **24** by assaying cell viability (SI Fig. 7A/B). Since KY02111 treatment did not alter cell viability after 48 h, we hypothesized that the decrease in SQS levels by SO2093, compounds **23** and **24** was caused by an unspecific mode of action (MoA) and therefore excluded the compounds from further testing.

### KY02111-induced SQS degradation is dose-, time– and proteasome-dependent

To further characterize the reduction of SQS levels induced by KY02111 treatment, we performed dose-response and time-dependent experiments in HeLa cells (Fig. 3E and 3F, respectively). The dose response of KY02111 yielded a DC_50_ of 1.4 µM and a D_max_ of 68 %. However, we were not able to induce complete degradation even when increasing the concentration to as high as 20 µM (Fig. 3E). To exclude the possibility of a cell line dependent effect, we treated U2OS cells with KY02111 and observed a similar concentration-dependent reduction in SQS levels (SI Fig. 8). Additionally, we found that the effect on SQS was time-dependent, with maximum levels of degradation reached between 8 and 24 hours, after which protein levels slowly recover (Fig. 3F).

Next, we sought to determine which cellular degradation machineries were responsible for the loss of SQS protein. HeLa cells were pre-incubated for 2 h with either MG132 (proteasomal inhibitor), MLN4924 (neddylation inhibitor) or chloroquine (CQ, lysosomal inhibitor) followed by co-incubation with 10 µM KY02111 for 18 h (Fig. 4B). MG132 was clearly able to block the KY02111-induced SQS degradation, whereas MLN4924 could not rescue protein levels. Interestingly, co-incubation with chloroquine (CQ) led to an increase in the SQS band intensity. However, compared to CQ alone, KY02111 was still able to markedly reduce SQS levels. Collectively, these observations indicate that KY02111-induced degradation of SQS requires functioning proteasomes, but is not dependent on an E3 ligase which requires activation by neddylation, such as those in the Cullin RING ligase family.[29] Having established a correlation between KY02111-treatment, SQS-binding and proteasomal degradation, we wanted to further investigate and understand how KY02111 precisely mediated this process.

**Figure 4:**
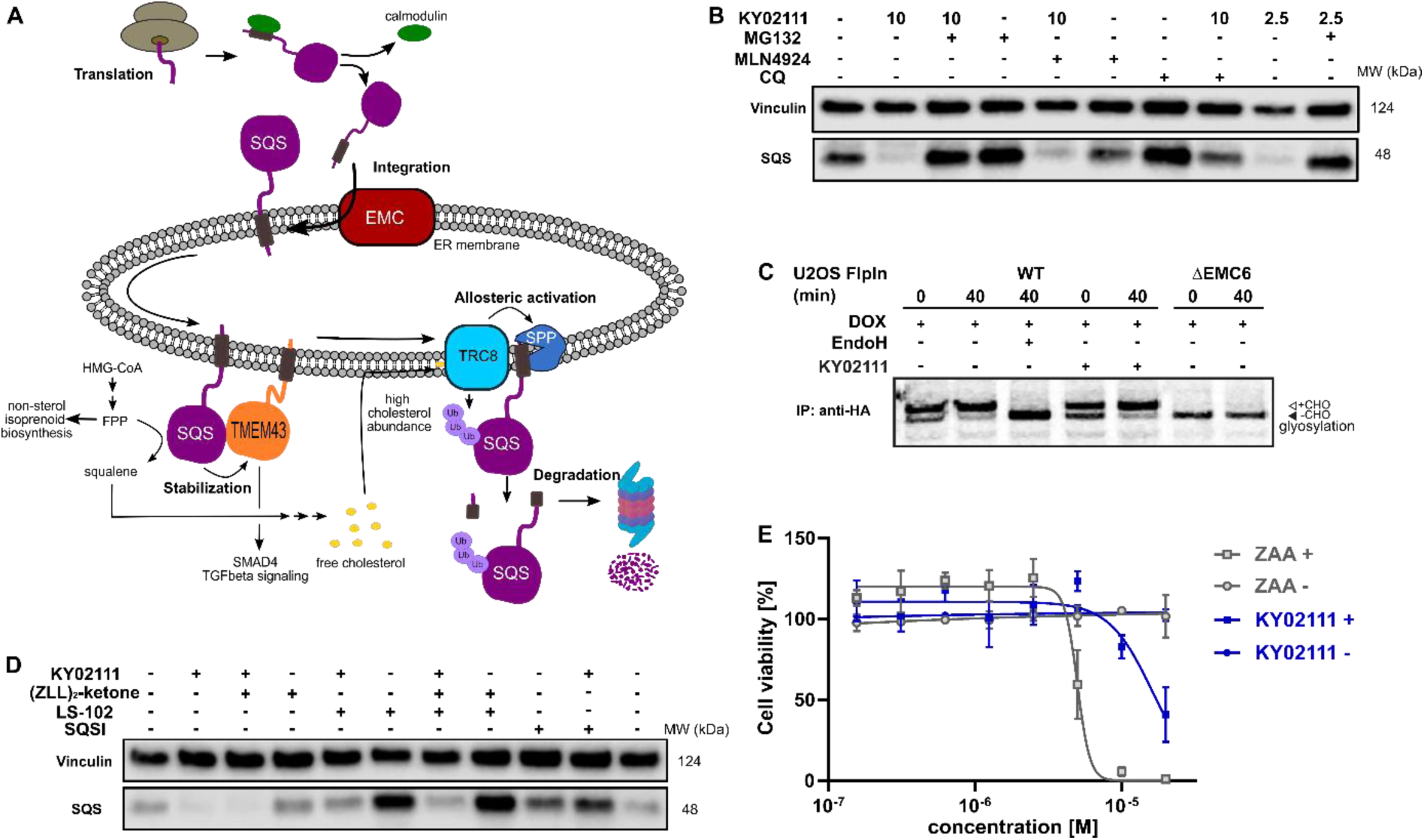
Characterization and evaluation of KY02111-mediated SQS degradation. **A**) “Lifecycle” of SQS based on literature reports. **B)** KY02111-induced SQS reduction is proteasome, and to a lesser degree lysosome, dependent. HeLa cells were pre-incubated with either MG132 (20 µM) or chloroquine (CQ, 20 µM) or MLN4924 (5 µM) for 2 h prior to addition of KY02111 at indicated concentrations (n = 2). **C)** ^35^S-Met/Cys pulse-chase insertion assay of HA:SQS_opsin_ inducibly expressed in U2OS Flp-In^TM^ T-Rex^TM^ WT and ΔEMC6 cells. HA:SQS_opsin_ expression was induced by the pretreatment and continuous presence of doxycycline (DOX, 10 ng/mL, 18 h) throughout the assay. Where indicated, KY02111 (10 µM) was also continuously present during the assay but only after 1 hr pretreatment. Radiolabelled HA:SQS_opsin_ from each timepoint (0 and 40 min) was immunoprecipitated and separated by SDS-PAGE. Where indicated, WT eluate was treated with EndoH_f_ to confirm opsin glycosylation. Core glycosylated (+CHO) and unglycosylated (-CHO) HA:SQS_opsin_ are shown. **D)** Co-incubation of KY02111 with tool compounds inhibiting natural SQS degradation. HeLa cells were co-incubated with KY02111 (10 µM) and either (ZLL)_2_-ketone (10 µM), LS-102 (10 µM) or SQSI (1 µM) for 18 h (n = 3). E) Mild cholesterol auxotrophy after treatment of HeLa cells with KY02111 (20 µM) in the presence of MBCD (3 mM) (n = 3, mean ± SD of one biological replicate shown). **F)** Co-treatment of HeLa cells with KY02111 (10 µM) and CB-5083 (5 µM) does not rescue SQS levels. Hela cells were co-incubated with both compounds for 8 h (n = 3).

### KY02111 does not interfere with SQS insertion into the ER membrane

To become functionally active, SQS must be inserted into the ER membrane via its C-terminal transmembrane domain. Insertion of the SQS “tail anchor” occurs post-translationally and is facilitated by the multi-subunit assembly known as ER membrane protein complex (EMC, Fig. 4A).[30] Without the EMC, SQS tail anchor insertion into the ER membrane is inefficient and rather than accumulating in the cytoplasm, SQS is degraded by proteasomes in a process differing from the canonical ERAD pathway.[23] Given this, we hypothesized that the KY02111-SQS interaction might inhibit its tail anchor insertion into the ER by indirectly preventing it from accessing the EMC (Fig. 4A).

This MoA is consistent with our earlier *in vitro* binding studies, which showed that KY02111 can interact with a recombinant, soluble form of SQS lacking its tail anchor (aa 31-370). Moreover, polymerization triggered by small molecules has been described previously.[31,32] To test this, we performed radiolabeling pulse-chase insertion assays using a doxycycline-inducible (DOX) HA-tagged SQS-construct containing a C-terminal opsin tag after the tail anchor. The opsin tag encodes a sequence that is glycosylated only upon exposure to the ER lumen.[23,33] After 40 min, a shift to the glycosylated form of SQS can be observed in wild-type cells which is sensitive to the deglycosidase EndoH (Fig 4C). Without the EMC (ΔEMC6), SQS remains unglycosylated throughout the chase indicating insertion failure. Pre-treatment (1hr) with KY02111 and its inclusion throughout did not compromise insertion of the SQS tail anchor and resembled untreated cells. This indicates that KY02111 did not cause aggregation and therefore was not acting by attenuating or disrupting normal SQS biogenesis.

Additionally, we investigated whether KY02111 could cause SQS to aggregate and therefore performed *in vitro* differential light scanning (DLS) experiments as well as size exclusion chromatography in the presence of KY02111 (SI Fig. 9). We determined the diameter (d) of recombinant SQS to be 4-6 nm, which is in agreement with protein sizes generally determined via DLS, but could not detect any change in diameter upon incubation with KY02111 (SI Fig. 9 A/B).

### Partial chemical SQS knockdown does lead to mild cholesterol auxotrophy

We found that the viability of HeLa cells treated with KY02111 was indistinguishable from DMSO-treated when cells were grown in standard FBS-containing media for up to 48 h (SI Fig. 7A/B). However, it has previously been reported that SQS depletion resulting from loss of the EMC led to a cholesterol auxotrophic effect and increased cell death upon cholesterol depletion when using either lipoprotein deficient serum (LPDS) or methyl β-cyclodextrin (MBCD). To test whether a loss of SQS induced by KY02111 (∼65-70%) would compromise cell viability similarly, we grew HeLa cells in FBS-containing media supplemented with MBCD for 72 h and treated with different concentrations of either KY02111 or the potent active site inhibitor ZAA (Fig. 4E). We determined the non-toxic concentration of MBCD to be 3 mM for HeLa cells (SI Fig. 10 A/B). As reported previously, high concentrations of ZAA (5 and 10 µM) were sufficient to completely abolish cell viability. KY02111 also reduced cell viability at high concentrations (20 µM) although not to the same degree as ZAA. When cells were grown in standard conditions (DMEM 10% FBS), neither ZAA nor KY02111 adversely affected cell viability at 72 h. This indicates that reducing levels of SQS through chemically induced degradation partially phenocopies catalytic inhibition, compromising cholesterol biosynthesis through the mevalonate pathway.

### Conventional SQS degradation pathways are not enhanced by KY02111

As SQS is a critical component in the mevalonate pathway downstream of FPP, its abundance is tightly controlled. Rapid degradation is facilitated by either signal peptide peptidase (SPP), which cleaves the TMD of SQS at high cholesterol levels sensed by the ER-resident ubiquitin ligase (E3) TRC8, or by HRD1, another ER-E3 associated with ERAD (Fig. 4A).[2] To investigate whether KY02111 treatment might be enhancing natural degradation of SQS degradation, we co-incubated HeLa cells with tool compounds targeting these pathways (Fig. 4D). We found that neither the SPP inhibitor (ZLL)_2_-ketone, the HRD1 inhibitor LS-102, nor the combination of both could prevent a reduction in SQS band intensity in the presence of KY02111, when compared to the individually treated samples. Interestingly, LS-102 treatment led to a marked accumulation of SQS, which suggests that HRD1 is an important factor facilitating basal turnover. When LS-102 treatment was combined with KY02111, SQS no longer accumulated. Yet, the levels were still greater than that of KY02111 alone, suggesting that KY02111-induced SQS degradation results from a combination effect of compound treatment and regular turnover. Collectively, these data support a mode of action for KY02111 that is independent of the natural SQS degradation processes.

Since our earlier FP measurements showed competition between active-site inhibitor SQSI and the KY02111-based probe **27,** and PROTAC **18** vs probe **26** (Fig. 2B), we next wanted to test if KY02111-mediated degradation can be blocked by SQSI. Surprisingly, SQSI was able to rescue SQS levels in the presence of KY02111, even when incubated at a 10-fold lower concentration (1 µM), reflecting the stronger binding affinity of SQSI compared to KY02111. Data from Takemoto *et al.* suggested that KY02111 does not inhibit the catalytic activity of SQS and therefore, binding should occur outside of the active site.[3] We confirmed this using a similar SREBP reporter gene assay with KY02111, finding that SREBP target genes are not activated in response to KY02111 treatment (SI Fig. 6).[34] SREBP is the transcriptional regulator of SQS and cholesterol biosynthesis in general, its activity can be correlated to impaired cholesterol synthesis and SQS activity. [35,36] Reconsidering the FP and cellular competition experiments, our data suggests that KY02111 binds close to the SQS catalytic site without inhibiting the enzymatic activity. To fully elucidate the binding interaction between KY02111 and SQS we attempted to generate a crystal structure of the compound-protein complex, but those efforts have been unsuccessful so far.

### KY02111 is a selective degrader of SQS

To further our investigation into the mechanism of degradation induced by KY02111, we performed global proteomic studies using isobaric tandem mass tag labeling (TMT, 16-plex) coupled to mass spectrometry analysis (MS). To this end, we incubated HeLa cells with either KY02111 (10 µM, Fig. 5A), compound **18** (5 µM, Fig. 5B) and SQSI (1 µM, Fig. 5C) for 18 h. The concentrations were chosen with our earlier WB results in mind to ensure robust detection of changes in SQS protein level. We specifically sought to detect changes protein levels of enzymes within cholesterol/lipid metabolism, ERAD or EMC clients as well as TMEM43, a potential PPI interaction-partner of SQS.

**Figure 5:**
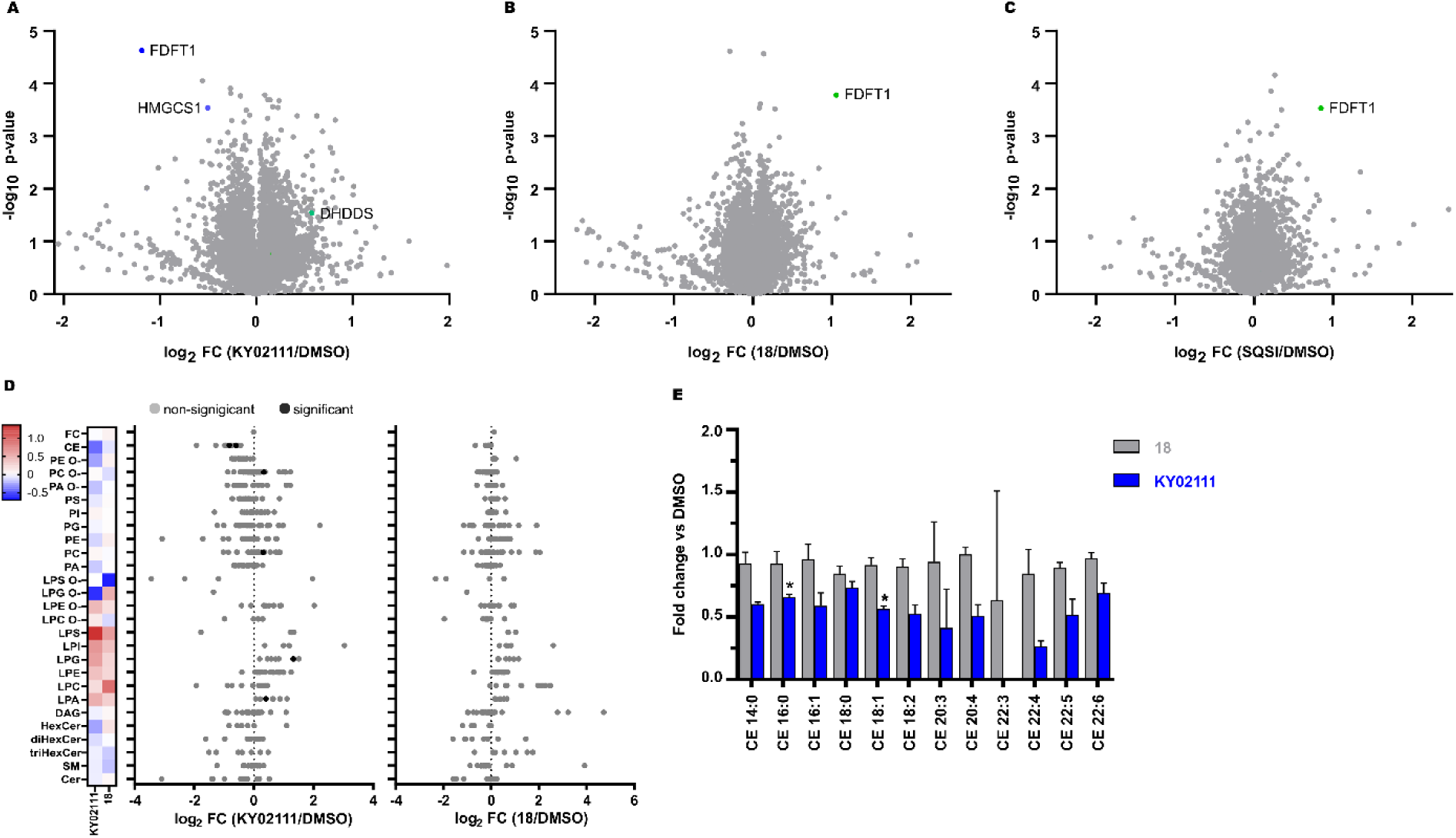
Global proteomic and lipidomic analysis of KY02111– and **18**-treatment in HeLa cells. **A/B/C**) Proteomic profile of HeLa cells treated with KY02111 (10 µM, A), PROTAC **18** (5 µM, B) or SQSI (1 µM, C) for 18 h. The soluble fraction of the lysates was labeled with 16-plex TMT labels and subjected to MS/MS analysis. The analysed data was plotted as –log_10_ p-value vs log_2_ FC (FC = fold change, n = 3). Please see extended dataset 1 for complete proteomics data and analysis; **D)** Changes in lipidomic species of HeLa cells treated with KY02111 (10 µM) or **18** (5 µM) for 18 h. Significantly altered species are highlighted, Holm-Šídák correction was used to determine adjusted p-values (n = 3, p < 0.005). **E)** Cholesteryl ester (CE) class overview of HeLa cells treated with KY02111 (10 µM) or PROTAC **18** (5 µM) for 18 h. Significantly altered species are highlighted with * (n = 3, p < 0.005). Please see extended dataset 2 for complete lipidomics data and analysis.

Importantly, quantitative proteomic analyses were able to reproduce our WB results, identifying SQS (= FDFT1) as significantly reduced (KY02111, log_2_ FC = –1.19, –log_10_ p-value = 4.63) or enriched (**18**, log_2_ FC = 1.05, –log_10_ p-value = 3.78; SQSI, log_2_ FC = 0.85, –log_10_ p-value = 3.53) in response to their respective treatments (extended dataset 1). Surprisingly, we did not detect a general increase in cholesterol biosynthesis enzyme protein levels as a result of SQS degradation, stabilization or inhibition. It should be noted that we could not detect HMGR in all of our MS data sets. Loss of cholesterol biosynthetic capacity might be compensated for by an increase in cholesterol uptake from the medium (standard growth conditions, 10 % FBS) or by utilizing stored cholesteryl esters. KY02111 treatment only reduces SQS protein levels to 30– 35% and the remaining fraction may be sufficient to supply cells with cholesterol if needed. We speculated that reducing SQS levels by nearly two-thirds could lead to accumulation of its substrate FPP, accompanied by changes to products within the non-sterol isoprenoid pathway like dolichol, ubiquinone and general protein prenylation.[37,38] We believe this is indicated by a significant decrease in HMG-CoA synthase (log_2_ FC = –0.51, –log_10_ p-value = 3.54) protein levels, which is consistent with our SREBP reporter gene assay data. Here KY02111 led to a reduction of a luciferase reporter signal connected to a HMGCS promotor region (SI Fig. 6). Moreover this was accompanied by an increase of dehydrodolichyl diphosphate synthase (DHDDS, log_2_ FC = 0.57, –log_10_ p-value = 1.53), which uses FPP as a direct substrate.[39,40]

This hypothesis is also supported by studies of mice lacking SQS in the liver,[4] which show significant increase in FPP levels.[15] Since Uesugi *et al.* reported inhibition of the SQS-TMEM43 PPI by KY02111 and showed that siRNA-SQS KD led to a decrease of TMEM43, we also anticipated a reduction of TMEM43 in cells with chemically reduced SQS levels. To our surprise we did not detect any significant level changes for TMEM43 (log_2_FC = 0.13, –log_10_ p-value = 0.78) in KY02111-treated samples, which we attributed to numerous differences in the experimental setup.

### Cholesteryl ester levels are downregulated as a result of SQS degradation

To determine whether KY02111-induced reduction of SQS protein level is able to attenuate cholesterol biosynthesis and concomitantly decrease cholesterol content, we performed a quantitative shotgun lipidomics analysis of HeLa cells, which covered 27 lipid classes and over 446 species (Fig. 5D, SI Fig. 11 and extended dataset 2). Most notably, we detected a significant reduction in cholesteryl ester levels of CE 16:0 and 18:1 (Fig. 5E) in cells treated with KY02111. Together, the reduction of these two species alone accounted for an overall change larger than 1 mol% of the whole lipidome. Additional CE species were also affected, albeit not to a statistically significant level. However, we observed an overall trend in reduction of CE content upon SQS degradation compared to cells treated with DMSO or compound **18** by about 40%. Generally, compound **18** did not have any significant effect on cellular lipid levels (Fig. 5D). CE’s are the storage form of cholesterol inside cells and are usually produced from acyl-CoA by Sterol-O-acyltransferase (SOAT1).[41] As unesterified cholesterol levels are not altered, we believe that the cells compensate for a lower amount of SQS protein and activity not by upregulating cholesterol biosynthesis but by utilizing available cholesteryl esters. This could either occur by hydrolyzing and liberating stored cholesteryl esters or by downregulating ester production. Phopsphatidylcholines (PC 32:1) were also altered significantly, which could be increased (≥ 1 mol%) to compensate for a greater excess of acyl-CoA, since it is not used to produce CE’s. We also detected a significant increase in lysophosphatidic acid (LPA 18:0) and lysophosphatidyl glycerol (LPG 16:0). Whether these changes directly relate to KY02111-induced SQS degradation has yet to be determined.

## Discussion

SQS is a catalytic enzyme dictating the flow of FPP and a long-investigated drug target. Herein we sought to identify chemical strategies to degrade SQS utilizing the TPD design principle. Our efforts led us to identify the known SQS ligand KY02111 as a degrader, whereas bifunctional molecules containing E3 ligase-recruiting ligands generally led to an increase in SQS levels. To investigate this phenomenon we used compound **18** as a model which does not bind the active site, and found that these molecules likely act by shielding the protein from its natural degradation.

It is intriguing that SQS degradation in response to KY02111 treatment was not observed previously.[3] However, this may be attributable to differences in the experimental setup. We monitored endogenous SQS expressed in HeLa cells, whereas Takemoto *et al.* used FLAG-tagged SQS constitutively overexpressed in HEK293 cells.[42] Since the ability for KY02111 to degrade SQS plateaus at 70%, the constant overexpression of FLAG-SQS may mask the underlying degradation. Similar reasoning can be applied when comparing the effects of siRNA KD of SQS (48 h, 100%, Takemoto *et al.*) to chemical KD with KY02111 (18 h, 70%, this work). Takemoto *et al.* found TMEM43 to be destabilized when SQS was knocked down, which phenocopies a S358L TMEM43 mutation in arrhythmogenic right ventricular cardiomyopathy (ARVC). In our case, the 30% of SQS protein remaining is likely sufficient to stabilize the TMEM43 protein after 18 h, which could explain that we did not observe a decrease in TMEM43 abundance in our proteomic analysis. If KY02111 directly inhibits the SQS-TMEM43 PPI, as was proposed by co-immunoprecipitation, or if SQS degradation indirectly leads to a decrease of TMEM43 remains unclear. To answer this question and to unequivocally determine the binding mode of KY02111 and derived molecules to SQS, we initiated attempts to generate a crystal structure of the compounds bound to the SQS protein but have been unsuccessful so far.

Our data suggests that KY02111 binds in, or at least very close to the active site of SQS without altering its catalytic activity. FP experiments showed that a fluorescent 5-TAMRA labeled probe based on a KY02111 analogue (**27**) was competed off by SQSI, a known active site binder of SQS. The finding that SQS degradation was rescued when co-incubated with SQSI is consistent with our current hypothesis (Fig. 4D). We also observed a general trend where attachment of a linker and a degrader moiety led to higher binding affinity for SQS, possibly by altering the KY02111-scaffold binding mode. This could be the case for compound **18**, a designed “PROTAC” with a VHL recruiting modality, which we later identified as a stabilizer of SQS. Compound **18** showed the highest binding affinity for SQS when competed against both fluorescent probes (Fig. 2) as well as the highest degree of stabilization in a DSF thermal shift assay (SI Fig. 4). We rationalized that this high affinity is the basis for the stabilization SQS by compound **18** in a concentration-dependent manner. SQS stabilizers might have applications in settings where SQS is expressed in low amounts. Recently, Coman *et al.*[5] identified three human patients harboring pathogenic FDFT1 gene variants resulting in dysregulated splicing and transcription and ultimately lower or no expression of SQS. We believe this rare disease setting could be a potential application for molecular stabilizers like compound **18**, which can lead to increased cellular concentrations of SQS.

We established that KY02111-mediated SQS degradation is dose-, time– and proteasome-dependent and explored several possible MoA’s of this phenotype. We were most intrigued by the idea that KY02111 might interfere with the insertion of SQS into the ER membrane, since we found several characteristic similarities between our chemical KD and an ΔEMC cell line developed and investigated by Volkmar *et al*.[23] When cells fail to insert SQS into the ER membrane, the protein is degraded via the proteasome in manner that appears independent of ERAD. As in ΔEMC cells, our chemical KD causes cells to utilize CE’s under normal growth conditions (Fig. 5D/E), as less free cholesterol can be synthesized. When grown in cholesterol depleted medium (Fig. 4E), we were able to observe a mild auxotrophic effect at higher concentrations. However, we found that KY02111 did not lead to aggregation of SQS, and insertion into the ER membrane was not affected (Fig. 4B). Additional DLS and SEC experiments did not suggest formation of aggregates of SQS upon KY02111-treatment *in vitro* (SI Fig. 9). Even though not a common MoA, recent examples show that small molecule-induced aggregation is possible.[31,32] Ultimately we were not able to fully decipher the fundamental mechanism of SQS-degradation by KY02111.

It is important to note that KY02111 was originally reported as a Wnt-pathway inhibitor and frequently used for the induction of hemi pluripotent stem cells (hPSC) in combination with other small molecules.[43–45] Importantly, in their work Takemoto *et al.* demonstrate that the Wnt activity of KY02111 may be an artifact, where KY02111 co-precipitates with the Wnt activator (2’Z,3’E)-6-bromoindirubin-3’-oxime (BIO). BIO is a known and widely used Wnt activator containing two linked indole fragments.[46] This further led to the conclusion that KY02111 does not have a direct effect on endogenous Wnt-signaling but acts within TGFβ signaling by destabilizing TMEM43. So far, it remains unresolved if SQS degradation and its potential effect on TGFβ signaling might be the underlying mechanism behind KY02111’s action in hSPCs. Early results[15] also indicate additional SQS functions in development. Homozygous SQS knockout mice are embryonic lethal, even when supplemented with squalene and cholesterol. The afore-mentioned study by Coman *et al.* supports this notion. We have shown that even after 18 h treatment, SQS degradation by KY02111 leads to a decrease in CE’s. Hence it would be premature to dismiss potential connections between the Wnt-inhibition activity annotated for KY02111 and SQS degradation.

To conclude, some general considerations on the design and development of optimized PROTACs/degraders for either SQS or any sterol biosynthetic enzymes are warranted. Cholesterol biosynthesis is a tightly regulated process, where overall activity depends on cholesterol homeostasis and localization inside the cell, as well as extracellular availability, which makes specific perturbations a challenging task. A recent example of this is the development of atorvastatin-based PROTACs for HMGR by Li *et al.*, who chose SRD15 cells as their *in vitro* model system.[47] These cells are lacking Insig 1 and 2, which normally facilitate sterol-regulated degradation of HMGR.[48] Thereby, natural degradation and concomitant accumulation of HMGR in response to statins was eliminated, simplifying evaluation of potential degradation. Yet when regular Huh7 cells were treated with the optimized PROTACs, the protein levels were close to the DMSO control. Furthermore, HMGR-PROTACs led to increased transcription of HMGR mRNA. Similarly, potent degradation of SQS might lead to feedback activation of SREBP and concomitant upregulation, which would need to be competed and overcome by the degrader.[11,37] As we were able to show that overall cholesterol levels can already be lowered by moderate SQS degradation, one might consider-the potential benefits of this over complete chemical knockdown. Regardless, the exact cellular responses to absolute SQS degradation via a chemical probe remain to be investigated and our discovery of KY02111 as a SQS degrader lays the groundwork towards this goal.

Ultimately our study highlights that SQS degradation is possible despite multiple biological challenges including moderate protein turnover (t_1/2_ ≈ 5h) and potential feedback regulation within the cholesterol biosynthesis pathway. Most importantly, we were able to show that SQS degraders have overall cholesterol lowering abilities and consequently believe there is large potential for further optimized probes, which fall outside of the generally accepted PROTAC paradigm. Compounds designed following traditional TPD-principles generally seem to act as stabilizers of SQS by shielding the enzyme from its natural degradation. Though not designed for this purpose, these previously unknown molecules could serve as probes to further study SQS biology in rare genetic diseases where correct SQS expression is markedly reduced.

## Supporting information

Extended Dataset 1 Proteomics

Extended Dataset 2 Lipidomics

## Acknowledgements

We thank the Novo Nordisk Foundation (NNF19OC0058183 and NNF21OC0067188) and the Independent Research Fund Denmark (9041-00241B) for funding to LL. JCC is supported by a Senior Cancer Research Fellowship from Cancer Research UK (C68569/A29217).

## Conflict of interest

The authors have no conflicts to declare.

## Methods

### Key Resource Table

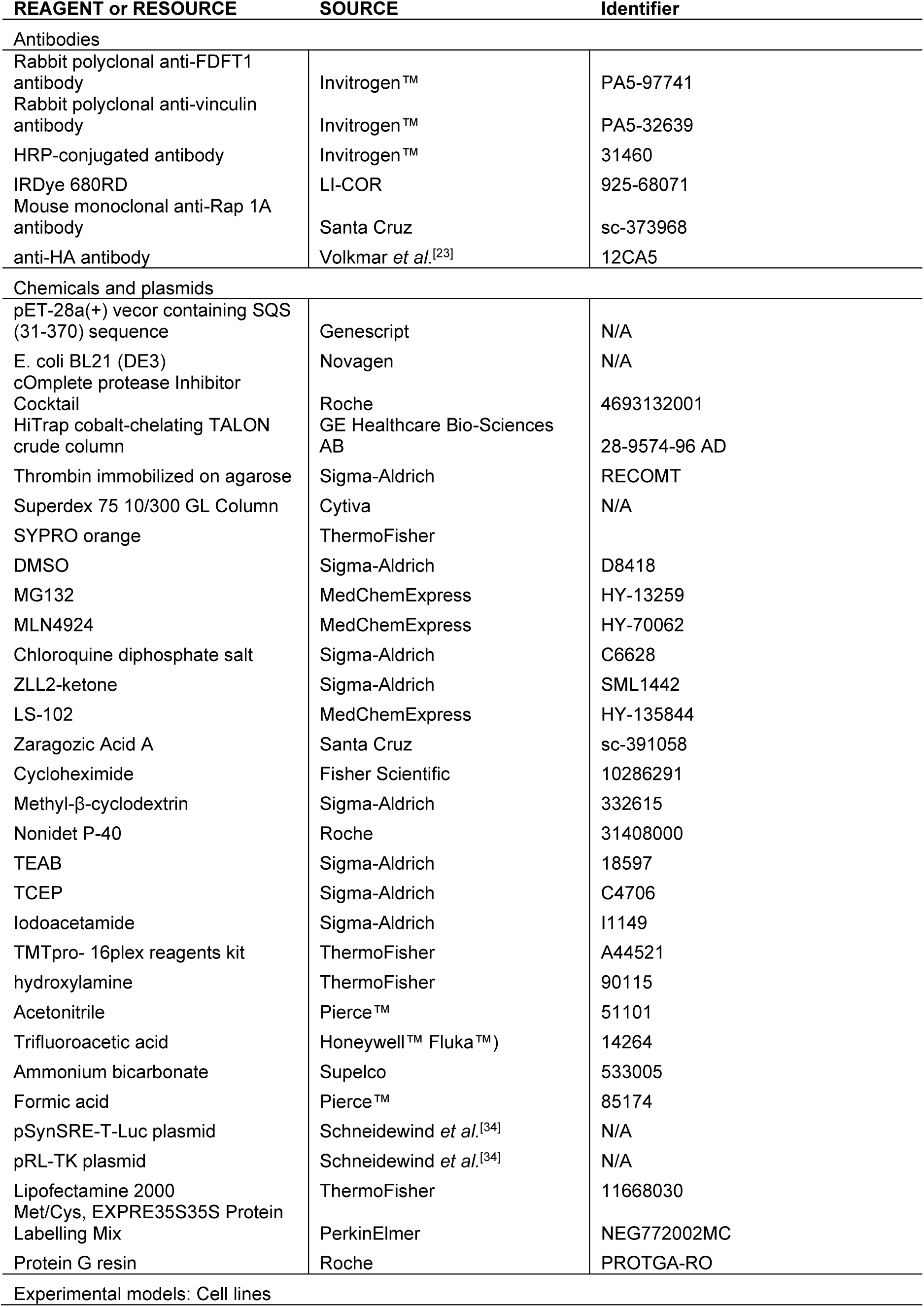
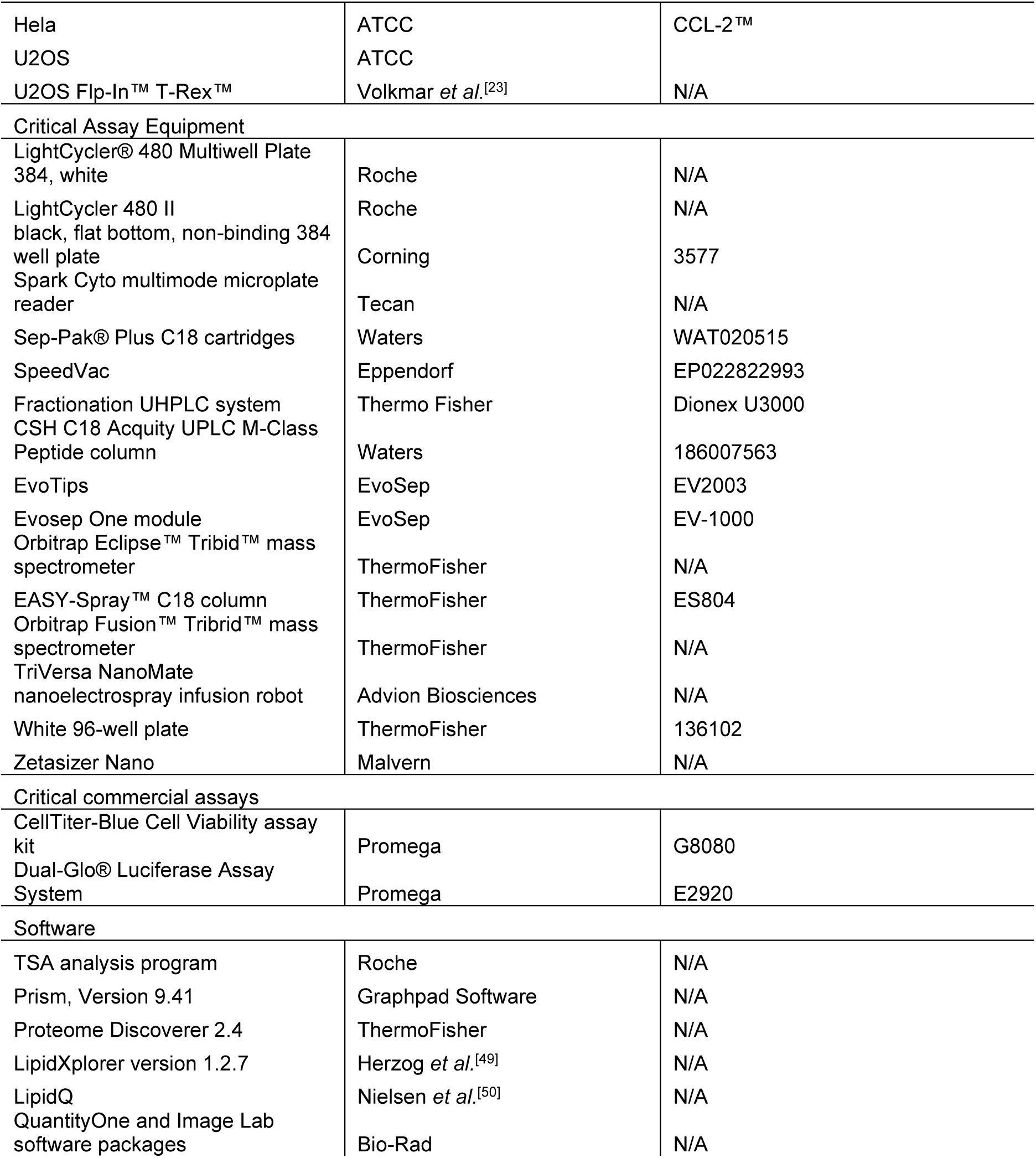

The

### Chemical Synthesis

The chemical synthesis of the SQS degrader library as well as tested compounds is described in the supporting information. Tool compounds were bought from the respective suppliers indicated in the key resource table and stored according to the manufacturer’s instructions.

### Protein expression, purification and thrombin cleavage

The expression of the doubly truncated SQS protein was conducted following a reported procedure by Song *et al*.[51] with minor adjustments. In short, a bacterial expression vector pET-28a(+) containing a sequence corresponding to AA 31-370 of the SQS enzyme was bought from Genescript. The expression vector was used to transform *E. coli* BL21 (DE3)RP strain (Novagen) for overexpression. The resulting construct contained a *N*-terminal His_6_-tag with a thrombin cleavage site. Bacteria expressing the constructs were cultured in LB medium supplemented with kanamycin (30 *μ*g/mL) at 37 °C, until the cells reached an OD of 0.7 at 600 nm, and were then induced at 37 °C for 4 h by incubation with 0.1 mM isopropyl-1-thio-*D*– galactopyranoside.

Cells were harvested by centrifugation (10 min, 4000 rpm) and resuspended in 40 mL of lysis/elution buffer (20 mM NaH_2_PO_4_, pH 7.4, 10 mM CHAPS, 2 mM MgCl_2_, 10% glycerol, 2 mM DTT, 500 mM NaCl and a protease inhibitor cocktail (Roche), disrupted by sonication, and centrifuged at 16000 rpm for 30 min. The supernatant was then applied to a HiTrap cobalt-chelating TALON crude column (GE Healthcare Bio-Sciences AB). Enzyme purification was performed according to the manufacturer’s instructions using an Äkta FPLC system. Unbound protein was washed off with 5 mM imidazole. Then the His_6_-SQS was eluted with 150 mM imidazole. Purity was confirmed by SDS-PAGE electrophoresis. Fractions containing the pure enzyme were pooled and dialyzed against buffer A (25 mM sodium phosphate, pH 7.4, 20 mM NaCl, 2 mM DTT, 1 mM EDTA, 10% glycerol, 10% methanol), concentrated (2.37 mg/mL) and then stored at –80 °C for *in vitro* assays.

For cleavage of the His_6_-tag, thrombin immobilized on agarose (RECOMT) was purchased from Sigma-Aldrich and used following the manufacturers instructions. Shortly, His_6_-*h*SQS was transferred into a thrombin-cleavage buffer (50 mM Tris-HCl, pH 8.0, 10 mM CaCl_2_) and the His_6_-tag was cleaved after incubation at room temperature overnight. After centrifugation recovery, the cleaved protein was subjected to a second HiTrap cobalt-chelating TALON crude column to separate the cleaved His_6_-tag. Successful cleavage was confirmed by SDS-PAGE analysis. The protein was concentrated using Amicon Ultra Centrifugal Filters (10.000 MWCO) and then stored at –80 °C. Before crystallization attempts, the thawed protein was subjected to a SEC column and transferred into the crystallization buffer (20 mM Tris-HCl, pH 7.3, 2 mM MgCl_2_, 0.1 mM EDTA, 1 mM DTT, final protein concentration > 8mg/mL).

### Differential scanning fluorimetry

Differential scanning fluorimetry experiments were performed in a buffer composed of 50 mM HEPES pH 7.5 and 5 mM MgCl_2_ in Milli-Q water. His_6_-*h*SQS was diluted to a concentration of 0.5 mg/mL and 8 µL/well of the resulting solution were transferred to a 384-well plate (LightCycler® 480 Multiwell Plate 384, white). Subsequently, SYPRO orange (Thermo Fisher) was added (final concentration 1× SYPRO orange) followed by the addition of the tested compounds at the indicated concentrations (50, 25, 12.5 µM) to a total final volume of 10 μL. The fluorescence intensity was measured in a Roche LightCycler 480 II with an initial incubation at 30 °C for 1 minute followed by acquisition steps of 0.1 °C up to 95 °C with incubation for 1 second at each step. Melting temperatures were calculated with the Roche TSA analysis program. Exemplary melting curves were plotted using Prism (Graphpad Software, Inc. Version 9.41).

### Fluorescence Polarization

Fluorescence polarization experiments were performed at room temperature in a buffer composed of 50 mM HEPES pH 7.5, 5 mM MgCl_2_ in a final volume of 30 μl in black, flat-bottom, non-binding 384-well plates (Corning). For competition experiments 20 nM 5-TAMRA-SQSI (or 5 nM 5-TAMRA-KY02111) was mixed with 200 nM of His_6_-SQS and incubated for 30 minutes. Meanwhile, a 10-point 1:1 dilution series for the tested compounds was performed from 20 µM in the assay buffer. After 30 minutes, the screening compounds were added at the indicated conentrations and the plate was incubated for 45 minutes. The fluorescence polarization signal was measured using a Spark Cyto multimode microplate reader (Tecan) with filters set at 530 ± 25 nm for excitation and at 590 ± 20 nm for emission. The data was analyzed using GraphPad Prism 9. Curves were fitted by non-linear regression to allow the determination of IC_50_ values with GraphPad Prism 9. The *k_d_* values for the FP probes were determined by curve fitting of a 1:3 protein titration series against 20 nM/5 nM of the respective FP probes using nonlinear regression. The determined values were then used to calculate the *k_i_* values for the respective compounds by applying equations (1) and (2)[52]:

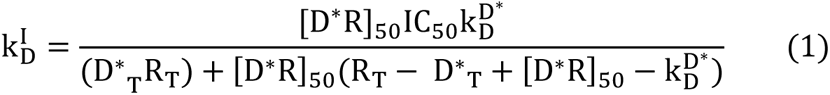

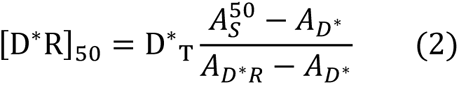

Where R_T_ is the total amount of protein, D*_T_ the total amount of the FP probe and k_D_^D*^ the *k_d_* value for the respective FP probe previously determined via Prism. A_D*R_ is the anisotropy for the protein + fluorophore control well whereas A_D*_ is the anisotropy for the fluorophore control well. A^50^_S_ is the anisotropy at IC_50_ and can be calculated using the values generated by nonlinear regression (3):

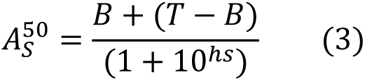

Where B is the bottom value and T is the top value. hs is the hill slope.

### Cell culture and compound incubation

HeLa or U2OS cells were purchased from ATCC and cultured in DMEM medium supplemented with 10% FBS and 1% Penicillin/Streptomycin at 37°C in 5% CO_2_ atmosphere. For compound testing, cells where harvested from a T175 flask (confluence 90%, passage 3-20) by trypsination and diluted to 2*10^5 cells/mL, and finally transferred to 6-well plate with a final cell count of 4*10^5 cells per well. After attachment overnight, the medium was exchanged and compounds were added to the final indicated concentrations followed by incubation for the indicated time points. For tool compound co-incubation experiments, MG132, MLN4294 and Chloroquine were pre-incubated for 2 hours, Cycloheximide was pre-incubated for 1 hour, and (ZLL)_2_ketone and LS-102 were simultaneously incubated with the tested compounds. Upon completed incubation, the medium was removed and the cells were washed with 2 mL of ice-cold PBS before being lysed with 150 µL SDS lysis buffer (100 mM Tris–HCl, pH 6.8, 4% (w/v) SDS, 20% (v/v) glycerol). The cell lysates were collected, sonicated (3 x 20 cycles (1/second)) and the total protein concentration was determined using a nanodrop (A_280_). Finally, cell lysates were flash frozen in liquid N_2_ and stored at –80 °C.

### SDS-PAGE and Western Blot analysis

Cell lysates were thawed on ice and diluted with 2× loading buffer (100 mM Tris–HCl, pH 6.8, 4% (w/v) SDS, 20% (v/v) glycerol) to a final concentration of 4 mg/mL (max, or the highest possible concentration in a lysate set). To each sample, 2× loading buffer containing bromophenol blue (0.2% (w/v)) and DTT (200 mM) was added to a final ratio of 2:1 (v/v, sample:loading buffer) and heated at 95°C for 5 min for complete protein denaturation. Samples were then loaded onto 4–15% Mini-PROTEAN TGX Protein Gels (4561086, Bio-Rad) and run at 120 V for 1 hour. Protein transfer from gels to polyvinylidene difluoride (PVDF) membranes was carried out using the Trans-Blot Turbo Transfer Kit (1704274, Bio-Rad) according to the manufacturer’s instructions. Membranes were subsequently incubated with the blocking solution (5% Skim Milk Powder (10651135, Fisher Scientific) added to tris-buffered saline (TBS) solution (150 mM NaCl (S7653, Sigma-Aldrich), 50 mM Tris pH 7.6, 0.1% Triton™ X-100 (T8787, Sigma-Aldrich)) before the incubation overnight at 4 °C with the primary antibodies for SQS and vinculin. The following day membranes were incubated for 1 h with the HRP-conjugated secondary antibody at room temperature. Successively, the SuperSignal™West Femto Chemiluminescent substrate (10391544, Thermo Scientific™) was added to the membranes before imaging with a Odyssey Fc imaging system (Licor). Image acquisition was performed with Image Studio (LI-COR), quantification of the chemiluminescent intensities of the bands were performed with Empiria studio (LI-COR version 1.3).

### Cycloheximide-chase assay

HeLa cells were cultured as described above. One day before compound incubation, 2 mL of a 2.0×10^5 cells/mL solution were added to single cell culture dishes (EP0030700112, Sigma-Aldrich) and incubated overnight. The next day, the medium was refreshed and cells were incubated with either Cycloheximide (100 µg/mL, 355 µM) alone or together with **18** (3 µM). After the indicated time points, cells were lysed by addition of 100 µL SDS-lysis buffer and analyzed via western blotting.

### Cell viability assay ± MBCD

HeLa cells were harvested from a T175 flask (confluence 90%) by trypsination and diluted to 3*10^4 cells/mL with medium. 100 µL/well of the diluted cell solution were transferred into a 96 well plate (Nunc™ Edge™, clear, 167425, Thermo Fisher) and cells were allowed to attach overnight (37 °C, 5% CO_2_). The next day, a dilution series starting from the highest concentration was prepared for the tested compounds in DMEM medium (10 % FBS, 1 % PS), or in MBCD-containing DMEM medium (10 % FBS, 1 % PS, 4 mM MBCD). The medium was removed from the cells and 100 µL of the medium including the compounds at indicated concentrations was added (technical triplicates). Alternatively, the cells were washed with pre-warmed PBS before addition of MBCD-containing medium. The cells were incubated for the indicated time points, after which the wells were checked for potential compound precipitation. Cell viability was determined by using the Cell Titer Blue Cell Viability assay kit (G8080, Promega) according to the manufacturer’s instructions. Shortly, 10 µL of equilibrated CellTiter-Blue reagent were added per well and the plates were incubated for 60 min at 37 °C. The fluorescence intensity signal was measured using a Spark Cyto multimode microplate reader (Tecan) with filters set at 560 ± 20 nm for excitation and at 590 ± 20 nm for emission. The background (well without cells) was subtracted from the sample wells and the signals were normalized on DMSO controls. The obtained values were plotted in GraphPad Prism 9.

### PROTEOMIC sample preparation

HeLa cells were cultured and incubated as described above. Here, cells were lysed by addition of 200 µL Nonidet P-40 (NP-40) lysis buffer (0.4 % NP-40 in PBS) followed by three consecutive freeze-thaw cycles in liquid N_2_ and storage at –80 °C. The next day, the lysates were thawed on ice and ultracentifuged for 25 minutes at 4 °C and 30000 rpm. The concentration of the supernatant was determined by using the DC assay kit (5000112, Bio-Rad) according to the manufacturer’s instructions and the samples were diluted to a final concentration of 2 mg/mL in 75 µL NP-40 lysis buffer. For protein reduction and alkylation, 75 µL of a 100 mM TEAB (1:1, 18597, Sigma-Aldrich, prepared in pre-filtered MilliQ water) solution was added to the samples followed by the addition of 7.5 µL 200 mM TCEP (1:10, C4706, Sigma-Aldrich, prepared in solution of 140 µL TCEP 0.5 M, 140 µL MilliQ water, 70 µL 1 M TEAB). The prepared samples were incubated in a thermoblock at 55 °C for 1 hour and cooled down to room temperature before 7.5 µL of a 375 mM iodoacetamide solution (1:10, I1149, Sigma-Aldrich, prepared in 300 µL of MilliQ water and 75 µL 1 M TEAB) was added followed by incubation in the dark for 30 minutes. Finally, 900 µL of ice-cold acetone was added and each sample was left at –20 °C overnight for protein precipitation. For protein digestion, the thawed samples were centrifuged for 10 minutes at 4 °C and 8000 g. The samples were kept on ice while removing the supernatant and the pellets were dried for 4 hours. After completed drying, 150 µL of a freshly prepared 100 mM TEAB solution were added to solubilize the protein pellets, before the addition of 115 µL of a 0.03 µg/mL trypsin solution (1:80 enzyme:substrate ratio). The samples were incubated at 37 °C overnight at 500 rpm. The next day, the samples were placed in the freezer for 1 minute to stop the digestion.

Half of the sample volume corresponding to 75 µg of peptides were successively labelled with TMTpro 16-plex reagents kit. TMT reaction was allowed for 2 hours and quenched with 5% hydroxylamine (90115, Thermo Fisher Scientific). All the samples were pooled and dried in a SpeedVac (EP022822993, Eppendorf) before desalting with Sep-Pak® Plus C18 cartridges (WAT020515, Waters). The peptides were eluted with 40% and 60% of acetonitrile (51101, Pierce™) in 0.1% of trifluoroacetic acid (TFA) (14264, Honeywell™ Fluka™) and dried before injection of 30 µg to the UHPLC system (Dionex U3000) for high-pH fractionation.

The separation of the peptides was carried out at a constant flowrate of 5 µl min^-1^ on a CSH C18 Acquity UPLC M-Class Peptide column, 130 Å, 1.7 µm, 300 µm x 150 mm (186007563, Waters) using a 100 min linear gradient from 5 to 35% of mobile phase B (acetonitrile) with a subsequent 15 min gradient to 70%, before 5 min re-equilibration with 95% of mobile phase A (5mM ammonium bicarbonate (533005, Supelco), pH 10). 60 time-based fractions were pooled in 30 fractions in the collection plates. Clean-up of the fractions was performed by EvoTip according to the manufacturer’s instructions.

### Global proteomic mass spectrometry (MS) analysis

For MS sample analysis, the EvoTips (EV2003, EvoSep) were loaded on the Evosep One module (EV-1000, EvoSep) coupled to an Orbitrap Eclipse™ Tribid™ mass spectrometer (Thermo Fisher Scientific). The peptides were loaded onto the EASY-Spray™ C18 column, 2 µm, 100 Å, 75 µm x 15 cm (ES804, Thermo Fisher Scientific) using the standard “30 samples per day” Evosep method. The method eluted the peptides with a 44 min gradient ranging from 5% to 90% acetonitrile with 0.1% formic acid (85174, Pierce™). The MS acquisition was performed in data dependent-MS3 with real-time-search (RTS) and a FAIMS interface switching between CVs of −50 V and −70 V with cycle times of 2 s and 1.5 s, respectively. The data dependent acquisition mode was run in a MS1 scan range between 375 and 1500 *m/z* with a resolution of 120000, and a normalized gain control (AGC) *target* of 100%, with a maximum injection time of 50 ms. RF Lens set at 30%. Filtering of the precursors was performed using peptide monoisotopic peak selection (MIPS), including charge states from 2 to 7, dynamic exclusion of 120 s with ±10 ppm tolerance excluding isotopes, and a precursor fit of 70% in a window of 0.7 m/z with an intensity threshold of 5000. Selected precursors for further MS2 analysis were isolated with a window of 0.7 m/z in the quadrupole. The MS2 scan was performed over a range of 200-1400 *m/z*, collecting ions with a maximum injection time of 35 ms and normalized AGC target of 300% MS2 fragmentation was operated with normalized HCD collision energy at 30%. Fragmentation spectra were searched against the fasta files from the human Uniprot database (reviewed) in the RTS, set with tryptic digestion, TMTpro-16plex as fixed modification on Lysine (K) and N-Terminus together with cysteine (C) carbamidomethylation, and oxidation of methionine (M) as variable modification. 1 missed cleavage and 2 variable modifications were allowed with a maximum search time of 35 ms.

FDR filtering was enabled with 1 as Xcorrelation and 5 ppm of precursor tolerance. Precursors identified via RTS in the MS2 scan were further isolated in the quadrupole with a 2 m/z window, maximum injection time of 86 ms and normalized AGC target of 300%. The further MS3 fragmentation was operated with a normalized HCD collision energy at 50% and fragments were scanned with a resolution of 50000 in the range of 100 to 500 m/z. The MS performances were monitored by quality control of an in-house standard of HeLa-cell lysate, both at the beginning and the end of each sample set.

### Proteomic data analysis

Mass spectrometric raw files were analyzed with Proteome Discoverer 2.4 (Thermo Fisher Scientific) by using the built-in TMTpro Reporter ion standard quantification workflows. The search was run setting trypsin as enzyme (allowing maximum 2 missed cleavages), TMTpro16plex and carbamidomethylation of cysteine (C) as fixed modifications, while methionine (M) oxidation and acetylation of protein N-termini as variable modifications. Sequest search engine was used to match the MS^3^ spectra in the Uniprot homo sapiens database (Swiss-Prot reviewed including isoforms) with a precursor mass tolerance of 10 ppm and fragment mass tolerance of 0.6 Da. Percolator was used to score the results and to filter at 1% FDR. Reporter ion quantification was performed on MS^3^ spectra by applying isotopic error correction. Normalization and scaling were not included in the Proteome Discovery analysis and were performed successively on the protein result table. Here, proteins and proteins not identified as Master Protein or as Master Protein Candidate were removed first. Moreover, all the proteins identified with a sum of Unique + Razor peptides below 2 were removed. Loading normalization was performed summing the intensities for each TMT channel and calculating the respective correction factor on the average of the summed intensities. Hence, each protein intensity was normalized for the respective channel correction factor. Normalized intensities were further used to calculate the average among the replicates for one label, which was then used to obtain the fold changes (FC) of compound treated samples compared to DMSO control samples. The obtained values were log_2_ transformed. A two-sided T-test was performed on the normalized data and obtained p-values were –log_10_ transformed. Volcano plots were obtained plotting the log_2_ FC vs –log_10_ (p-value) in both Prism 9 (Graphpad Software, Inc. Version 9.41) and VolcanoseR.

### Lipidomic sample preparation

HeLa cells were harvested from multiple T175 flasks (confluence 90%) by trypsination and diluted to 2*10^5 cells/mL with DMEM medium (10 % FBS, 1 % P/S). Then the cells were plated in 10 cm dishes with a final cell count of 2*10^6 cells. After attachment overnight, the medium was replaced with fresh medium followed by compound incubation at the indicated concentrations for 18 h. For sample collection, the medium was removed and the cells were washed with 1× PBS, followed by trypsination. The detached cells were collected with medium and subsequently diluted to a concentration of 3*10^6 cells/mL. Then, each sample was centrifuged and washed with cold ammonium bicarbonate buffer (1 mL, 155 mM) twice. Lastly, the cells were resuspended in ammonium bicarbonate (1 mL, 155 mM) and stored at –80 °C.

### Lipidomics analysis

To Eppendorf tubes with 2*10^5 HeLa cells in 200µl 155mM ammonium bicarbonate were added 1000 µl chloroform/methanol (2:1, v/v, Rathburn Chemicals Ltd) and spiked with 12.5 µl of synthetic lipid standards (avanti polar lipids, details in table 1). Lipid extraction was executed on ice or 4°C according to our previously reported protocol.[50] Extracted lipid were subjected to FT MS and FT MS/MS analysis on an Orbitrap Fusion Tribrid mass spectrometer (Thermo Fisher Scientific) coupled to TriVersa NanoMate (Advion Biosciences, USA) a direct nanoelectrospray infusion robot. The lipid extract were mixed with positive and negative ionization mode solvents according to our previously reported protocol.[50] Mass spectrometric settings in positive and negative mode: The FT MS analysis operated with R*_m/z_*_200_ = 500,000; AGC value of 1 × 10^5^; maximum injection time of 50 ms; three microscan and FT MS/MS operated with R*_m/z_*_200_ = 15,000; AGC value of 2.5 × 10^4^; maximum injection time of 66 ms; one microscan, while the direct infusion settings of the robot follows our according to our previously reported protocol.[50] Lipid identification was performed with LipidXplorer version 1.2.7 and absolute quantification was performed using homemade software LipidQ.[50]

### Lipid annotation

Lipid species are annotated according to their sum composition, where glycerolipid and glycerophospholipid species denoted as: <lipid class>< total number of C in fatty acid moieties>:<total number of double bonds in fatty acid moieties>(e.g., PS 34:1). Ether glycerophospholipid species are denoted with “O-”(e.g., PS O-34:1). While sphingolipid species are denoted as <lipid class><total number of C in the long-chain base and fatty acid moiety>:<total number of double bonds in the long-chain base and fatty acid moiety>;<total number of OH groups in the long-chain base and fatty acid moiety>(e.g., Cer 34:1;2).[53–56]

### SREBP reporter gene assay

The SREBP reporter gene assay was performed by applying a modified procedure.[34] Shortly, a pSynSRE-T-Luc plasmid, which contains a promotor for 3-hydroxy-3-methylglutaryl-CoenzymA (HMG-CoA) synthase containing the SREBP responsive region linked to a firefly luciferase[57], was utilized. Additionally, a pRL-TK (*Renilla* luciferase) vector was used for normalization. Both plasmids were gifts from Prof. Timothy Osborne. Shortly, for transient transfection, HeLa cells cultured in DMEM (10 % FBS) without antibiotics were transfected by means of lipofection using Lipofectamine 2000 (11668030, Thermo Fisher Scientific). 1.5*10^6 cells per T25 flask were transfected with 3 µg of pSyn-SRE-T-luc and 1 µg of pRL-TK using a DNA:LF ratio of 1:3 (total amounts). The cells were incubated for 24 h and afterwards replated in a white 96-well plate (136102, Nunclon®, Thermo Fisher Scientific) with a final cell count of 2.5*10^4 cells per well. Cell attachment was allowed for 1 h before compound addition with the indicated concentrations and incubation for 18 h. The Luciferase activities were measured using the Dual-Glo® Luciferase Assay System (E2920, Promega) according to the manufacturer’s instructions. The signal for the firefly luciferase was divided by the *Renilla* luciferase signal to obtain normalized signal ratios which were then related to DMSO.

### Differential Light Scattering (DLS) assay

DLS experiments were performed to detect possible *in vitro* aggregation of SQS upon treatment with KY02111. Shortly, stored recombinant SQS (without His_6_-tag) was filtered using a Superdex 75 10/300 GL Column (Cytiva) to remove any aggregates formed by the freeze-thawing process. Freshly filtered protein was then diluted to 0.25 mg/mL in buffer (20 mM Tris-HCl, pH 7.3, 2 mM MgCl_2_, 0.1 mM EDTA), slowly filtered through 0.2 µm pore size filter (6876-2502, Whatman®, GE Healthcare Life Sciences)) and added to a polystyrene cuvette (67.741, Sarstedt). The sample was allowed to equilibrate to room temperature for 2 min before Zetasizer Nano (Malvern) was used to measure and analyse the particle size. SQS alone was found to have a diameter (d) of about 6 nm. This measurement was performed for every new sample before compound addition at the indicated concentrations. The data was plotted as either intensity or volume vs size with Prism 9 (Graphpad Software, Inc. Version 9.41).

### ER membrane insertion assay

Pulse-chase assays of U2OS Flp-In™ T-Rex™ cells (WT/ΔEMC6) expressing HA:SQS_opsin_ were carried out as described previously by Volkmar *et al.*[23] Briefly, cells were starved in methionine/cysteine-free DMEM (Lonza) plus 10% dialyzed FCS (10 min) and metabolically labelled by adding ^35^S-methionine/cysteine [Met/Cys, EXPRE^35^S^35^S Protein Labelling Mix (PerkinElmer); 150 μCi/10 plate] for 10 min. After removal of media, cells were rinsed thrice with PBS and incubated in DMEM plus 10% FCS and methionine/cysteine (50 mM each) for the indicated times (0 and 40 min). Cells were collected and resuspended in lysis buffer (50 mM Tris-HCl pH 7.5, 150 mM NaCl, 5 mM EDTA, 2 mM NEM, 1× cOmplete™ Protease Inhibitor Cocktail (Roche)) + 1% Triton X-100 (v/v) and post-nuclear fractions pre-cleared overnight using unconjugated Sepharose beads. HA:SQS_opsin_ was immunoprecipitated from the detergent-soluble fraction using 70 μl of 50% protein G resin (Roche) slurry and anti-HA antibody (12CA5). Immunoprecipitated material was resuspended in 2× Laemmli buffer plus 20 mM DTT, separated by 4-15% gradient SDS-PAGE, and the radiolabelled proteins visualised using a phosphoimager and the QuantityOne and Image Lab software packages (Bio-Rad). To deglycosylate samples, EndoHf (NEB, 500 units) was added to Laemmli eluates and incubated for 30 min at 37°C prior to loading.

## Notes

### Competing Interest Statement

The authors have declared no competing interest.

