## Supporting Information for "Identification of non-conventional small molecule degraders and stabilizers of squalene synthase"

### Supplementary Information

**Table of contents**

| | |
|---|---|
| Supporting figures 1-11 | 1-7 |
| Chemical synthesis and compound characterization | 8-42 |
| References | 43 |
| NMR data | 44-70 |
| LCMS purity of SQSI, KY02111 and 18 | 71-72 |

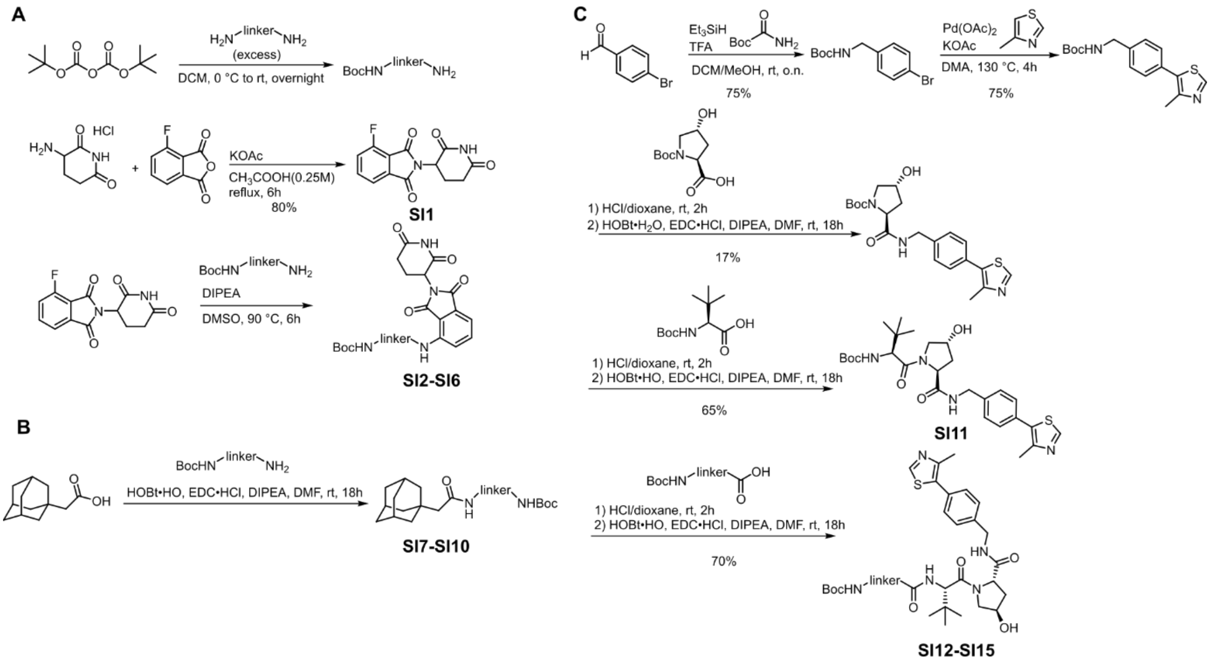


**SI Figure 1: Synthesis of degrader modalities.** Linker is representative for all linker-degrader sets. **A)** Linker-pomalidomide conjugates.[1] **B)** Linker-adamantane conjugates.[2] **C)** Linker-VH032 conjugates.[3]

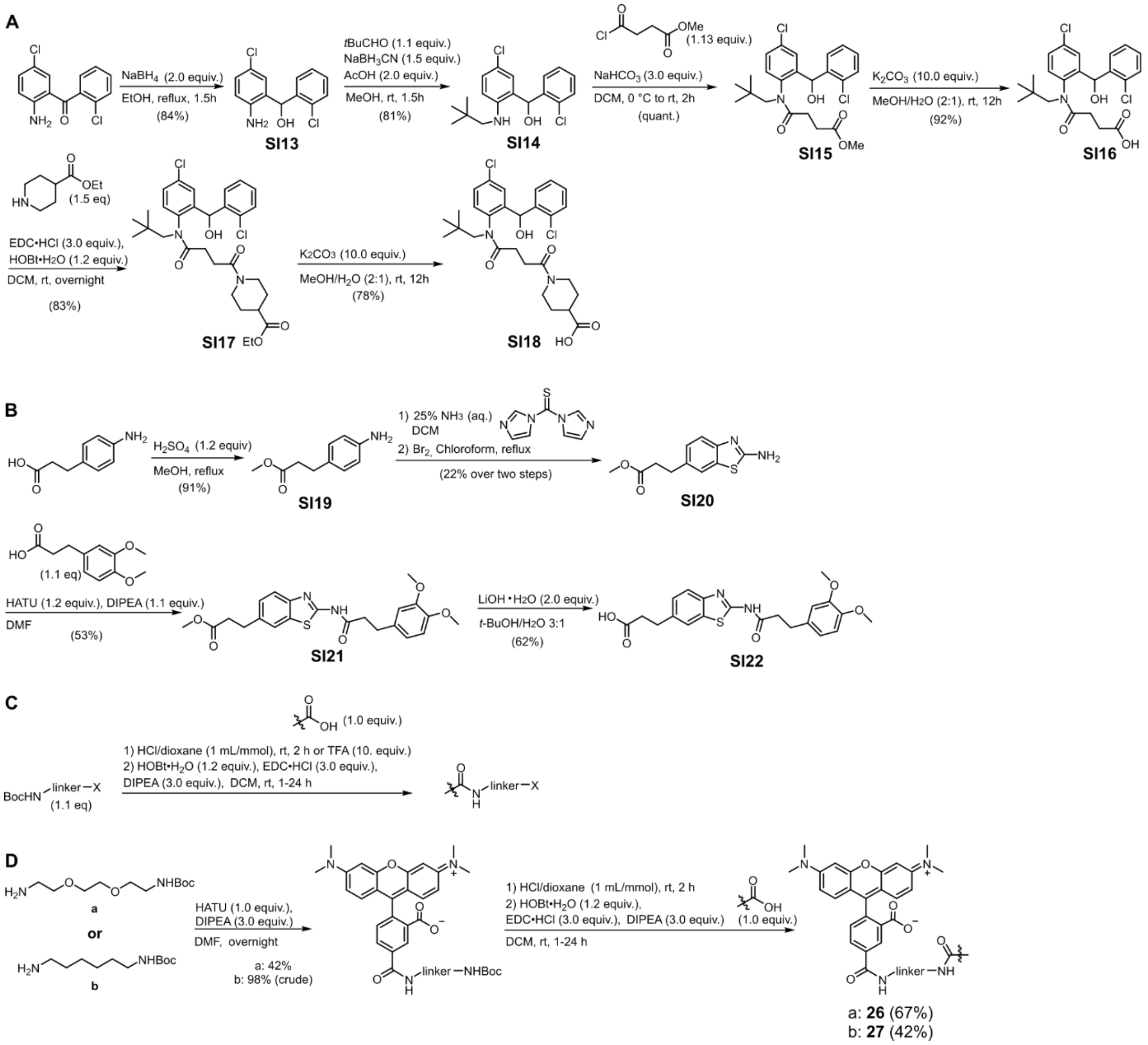

**SI Figure 2: Synthesis of SQS-degrader library molecules. A)** Synthesis of a SQSI-derivative, a carboxylic acid coupled to linker-degrader conjugates.[4] **B)** Synthesis of a SO2093 derivative, a carboxylic acid coupled to linker-dergrader conjugates.[5] **C)** Amide coupling procedure for the synthesis of SQS-degraders. **D)** Synthesis of FP tracers 26 and 27. 5-TAMRA was synthesized as reported before (not shown).[6]

A

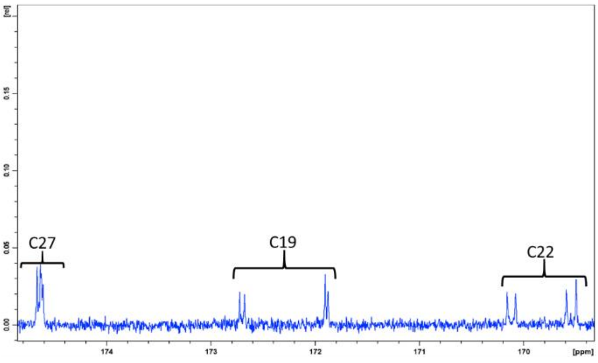


B

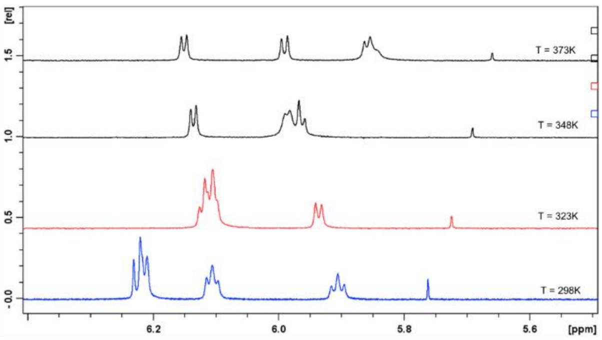


C

| Temperature | CH ratio (H7) I2/I1 | t-butyl CH3 ratio (H17) I2/I1 |
|---|---|---|
| 298K | 1.1/0.8 = 1.4 | 9.0/6.4 = 1.4 |
| 323K | Not resolved (overlap with H8) | 9.0/6.9 = 1.3 |
| 348K | Not resolved (overlap with H8) | 9.0/7.6 = 1.2 |
| 373K | 1.0/0.9 = 1.1 | 9.0/8.3 = 1.1 |

D

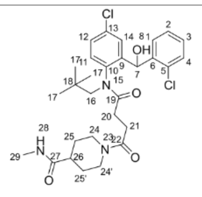

Isomer 1

| Label | 13C | 1H | HMBC | ROESY |
|---|---|---|---|---|
| 1 | 129.46 | 7.56 | 3, 5, 7 | 7(0.5), 8(2), 14(1) |
| 2 | 127.85 | 7.44 | | |
| 3 | 129.88 | 7.39 | | |
| 4 | 130.02 | 7.48 | | |
| 5 | 132.4 | - | | |
| 6 | 140.06 | - | | |
| 7 | 67.63 | 5.9046 | 5, 6, 9, 14 | 1, 8, 14 16a(w), 16b, 20a, 21b |
| 8 | **OH** | 6.2240 | 5, 7, 9 | 1, 7, 14, 16b, 21b |
| 9 | 142.75 | | | |
| 10 | 141.3 | | | |
| 11 | 134.6 | 7.61 | **15** | 21a(w) |
| 12 | 128.56 | 7.487 | | |
| 13 | 132.9 | | | |
| 14 | 128.91 | 7.04 | 7, 10, 12 | |
| 15 | **N** 128.2 | | | |
| 16 | 57.94 | 4.215/2.575 | A: 10, **15**, 19 B: 10, 19 | A: 7, 16b, 17 B: 7, 8, 16a, 17 |
| 17 | 28.84 | 0.761 | | 11, 16a, 16b |
| 18 | 34.2 | - | | |
| 19 | 172.7 | - | | |
| 20 | 28.53 | 2.58/2.4 | A: 19, 22 B: 19, 22 | A: 7, 20b B: 20a, 21b |
| 21 | 29.73 | 2.34/2.02 | A: 19, 22 B: 19, 22 | A: 11, 21b B: 7(w), 8 20b, 21a |
| 22 | 170.11 | - | | |
| 23 | na | | | |
| 24 24' | 44.78 41.21 | 3.85/2.98 4.3/2.53 | | |
| 25 25' | 29.27 28.64 | 1.685/1.455 1.54/1.26 | | |
| 26 | 42.19 | 2.31 | | |
| 27 | 174.65 | | | |
| 28 | **N** 101.2 | 7.72 | | 26, 29 |
| 29 | 25.96 | 2.56 | | |

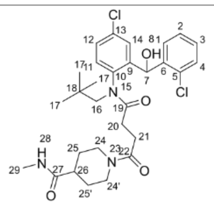

Isomer 2

| Label | 13C | 1H | HMBC | ROESY |
|---|---|---|---|---|
| 1 | 130.06 | 7.288 | 3, 5, 7 | 7(0.7), 8(1.4), 14(1) |
| 2 | 129.85 | 7.40 | | |
| 3 | 130.05 | 7.31 | | |
| 4 | 129.75 | 7.32 | | |
| 5 | 132.8 | | | |
| 6 | 139.95 | | | |
| 7 | 66.59 | 6.1064 | 1, 5, 6, 9, 14 | 1, 8, 14, 16a(w), 16b, 17 |
| 8 | **OH** | 6.2110 | 5, 7, 9 | 1, 7, 14, 16b, 17 |
| 9 | 142.94 | - | | |
| 10 | 140.2 | - | | |
| 11 | 134.3 | 7.61 | **15** | 21a |
| 12 | 128.2 | 7.46 | | |
| 13 | 132.9 | | | |
| 14 | 128.55 | 7.65 | | 8 |
| 15 | **N** 128.8 | - | | |
| 16 | 58.66 | 4.4065/3.02 | A: 10, 19 B: 10 ,19 | A: 7, 16b 17 B: 7, 8, 16a 17 |
| 17 | 29.1 | 0.84 | | 11, 16a, 16b |
| 18 | 34.6 | - | | |
| 19 | 171.9 | - | | |
| 20 | 28.41 | 2.375/1.703 | A: 22 B: 19, 22 | A: 20b B: 20a ,21b |
| 21 | 29.37 | 2.155/1.004 | A: 19, 22 B: 19, 22 | A: 11, 21b B: 20b, 21a |
| 22 | 169.46 | - | | |
| 23 | na | - | | |
| 24 24' | 44.7 41.15 | 3.66/2.89 4.23/2.48 | | |
| 25 25' | 29.2 28.6 | 1.64/1.396 | | |
| 26 | 42.16 | 2.29 | | |
| 27 | 174.65 | - | | |
| 28 | **N** 101.2 | 7.72 | | 26, 29 |
| 29 | 25.96 | 2.56 | | |

**SI Figure 3:** The utilized SQS-ligand is known to produce a set of atropoisomers due to hindered rotation around the C3-N bond in the presence of the *t*-butyl group. Additionally we identified a second pair of atropoisomers in DMSO-d6. **A-D)** A full spectra assignment for SQSI can be made based on 1D and 2D NMR spectra. However, it is evident that two major isomers are present in the sample, termed isomer 1 and isomer 2. The chemical shift assignment is shown in **D)**. In addition, each of the isomers then also exhibits a second type of isomerism, and thus at least four structural isomers are identified. This is most clear in the 1D 13C spectrum in-which most carbon sites show four distinct NMR signals – illustrated in **A)** for the three amide carbon atoms. Similar evidence can be seen in **B)**, the 1H spectrum, where a total of four doublets are observed for H7, and H16a and H16b for I2 are both split into two doublets. Interestingly, H16a and H16b for I1 does not show such a splitting, and both appear as single doublets. **B/C)** $^{1}$H NMR spectra has been measured at 298K, 323K, 348K and 373K for the sample dissolved in DMSO-d6. It is seen that the set of "triplets" for H7 collapses into doublets already at 323K, confirming a relatively low energy barrier for the first type of isomerism (which is also suggested from the small $^{13}$C chemical shift differences). In contrast, the two doublets only show minor signs of collapsing into a single doublet and stay well-separated even at 373K ($\Delta\delta$ (298K) = 0.20 ppm, $\Delta\delta$ (323K) = 0.19 ppm, $\Delta\delta$ (348K = 0.17 ppm and $\Delta\delta$ (373K

= 0.16 ppm). The relative integral area for both H7 and H17 has been determined at the four temperatures, and show that equilibrium is moving towards I1 as the temperature is increased.

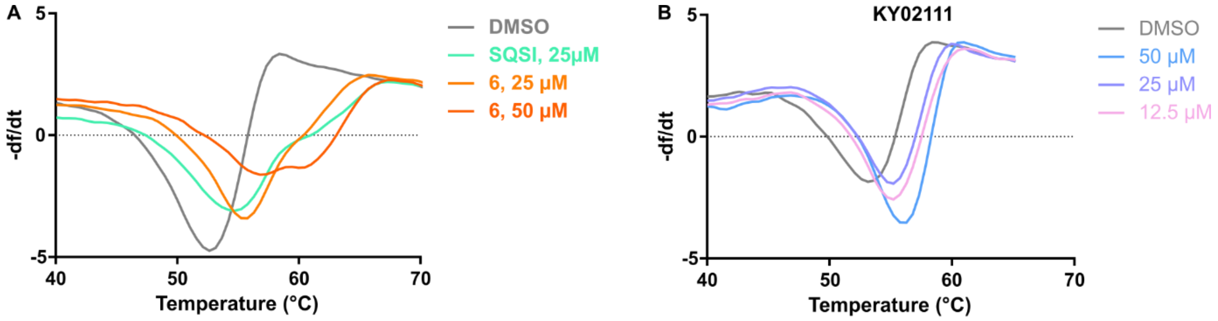


C

| Compound | dTm [°C] 12.5 µM | 25 µM | 50 µM |
|---|---|---|---|
| SQSI | 0.9 ± 0.8 | 1.4 ± 0.2 | 6.1 ± 1.0 |
| KY02111 | 2.1 ± 0.4 | 1.8 ± 0.4 | 2.3 ± 0.1 |
| SO2093 | 2.3 ± 0.5 | 2.2 ± 0.3 | 2.0 ± 0.5 |
| 6 | 0.7 ± 0.1 | 2.8 ± 0.3 | 6.3 ± 1.0 |
| 7 | 1.6 ± 0.1 | 3.0 ± 0.2 | 6.2 ± 1.6 |
| 8 | 1.5 ± 0.6 | 2.3 ± 0.6 | 5.3 ± 1.5 |
| 13 | 2.1 ± 0.7 | 2.4 ± 0.3 | 1.9 ± 0.1 |
| 18 | 2.9 ± 0.4 | 3.6 ± 0.3 | 3.3 ± 0.9 |
| 22 | 1.9 ± 0.6 | 2.6 ± 0.3 | 2.9 ± 0.3 |
| 23 | 2.0 ± 0.6 | 2.1 ± 0.4 | 1.9 ± 0.7 |

**SI Figure 4: Differential scanning fluorimetry (DSF) data for selected members of the SQS degrader library. A)/B)** First derivative of the melting curves for SQSI, **6** and KY02111. Recombinant SQS (His6-31-370, 0.5 mg/mL) was incubated with sypro orange (1x) and either DMSO, 6, or KY02111 at the indicated concentrations. The fluorescence intensity was measured in a Roche LightCycler 480 II. **C)** Overview of melting temperatures ΔTm [°C] for tested members of the SQS degrader library (n = 3, mean ± SD shown).

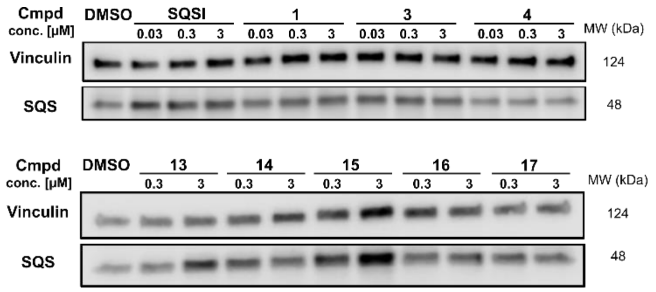


**SI Figure 5:** ***In cellulo*** **SQS-degrader screen: Complimentary data for library compounds 1-5 and 13-17.** Western blots showing the changes in SQS protein levels after 18 h treatment of HeLa cells with compounds at indicated concentrations (n = 2 or 3).

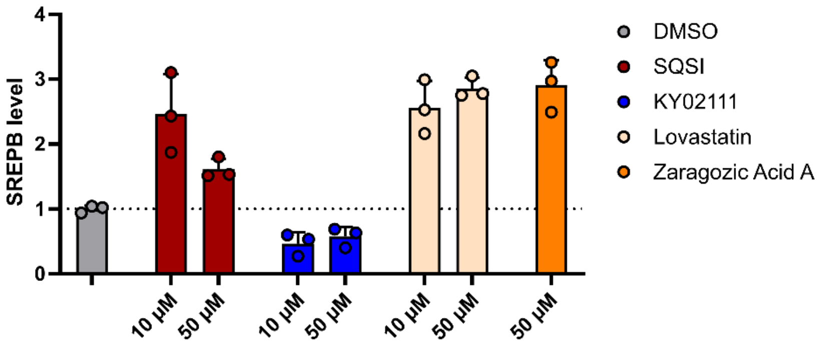


**SI Figure 6: B)** SREBP reporter gene assay to see indirect upregulation of cholesterol biosynthesis.[7] HeLa cells were transfected with a pSynSRE-T-Luc plasmid, containing the promotor for HMG-CoA synthase (HMGCS) containing the SREBP responsive region linked to a firefly luciferase and a pRL-TK (*Renilla* luciferase) vector. For transient transfection, HeLa cells were transfected by means of lipofection using Lipofectamine 2000 for 24 h. Afterwards the cells were re-plated and incubated with compounds for 18 h. The Promega Dual-Glo Luciferase assay system was used for reading out the firefly and renilla luciferase activity in the same sample (n = 2/3 (biol./tech.), mean ± SD shown).

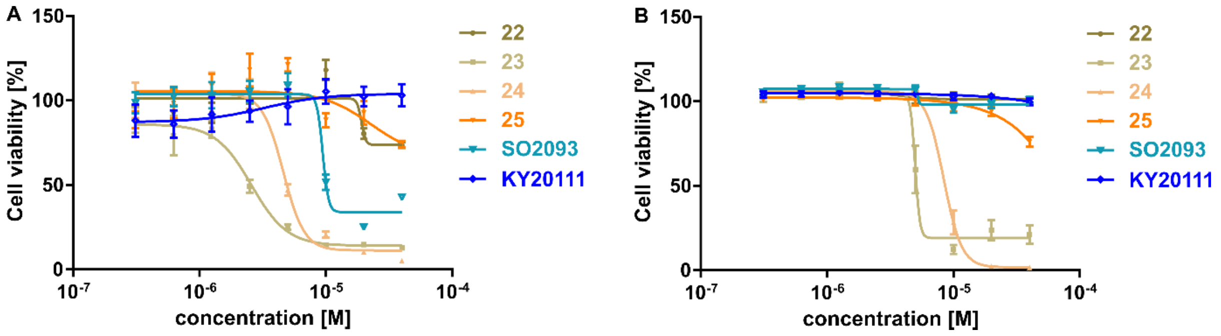


**SI Figure 7**: Cell viability data for selected compounds in HeLa cells. Viability was analyzed after the indicated time points utilizing the Cell-Titer Blue assay system by Promega. **A)/B)** The indicated compounds were used to incubate HeLa cells for 18 (A) and 48 (B) h. The dilution series was started at 40 µM (n = 3 (tech. and biol.), mean ± SD shown).

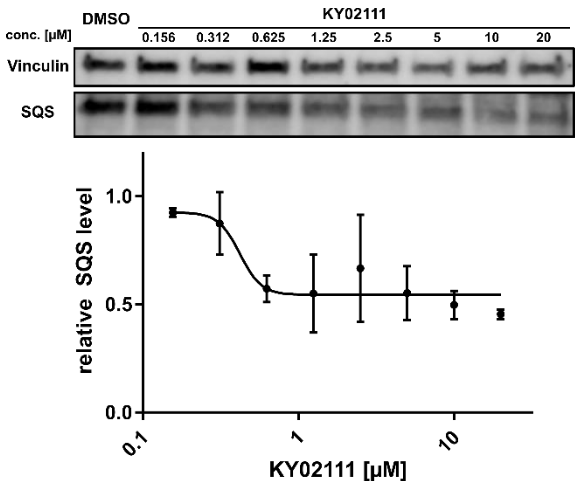


**SI Figure 8:** KY02111 dose-response in U2OS cells incubated for 18 h (n = 2, mean ± SEM shown).

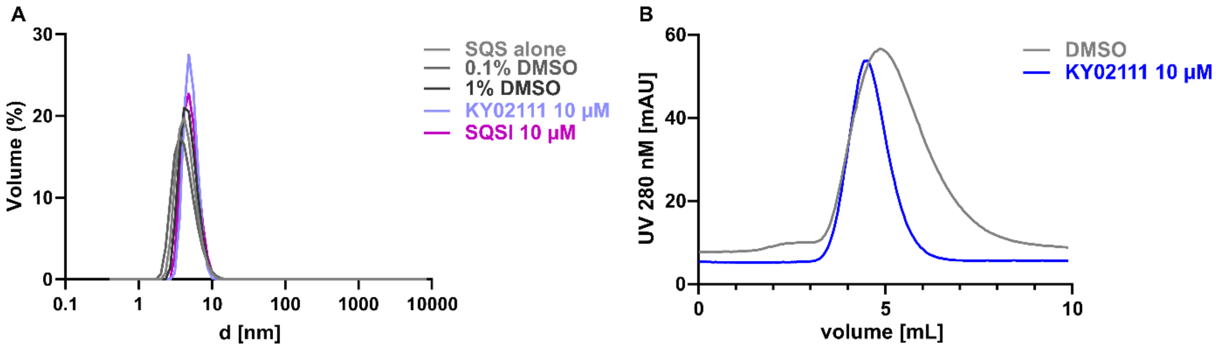


**SI Figure 9:** Investigation of potential SQS aggregation by KY02111. **A)** Differential light scattering (DLS) data for the indicated conditions. Recombinant SQS (31-370) was filtered by size-exclusion chromatography (SEC) and then diluted to 0.25 mg/mL). The sample was then filtered through a 0.2 µm syringe filter and added to a cuvette. After an initial measuremnt with SQS alone, compounds were added at the indicated concentrations. Each sample was prepared individually. **B)** UV peak for SEC purification of recombinant SQS (31-370, 0.25 mg/mL) pre-incubated with either DMSO or KY02111 (10 µM) for 1 h.

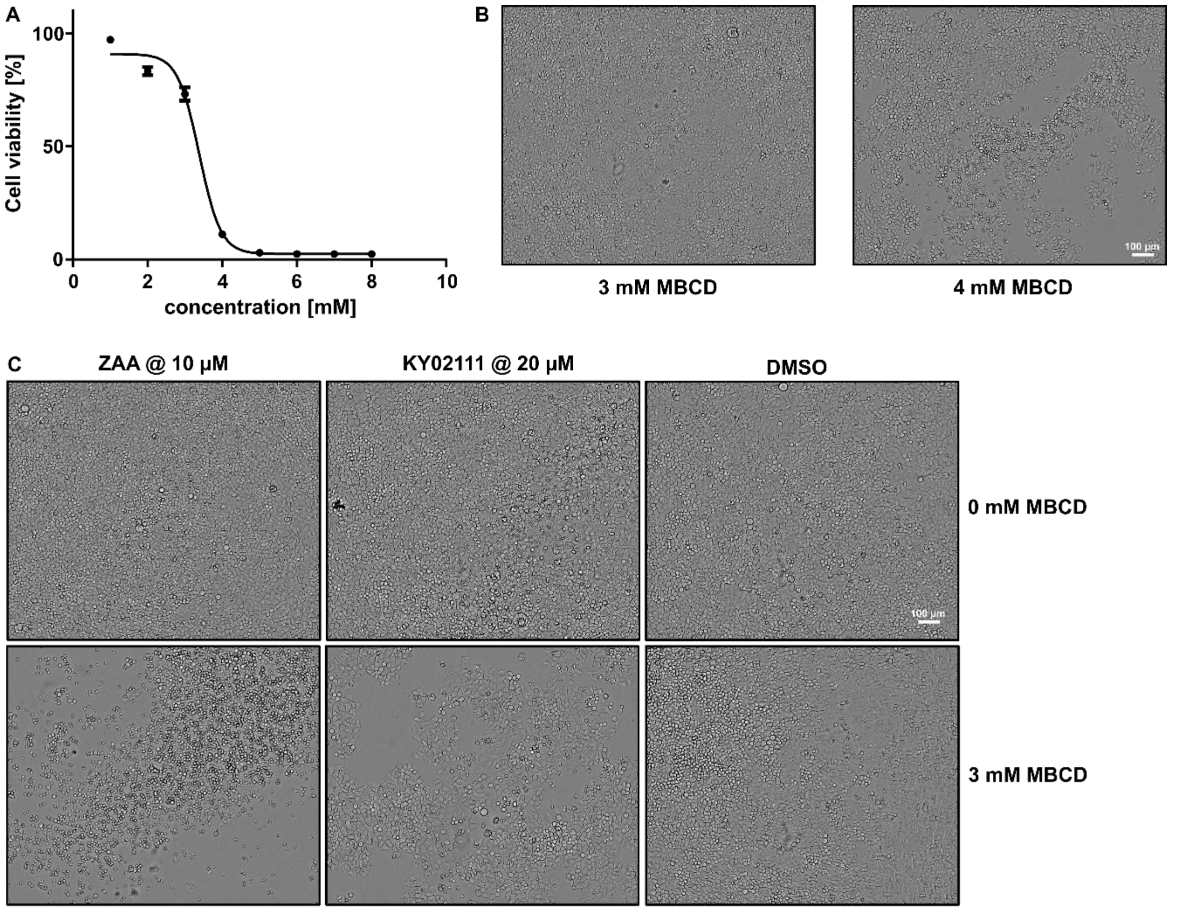


**SI Figure 10:** Cell viability data for selected compounds in HeLa cells. Viability was analyzed after the indicated time points utilizing the Cell-Titer Blue assay system by Promega. **A)/B)** Cells were titrated with different concentrations of MBCD for 72 h (in standard medium, 10% FBS, 1% PS). Pictures were also taken after 72 h. 3 mM MBCD was chosen as a non-toxic concentration for follow-up experiments. C) Comparison in cell viability between cells incubated with ZAA (10 µM), KY02111 (20 µM) and DMSO, grown with (3 mM) or without MBCD for 72 h.

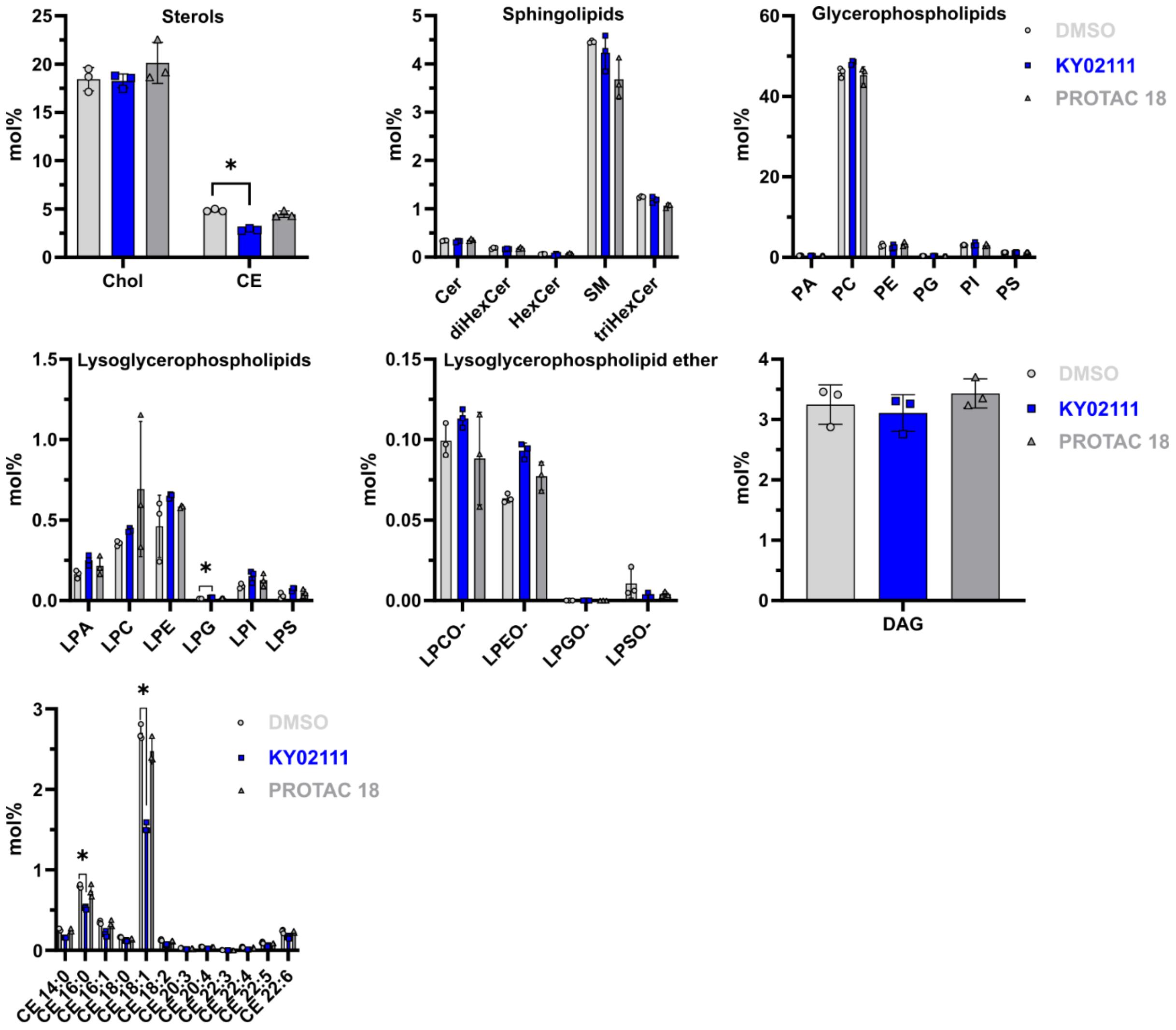


**SI Figure 11:** Bar plots showing the mol% changes for respective lipid classes after treatment of HeLa cells with KY02111 (10 µM) or PROTAC 18 (5 µM) after 18 h ($n = 3$, Holm-Šídák correction was used to determine adjusted p-values $p < 0.005$). For changes of specific species within the lipid classes the reader is referred to the supplementary excel file containing the full data set.

### General directions

All reactions were run under a $N_2$ atmosphere unless otherwise specified and were monitored by thin layer chromatography (TLC) and/or reversed-phase ultra-performance liquid chromatography mass spectrometry (RP-UPLC-MS). Commercially available reagents were purified according to standard procedures or were used as received from Sigma Aldrich, Alfa Aesar, Acros Organics, Combi-Blocks, Fisher Scientific, Strem, and Merck. All solvents used were of HPLC quality and dry solvents (DCM, Et2O, THF, and Toluene) were obtained from a PureSolv system (Innovative Technology, Tronyx). Methanol was stored over activated 3 Å molecular sieves before use. Analytical TLC was conducted on Merck aluminium sheets covered with silica (C60). The plates were either visualized under UV-light or stained by dipping in a developing agent followed by heating. $KMnO_4$ [$KMnO_4$ (3 g) in water (300 mL), $K_2CO_3$(20 g) and 5% aqueous NaOH (5 mL)] and cerium molybdate [$Ce(NH_4)_2(NO_3)_6$ (0.5g), $(NH_4)_6Mo_7O_{24}{\cdot}4H_2O$ (24.0 g), and $H_2SO_4$ (24.0 g)] were used as developing agents. Flash column chromatography was performed using Merck Geduran® Si 60 (40-63 μm) silica gel. All new compounds were characterized by NMR, MS (ESI) and HRMS (ESI) (byproducts were not fully characterized). Structural assignments were made when possible for new compounds using COSY, HSQC, HMBC, H2BC, and NOESY spectra where appropriate. For the recording of $^1$H NMR and $^{13}$C NMR a Bruker Ascend with a Prodigy cryoprobe (operating at 400 MHz for proton and 100 MHz for carbon) was used. The chemical shifts (δ) are reported in parts per million (ppm) and the coupling constants (J) in Hz. Spectra were referenced using the residual solvent peaks of the respective solvent; DMSO (δ 2.50 ppm for $^1$H NMR and δ 39.52 ppm for $^{13}$C NMR), $CDCl_3$ (δ 7.26 ppm for 1H NMR and δ 77.16 ppm for $^{13}$C NMR), $CD_3OD$ (δ 3.31 ppm for $^1$H NMR and δ 49.00 ppm for $^{13}$C NMR). The following abbreviations were used to report peak multiplicities: s = singlet, d = doublet, t = triplet, q = quartet, dd = doublet of doublets, sept = septet, m = multiplet, bs = broad singlet. Analytical RP-UPLC-MS (ESI) analysis was performed on a S2 Waters AQUITY RP-UPLC system equipped with a diode array detector using an Thermo Accucore C18 column (d 2.6 μm, 2.1 x 50 mm; column temp: 50 °C; flow: 1.0 mL/min). Eluents A (0.1% $HCO_2H$ in $H_2O$) and B (0.1% $HCO_2H$ in MeCN) were used in a linear gradient (5% B to 100% B) in 2.4 min and then held for 0.1 min at 100% B (total run time: 2.6 min). The LC system was coupled to a SQD mass spectrometer. Preparative RP-HPLC was carried out on a Waters Alliance reversed-phase HPLC system consisting of a Waters 2545 Binary Gradient Module equipped with an xBridge BEH C18 OBD Prep Column (130 A, 5 μm, 30 x 150 mm) operating at 20 °C and a flow rate of 20 mL/min, a Waters Photodiode Array Detector (detecting at 210-600 nm), a Waters UV Fraction Manager, and a Waters 2767 Sample Manager. Eluents A1 (0.1% $HCO_2H$ in $H_2O$) and B1 (0.1% $HCO_2H$ in MeCN) were used. Analytical LC-HRMS (ESI) analysis was performed on a Waters Alliance

2695 system. Samples were injected directly and the LC system was coupled to a Waters LCT Premier XE Micromass equipped with a Lock Mass probe operating in positive electrospray mode. Eluents A (0.1%$HCO_2H$ in H2O) and B (0.1% $HCO_2H$ in MeCN) were used in a 1:1 ratio for a total run time of 2 min. 17 Chiral HPLC was run on a Waters 2695 Alliance Separations module with a Waters 2996 PhotoDiode Array detector, equipped with a ChiralPak AD-H column (5 µM, 250 x 4.6 mm).

**Note for active site binding compounds:** The utilized SQS ligand is known to produce a set of atropoisomers due to hindered rotation around the C3-N bond in the presence of the *t*-butyl group. Additionally we identified a second pair of atropoisomers in DMSO-$d_6$ which we analyzed via variable temperature (VT) NMR and 2D NMR. The data is presented and discussed in SI Fig. 3. Due to the presence of 4 isomers in DMSO-$d_6$, we could not clearly assign all observed peaks for compounds based on SQSI. Therefore only characteristic peaks are highlighted in the data, where all 4 isomers are included, SQSI was fully assigned (SI Fig. 3D).

### Chemical synthesis

Pomalidomide as well as Boc-protected pomalidomide-linker conjugates were synthesized and characterized as previously described[1]. VH032 as well as Boc-protected VH032-linker conjugates were synthesized by a slightly modified reported procedure (see general procedures: amide coupling A).[3] Boc-protected adamantane-linker conjugates were synthesized via the general amide coupling A procedure and characterized by comparison with previously reported data.[8] New compounds were fully characterized.

### Pomalidomide

2-(2,6-dioxopiperidin-3-yl)-4-fluoroisoindoline-1,3-dione **SI1**

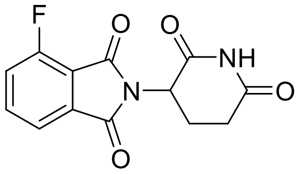


**$^1$H NMR** (400 MHz, DMSO-$d_6$): δ 11.16 (s, 1H), 7.96 (td, $J$ = 7.9, 4.5 Hz, 1H), 7.80 (d, $J$ = 7.3 Hz, 1H), 7.75 (t, $J$ = 8.9 Hz, 1H), 5.17 (dd, $J$ = 12.9, 5.4 Hz, 1H), 2.96 – 2.82 (m, 1H), 2.69 – 2.53 (m, 2H), 2.12 – 2.01 (m, 1H) ppm. **LCMS** (ESI) *m/z* [M + H] calcd for $C_{13}H_{10}FN_2O_4^+$: 278.1; found: 276.9.

**$S_NAr$ for the synthesis of pomalidomide-linker conjugates**

The mono-Boc protected linker (1.0 equiv.) was dissolved in dry DMSO (15 mL) and DIPEA (4.0 eq) was added. Then, 4-fluoro-thalidomide (1.0 eq) was added, and the mixture was stirred at 90 °C for 6 h. After cooling to rt, the yellow or green mixture was partitioned between half-saturated brine (100 mL) and EtOAc (3 × 50 mL). The combined organic layers were further washed with saturated $NH_4Cl$ solution, 5% LiCl solution, and brine (each 50 mL). The organic layers were dried over $Na_2SO_4$, filtered and concentrated *in vacuo*. The crude product was purified by column chromatography (hexane/EtOAc 1:1 to 1:2) to give a yellow solid.

*tert*-butyl (6-((2-(2,6-dioxopiperidin-3-yl)-1,3-dioxoisoindolin-4-yl)amino)hexyl)carbamate **SI2**

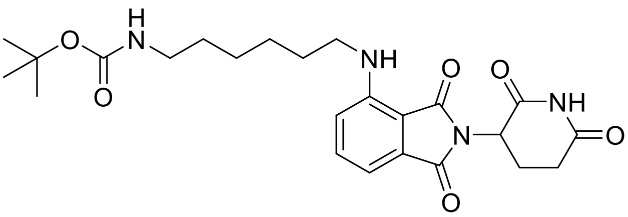

**$^1$H NMR** (400 MHz, $CDCl_3$): δ 7.99 (s, 1H), 7.54 – 7.49 (m, 1H), 7.12 (d, *J* = 7.1 Hz, 1H), 6.90 (d, *J* = 8.6 Hz, 1H), 4.97 – 4.90 (m, 1H), 4.53 (s, 1H), 3.29 (t, *J* = 7.0 Hz, 2H), 3.14 (s, 2H), 2.96 – 2.71 (m, 3H), 2.08 – 1.97 (m, 1H), 1.75 – 1.57 (m, 4H), 1.53 (q, *J* = 7.3 Hz, 2H), 1.47 (s, 9H), 1.45(m, 2H) ppm. One signal for NH is missing due to H-D exchange. **LCMS** (ESI) *m/z* [M + H] calcd for $C_{24}H_{33}N_4O_6^+$: 473.2; found: 473.0.

*tert*-butyl(2-(2-(2-((2-(2,6-dioxopiperidin-3-yl)-1,3-dioxoisoindolin-4-yl)amino)ethoxy)ethoxy)ethyl)carbamate **SI3**

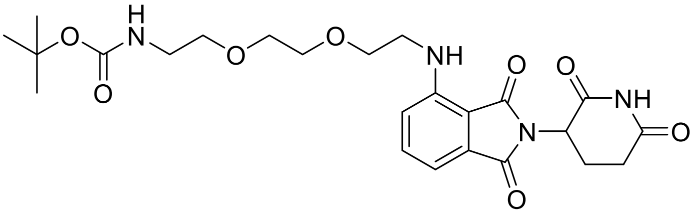

**$^1$H NMR** (400 MHz, $CDCl_3$): δ 8.48 (s, 1H), 7.51 (dd, *J* = 8.5, 7.1 Hz, 1H), 7.13 (d, *J* = 7.1 Hz, 1H), 6.93 (d, *J* = 8.5 Hz, 1H), 5.09 (s, 1H), 4.97 (s, 1H), 3.74 (t, *J* = 5.3 Hz, 2H), 3.67 (hept, *J* = 2.7 Hz, 4H), 3.58 (t, *J* = 5.2 Hz, 2H), 3.49 (t, *J* = 5.3 Hz, 2H), 3.33 (t, *J* = 5.2 Hz, 2H), 2.93 – 2.71 (m, 3H), 2.08 – 1.97 (m, 1H), 1.45 (s, 9H) ppm. One signal for NH is missing due to H-D exchange. **LCMS** (ESI) *m/z* [M + H] calcd for $C_{24}H_{33}N_4O_8^+$: 505.2; found: 505.1.

*tert*-butyl(3-(4-(3-((2-(2,6-dioxopiperidin-3-yl)-1,3-dioxoisoindolin-4-yl)amino)propoxy)butoxy)propyl)carbamate **SI4**

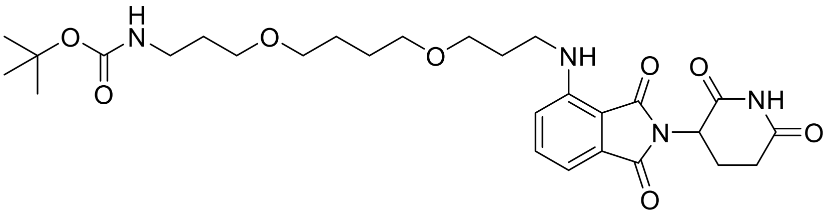

**¹H NMR** (400 MHz, $CDCl_3$): δ 8.25 (s, 1H), 7.51 (dd, *J* = 8.5, 7.1 Hz, 1H), 7.11 (d, *J* = 7.1 Hz, 1H), 6.94 (d, *J* = 8.5 Hz, 1H), 6.47 (s, 1H), 4.93 (dd, *J* = 12.0, 5.3 Hz, 1H), 3.56 (t, *J* = 5.7 Hz, 2H), 3.52 – 3.39 (m, 8H), 3.23 (q, *J* = 6.2 Hz, 2H), 2.96 – 2.70 (m, 3H), 2.08 – 1.97 (m, 1H), 1.94 (p, *J* = 6.2 Hz, 2H), 1.76 (p, *J* = 6.3 Hz, 2H), 1.72 – 1.61 (m, 4H), 1.46 (s, 9H) ppm. One signal for NH is missing due to H-D exchange. **LCMS** (ESI) *m/z* [M + H] calcd for $C_{28}H_{41}N_4O_8^+$: 561.3; found: 561.2.

*tert*-butyl (3-(2-(2-(3-((2-(2,6-dioxopiperidin-3-yl)-1,3-dioxoisoindolin-4-yl)amino)propoxy)ethoxy)ethoxy)propyl)carbamate **SI5**

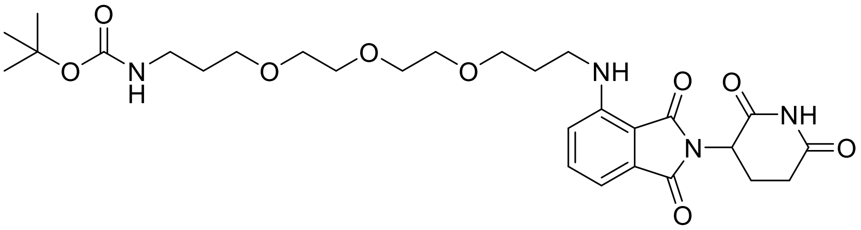

**¹H NMR** (400 MHz, $CDCl_3$): δ 8.17 (s, 1H), 7.51 (dd, *J* = 8.5, 7.1 Hz, 1H), 7.11 (d, *J* = 7.1 Hz, 1H), 6.96 (d, *J* = 8.5 Hz, 1H), 4.93 (dd, *J* = 12.1, 5.4 Hz, 1H), 3.58 – 3.41 (m, 10H), 3.55 (t, *J* = 6.0 Hz, 2H), 3.43 (t, *J* = 6.6 Hz, 2H), 3.24 (q, *J* = 5.8 Hz, 2H), 2.96 – 2.68 (m, 3H), 2.08 – 1.97 (m, 1H), 1.96 (p, *J* = 6.2 Hz, 2H), 1.77 (p, *J* = 6.2 Hz, 2H), 1.46 (s, 9H) ppm. Two signals for NH are missing due to H-D exchange. **LCMS** (ESI) *m/z* [M + H] calcd for $C_{28}H_{41}N_4O_9^+$: 577.3; found: 577.3.

*tert*-butyl (10-((2-(2,6-dioxopiperidin-3-yl)-1,3-dioxoisoindolin-4-yl)amino)decyl)carbamate **SI6**

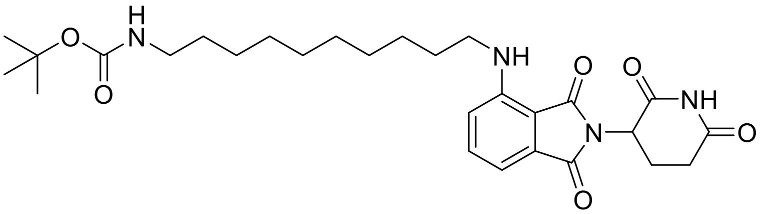

**¹H NMR** (400 MHz, $CDCl_3$): δ 8.08 (s, 1H), 7.51 (dd, *J* = 8.5, 7.1 Hz, 1H), 7.11 (d, *J* = 7.1 Hz, 1H), 6.91 (d, *J* = 8.5 Hz, 1H), 4.98 – 4.89 (m, 1H), 4.54 (s, 1H), 3.28 (t, *J* = 7.0 Hz, 2H), 3.12 (t, *J* = 7.1 Hz, 2H), 2.98 – 2.68 (m, 3H), 2.08 – 1.97 (m, 1H), 1.68 (p, *J* = 7.1 Hz, 4H), 1.47 (s, 9H), 1.42 (d, *J* = 8.3 Hz, 2H), 1.37 – 1.26 (m, 10H) ppm. One signal for NH is missing due to H-D exchange. **LCMS** (ESI) *m/z* [M + H] calcd for $C_{28}H_{41}N_4O_6^+$: 529.3; found: 529.3.

**VH032**

*tert*-butyl ((*S*)-1-((2*S*,4*R*)-4-hydroxy-2-((4-(4-methylthiazol-5-yl)benzyl)carbamoyl)pyrrolidin-1-yl)-3,3-dimethyl-1-oxobutan-2-yl)carbamate **SI11**

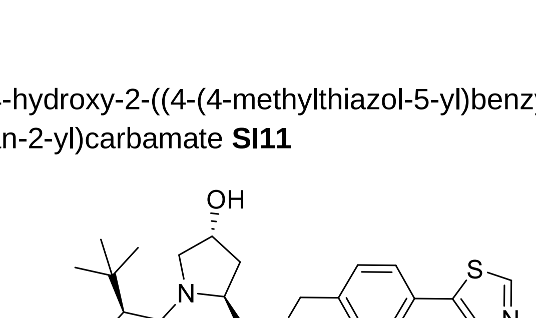

**¹H NMR** (400 MHz, DMSO): δ 8.98 (s, 1H), 8.58 (t, *J* = 6.1 Hz, 1H), 7.44 – 7.33 (m, 4H), 6.46 (d, *J* = 9.3 Hz, 1H), 5.15 (d, *J* = 3.5 Hz, 1H), 4.49 – 4.33 (m, 3H), 4.23 (dd, *J* = 15.8, 5.6 Hz,

1H), 4.15 (d, *J* = 9.3 Hz, 1H), 3.70 – 3.52 (m, 2H), 2.44 (s, 3H), 2.09 – 1.99 (m, 1H), 1.95 – 1.84 (m, 1H), 1.38 (s, 9H), 0.93 (s, 9H) ppm. **LCMS** (ESI) *m/z* [M + H] calcd for $C_{27}H_{39}N_4O_5S^+$: 531.3; found: 531.6.

*tert*-butyl (10-(((*S*)-1-((2*S*,4*R*)-4-hydroxy-2-((4-(4-methylthiazol-5-yl)benzyl)carbamoyl)pyrrolidin-1-yl)-3,3-dimethyl-1-oxobutan-2-yl)amino)-10-oxodecyl)carbamate **SI12**

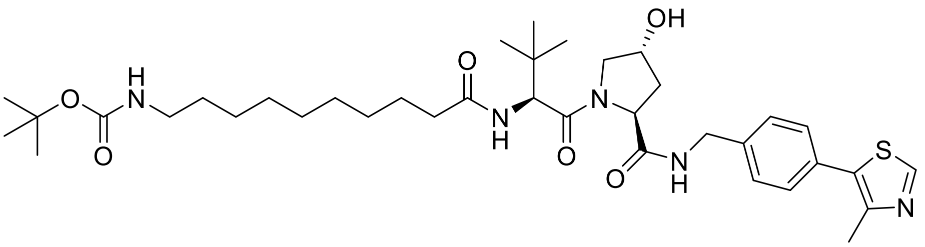

**$^1$H NMR** (400 MHz, DMSO): δ 8.98 (s, 1H), 8.56 (t, *J* = 6.1 Hz, 1H), 7.84 (d, *J* = 9.3 Hz, 1H), 7.40 (q, *J* = 8.2 Hz, 4H), 6.74 (t, *J* = 5.8 Hz, 1H), 5.12 (d, *J* = 3.4 Hz, 1H), 4.54 (d, *J* = 9.3 Hz, 1H), 4.47 – 4.38 (m, 2H), 4.37 – 4.33 (m, 1H), 4.21 (dd, *J* = 15.9, 5.5 Hz, 1H), 3.71 – 3.60 (m, 2H), 2.87 (q, *J* = 6.6 Hz, 2H), 2.44 (s, 3H), 2.31 – 2.18 (m, 1H), 2.15 – 1.98 (m, 2H), 1.95 – 1.84 (m, 1H), 1.56 – 1.39 (m, 2H), 1.36 (s, 9H), 1.22 (s, 12H), 0.93 (s, 9H) ppm. **LCMS** (ESI) *m/z* [M + H] calcd for $C_{37}H_{58}N_4O_5S^+$: 700.4; found: 700.8.

*tert*-butyl (2-(2-(2-(((*S*)-1-((2*S*,4*R*)-4-hydroxy-2-((4-(4-methylthiazol-5-yl)benzyl)carbamoyl)pyrrolidin-1-yl)-3,3-dimethyl-1-oxobutan-2-yl)amino)-2-oxoethoxy)ethoxy)ethyl)carbamate **SI13**

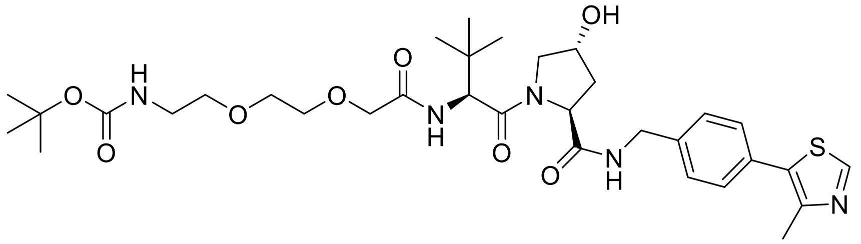

**$^1$H NMR** (400 MHz, DMSO): δ 8.98 (s, 1H), 8.57 (t, *J* = 6.0 Hz, 1H), 7.48 – 7.37 (m, 4H), 6.76 (t, *J* = 5.8 Hz, 1H), 5.15 (d, *J* = 3.4 Hz, 1H), 4.57 (d, *J* = 9.5 Hz, 1H), 4.46 – 4.33 (m, 3H), 4.26 (dd, *J* = 15.8, 5.7 Hz, 1H), 3.96 (s, 2H), 3.69 – 3.50 (m, 7H), 3.40 (dt, *J* = 6.9, 3.8 Hz, 2H), 3.08 (q, *J* = 6.0 Hz, 2H), 2.44 (s, 3H), 2.11 – 2.01 (m, 1H), 1.96 – 1.85 (m, 1H), 1.35 (s, 9H), 0.94 (s, 9H) ppm. **LCMS** (ESI) *m/z* [M + H] calcd for $C_{33}H_{50}N_4O_8S^+$: 676.3; found: 676.8.

*tert*-butyl ((*S*)-17-((2*S*,4*R*)-4-hydroxy-2-((4-(4-methylthiazol-5-yl)benzyl)carbamoyl)pyrrolidine-1-carbonyl)-18,18-dimethyl-15-oxo-3,6,9,12-tetraoxa-16-azanonadecyl)carbamate **SI14**

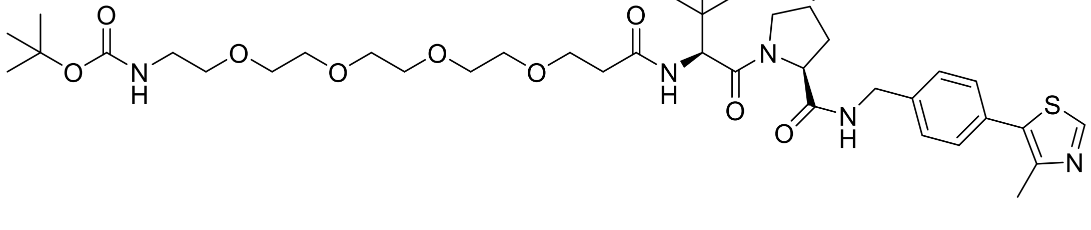

**$^1$H NMR** (400 MHz, DMSO): δ 8.98 (s, 1H), 8.56 (t, *J* = 6.1 Hz, 1H), 7.92 (d, *J* = 9.4 Hz, 1H), 7.40 (q, *J* = 8.4 Hz, 4H), 6.74 (t, *J* = 5.8 Hz, 1H), 5.13 (d, *J* = 3.5 Hz, 1H), 4.55 (d, *J* = 9.4 Hz, 1H), 4.47 – 4.38 (m, 2H), 4.38 – 4.31 (m, 1H), 4.21 (dd, *J* = 15.9, 5.5 Hz, 1H), 3.72 – 3.54 (m, 4H), 3.49 (d, *J* = 3.1 Hz, 12H), 3.36 (t, *J* = 6.1 Hz, 2H), 3.05 (q, *J* = 6.0 Hz, 2H), 2.60 – 2.50 (m, 1H), 2.44 (s, 3H), 2.41 – 2.29 (m, 1H), 2.09 – 1.96 (m, 1H), 1.95 – 1.84 (m, 1H), 1.36 (s, 9H), 0.93 (s, 9H) ppm. **LCMS** (ESI) *m/z* [M + H] calcd for $C_{38}H_{60}N_5O_{10}S^+$: 778.4; found: 777.8.

*tert*-butyl (7-(((*S*)-1-((2*S*,4*R*)-4-hydroxy-2-((4-(4-methylthiazol-5-yl)benzyl)carbamoyl)pyrrolidin-1-yl)-3,3-dimethyl-1-oxobutan-2-yl)amino)-7-oxoheptyl)carbamate **SI15**

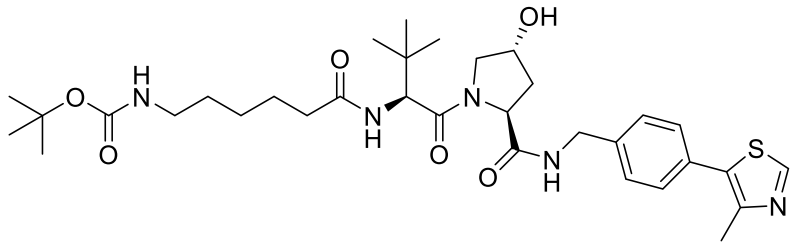

**$^1$H NMR** (400 MHz, DMSO): δ 8.98 (s, 1H), 8.56 (t, *J* = 6.1 Hz, 1H), 7.84 (d, *J* = 9.3 Hz, 1H), 7.40 (q, *J* = 8.3 Hz, 4H), 6.74 (t, *J* = 5.7 Hz, 1H), 5.12 (d, *J* = 3.5 Hz, 1H), 4.54 (d, *J* = 9.3 Hz, 1H), 4.50 – 4.39 (m, 2H), 4.39 – 4.30 (m, 1H), 4.21 (dd, *J* = 15.9, 5.4 Hz, 1H), 3.71 – 3.60 (m, 2H), 2.87 (q, *J* = 6.6 Hz, 2H), 2.44 (s, 3H), 2.30 – 2.18 (m, 1H), 2.16 – 1.96 (m, 2H), 1.95 – 1.84 (m, 1H), 1.55 – 1.39 (m, 2H), 1.36 (s, 12H), 1.26 – 1.12 (m, 3H), 0.93 (s, 9H) ppm. **LCMS** (ESI) *m/z* [M + H] calcd for $C_{34}H_{52}N_5O_6S^+$: 657.4; found: 657.4.

**Hydrophobic tags**

*tert*-butyl (2-(2-(2-(2-((3r,5r,7r)-adamantan-1-yl)acetamido)ethoxy)ethoxy)ethyl)carbamate **SI7**

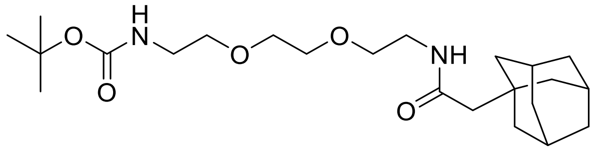

**$^1$H NMR** (400 MHz, $CDCl_3$): δ 5.90 (s, 1H), 5.02 (s, 1H), 3.69 – 3.54 (m, 8H), 3.48 (q, *J* = 5.1 Hz, 2H), 3.34 (s, 2H), 2.02 – 1.95 (m, 5H), 1.76 – 1.61 (m, 12H), 1.48 (s, 9H) ppm. **LCMS** (ESI) *m/z* [M + H] calcd for $C_{23}H_{41}N_2O_5^+$: 425.3; found: 425.3.

*tert*-butyl (3-(4-(3-(2-((3r,5r,7r)-adamantan-1-yl)acetamido)propoxy)butoxy)propyl)carbamate **SI8**

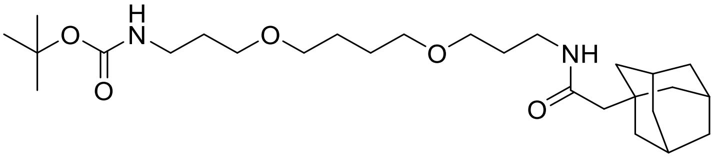

The adamantane-carbaxylic acid (1.0 equiv., 1.0 mmol) was dissolved in DCM (2 mL) and EDC x HCl (3.0 equiv.) and HOBt x $H_2O$ (1.2 equiv.) were added. After stirring for 5 min, the respective mono-Boc protected linker (1.0 equiv., 1.0 mmol) and DIPEA (3.0 equiv.) were dissolved in DCM (4 mL) and added to the ligand. The combined mixture was stirred at rt overnight. After the reaction was finished as indicated by TLC (DCM/MeOH 9:1), the mixture was diluted with water (10 ml) and the organics were extracted with DCM (3 x 10 ml). The combined organic layers were washed with brine, dried over $Na_2SO_4$ and concentrated under reduced pressure. The resulting crude was purified by flash silica gel column chromatography (DCM/MeOH 98:2 to 95:5) to give the title compound as a colorless oil (344 mg, 70%). **Rf** = 0.57 (DCM/MeOH 9:1). **$^1$H NMR** (400 MHz, DMSO): δ 7.63 (t, *J* = 5.6 Hz, 1H), 6.75 (t, *J* = 5.7 Hz, 1H), 3.53 – 3.43 (m, 1H), 3.41 – 3.31 (m, 7H), 3.05 (q, *J* = 6.6 Hz, 2H), 2.95 (q, *J* = 6.6 Hz, 2H), 1.90 (s, 3H), 1.79 (s, 2H), 1.68 – 1.46 (m, 20H), 1.37 (s, 9H) ppm. **$^{13}$C NMR** (101 MHz, DMSO): δ 170.2, 155.4, 132.7, 70.3, 68.2, 50.6, 42.6, 37.7, 36.9, 36.1, 32.6, 30.2, 30.0, 28.7, 28.5, 26.5 ppm. **LCMS** (ESI) *m/z* [M + H] calcd for $C_{27}H_{49}N_2O_5^+$: 481.4; found: 481.3.

*tert*-butyl (1-((3r,5r,7r)-adamantan-1-yl)-2-oxo-7,10,13-trioxa-3-azahexadecan-16-yl)carbamate **SI9**

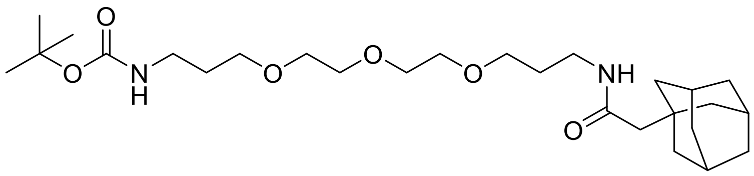


The adamantane-carbaxylic acid (1.0 equiv., 1.0 mmol) was dissolved in DCM (2 mL) and EDC x HCl (3.0 equiv.) and HOBt x $H_2O$ (1.2 equiv.) were added. After stirring for 5 min, the respective mono-Boc protected linker (1.0 equiv., 1.0 mmol) and DIPEA (3.0 equiv.) were dissolved in DCM (4 mL) and added to the ligand. The combined mixture was stirred at rt overnight. After the reaction was finished as indicated by TLC (DCM/MeOH 9:1), the mixture was diluted with water (10 ml) and the organics were extracted with DCM (3 x 10 ml). The combined organic layers were washed with brine, dried over $Na_2SO_4$ and concentrated under reduced pressure. The resulting crude was purified by flash silica gel column chromatography (DCM/MeOH 98:2 to 95:5) to give the title compound as a colorless oil (432 mg, 85%). **Rf** = 0.57 (DCM/MeOH 9:1). **$^1$H NMR** (400 MHz, DMSO): δ 7.63 (t, *J* = 5.7 Hz, 1H), 6.75 (t, *J* = 5.7 Hz, 1H), 3.54 – 3.42 (m, 7H), 3.38 (q, *J* = 6.6 Hz, 4H), 3.05 (q, *J* = 6.5 Hz, 2H), 2.95 (q, *J* = 6.6 Hz, 2H), 1.93 – 1.87 (m, 3H), 1.80 (s, 2H), 1.69 – 1.51 (m, 17H), 1.37 (s, 9H) ppm. **$^{13}$C NMR** (101 MHz, DMSO): δ 170.2, 156.0, 77.8, 70.2, 70.0, 70.0, 68.6, 68.6, 50.5, 42.6, 37.7, 36.9, 36.1, 32.6, 30.2, 30.0, 28.7, 28.5 ppm. **HRMS** (ESI) *m/z* [M + Na] calcd for $C_{27}H_{49}N_2O_7Na^+$: 496.3512; found: 519.3408.

*tert*-butyl (6-(2-((3r,5r,7r)-adamantan-1-yl)acetamido)hexyl)carbamate **SI10**

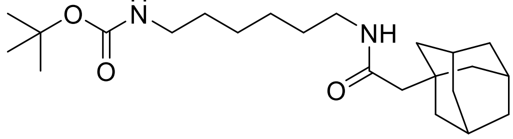


**$^1$H NMR** (400 MHz, DMSO): δ 7.61 (t, *J* = 5.6 Hz, 1H), 6.75 (t, *J* = 5.7 Hz, 1H), 2.99 (q, *J* = 6.7 Hz, 2H), 2.88 (q, *J* = 6.6 Hz, 2H), 1.93 – 1.87 (m, 3H), 1.79 (s, 2H), 1.68 – 1.61 (m, 3H), 1.61 – 1.51 (m, 9H), 1.36 (s, 9H), 1.37 – 1.30 (m, 4H), 1.29 – 1.15 (m, 4H) ppm. **LCMS** (ESI) *m/z* [M + H] calcd for $C_{23}H_{41}N_2O_3^+$: 393.3; found: 393.3.

**SQSI Ligands**

SQSI was synthesized following a previously reported procedure (SI Fig.1).[4]

(2-amino-5-chlorophenyl)(2-chlorophenyl)methanol **SI13**

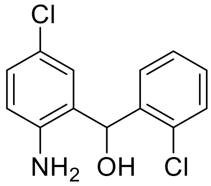


**Rf** = 0.51 (hexane/EtOAc); **$^{1}$H NMR** (400 MHz, $CDCl_3$): δ 7.47 (dd, *J* = 7.0, 2.4 Hz, 1H), 7.41 (dd, *J* = 7.0, 2.2 Hz, 1H), 7.37 – 7.23 (m, 2H), 7.07 (dd, *J* = 8.5, 2.5 Hz, 1H), 6.89 (d, *J* = 2.5 Hz, 1H), 6.64 (d, *J* = 8.4 Hz, 1H), 6.15 (s, 1H) ppm. OH and NH2 peaks missing due to proton-deuterium exchange.; **$^{13}$C NMR** (101 MHz, $CDCl_3$): δ 143.4, 138.4, 133.0, 129.8, 129.5, 128.8, 128.6, 127.6, 127.4, 127.3, 123.2, 117.8, 70.4 ppm. **LCMS** (ESI): *m/z* $[M + H]^+$ calcd for $C_{13}H_{12}Cl_2O^+$: 268.0; found: 268.4.

(5-chloro-2-(neopentylamino)phenyl)(2-chlorophenyl)methanol **SI14**

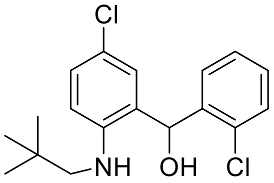


**Rf** = 0.54 (hexane/EtOAc 9:1); **$^{1}$H NMR** (400 MHz, $CDCl_3$): δ 7.44 (dt, *J* = 5.7, 3.9 Hz, 2H), 7.37 – 7.27 (m, 2H), 7.18 (dd, *J* = 8.7, 2.5 Hz, 1H), 6.99 (d, *J* = 2.5 Hz, 1H), 6.67 (d, *J* = 8.7 Hz, 1H), 6.18 (s, 1H), 2.87 (s, 2H), 0.96 (s, 9H) ppm. OH and NH peaks missing due to proton-deuterium exchange.; **$^{13}$C NMR** (101 MHz, $CDCl_3$): δ 145.3, 138.5, 133.4, 130.0, 129.8, 129.0, 128.9, 127.5, 127.5, 126.9, 121.6, 112.5, 70.8, 56.1, 31.7, 27.8 ppm. **LCMS** (ESI): *m/z* $[M + H]^+$ calcd for $C_{13}H_{12}Cl_2O^+$: 338.1; found: 338.2.

Methyl 4-((4-chloro-2-((2-chlorophenyl)(hydroxy)methyl)phenyl)(neopentyl)amino)-4-oxobutanoate **SI15**

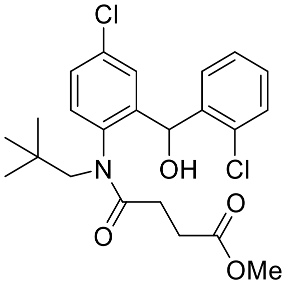


**Rf** = 0.51 (hexane/EtOAc 2:1); **$^{1}$H NMR** (400 MHz, $CDCl_3$): δ 7.81 (dd, *J* = 7.5, 2.0 Hz, 1H), 7.39 (d, *J* = 7.7 Hz, 2H), 7.39 – 7.17 (m, 3H), 7.03 (d, *J* = 1.9 Hz, 1H), 6.15 (6.34) (s, 1H), 4.47 (4.53) (d, *J* = 13.6 Hz, 1H), 3.68 (3.58) (s, 3H), 3.06 (3.02) (d, *J* = 13.7 Hz, 1H), 2.94 – 2.79

(m, 2H), 2.47 – 2.37 (m, 2H), 2.31 – 2.17 (1.83) (m, 1H), 0.89 (0.91) (s, 9H) ppm. **LCMS** (ESI): *m/z* $[M + H]^+$ calcd for $C_{23}H_{28}Cl_2NO_4^+$: 452.1; found: 452.6.

4-((4-chloro-2-((2-chlorophenyl)(hydroxy)methyl)phenyl)(neopentyl)amino)-4-oxobutanoic acid **SI16**

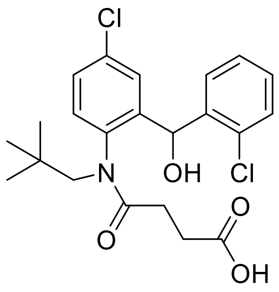


**$^1$H NMR** (400 MHz, DMSO): δ 11.99 (s, 1H), 7.61 – 7.24 (m, 6H), 7.06 (d, *J* = 2.5 Hz, 1H), 6.08 (6.23) (d, *J* = 4.7 Hz, 1H), 5.88 (6.18) (s, 1H), 4.40 (4.20) (d, *J* = 13.6 Hz, 1H), 3.03 (2.52) (d, *J* = 13.7 Hz, 1H), 2.49 – 2.39 (m, 1H), 2.39 – 2.09 (m, 3H), 2.08 – 2.00 (m, 1H), 1.76 – 1.63 (m, 1H), 0.76 (0.83) (s, 9H) ppm. **LCMS** (ESI): *m/z* $[M + H]^+$ calcd for $C_{22}H_{26}Cl_2NO_4^+$: 438.1; found: 438.6.

Ethyl 1-(4-((4-chloro-2-((2-chlorophenyl)(hydroxy)methyl)phenyl)(neopentyl)amino)-4-oxobutanoyl)piperidine-4-carboxylate **SI17**

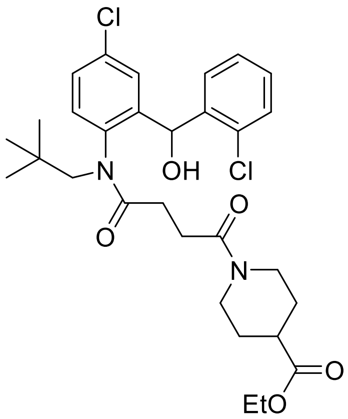


**$R_f$** = 0.33 (DCM/MeOH 99:1); **$^1$H NMR** (400 MHz, DMSO): δ 7.65 – 7.52 (m, 2H), 7.51 – 7.32 (m, 3H), 7.31 – 7.24 (m, 1H), 7.03 (dd, *J* = 4.1, 2.5 Hz, 1H), 6.21 (d, *J* = 5.7 Hz, 1H), 5.90 (6.09) (d, *J* = 5.8 Hz, 1H), 4.40 (4.21) (d, *J* = 13.6 Hz, 1H), 4.18 – 4.12 (m, 1H), 4.07 (q, *J* = 7.1 Hz, 2H), 3.79 (3.61) (dd, *J* = 13.8, 3.9 Hz, 1H), 3.17 (d, *J* = 5.2, 1H), 3.12 – 2.96 (m, 1H), 2.74 – 2.53 (m, 3H), 2.46 – 2.28 (m, 2H), 2.07 – 1.95 (m, 1H), 1.88 – 1.67 (m, 2H), 1.54 – 1.22 (1.00) (m, 2H), 1.18 (1.05) (t, *J* = 7.1 Hz, 3H), 0.76 (0.83) (s, 9H) ppm. **LCMS** (ESI): *m/z* $[M + H]^+$ calcd for $C_{30}H_{39}Cl_2N_2O_5^+$: 577.2; found: 577.7.

1-(4-((4-chloro-2-((2-chlorophenyl)(hydroxy)methyl)phenyl)(neopentyl)amino)-4-oxobutanoyl)piperidine-4-carboxylic acid **SI18**

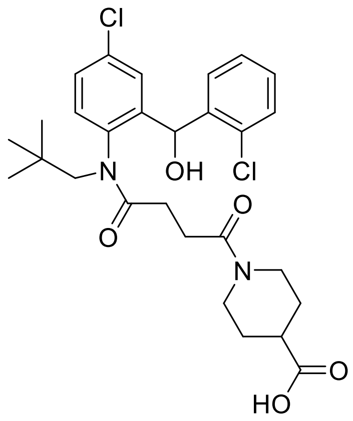

**R$_f$** = 0.01 (DCM/MeOH 99:1); **$^1$H NMR** (400 MHz, DMSO): δ 12.25 (s, 1H), 7.65 – 7.52 (m, 2H), 7.51 – 7.34 (m, 4H), 7.28 (q, *J* = 4.2 Hz, 1H), 7.06 – 7.01 (m, 1H), 6.22 (d, *J* = 5.8 Hz, 1H), 6.10 (s, 1H), 5.93 – 5.87 (m, 1H), 4.40 (d, *J* = 13.6 Hz, 1H), 4.21 (d, *J* = 13.7 Hz, 1H), 4.16 (d, *J* = 13.4 Hz, 1H), 4.12 – 4.04 (m, 1H), 3.78 (d, *J* = 13.6 Hz, 1H), 3.60 (d, *J* = 13.7 Hz, 1H), 3.12 – 2.91 (m, 1H), 2.76 – 2.52 (m, 3H), 2.49 – 2.28 (m, 2H), 2.17 (dt, *J* = 15.8, 7.6 Hz, 1H), 2.08 – 1.94 (m, 1H), 1.87 – 1.67 (m, 3H), 1.55 – 1.20 (m, 2H), 1.04 – 0.93 (m, 1H), 0.83 (s, 3H), 0.76 (s, 6H) ppm. **LCMS** (ESI): *m/z* [M + H]$^+$ calcd for $C_{28}H_{35}Cl_2N_2O_5^+$: 549.2; found: 549.1.

1-(4-((4-chloro-2-((2-chlorophenyl)(hydroxy)methyl)phenyl)(neopentyl)amino)-4-oxobutanoyl)-N-methylpiperidine-4-carboxamide **(SQSI)**

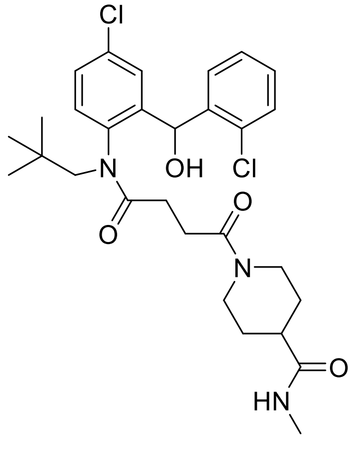

**R$_f$** = 0.59 (DCM/MeOH 95:5); NMR assignment can be found in SI Fig. 3. **HRMS** (ESI): *m/z* [M + Na]$^+$ calcd for $C_{29}H_{37}Cl_2N_3NaO_4^+$: 584.2054; found: 584.2056.

Methyl 3-(4-aminophenyl)propanoate **SI19** [5>

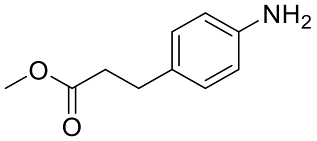

**$^1$H NMR** (400 MHz, $CDCl_3$): δ 7.02 – 6.95 (m, 2H), 6.66 – 6.59 (m, 2H), 3.66 (s, 3H), 3.51 ( br s, 2H), 2.84 (t, *J* = 7.8 Hz, 2H), 2.60 – 2.55 (m, 2H). **$^{13}$C NMR** (101 MHz, $CDCl_3$): δ 173.7, 144.6, 130.7, 129.2, 115.5, 51.7, 36.3, 30.3 ppm. **LCMS** (ESI): *m/z* $[M + H]^+$ calcd for $C_{10}H_{14}NO_2^+$: 180.1; found: 179.9.

Methyl 3-(2-aminobenzo[*d*]thiazol-6-yl)propanoate **SI20**

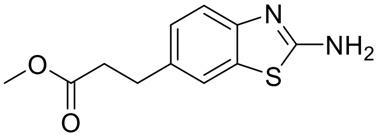

**$^1$H NMR** (400 MHz, $CDCl_3$): δ 7.46 (d, *J* = 8.3 Hz, 1H), 7.43 (d, *J* = 1.8 Hz, 1H), 7.17 (dd, *J* = 8.2, 1.8 Hz, 1H), 3.67 (s, 3H), 3.00 (t, *J* = 7.6 Hz, 2H), 2.65 (t, *J* = 7.7 Hz, 2H) ppm. **$^{13}$C NMR** (101 MHz, $CDCl_3$): δ 173.5, 166.6, 159.3, 147.0, 136.0, 127.2, 121.1, 118.4, 51.9, 36.0, 30.8 ppm. **LCMS** (ESI): *m/z* $[M + H]^+$ calcd for $C_{11}H_{16}N_2O_2S^+$: 237.1; found: 236.9.

Methyl 3-(2-(3-(3,4-dimethoxyphenyl)propanamido)benzo[*d*]thiazol-6-yl)propanoate **SI21**

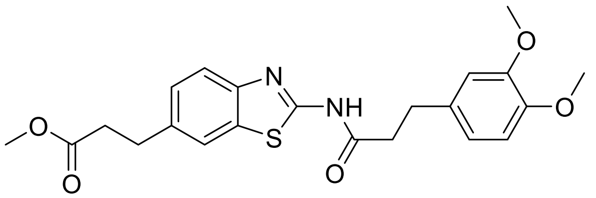

**$^1$H NMR** (400 MHz, DMSO): δ 12.28 (s, 1H), 7.80 (d, *J* = 1.7 Hz, 1H), 7.62 (d, *J* = 8.3 Hz, 1H), 7.29 (dd, *J* = 8.3, 1.8 Hz, 1H), 6.88 – 6.81 (m, 2H), 6.74 (dd, *J* = 8.2, 2.1 Hz, 1H), 3.72 (s, 3H), 3.70 (s, 3H), 3.58 (s, 3H), 2.95 (t, *J* = 7.6 Hz, 2H), 2.88 (t, *J* = 7.2 Hz, 2H), 2.80 – 2.74 (m, 2H), 2.68 (t, *J* = 7.6 Hz, 2H) ppm. **$^{13}$C NMR** (101 MHz, DMSO): δ 172.6, 171.5, 157.3, 148.6, 147.2, 147.0, 136.1, 133.1, 131.6, 126.7, 120.8, 120.2, 119.9, 112.2, 111.9, 55.5, 55.4, 51.3, 37.1, 35.2, 30.2, 30.0 ppm. **LCMS** (ESI): *m/z* $[M + H]^+$ calcd for $C_{22}H_{25}N_2O_5S^+$: 429.2; found: 429.0.

3-(2-(3-(3,4-dimethoxyphenyl)propanamido)benzo[*d*]thiazol-6-yl)propanoic acid **SI22**

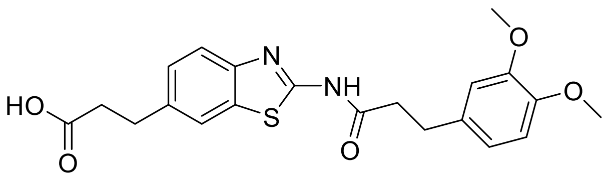

**$^1$H NMR** (400 MHz, DMSO): δ 12.27 (s, 1H), 12.13 (s, 1H), 7.80 (d, *J* = 1.8 Hz, 1H), 7.62 (d, *J* = 8.3 Hz, 1H), 7.29 (dd, *J* = 8.3, 1.8 Hz, 1H), 6.88 – 6.80 (m, 2H), 6.74 (dd, *J* = 8.2, 2.0 Hz, 1H), 3.72 (s, 3H), 3.70 (s, 3H), 2.90 (dt, *J* = 15.6, 7.5 Hz, 4H), 2.81 – 2.73 (m, 2H), 2.58 (t, *J* = 7.6 Hz, 2H) ppm. **$^{13}$C NMR** (101 MHz, DMSO): δ 173.7, 171.5, 157.3, 148.6, 147.2, 146.9, 136.5, 133.1, 131.5, 126.7, 120.8, 120.2, 120.0, 112.2, 111.9, 55.5, 55.4, 37.2, 35.5, 30.3, 30.0 ppm. **LCMS** (ESI): *m/z* $[M + H]^+$ calcd for $C_{21}H_{23}N_2O_5S^+$: 415.1; found: 415.0.

*N*-butyl-3-(2-(3-(3,4-dimethoxyphenyl)propanamido)benzo[d]thiazol-6-yl)propanamide **SO2093**[5]

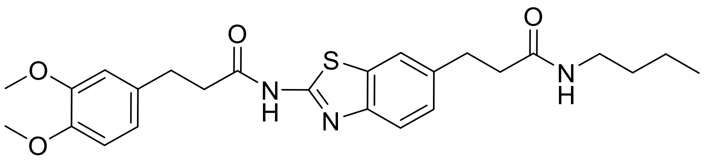

**R$_f$** = 0.63 (DCM/MeOH 9:1); **$^1$H NMR** (400 MHz, DMSO): δ 12.26 (s, 1H), 7.78 – 7.71 (m, 2H), 7.61 (d, *J* = 8.3 Hz, 1H), 7.25 (dd, *J* = 8.3, 1.8 Hz, 1H), 6.88 – 6.81 (m, 2H), 6.74 (dd, *J* = 8.2, 2.0 Hz, 1H), 3.71 (s, 3H), 3.69 (s, 3H), 3.00 (td, *J* = 6.9, 5.6 Hz, 2H), 2.89 (dt, *J* = 9.9, 7.6 Hz, 4H), 2.83 – 2.72 (m, 2H), 2.39 (dd, *J* = 8.3, 6.9 Hz, 2H), 1.35 – 1.24 (m, 2H), 1.22 – 1.12 (m, 2H), 0.80 (t, *J* = 7.3 Hz, 3H) ppm. **$^{13}$C NMR** (101 MHz, DMSO): δ 171.5, 171.0, 157.2, 148.6, 147.2, 146.8, 136.9, 133.1, 131.5, 126.7, 120.7, 120.1, 120.0, 112.2, 111.9, 55.5, 55.4, 38.0, 37.3, 37.2, 31.2, 31.1, 30.0, 19.5, 13.6 ppm. **HRMS (ESI):** m/z [M + H]$^+$ calcd for $C_{25}H_{31}N_3O_4S^+$ 470.2113; found 470.2114.

*N*-(6-chlorobenzo[d]thiazol-2-yl)-3-(3,4-dimethoxyphenyl)propanamide **KY02111**[5]

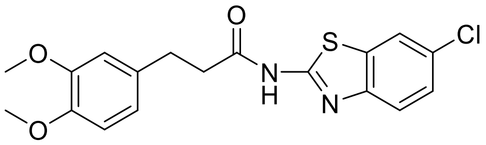

**Rf** = 0.50 (Hex/EtOAc 1:1); **$^1$H NMR** (400 MHz, DMSO): δ 12.43 (s, 1H), 8.11 (d, *J* = 2.2 Hz, 1H), 7.72 (d, *J* = 8.6 Hz, 1H), 7.44 (dd, *J* = 8.6, 2.2 Hz, 1H), 6.88 – 6.81 (m, 2H), 6.74 (dd, *J* = 8.2, 2.0 Hz, 1H), 3.72 (s, 3H), 3.70 (s, 3H), 2.92 – 2.84 (m, 2H), 2.82 – 2.76 (m, 2H) ppm. **$^{13}$C NMR** (101 MHz, DMSO): δ 171.8, 158.7, 148.6, 147.4, 147.2, 133.1, 133.0, 127.5, 126.4, 121.7, 121.4, 120.0, 112.2, 111.9, 55.5, 55.4, 37.1, 29.9 ppm. **HRMS (ESI):** m/z [M + H]$^+$ calcd for $C_{18}H_{18}ClN_2O_3S^+$ 377.0721; found 377.0718.

**SQS degrader library**

**General Procedure A** for the amide coupling of linker-pomalidomide conjugate to the SQS-ligands:

The Boc-protected linker-pomalidomide conjugate (1.1 equiv.) was dissolved in DCM (2 mL/0.1mmol) and treated with trifluoroacetic acid (0.75 mL). The reaction mixture was stirred for 30min to 2h at rt. After the reaction was completed as confirmed by TLC ($Et_2O$/EtOAc 1:1), the reaction was slowly quenched by the addition of sat. $NaHCO_3$ (15 ml). The aqueous solution was extracted with DCM until the organic phase was not fluorescent anymore (ca. 6 × 20 ml). The combined organic phases were dried over $Na_2SO_4$ and the solvent was removed under reduced pressure. The oily residue was further dried in high vacuum. In a separate flask, the respective SQS ligand (1.0 equiv.) was dissolved in DCM (2 mL) and EDC x HCl (3.0 equiv.) and HOBt x $H_2O$ (1.2 equiv.) were added. After stirring for 5 min, the deprotected amine derivative and DIPEA (3.0 equiv.) were dissolved in DCM (4 mL) and added to the ligand. The combined mixture was stirred at rt for 1-18 h. After the reaction was finished as indicated by TLC (DCM/MeOH 9:1), the mixture was diluted with water (10 ml) and the organics were extracted with DCM (3 × 10 ml). The combined organic layers were washed with brine, dried over $Na_2SO_4$ and concentrated under reduced pressure. The resulting crude was purified by flash silica gel column chromatography (DCM/MeOH 98:2 to 95:5) to give the title compound as a yellow amorph solid.

**General Procedure B** for the amide coupling of linker-VH032 and linker-adamantane conjugates to the SQS-ligands:

The Boc-protected linker-VHL or linker-adamantane conjugate (1.2 equiv.) was dissolved in DCM (2 mL/0.1mmol) and treated with 4M HCL in dioxane (1 ml/mmol). The reaction mixture was stirred for 30min to 2h at rt. After the reaction was completed as confirmed by TLC (DCM/MeOH 9:1), the solvent was evaporated in reduced pressure and co-evaporated with $Et_2O$ (3 × 20 mL). The precipitate was further dried in high vacuum. In a separate flask, the respective SQS ligand (1.0 equiv.) was dissolved in DCM (2 mL) and EDC x HCl (3.0 equiv.) and HOBt x $H_2O$ (1.2 equiv.) were added. After stirring for 5 min, the deprotected amine derivative and DIPEA (3.0 equiv.) were dissolved in DCM (4 mL) and added to the ligand. The combined mixture was stirred at rt for 1-18 h. After the reaction was finished as indicated by TLC (DCM/MeOH 9:1), the mixture was diluted with water (10 ml) and the organics were extracted with DCM (3 x 10 ml). The combined organic layers were washed with brine, dried over $Na_2SO_4$ and concentrated under reduced pressure. The resulting crude was purified by flash silica gel column chromatography (DCM/MeOH 98:2 to 95:5) to give the title compound as a white amorph solid.

**Active-site binding compounds**

1-(4-((4-chloro-2-((2-chlorophenyl)(hydroxy)methyl)phenyl)(neopentyl)amino)-4-oxobutanoyl)-N-(2-(2-(2-((2-(2,6-dioxopiperidin-3-yl)-1,3-dioxoisoindolin-4-yl)amino)ethoxy)ethoxy)ethyl)piperidine-4-carboxamide **1**

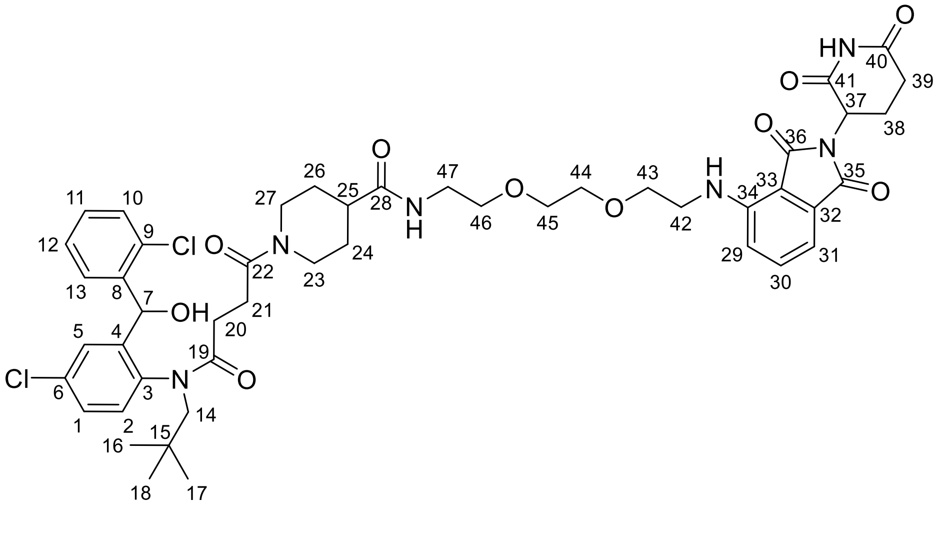

According to general procedure A, SI18 (0.091 mmol) and SI3 (0.14 mmol) provided the title compound as a yellow solid (77.00 mg, 90%). $\mathbf{R_f}$ = 0.48 (DCM/MeOH 95:5); **$^1$H NMR** (400 MHz, DMSO): δ 11.09 (s, 1H, **C40-NH-C41**), 7.82 (t, *J* = 5.9 Hz, 1H), 7.67 – 7.53 (m, 3H), 7.52 – 7.34 (m, 3H), 7.34 – 7.23 (m, 2H), 7.14 (d, *J* = 8.6 Hz, 1H), 7.04 (d, *J* = 7.1 Hz, 1H), 6.61 (t, *J* = 5.8 Hz, 1H), 6.22 (d, *J* = 5.6 Hz, 1H, **OH-8**), 6.10 (t, *J* = 4.3 Hz, 1H, **H7**), 5.90 (t, *J* = 4.9 Hz, 1H, **H7**), 5.06 (dd, *J* = 12.9, 5.4 Hz, 1H, **H37**), 4.40 (d, *J* = 13.6 Hz, 1H), 4.28 (d, *J* = 13.0 Hz, 1H), 4.21 (d, *J* = 13.6 Hz, 1H), 3.83 (d, *J* = 13.6 Hz, 1H), 3.66 – 3.37 (m, 10H), 3.18 (q, *J*

= 5.8 Hz, 2H), 3.05 – 2.80 (m, 2H), 2.64 – 2.51 (m, 4H), 2.50 – 2.28 (m, 3H), 2.23 – 1.99 (m, 2H), 1.62 (q, *J* = 15.1, 14.4 Hz, 2H), 1.50 – 1.21 (m, 2H), 0.83 (s, 4H, **H16-18**), 0.76 (s, 5H, **H16-18**) ppm. **$^{13}$C NMR** (101 MHz, DMSO): δ 173.9, 172.8, 172.3, 171.1, 170.3, 170.1, 169.6 (d, *J* = 7.5 Hz), 169.0 (d, *J* = 7.7 Hz), 167.3, 146.4, 142.5, 142.3, 140.9, 139.8, 139.6, 136.2, 134.1, 133.8, 132.5, 132.3, 132.1, 131.9, 129.6, 129.4 (d, *J* = 7.6 Hz), 129.3, 129.0, 128.4, 128.1, 127.7, 127.4, 117.4, 110.7, 109.3, 69.7, 69.6, 69.1, 68.9, 67.2, 66.1, 58.2, 57.5, 48.6, 44.3, 41.7, 41.6, 40.7, 38.4, 34.2, 33.7, 31.0, 28.8 (**C16-18**), 28.6 (**C16-18**), 28.4, 28.1, 27.9, 22.1 ppm.; **HRMS (ESI):** m/z $[M+H]^+$ calcd for $C_{47}H_{57}Cl_2N_6O_{10}^+$ 935.3508; found 935.3513.

1-(4-((4-chloro-2-((2-chlorophenyl)(hydroxy)methyl)phenyl)(neopentyl)amino)-4-oxobutanoyl)-N-(3-(4-(3-((2-(2,6-dioxopiperidin-3-yl)-1,3-dioxoisoindolin-4-yl)amino)propoxy)butoxy)propyl)piperidine-4-carboxamide **2**

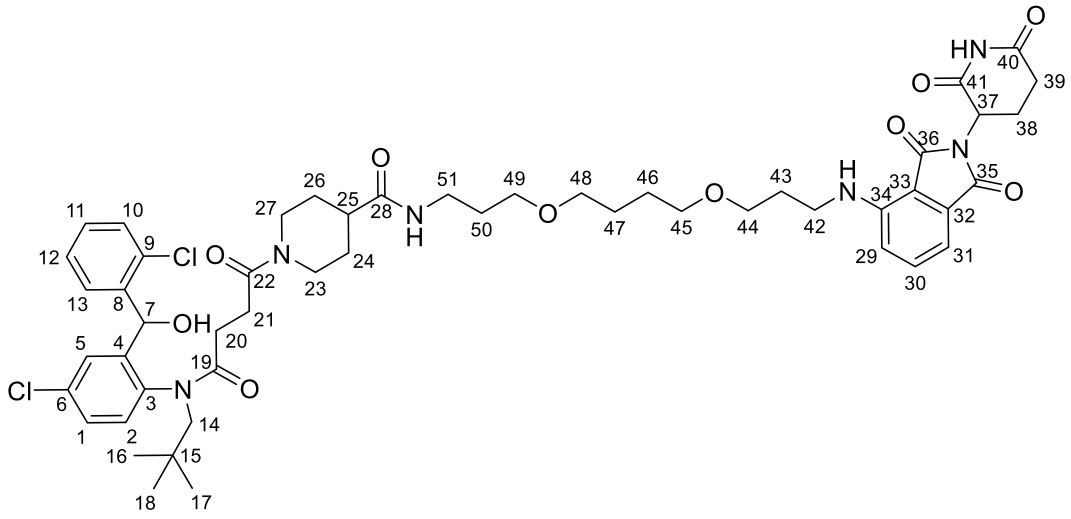

According to general procedure A, SI18 (0.091 mmol) and SI4 (0.14 mmol) provided the title compound as a yellow solid (55.90 mg, 62%). **R*f*** = 0.63 (DCM/MeOH 95:5); **$^{1}$H NMR** (400 MHz, DMSO): δ 11.10 (s, 1H, **C40-NH-C41**), 7.75 (d, *J* = 6.2 Hz, 1H), 7.69 – 7.52 (m, 3H), 7.51 – 7.35 (m, 3H), 7.35 – 7.24 (m, 2H), 7.09 (d, *J* = 8.6 Hz, 1H), 7.03 (d, *J* = 7.1 Hz, 1H), 6.67 (t, *J* = 5.9 Hz, 1H), 6.22 (t, *J* = 5.1 Hz, 1H, **OH-8**), 6.10 (t, *J* = 4.1 Hz, 1H, **H7**), 5.90 (t, *J* = 4.9 Hz, 1H, **H7**), 5.06 (dd, *J* = 12.9, 5.4 Hz, 1H, **H37**), 4.44 – 4.37 (m, 1H), 4.30 (d, *J* = 13.1 Hz, 1H), 4.26 – 4.18 (m, 1H), 3.85 (d, *J* = 13.5 Hz, 1H), 3.66 (d, *J* = 13.8 Hz, 1H), 3.46 (t, *J* = 5.9 Hz, 2H), 3.37 (q, *J* = 5.2 Hz, 8H), 3.06 (p, *J* = 6.1, 5.5 Hz, 2H), 3.03 – 2.95 (m, 1H), 2.89 (ddd, *J* = 17.4, 14.1, 5.4 Hz, 1H), 2.57 (tdd, *J* = 14.4, 5.4, 2.7 Hz, 2H), 2.48 – 2.23 (m, 4H), 2.16 (dt, *J* = 15.9, 7.7 Hz, 1H), 2.03 (dtt, *J* = 7.9, 5.7, 3.5 Hz, 1H), 1.81 (p, *J* = 6.3 Hz, 2H), 1.73 – 1.49 (m, 8H), 1.49 – 1.22 (m, 2H), 0.84 (s, 4H, **H16-18**), 0.76 (s, 5H **H16-18**) ppm. **$^{13}$C NMR** (101 MHz, DMSO): δ 174.1, 173.2, 172.7, 171.9, 170.5, 170.1 (d, *J* = 8.0 Hz), 169.5 (d, *J* = 9.4 Hz), 169.3, 167.8, 146.9, 142.9, 141.4, 140.2, 140.1, 140.0, 136.7, 134.6, 134.2, 132.9, 132.7, 132.4, 130.0, 129.9, 129.8, 129.7, 129.4, 128.9, 128.5, 128.2, 127.8, 117.5, 110.8, 109.5, 70.5, 70.3, 68.3, 68.1, 67.6, 66.5, 58.6, 57.9, 49.0, 44.7, 42.2, 41.2, 36.2, 34.6, 34.2, 31.4, 29.9, 29.3, 29.3, 29.0 (**C16-18**), 28.9 (**C16-18**), 28.6, 28.4, 26.5, 26.4, 22.6 ppm one linker-$CH_2$ group missing due to superimposition with DMSO-d6). **HRMS (ESI):** m/z [M+H]$^{+}$ calcd for $C_{51}H_{65}Cl_2N_6O_{10}^{+}$ 991.4134; found 991.4140.

1-(4-((4-chloro-2-((2-chlorophenyl)(hydroxy)methyl)phenyl)(neopentyl)amino)-4-oxobutanoyl)-N-(3(2-(2-(3-((2-(2,6-dioxopiperidin-3-yl)-1,3-dioxoisoindolin-4-yl)amino)propoxy)ethoxy)ethoxy)propyl)piperidine-4-carboxamide **3**

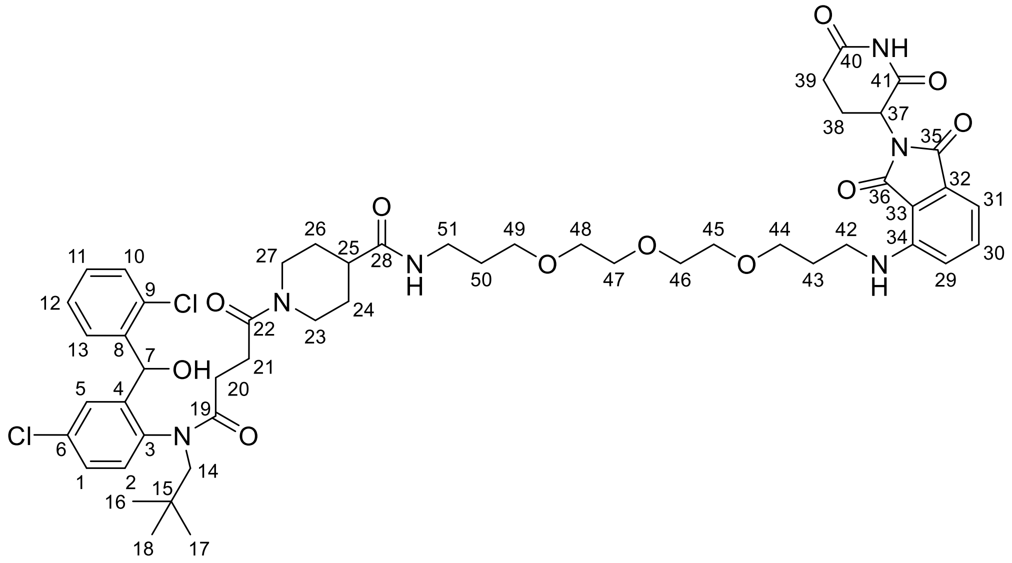

According to general procedure A, SI18 (0.091 mmol) and SI5 (0.14 mmol) provided the title compound as a yellow solid (45.00 mg, 49%). **R*f*** = 0.55 (DCM/MeOH 95:5); **$^1$H NMR** (400 MHz, DMSO) δ 11.09 (s, 1H), 7.79 – 7.70 (m, 1H), 7.67 – 7.54 (m, 3H), 7.52 – 7.35 (m, 2H), 7.30 (d, *J* = 5.4 Hz, 1H), 7.11 (d, *J* = 8.6 Hz, 1H), 7.02 (d, *J* = 7.0 Hz, 1H), 6.71 – 6.64 (m, 1H), 6.26 – 6.19 (m, 1H, **OH-8**), 6.14 – 6.07 (m, 1H, **H7**), 5.91 (d, *J* = 4.9 Hz, 1H, **H7**), 5.11 – 5.01 (m, 1H, **H37**), 4.41 (d, *J* = 14.3 Hz, 1H), 4.30 (d, *J* = 13.0 Hz, 1H), 4.22 (d, *J* = 13.5 Hz, 1H), 3.86 (s, 1H), 3.66 (d, *J* = 12.9 Hz, 1H), 3.60 – 3.44 (m, 10H), 3.37 (tt, *J* = 6.7, 3.4 Hz, 4H), 3.12 – 2.82 (m, 4H), 2.58 (dd, *J* = 13.6, 3.8 Hz, 4H), 2.49 – 2.23 (m, 3H), 2.22 – 2.12 (m, 1H), 2.09 – 1.96 (m, 2H), 1.82 (q, *J* = 6.4 Hz, 2H), 1.77 – 1.55 (m, 5H), 1.52 – 1.21 (m, 2H), 0.84 (s, 4H, **H16-18**), 0.76 (s, 5H, **H16-18**). **$^{13}$C NMR** (101 MHz, DMSO): δ 174.1, 173.3, 172.7, 171.9, 170.5, 169.7, 169.5, 169.3, 167.8, 146.9, 142.9, 141.5, 140.2, 140.1, 136.7, 134.6, 134.2, 132.9, 132.6, 132.4, 130.0, 129.9, 129.8, 129.7, 128.9, 128.5, 128.2, 127.8, 117.5, 110.8, 109.5, 70.2, 70.2, 70.1, 70.0, 68.7, 68.5, 67.6, 66.5, 58.6, 57.9, 49.0, 44.7, 42.2, 41.2, 36.2, 34.6, 34.2, 31.4, 29.8, 29.3, 29.0 (**C16-18**), 28.9 (**C16-18**), 28.6, 28.4, 22.6 ppm (one linker CH2 group missing due to superimposition with DMSO-d6). **HRMS (ESI):** m/z $[M+H]^+$ calcd for $C_{51}H_{65}Cl_2N_6O_{11}^+$ 1007.4083; found 1007.4091.

1-(4-((4-chloro-2-((2-chlorophenyl)(hydroxy)methyl)phenyl)(neopentyl)amino)-4-oxobutanoyl)-N-(6-((2-(2,6-dioxopiperidin-3-yl)-1,3-dioxoisoindolin-4-yl)amino)hexyl)piperidine-4-carboxamide **4**

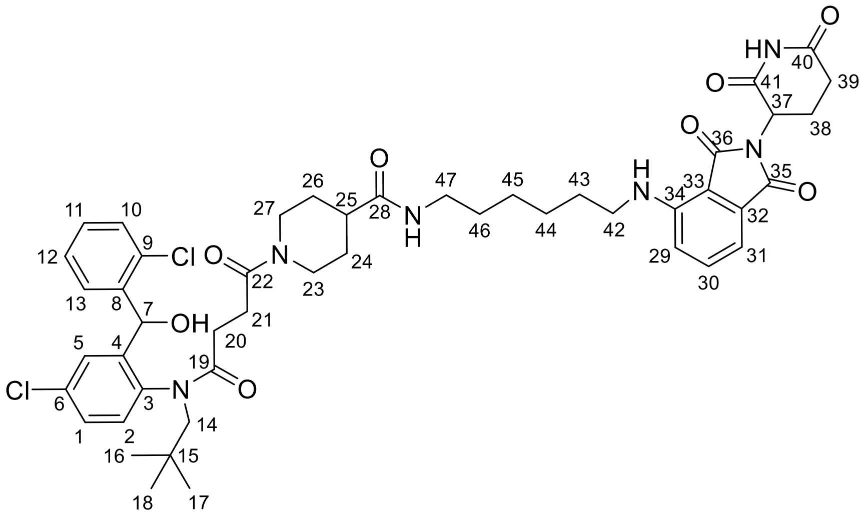

According to general procedure A, SI18 (0.091 mmol) and SI1 (0.14 mmol) provided the title compound as a yellow solid (88.00 mg, 54%). **R*f*** = 0.61 (DCM/MeOH 95:5); **$^1$H NMR** (400 MHz, DMSO): δ 11.10 (s, 1H, **C40-NH-C41**), 7.75 (q, *J* = 7.1, 6.4 Hz, 1H), 7.66 – 7.54 (m, 3H), 7.47 (ddd, *J* = 11.4, 7.5, 1.6 Hz, 2H), 7.43 (s, 1H), 7.35 – 7.25 (m, 2H), 7.09 (d, *J* = 8.6 Hz, 1H), 7.05 – 7.00 (m, 2H), 6.54 (t, *J* = 6.0 Hz, 1H), 6.23 (d, *J* = 5.6 Hz, 1H, **OH-8**), 6.11 (d, *J* = 4.1 Hz, 1H, **H7**), 5.91 (t, *J* = 4.7 Hz, 1H, **H7**), 5.06 (dd, *J* = 12.9, 5.4 Hz, 1H, **H37**), 4.41 (d, *J* = 13.6 Hz, 1H), 4.30 (d, *J* = 13.1 Hz, 1H), 4.22 (d, *J* = 13.6 Hz, 1H), 3.85 (d, *J* = 13.5 Hz, 1H), 3.66 (d, *J* = 13.4 Hz, 1H), 3.29 (q, *J* = 6.7 Hz, 2H), 3.08 – 2.81 (m, 5H), 2.64 – 2.52 (m, 3H), 2.49 – 2.24 (m, 3H), 2.16 (dt, *J* = 15.8, 7.6 Hz, 1H), 2.03 (dtd, *J* = 11.6, 5.4, 4.3, 2.5 Hz, 1H), 1.78 – 1.49 (m, 5H), 1.46 – 1.21 (m, 9H), 0.84 (s, 4H, **H16-18**), 0.76 (s, 5H, **H16-18**) ppm. **$^{13}$C NMR** (101 MHz, DMSO): δ 174.0, 173.3, 172.7, 171.9, 170.6, 170.1, 169.5, 169.4, 167.7, 146.9, 142.9, 141.3, 140.2, 140.1, 136.7, 134.6, 134.2, 132.9, 132.7, 132.6, 132.4, 130.0, 129.9, 129.8, 129.7, 129.4, 128.9, 128.6, 128.2, 127.8, 117.6, 110.8, 109.5, 67.6, 66.5, 58.6, 57.9, 49.0, 44.7, 42.2, 41.2, 38.7, 34.6, 34.2, 31.4, 31.1, 29.5, 29.3, 29.1, 29.0 (**C16-18**), 28.9 (**C16-18**), 28.7, 28.4, 26.5, 26.5, 22.6 ppm. **HRMS (ESI):** m/z $[M+H]^+$ calcd for $C_{47}H_{58}Cl_2N_6O_8^+$ 903.3609; found 903.3614.

1-(4-((4-chloro-2-((2-chlorophenyl)(hydroxy)methyl)phenyl)(neopentyl)amino)-4-oxobutanoyl)-N-(10-((2-(2,6-dioxopiperidin-3-yl)-1,3-dioxoisoindolin-4-yl)amino)decyl)piperidine-4-carboxamide **5**

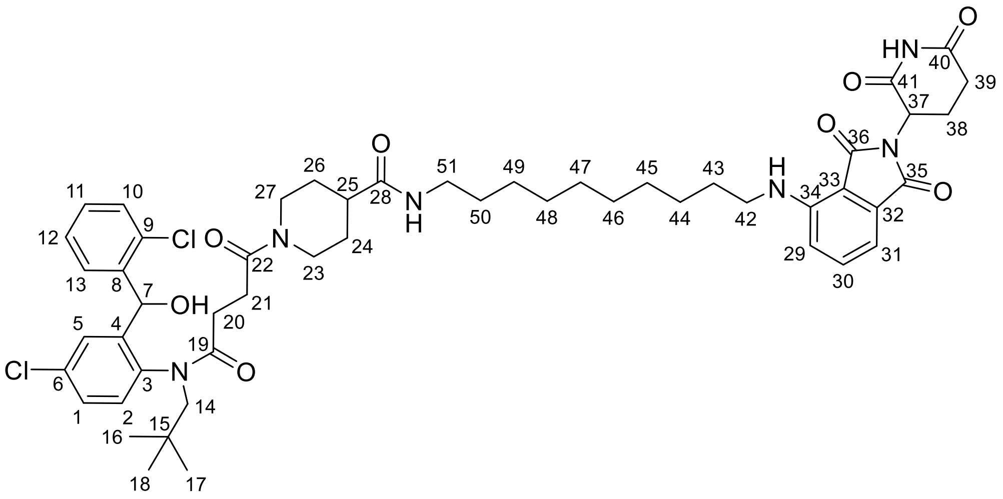

According to general procedure A, SI18 (0.091 mmol) and SI6 (0.14 mmol) provided the title compound as a yellow solid (96.70 mg, 74%). **$R_f$** = 0.60 (DCM/MeOH 95:5); **$^{1}$H NMR** (800 MHz, DMSO): δ 11.09 (s, 1H, **C40-NH-H41**), 7.76 – 7.70 (m, 1H), 7.66 – 7.54 (m, 3H), 7.51 – 7.37 (m, 3H), 7.33 – 7.25 (m, 2H), 7.09 (d, *J* = 8.6 Hz, 1H), 7.06 – 7.01 (m, 1H), 6.52 (t, *J* = 5.9 Hz, 1H), 6.24 – 6.19 (m, 1H, **OH-8**), 6.10 (dd, *J* = 8.0, 5.1 Hz, 1H, **H7**), 5.90 (dd, *J* = 8.2, 5.8 Hz, 1H, **H7**), 5.05 (dd, *J* = 12.9, 5.5 Hz, 1H), 4.41 (dd, *J* = 13.7, 4.3 Hz, 1H), 4.33 – 4.27 (m, 1H), 4.22 (ddd, *J* = 13.6, 10.0, 3.1 Hz, 1H), 3.85 (d, *J* = 13.6 Hz, 1H), 3.66 (d, *J* = 12.9 Hz, 1H), 3.29 (q, *J* = 6.7 Hz, 2H), 3.04 – 2.94 (m, 3H), 2.92 – 2.85 (m, 2H), 2.63 – 2.51 (m, 2H), 2.49 – 2.44 (m, 1H), 2.44 – 2.25 (m, 2H), 2.19 – 2.13 (m, 1H), 2.03 (dtd, *J* = 13.1, 5.4, 2.4 Hz, 1H), 1.74 – 1.54 (m, 5H), 1.50 – 1.19 (m, 16H), 0.84 (s, 5H), 0.76 (s, 4H) ppm. **$^{13}$C NMR** (201 MHz, DMSO) δ 174.1, 173.2, 172.7, 171.9, 170.5, 170.1, 170.1, 169.6, 169.5, 169.4, 167.8, 146.9, 142.9, 142.8, 141.4, 141.3, 140.3, 140.2, 140.1 (d, *J* = 3.4 Hz), 140.0, 136.7, 134.6, 134.2, 132.9 (d, *J* = 3.9 Hz), 132.8, 132.7, 132.6, 132.4, 130.0, 129.9, 129.8, 129.7, 129.4, 128.9, 128.5, 128.2, 127.8, 117.6, 110.8, 109.5, 67.6 (d, *J* = 7.8 Hz), 66.6, 58.6, 57.9, 49.0, 44.8 – 44.6 (m), 42.3, 42.2, 41.3 – 41.0 (m), 38.8, 34.6, 34.2, 31.4, 29.7, 29.5, 29.4, 29.3, 29.2, 29.1, 29.1, 29.0 (**C16-18**), 28.9 (**C16-18**), 28.7, 28.5, 28.4, 26.8, 22.6. **HRMS (ESI):** m/z $[M+H]^+$ calcd for $C_{51}H_{65}Cl_2N_6O_8^+$ 959.4235; found 959.4241.

1-(4-((4-chloro-2-((2-chlorophenyl)(hydroxy)methyl)phenyl)(neopentyl)amino)-4-oxobutanoyl)-N-(2-(2-(2-(((S)-1-((2S,4R)-4-hydroxy-2-((4-(4-methylthiazol-5-yl)benzyl)carbamoyl)-pyrrolidin-1-yl)-3,3-dimethyl-1-oxobutan-2-yl)amino)-2-oxoethoxy)ethoxy)ethyl)piperidine-4-carboxamide **6**

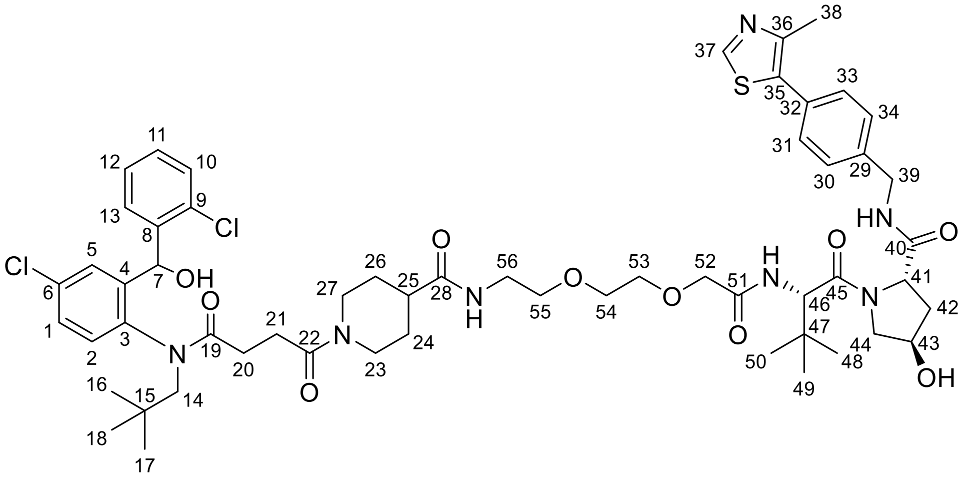

According to general procedure B, SI18 (0.091 mmol) and SI13 (0.14 mmol) provided the title compound as a white solid (114.40 mg, 81%). **Rf** = 0.43 (DCM/MeOH 9:1); **$^{1}$H NMR** (400 MHz, DMSO): δ 8.98 (s, 1H), 8.58 (t, *J* = 6.0 Hz, 1H), 7.89 – 7.80 (m, 1H), 7.67 – 7.52 (m, 2H), 7.51 – 7.41 (m, 3H), 7.39 (s, 5H), 7.32 – 7.24 (m, 2H), 7.03 (d, *J* = 2.5 Hz, 1H), 6.22 (d, *J* = 5.9 Hz, 1H, **OH-8**), 6.10 (s, 1H, **H7**), 5.90 (s, 1H, **H7**), 5.16 (d, *J* = 3.4 Hz, 1H), 4.58 (d, *J* = 9.6 Hz, 1H), 4.50 – 4.33 (m, 4H), 4.31 – 4.17 (m, 2H), 3.97 (s, 2H), 3.70 – 3.50 (m, 7H), 3.43 (t, *J* = 5.8 Hz, 2H), 3.30 – 3.13 (m, 2H), 3.05 – 2.82 (m, 2H), 2.61 – 2.53 (m, 1H), 2.44 (s, 3H), 2.40 – 2.26 (m, 1H), 2.22 – 1.84 (m, 2H), 1.61 (d, *J* = 16.6 Hz, 2H), 1.43 – 1.21 (m, 2H), 0.95 (s, 9H, **H48-H50**), 0.83 (s, 5H, **H16-H18**), 0.75 (s, 4H, **H16-H18**) ppm. **$^{13}$C NMR** (101 MHz, DMSO): δ 173.9, 171.6, 171.4, 169.1, 168.6, 151.4, 147.7, 142.5, 139.7, 139.5, 139.3, 134.0, 132.4, 131.8, 131.1, 129.6, 129.5, 129.4, 129.3, 128.6, 128.1, 127.7, 127.4, 127.3, 70.3, 69.5, 69.3, 69.1, 68.8, 66.0, 58.7, 58.2, 56.6, 55.6, 41.6, 41.5, 40.4, 38.3, 37.9, 35.7, 34.1, 33.7, 29.2 (**C16-18**), 28.8 (**C16-18**), 28.5, 28.3, 28.0, 27.9, 26.1 (**C-48-50**), 15.9 ppm. **HRMS (ESI):** m/z $[M+H]^+$ calcd for $C_{56}H_{74}Cl_2N_7O_{10}S^+$ 1106.4595; found 1106.4605.

1-(4-((4-chloro-2-((2-chlorophenyl)(hydroxy)methyl)phenyl)(neopentyl)amino)-4-oxobutanoyl)-N-(6-(((S)-1-((2S,4R)-4-hydroxy-2-((4-(4-methylthiazol-5-yl)benzyl)carbamoyl)pyrrolidin-1-yl)-3,3-dimethyl-1-oxobutan-2-yl)amino)-6-oxohexyl)piperidine-4-carboxamide **7**

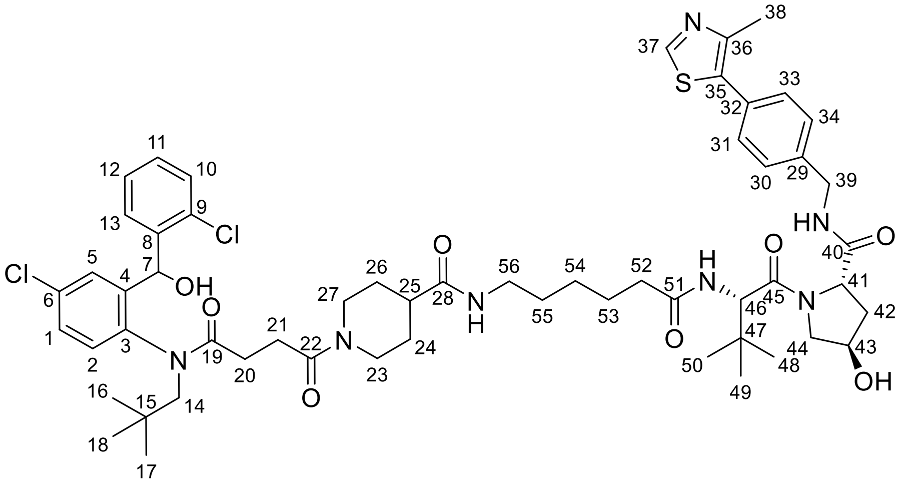

According to general procedure B, SI18 (0.091 mmol) and SI14 (0.14 mmol) provided the title compound as a white solid (98.00 mg, 86%). **Rf** = 0.50 (DCM/MeOH 9:1); **$^1$H NMR** (800 MHz, DMSO): δ 8.99 (s, 1H), 8.56 (t, *J* = 6.1 Hz, 1H), 7.84 (d, *J* = 9.4 Hz, 1H), 7.77 – 7.71 (m, 1H), 7.67 – 7.54 (m, 2H), 7.49 – 7.37 (m, 7H), 7.33 – 7.26 (m, 2H), 7.04 (t, *J* = 3.0 Hz, 1H), 6.24 – 6.20 (m, 1H, **OH-8**), 6.10 (dd, *J* = 8.6, 5.1 Hz, 1H, **H7**), 5.91 (dd, *J* = 8.7, 5.9 Hz, 1H, **H7**), 5.12 (d, *J* = 3.6 Hz, 1H), 4.55 (d, *J* = 9.4 Hz, 1H), 4.46 – 4.39 (m, 3H), 4.36 (h, *J* = 3.7 Hz, 1H), 4.32 – 4.28 (m, 1H), 4.26 – 4.20 (m, 2H), 3.85 (d, *J* = 13.5 Hz, 1H), 3.70 – 3.63 (m, 3H), 3.01 (td, *J* = 13.4, 12.9, 9.1 Hz, 3H), 2.89 (q, *J* = 13.6 Hz, 1H), 2.62 – 2.52 (m, 1H), 2.45 (s, 3H), 2.43 – 2.22 (m, 4H), 2.14 (dddd, *J* = 34.6, 14.4, 8.4, 6.0 Hz, 1H), 2.06 – 2.01 (m, 1H), 1.94 – 1.88 (m, 1H), 1.75 – 1.57 (m, 2H), 1.53 – 1.41 (m, 2H), 1.38 (p, *J* = 7.3 Hz, 2H), 1.34 – 1.18 (m, 3H), 1.05 – 0.97 (m, 1H), 0.94 (s, 9H, **H48-50**), 0.84 (s, 5H **H16-18**), 0.76 (s, 4H, **H16-18**) ppm. **$^{13}$C NMR** (201 MHz, DMSO): δ 174.0, 172.7, 172.5, 172.4, 171.9, 170.2, 170.1, 169.5, 151.9, 148.2, 142.9, 142.8, 141.4, 140.2, 140.1, 140.0, 134.6, 134.2, 132.9, 132.7, 132.4, 131.6, 130.1, 130.0, 129.9, 129.8, 129.7, 129.4, 129.1, 128.9, 128.6, 128.2, 127.9, 127.8, 69.3, 67.6, 66.6, 59.1, 58.6, 57.9, 56.8, 56.7, 55.4, 45.1, 42.2, 42.1, 41.2, 38.7, 38.4, 35.7, 35.3, 34.6, 34.2, 29.8, 29.6, 29.3, 29.3, 29.2, 29.0 (**C16-18**), 28.9 (**C16-18**), 28.7, 28.5, 28.4, 26.8 (**C48-50**), 26.5, 25.6, 16.4 ppm. **HRMS (ESI):** m/z $[M+H]^+$ calcd for $C_{56}H_{74}Cl_2N_7O_8S^+$ 1074.4696; found 1074.4699.

1-(4-((4-chloro-2-((2-chlorophenyl)(hydroxy)methyl)phenyl)(neopentyl)amino)-4-oxobutanoyl)-N-(10-(((S)-1-((2S,4R)-4-hydroxy-2-((4-(4-methylthiazol-5-yl)benzyl)carbamoyl)-pyrrolidin-1-yl)-3,3-dimethyl-1-oxobutan-2-yl)amino)-10-oxodecyl)piperidine-4-carboxamide **8**

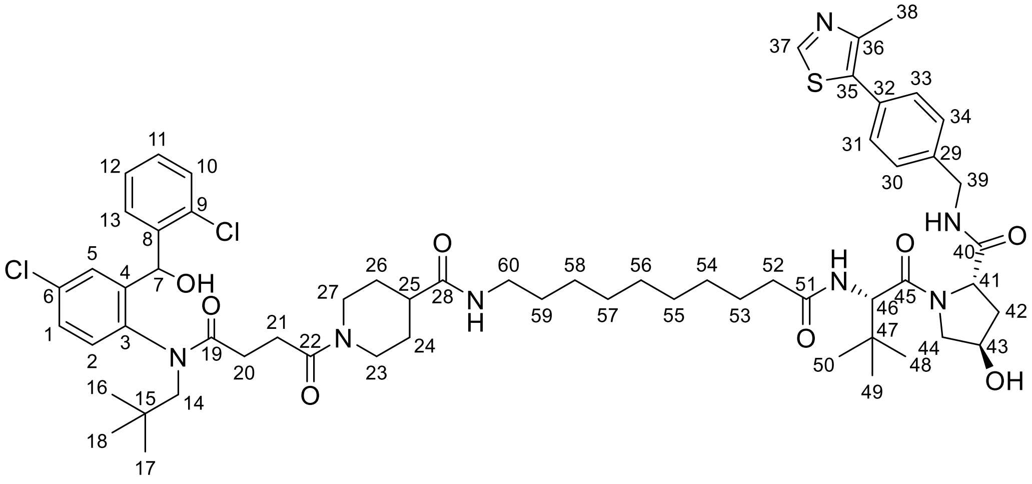

According to general procedure B, SI18 (0.091 mmol) and SI12 (0.14 mmol) provided the title compound as a white solid (94.00 mg, 65%). **Rf** = 0.56 (DCM/MeOH 9:1); **$^{1}$H NMR** (800 MHz, DMSO): δ 8.99 (s, 1H), 8.56 (t, *J* = 6.1 Hz, 1H), 7.83 (d, *J* = 9.4 Hz, 1H), 7.77 – 7.71 (m, 1H), 7.67 – 7.55 (m, 2H), 7.51 – 7.36 (m, 7H), 7.33 – 7.25 (m, 2H), 7.04 (dd, *J* = 4.6, 2.6 Hz, 1H), 6.24 – 6.20 (m, 1H, **OH-8**), 6.11 (dd, *J* = 8.2, 5.1 Hz, 1H, **H7**), 5.91 (dd, *J* = 8.6, 5.9 Hz, 1H, **H7**), 5.12 (d, *J* = 3.6 Hz, 1H), 4.55 (d, *J* = 9.4 Hz, 1H), 4.47 – 4.38 (m, 2H), 4.36 (dtd, *J* = 6.9, 4.3, 2.7 Hz, 1H), 4.32 – 4.28 (m, 1H), 4.26 – 4.19 (m, 2H), 3.85 (d, *J* = 13.5 Hz, 1H), 3.71 – 3.60 (m, 3H), 3.31 (d, *J* = 12.7 Hz, 1H), 3.05 – 2.95 (m, 3H), 2.89 (q, *J* = 13.2, 12.3 Hz, 1H), 2.61 – 2.52 (m, 1H), 2.45 (s, 3H), 2.43 – 2.23 (m, 2H), 2.16 (dtd, *J* = 16.0, 7.8, 3.1 Hz, 1H), 2.11 (ddd, *J* = 14.1, 8.0, 6.1 Hz, 1H), 2.06 – 1.98 (m, 1H), 1.91 (ddd, *J* = 12.9, 8.6, 4.6 Hz, 1H), 1.74 – 1.56 (m, 2H), 1.48 (ddd, *J* = 47.1, 13.3, 6.5 Hz, 1H), 1.42 – 1.27 (m, 2H), 1.26 – 1.21 (m, 13H), 1.04 – 0.97 (m, 2H), 0.94 (s, 9H, **H48-H50**), 0.84 (d, *J* = 1.6 Hz, 5H, **H16-18**), 0.76 (s, 4H, **H16-H18**) ppm. **$^{13}$C NMR** (201 MHz, DMSO): δ 174.1, 172.7, 172.5, 172.4, 172.0, 170.2, 170.1, 169.5, 151.9, 148.2, 142.9, 142.8, 141.4, 140.2, 140.1, 140.0, 134.6, 134.2, 132.9, 132.8, 132.4, 131.6, 130.1, 130.0, 129.9, 129.8, 129.7, 129.4, 129.3, 129.1, 128.9, 128.6, 128.5, 128.2, 127.9, 127.8, 69.3, 67.6, 66.6, 59.1, 58.6, 57.9, 56.8, 56.7, 55.4, 44.5, 42.2, 42.1, 41.3, 38.8, 38.4, 35.7, 35.3, 34.6, 34.2, 29.8, 29.5, 29.3, 29.2, 29.1, 29.0 (**C16-18**), 28.9 (**C16-18**), 28.7, 28.4, 26.8, 26.8 (**C-48-50**), 25.9, 16.4 ppm. **HRMS (ESI):** m/z $[M+H]^+$ calcd for $C_{60}H_{82}Cl_2N_7O_8S^+$ 1130.5322; found 1130.5329.

N-(2-(2-(2-(2-((1s,3s)-adamantan-1-yl)acetamido)ethoxy)ethoxy)ethyl)-1-(4-((4-chloro-2-((2-chlorophenyl)(hydroxy)methyl)phenyl)(neopentyl)amino)-4-oxobutanoyl)piperidine-4-carboxamide **9**

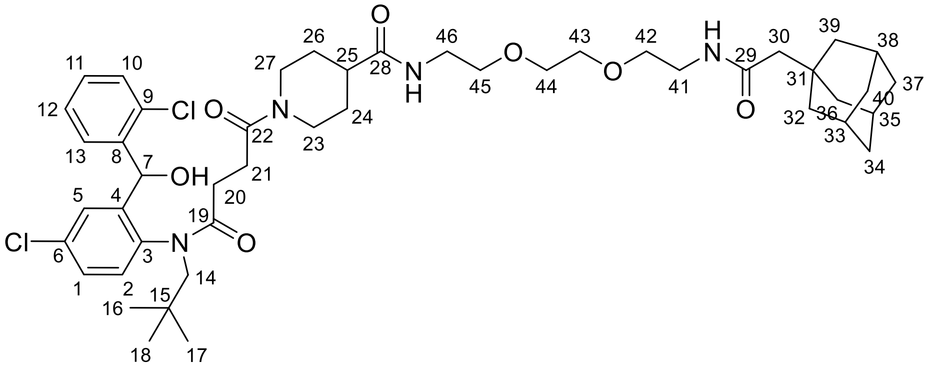

According to general procedure B, SI18 (0.091 mmol) and SI7 (0.14 mmol) provided the title compound as a white solid (61.40 mg, 79%). **Rf** = 0.65 (DCM/MeOH 9:1); **$^1$H NMR** (800 MHz, DMSO): δ 7.87 – 7.82 (m, 1H), 7.73 – 7.54 (m, 3H), 7.51 – 7.36 (m, 3H), 7.29 (dtd, *J* = 12.6, 9.0, 8.5, 4.7 Hz, 2H), 7.04 (dd, *J* = 4.4, 2.5 Hz, 1H), 6.24 – 6.20 (m, 1H, **OH-8**), 6.10 (dd, *J* = 8.3, 5.1 Hz, 1H, **H7**), 5.91 (dd, *J* = 8.7, 5.9 Hz, 1H, **H7**), 4.41 (dd, *J* = 13.7, 4.8 Hz, 1H), 4.33 – 4.28 (m, 1H), 4.22 (d, *J* = 13.5 Hz, 1H), 3.85 (d, *J* = 13.6 Hz, 1H), 3.67 (dd, *J* = 14.2, 4.5 Hz, 1H), 3.50 (s, 4H), 3.39 (td, *J* = 5.9, 2.6 Hz, 4H), 3.30 (s, 1H), 3.18 (dq, *J* = 11.9, 5.9 Hz, 4H), 3.05 – 2.94 (m, 1H), 2.89 (q, *J* = 13.7, 13.0 Hz, 1H), 2.60 – 2.56 (m, 2H), 2.49 – 2.45 (m, 1H), 2.43 – 2.30 (m, 2H), 2.16 (dtd, *J* = 15.8, 7.6, 4.3 Hz, 1H), 2.02 (ddt, *J* = 22.6, 16.2, 6.6 Hz, 1H), 1.90 (q, *J* = 3.2 Hz, 3H), 1.82 (s, 2H), 1.65 (d, *J* = 12.3 Hz, 4H), 1.59 – 1.52 (m, 10H), 1.51 – 1.21 (m, 2H), 1.00 (dddd, *J* = 24.7, 16.4, 8.0, 5.0 Hz, 1H), 0.84 (d, *J* = 2.4 Hz, 5H, **H16-18**), 0.77 (s, 4H, **H18**) ppm. **$^{13}$C NMR** (201 MHz, DMSO): δ 174.4, 172.7, 171.9, 170.4, 170.1, 169.6, 142.9, 142.8, 141.3, 140.2, 140.0, 134.6, 134.2, 132.9, 132.8, 132.7, 132.4, 130.0, 130.0, 129.9, 129.8, 129.7, 129.4, 128.9, 128.6, 128.5, 128.2, 127.8, 127.8, 70.0, 70.0, 69.6, 69.5, 67.6, 66.6, 58.6, 57.9, 50.4, 44.8, 42.5, 42.0, 41.3, 40.4, 38.9, 38.8, 36.9, 34.6, 34.2, 32.6, 29.8, 29.6, 29.3, 29.3, 29.1, 29.0 (**C16-18**), 28.9 (**C16-18**), 28.6, 28.6, 28.5, 28.5, 28.4, 28.4 ppm. **HRMS (ESI):** m/z [M + Na]$^+$ calcd for $C_{46}H_{64}Cl_2N_4O_7Na^+$ 877.4044; found 877.4049.

N-(3-(4-(3-(2-((1s,3s)-adamantan-1-yl)acetamido)propoxy)butoxy)propyl)-1-(4-((4-chloro-2-((2-chlorophenyl)(hydroxy)methyl)phenyl)(neopentyl)amino)-4-oxobutanoyl)piperidine-4-carboxamide **10**

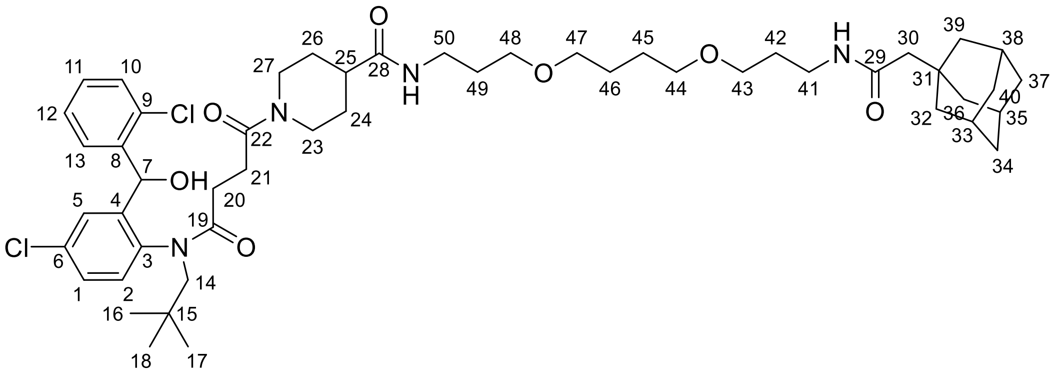

According to general procedure B, SI18 (0.091 mmol) and SI8 (0.14 mmol) provided the title compound as a light brown solid (56.40 mg, 68%). **Rf** = 0.65 (DCM/MeOH 9:1); **$^1$H NMR** (800

MHz, DMSO): δ 7.76 (dt, *J* = 13.2, 6.4 Hz, 1H), 7.68 – 7.56 (m, 3H), 7.52 – 7.37 (m, 3H), 7.34 – 7.25 (m, 2H), 7.04 (dd, *J* = 4.3, 2.5 Hz, 1H), 6.24 – 6.20 (m, 1H, **OH-8**), 6.11 (dd, *J* = 8.2, 5.1 Hz, 1H, **H7**), 5.91 (dd, *J* = 8.5, 5.9 Hz, 1H, **H7**) 4.41 (dd, *J* = 13.7, 5.0 Hz, 1H), 4.30 (dd, *J* = 12.9, 4.1 Hz, 1H), 4.23 (t, *J* = 13.4 Hz, 1H), 3.85 (d, *J* = 13.7 Hz, 1H), 3.67 (d, *J* = 13.1 Hz, 1H), 3.37 – 3.32 (m, 8H), 3.09 – 3.04 (m, 5H), 3.00 (ddd, *J* = 41.4, 13.5, 9.0 Hz, 1H), 2.89 (q, *J* = 13.8, 13.1 Hz, 1H), 2.60 – 2.56 (m, 1H), 2.60 – 2.53 (m, 1H), 2.49 – 2.44 (m, 1H), 2.43 – 2.26 (m, 2H), 2.16 (dtd, *J* = 15.9, 8.0, 2.8 Hz, 1H), 2.02 (tt, *J* = 16.3, 7.0 Hz, 1H), 1.92 – 1.89 (m, 3H), 1.80 (d, *J* = 1.3 Hz, 3H), 1.65 (d, *J* = 11.6 Hz, 5H), 1.63 – 1.53 (m, 19H), 1.51 (q, *J* = 3.0, 2.5 Hz, 5H), 1.43 (dddd, *J* = 48.8, 24.2, 12.0, 3.8 Hz, 1H), 1.34 – 1.21 (m, 2H), 0.84 (d, *J* = 2.1 Hz, 5H, **H16-18**), 0.77 (s, 4H, **H16-18**) ppm. **$^{13}$C NMR** (201 MHz, DMSO): δ 174.2, 172.6, 171.8, 170.2, 170.1, 169.6, 142.9, 142.8, 141.3, 140.3, 140.0, 134.6, 134.2, 132.9, 132.8, 132.4, 130.0, 130.0, 129.9, 129.8, 129.7, 129.4, 128.9, 128.6, 128.5, 128.2, 127.8, 70.3, 70.3, 68.6, 68.2, 68.1, 67.6, 66.6, 58.6, 57.9, 50.5, 44.9, 42.6, 42.2, 41.2, 40.4, 36.9, 36.2, 36.1, 34.6, 34.2, 32.6, 30.0, 29.8, 29.3, 29.2, 29.0 (**C16-18**), 28.9 (**C16-18**), 28.7, 28.6, 28.6, 28.5, 28.4, 26.5 ppm. **HRMS (ESI):** m/z [M + H]$^+$ calcd for $C_{50}H_{73}Cl_2N_4O_7^+$ 911.4851; found 911.4851.

N-(1-((3r,5r,7r)-adamantan-1-yl)-2-oxo-7,10,13-trioxa-3-azahexadecan-16-yl)-1-(4-((4-chloro-2-((2-chlorophenyl)(hydroxy)methyl)phenyl)(neopentyl)amino)-4-oxobutanoyl)piperidine-4-carboxamide **11**

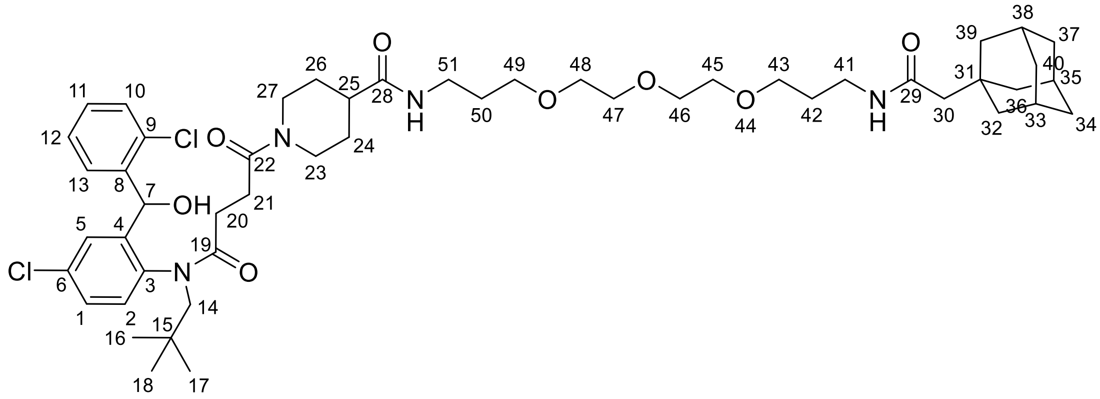


According to general procedure B, SI18 (0.091 mmol) and SI9 (0.14 mmol) provided the title compound as a white solid (59.60 mg, 71%). **Rf** = 0.65 (DCM/MeOH 9:1); **$^1$H NMR** (800 MHz, DMSO): δ 7.79 – 7.73 (m, 1H), 7.67 – 7.54 (m, 3H), 7.50 – 7.39 (m, 3H), 7.33 – 7.25 (m, 2H), 7.04 (dd, *J* = 4.1, 2.5 Hz, 1H), 6.22 (dd, *J* = 9.9, 5.6 Hz, 1H, **OH-8**), 6.10 (dd, *J* = 8.3, 5.0 Hz, 1H, **H7**), 5.91 (dd, *J* = 8.8, 5.9 Hz, 1H, **H7**), 4.41 (dd, *J* = 13.7, 5.0 Hz, 1H), 4.30 (dd, *J* = 13.5, 4.2 Hz, 1H), 4.26 – 4.19 (m, 1H), 3.85 (d, *J* = 13.5 Hz, 1H), 3.67 (d, *J* = 13.4 Hz, 1H), 3.53 – 3.50 (m, 3H), 3.49 – 3.45 (m, 7H), 3.39 (dt, *J* = 8.0, 6.3 Hz, 5H), 3.30 (d, *J* = 2.8 Hz, 1H), 3.06 (dp, *J* = 12.7, 6.5 Hz, 5H), 3.04 – 2.95 (m, 1H), 2.93 – 2.86 (m, 1H), 2.61 – 2.53 (m, 1H), 2.50 – 2.44 (m, 1H), 2.43 – 2.27 (m, 2H), 2.16 (dtd, *J* = 16.1, 7.8, 2.7 Hz, 1H), 2.02 (tt, *J* = 16.6, 6.8 Hz, 1H), 1.90 (q, *J* = 3.1 Hz, 4H), 1.80 (s, 2H), 1.74 – 1.51 (m, 17H), 1.51 – 1.36 (m, 1H), 1.35 – 1.22 (m, 1H), 1.05 – 0.96 (m, 1H), 0.84 (d, *J* = 2.4 Hz, 5H, **H16-18**), 0.77 (s, 4H, **H16-18**) ppm. **$^{13}$C NMR** (201 MHz, DMSO): δ 174.2, 172.7, 171.9, 170.2, 170.1, 169.5, 142.9, 142.8, 141.3, 140.3, 140.0, 134.6, 134.2, 132.9, 132.8, 132.4, 130.0, 129.9, 129.8, 129.7, 129.4, 128.9, 128.6, 128.5, 128.2, 127.8, 127.8, 70.2, 70.2, 70.0, 70.0, 68.6, 68.5, 67.6, 66.6, 58.6, 57.9, 50.5, 44.8, 42.6, 42.2, 41.2, 40.4, 36.9, 36.2, 36.0, 34.6, 34.2, 32.6, 31.4, 29.9, 29.8,

29.3, 29.3, 29.2, 29.0 (**C16-18**), 28.9 (**C16-18**), 28.7, 28.6, 28.6, 28.5, 28.5, 28.4, 22.5 ppm. **HRMS (ESI):** m/z $[M + Na]^+$ calcd for $C_{50}H_{73}Cl_2N_8O_7Na^+$ 949.4619; found 949.4625.

N-(6-(2-((1s,3s)-adamantan-1-yl)acetamido)hexyl)-1-(4-((4-chloro-2-((2-chlorophenyl)(hydroxy)methyl)phenyl)(neopentyl)amino)-4-oxobutanoyl)piperidine-4-carboxamide **12**

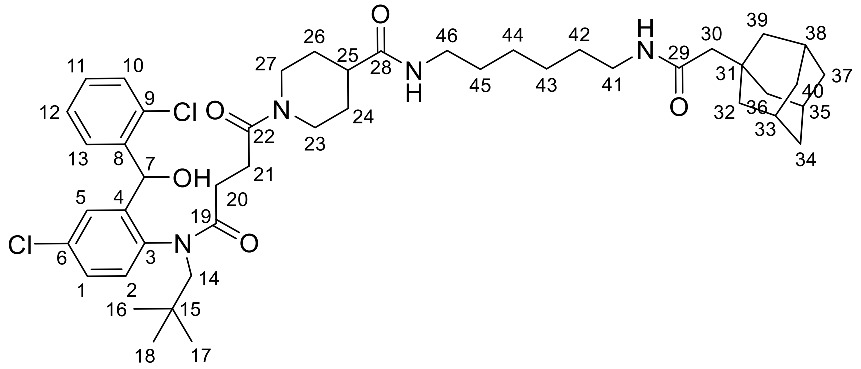

According to general procedure B, SI18 (0.091 mmol) and SI10 (0.14 mmol) provided the title compound as a white solid (65.00 mg, 86%). **Rf** = 0.65 (DCM/MeOH 9:1); **$^1$H NMR** (400 MHz, DMSO): δ 7.75 (q, *J* = 6.2 Hz, 1H), 7.67 – 7.55 (m, 3H), 7.52 – 7.36 (m, 3H), 7.30 (ddt, *J* = 6.5, 5.1, 3.2 Hz, 2H), 7.04 (t, *J* = 2.2 Hz, 1H), 6.22 (t, *J* = 5.0 Hz, 1H, **OH-8**), 6.11 (t, *J* = 4.5 Hz, 1H, **H7**), 5.91 (t, *J* = 4.9 Hz, 1H, **H7**), 4.45 – 4.37 (m, 1H), 4.30 (d, *J* = 13.1 Hz, 1H), 4.26 – 4.18 (m, 1H), 3.85 (d, *J* = 13.6 Hz, 1H), 3.67 (d, *J* = 13.5 Hz, 1H), 3.00 (p, *J* = 6.6 Hz, 5H), 2.89 (q, *J* = 11.8, 9.0 Hz, 1H), 2.58 (d, *J* = 13.5 Hz, 1H), 2.48 – 2.25 (m, 1H), 2.16 (dt, *J* = 15.8, 7.6 Hz, 1H), 2.09 – 1.97 (m, 1H), 1.90 (s, 3H), 1.80 (s, 2H), 1.65 (d, *J* = 12.2 Hz, 5H), 1.58 (s, 3H), 1.56 – 1.50 (m, 7H), 1.36 (p, *J* = 6.8 Hz, 4H), 1.28 – 1.21 (m, 5H), 1.07 – 0.94 (m, 1H), 0.84 (s, 5H, **H16-18**), 0.77 (s, 4H, **H16-18**) ppm. **$^{13}$C NMR** (101 MHz, DMSO): δ 174.0, 172.7, 171.9, 170.1, 169.5, 142.9, 142.5, 141.4, 140.2, 140.0, 134.6, 134.2, 132.9, 132.8, 132.4, 130.0, 129.9, 129.8, 129.7, 129.4, 128.9, 128.5, 128.2, 127.8, 67.6, 66.5, 58.6, 57.9, 50.5, 44.8, 42.6, 42.2, 41.2, 38.7, 38.6, 36.9, 34.6, 34.2, 32.6, 29.6, 29.5, 29.2, 29.0 (**C16-C18**), 28.9 (**C16-C18**), 28.6, 28.5, 26.6, 26.5 ppm. **HRMS (ESI):** m/z $[M + Na]^+$ calcd for $C_{46}H_{65}Cl_2N_4O_5Na^+$ 845.4146; found 845.3151.

5-((2-(2-(2-(1-(4-((4-chloro-2-((2-chlorophenyl)(hydroxy)methyl)phenyl)(neopentyl)amino)-4-oxobutanoyl)piperidine-4-carboxamido)ethoxy)ethoxy)ethyl)carbamoyl)-2-(6-(dimethylamino)-3-(dimethyliminio)-3H-xanthen-9-yl)benzoate **26**

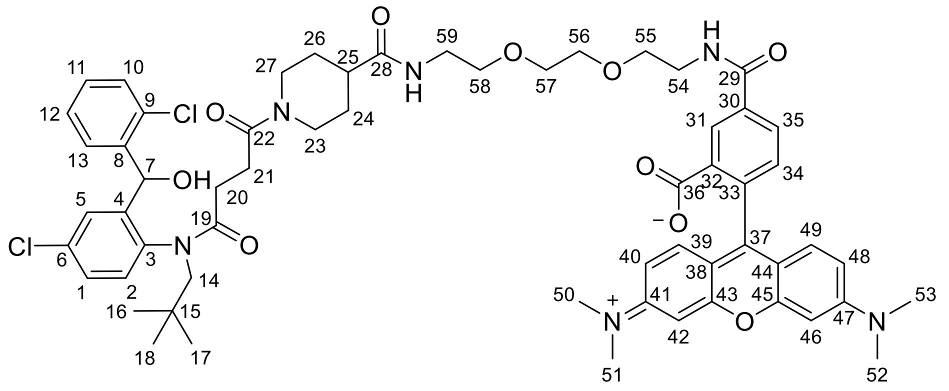

Dark violet solid (80.00 mg, 67%); **Rf** = 0.48 (DCM/MeOH 7:3); **$^1$H NMR** (800 MHz, DMSO): δ 8.89 (t, *J* = 5.6 Hz, 1H), 8.47 – 8.44 (m, 1H), 8.24 (dt, *J* = 7.9, 1.6 Hz, 1H), 7.84 (d, *J* = 16.0 Hz, 1H), 7.67 – 7.53 (m, 2H), 7.49 – 7.42 (m, 2H), 7.39 (ddt, *J* = 9.3, 4.7, 2.5 Hz, 1H), 7.35 – 7.24 (m, 2H), 7.04 (t, *J* = 3.1 Hz, 1H), 6.55 – 6.46 (m, 7H), 6.24 – 6.19 (m, 1H, **OH-8**), 6.10 (dd, *J* = 8.0, 5.0 Hz, 1H, **H7**), 5.92 – 5.88 (m, 1H, **H7**), 4.40 (dd, *J* = 13.7, 4.0 Hz, 1H), 4.28 (d, *J* = 13.0 Hz, 1H), 4.21 (d, *J* = 13.5 Hz, 1H), 3.84 (d, *J* = 13.5 Hz, 1H), 3.65 (d, *J* = 13.4 Hz, 1H), 3.60 – 3.55 (m, 5H), 3.53 (dd, *J* = 5.7, 3.4 Hz, 2H), 3.48 (q, *J* = 5.8 Hz, 2H), 3.41 (t, *J* = 6.0 Hz, 2H), 3.18 (d, *J* = 6.0 Hz, 2H), 3.04 – 3.00 (m, 1H), 2.95 (s, 12H, **H50-53**), 2.91 – 2.84 (m, 1H), 2.62 – 2.52 (m, 2H), 2.49 – 2.29 (m, 3H), 2.16 (dq, *J* = 14.7, 7.0 Hz, 1H), 2.01 (td, *J* = 17.5, 17.0, 7.2 Hz, 1H), 1.71 – 1.57 (m, 3H), 1.51 – 1.38 (m, 1H), 1.33 – 1.20 (m, 2H), 0.84 (s, 5H, **H16-18**), 0.76 (s, 4H, **H16-18**) ppm. **$^{13}$C NMR** (201 MHz, DMSO): δ 174.4, 172.4, 171.8, 170.5, 169.5, 168.8, 165.3, 155.3, 142.9, 140.1, 136.4, 134.9, 134.5, 134.2, 132.9, 132.4, 130.0, 129.9, 129.8, 129.7, 129.4, 128.9, 128.6, 128.2, 127.8, 127.3, 124.6, 123.6, 109.4, 106.0, 98.4, 85.2, 70.0, 69.5, 69.2, 67.6, 66.5, 58.6, 57.9, 55.4, 44.7, 42.0, 41.3, 38.9, 34.6, 34.2, 29.3, 29.1, 29.0 (**C-16-18**), 28.9 (**C16-18**), 28.7, 28.6, 28.4 ppm. One $CH_2$ group as well as $N(CH_3)_2$ groups are superimposing with the solvent peak. **HRMS (ESI):** m/z $[M + H]^+$ calcd for $C_{59}H_{69}Cl_2N_6O_{10}^+$ 845.4146; found 845.3151.

3-(3,4-dimethoxyphenyl)-*N*-(6-(3-((2-(2-(2-((2-(2,6-dioxopiperidin-3-yl)-1,3-dioxoisoindolin-4-yl)amino)ethoxy)ethoxy)ethyl)amino)-3-oxopropyl)benzo[d]thiazol-2-yl)propanamide **13**

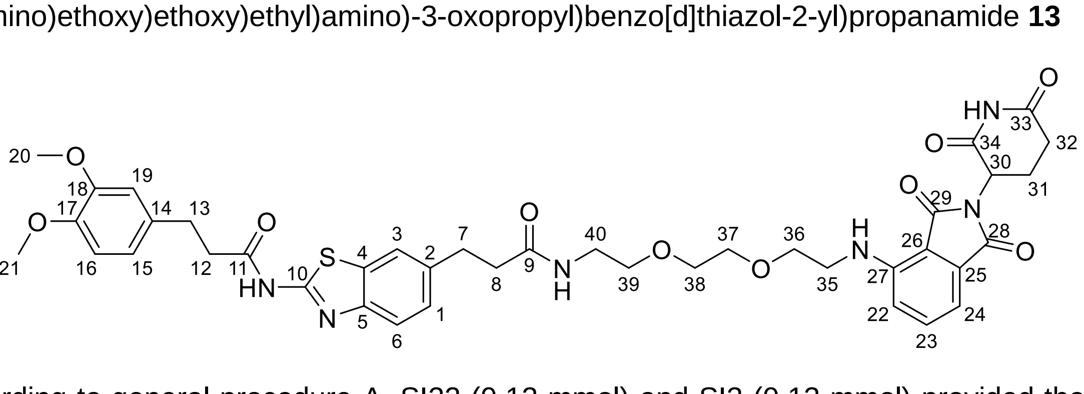

According to general procedure A, SI22 (0.12 mmol) and SI3 (0.13 mmol) provided the title compound as a yellow solid (46.00 mg, 66%). **Rf** = 0.50 (DCM/MeOH 9:1); **$^1$H NMR** (400 MHz, DMSO): δ 12.26 (s, 1H, C10-**NH**-C11), 11.09 (s, 1H, C33-**NH**-C34), 7.86 (t, *J* = 5.7 Hz, 1H,

C9-**NH**-C40), 7.75 (d, *J* = 1.7 Hz, 1H, **H3**), 7.64 – 7.52 (m, 2H, **H6, H23**), 7.24 (dd, *J* = 8.3, 1.8 Hz, 1H, **H1**), 7.12 (d, *J* = 8.6 Hz, 1H, **H22**), 7.03 (d, *J* = 7.0 Hz, 1H, **H24**), 6.88 – 6.81 (m, 2H, **H16, H19**), 6.74 (dd, *J* = 8.2, 2.0 Hz, 1H, **H15**), 6.59 (t, *J* = 5.8 Hz, 1H, C27-**NH**-C35), 5.05 (dd, *J* = 12.9, 5.3 Hz, 1H, **H30**), 3.71 (s, 3H, **H20**), 3.69 (s, 3H, **H21**), 3.60 (t, *J* = 5.4 Hz, 2H, **H36**), 3.53 (dd, *J* = 6.3, 3.7 Hz, 2H, **H38**), 3.50 – 3.41 (m, 4H, **H35, H37**), 3.36 (t, *J* = 5.8 Hz, 2H, **H39**), 3.17 (q, *J* = 5.8 Hz, 2H, **H40**), 2.88 (hept, *J* = 5.6, 4.9 Hz, 5H, **H7, H13, H32**), 2.80 – 2.72 (m, 2H, **H12**), 2.63 – 2.51 (m, 2H, **H31, H32**), 2.41 (t, *J* = 7.7 Hz, 2H, **H8**), 2.07 – 1.96 (m, 1H, **H31**) ppm. **$^{13}$C NMR** (101 MHz, DMSO): δ 172.8 (**C33**), 171.5 (**C11**), 171.3 (**C9**), 170.1 (**C34**), 168.9 (**C29**), 167.3 (**C28**), 157.2 (**C10**), 148.6 (**C17**), 147.2 (**C18**), 146.8 (**C5**), 146.4 (**C27**), 136.9 (**C2**), 136.2 (**C23**), 133.1 (**C14**), 132.1 (**C25**), 131.5 (**C4**), 126.7 (**C1**), 120.7 (**C3**), 120.1 (**C6**), 120.0 (**C15**), 117.4 (**C22**), 112.2 (**C19**), 111.9 (**C16**), 110.7 (**C24**), 109.3 (**C26**), 69.7 (**C37**), 69.6 (**C38**), 69.2 (**C39**), 68.9 (**C36**), 55.5 (**C20**), 55.4 (**C21**), 48.6 (**C30**), 41.7 (**C35**), 38.5 (**C40**), 37.2 (**C8**), 37.1 (**C12**), 31.1 (**C32**), 31.0 (**C7**), 30.0 (**C13**), 22.1 (**C31**) ppm. **HRMS (ESI):** m/z [M + H]$^+$ calcd for $C_{40}H_{45}N_6O_{10}S^+$ 801.2912; found 801.2918.

3-(3,4-dimethoxyphenyl)-*N*-(6-(3-((3-(4-(3-((2-(2,6-dioxopiperidin-3-yl)-1,3-dioxoisoindolin-4-yl)amino)propoxy)butoxy)propyl)amino)-3-oxopropyl)benzo[d]thiazol-2-yl)propanamide **14**

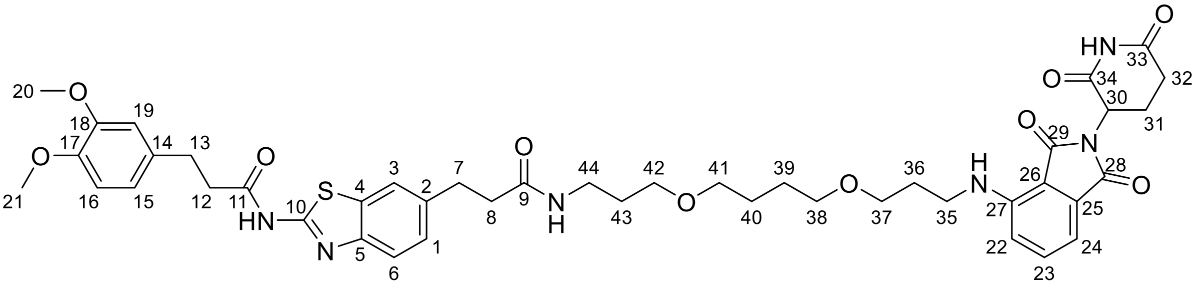

According to general procedure A, SI22 (0.12 mmol) and SI4 (0.13 mmol) provided the title compound as a yellow solid (87.00 mg, 84%). **Rf** = 0.50 (DCM/MeOH 9:1); **$^1$H NMR** (400 MHz, DMSO): δ 12.26 (s, 1H, C10-**NH**-C11), 11.09 (s, 1H, C33-**NH**-C34), 7.80 – 7.72 (m, 2H, C9-**NH**-C44, **H3**), 7.64 – 7.51 (m, 2H, **H6, H23**), 7.25 (dd, *J* = 8.3, 1.8 Hz, 1H, **H1**), 7.07 (d, *J* = 8.6 Hz, 1H, **H22**), 7.01 (d, *J* = 7.0 Hz, 1H, **H24**), 6.88 – 6.81 (m, 2H, **H16, H19**), 6.74 (dd, *J* = 8.2, 2.0 Hz, 1H, **H15**), 6.65 (t, *J* = 5.9 Hz, 1H, C27-**NH**-C35), 5.05 (dd, *J* = 12.9, 5.4 Hz, 1H, **H30**), 3.71 (s, 3H, **H20**), 3.69 (s, 3H, **H21**), 3.44 (t, *J* = 5.9 Hz, 2H, **H37**), 3.38 – 3.33 (m, 4H, **H35/H38**), 3.30 – 3.20 (m, 4H, **H41/H42**), 3.05 (q, *J* = 6.6 Hz, 2H, **H44**), 2.95 – 2.81 (m, 5H, **H7, H13, H32**), 2.80 – 2.72 (m, 2H, **H12**), 2.64 – 2.51 (m, 2H, **H31, H32**), 2.39 (t, *J* = 7.6 Hz, 1H, **H8**), 2.07 – 1.96 (m, 1H, **H31**), 1.80 (p, *J* = 6.3 Hz, 2H, **H36**), 1.60 – 1.43 (m, 6H **H39, H40, H43**) ppm. **$^{13}$C NMR** (101 MHz, DMSO): δ 172.8 (**C33**), 171.5 (**C11**), 171.1 (**C9**), 170.1 (**C43**), 168.8 (**C29**), 167.3 (**C28**), 157.1 (**C10**), 148.6 (**C17**), 147.2 (**C18**), 146.8 (**C5**), 146.4 (**C27**), 136.9 (**C2**), 136.2 (**C23**), 133.1 (**C14**), 132.2 (**C25**), 131.5 (**C4**), 126.7 (**C1**), 120.7 (**C3**), 120.1 (**C6**), 120.0 (**C15**), 117.0 (**C22**), 112.2 (**C19**), 111.9 (**C16**), 110.4 (**C24**), 109.1 (**C26**), 70.0 (**C38**), 69.8 (**C41**), 67.9 (**C42**), 67.6 (**C37**), 55.5 (**C20**), 55.3 (**C21**), 48.5 (**C30**), 37.3 (**C8**), 37.1 (**C12**), 35.7 (**C44**), 31.1 (**C32**), 31.0 (**C7**), 30.0 (**C13**), 29.4 (**C39/C40/C43**), 28.9 (**C36**), 26.0 (**C39/C40/C43**), 26.0 (**C39/C40/C43**), 22.2 (**C31**) ppm. (One CH2 group missing due to superimposition with DMSO-d6 (**C35**)).; **HRMS (ESI):** m/z [M + H]$^+$ calcd for $C_{44}H_{53}N_6O_{10}S^+$ 857.3544; found 857.3541.

3-(3,4-dimethoxyphenyl)-*N*-(6-(1-((2-(2,6-dioxopiperidin-3-yl)-1,3-dioxoisoindolin-4-yl)amino)-15-oxo-4,7,10-trioxa-14-azaheptadecan-17-yl)benzo[d]thiazol-2-yl)propanamide **15**

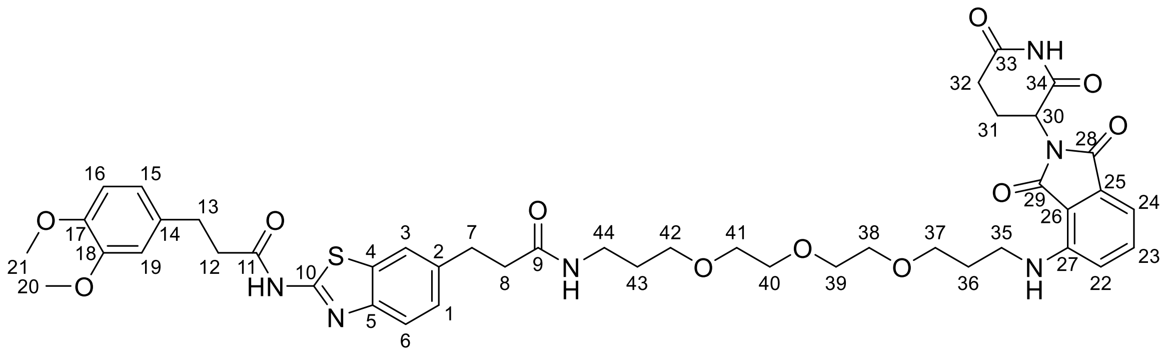

According to general procedure A, SI22 (0.12 mmol) and SI5 (0.13 mmol) provided the title compound as a yellow solid (76.00 mg, 72%). **Rf** = 0.56 (DCM/MeOH 9:1); **$^1$H NMR** (400 MHz, DMSO): δ 12.26 (s, 1H, C10-**NH**-C11), 11.09 (s, 1H, C33-**NH**-C34), 7.80 – 7.72 (m, 2H, C9-**NH**-C44, **H3**), 7.64 – 7.52 (m, 2H, **H6, H23**), 7.25 (dd, *J* = 8.3, 1.8 Hz, 1H, **H1**), 7.09 (d, *J* = 8.6 Hz, 1H, **H22**), 7.01 (d, *J* = 7.0 Hz, 1H, **H24**), 6.88 – 6.81 (m, 2H, **H16, H19**), 6.74 (dd, *J* = 8.2, 2.0 Hz, 1H, **H15**), 6.66 (t, *J* = 5.9 Hz, 1H, C27-**NH**-C35), 5.04 (dd, *J* = 12.9, 5.4 Hz, 1H, **H30**), 3.71 (s, 3H, **H20**), 3.69 (s, 3H, **H21**), 3.53 (dd, *J* = 6.3, 3.8 Hz, 2H, **H40**), 3.52 – 3.44 (m, 6H **H38, H39, H42**), 3.42 – 3.33 (m, 4H, **H35, H41**), 3.28 (t, *J* = 6.4 Hz, 2H, **H37**), 3.05 (q, *J* = 6.6 Hz, 2H, **H44**), 2.95 – 2.81 (m, 5H, **H7, H13, H32**), 2.81 – 2.72 (m, 2H, **H12**), 2.63 – 2.51 (m, 2H, **H31, H32**), 2.39 (t, *J* = 7.6 Hz, 1H, **H8**), 2.07 – 1.96 (m, 1H, **H31**), 1.80 (p, *J* = 6.4 Hz, 2H, **H36**), 1.55 (p, *J* = 6.7 Hz, 2H, **H43**) ppm. **$^{13}$C NMR** (101 MHz, DMSO): δ 172.8 (**C33**), 171.5 (**C11**), 171.1 (**C9**), 170.1 (**C34**), 168.8 (**C29**), 167.3 (**C28**), 157.2 (**C10**), 148.6 (**C17**), 147.2 (**C18**), 146.8 (**C5**), 146.4 (**C27**), 136.9 (**C2**), 136.3 (**C23**), 133.1 (**C14**), 132.2 (**C25**), 131.5 (**C4**), 126.7 (**C1**), 120.7 (**C3**), 120.1 (**C6**), 120.0 (**C15**), 117.1 (**C22**), 112.2 (**C19**), 111.9 (**C16**), 110.4 (**C24**), 109.0 (**C26**), 69.8 (**C38/C39/C40**), 69.7 (**C38/C39/C40**), 69.7 (**C38/C39/C40**), 69.5 (**C41**), 68.2 (**C42**), 68.0 (**C37**), 55.5 (**C20**), 55.4 (**C21**), 48.5 (**C30**), 37.3 (**C8**), 37.1 (**C12**), 35.7 (**C44**), 31.1 (**C32**), 31.0 (**C7**), 30.0 (**C13**), 29.3 (**C43**), 28.9 (**H36**), 22.2 (**C31**) ppm. (one CH2 group missing due to superimposition with DMSO-d6 (**C35**)). **HRMS (ESI):** m/z [M + H]$^+$ calcd for $C_{44}H_{53}N_6O_{11}S^+$ 873.3493; found 873.3491.

3-(3,4-dimethoxyphenyl)-*N*-(6-(3-((6-((2-(2,6-dioxopiperidin-3-yl)-1,3-dioxoisoindolin-4-yl)amino)hexyl)amino)-3-oxopropyl)benzo[d]thiazol-2-yl)propanamide **16**

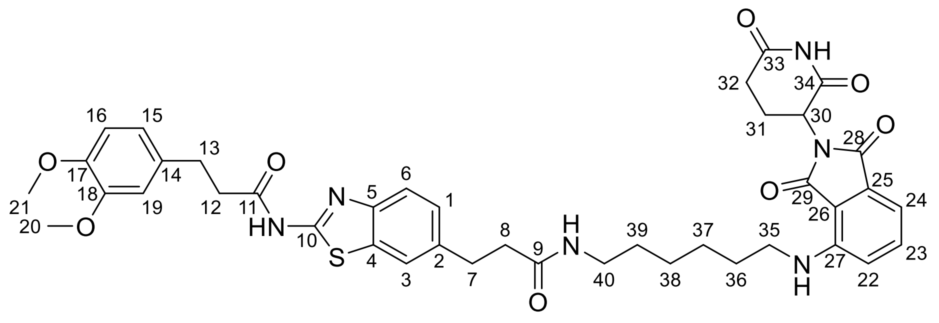

According to general procedure A, SI22 (0.07 mmol) and SI1 (0.08 mmol) provided the title compound as a yellow solid (25.00 mg, 45%). **Rf** = 0.55 (DCM/MeOH 9:1); **$^1$H NMR** (400 MHz, DMSO): δ 12.26 (s, 1H, C10-**NH**-C11), 11.09 (s, 1H, C33-**NH**-C34), 7.79 – 7.72 (m, 2H, C9-**NH**-C44, **H3**), 7.64 – 7.54 (m, 2H, **H6, H23**), 7.25 (dt, *J* = 8.3, 2.2 Hz, 1H, **H1**), 7.06 (d, *J* = 8.6

Hz, 1H, **H22**), 7.01 (d, *J* = 6.9 Hz, 1H , **H24**), 6.88 – 6.80 (m, 2H, **H16, H19**), 6.73 (dd, *J* = 8.2, 2.0 Hz, 1H, **H15**), 6.50 (t, *J* = 5.9 Hz, 1H, C27-**NH**-C35), 5.05 (dd, *J* = 12.9, 5.4 Hz, 1H, **H30**), 3.71 (s, 3H, **H20**), 3.69 (d, *J* = 1.0 Hz, 3H, **H21**), 3.31 – 3.16 (m, 2H, **H35**), 3.00 (q, *J* = 6.5 Hz, 2H, **H40**), 2.94 – 2.81 (m, 5H, **H7, H13, H32**), 2.80 – 2.71 (m, 2H, **H12**), 2.63 – 2.51 (m, 2H, **H31, H32**), 2.39 (t, *J* = 7.6 Hz, 2H, **H8**), 2.07 – 1.96 (m, 1H, **H31**), 1.50 (p, *J* = 7.2 Hz, 2H, **H36**), 1.38 – 1.14 (m, 6H, **H37, H38, H39**) ppm. $^{13}$**C NMR** (101 MHz, DMSO): δ 172.8 (**C33**), 171.5 (**C11**), 171.0 (**C9**), 170.1 (**C34**), 168.9 (**C29**), 167.3 (**C28**), 157.1 (**C10**), 148.6 (**C17**), 147.2 (**C18**), 146.8 (**C5**), 146.4 (**C27**), 136.9 (**C2**), 136.3 (**C23**), 133.1 (**C14**), 132.2 (**C25**), 131.5 (**C4**), 126.7 (**C1**), 120.7 (**C3**), 120.1 (**C6**), 120.0 (**C15**), 117.1 (**C22**), 112.2 (**C19**), 111.9 (**C16**), 110.4 (**C24**), 109.0 (**C26**), 55.5 (**C20**), 55.3 (**C21**), 48.5 (**C30**), 41.8 (**C35**), 38.3 (**C40**), 37.3 (**C8**), 37.1 (**C12**), 31.1 (**C32**), 31.0 (**C7**), 30.0 (**C13**), 29.1 (**C39**), 28.6 (**C36**), 26.1 (**C37/C38**), 26.1 (**C37/C38**), 22.1 (**C31**) ppm. **HRMS (ESI):** m/z $[M + H]^+$ calcd for $C_{40}H_{45}N_6O_8S^+$ 769.3019; found 769.3020.

3-(3,4-dimethoxyphenyl)-*N*-(6-(3-((10-((2-(2,6-dioxopiperidin-3-yl)-1,3-dioxoisoindolin-4-yl)amino)decyl)amino)-3-oxopropyl)benzo[d]thiazol-2-yl)propanamide **17**

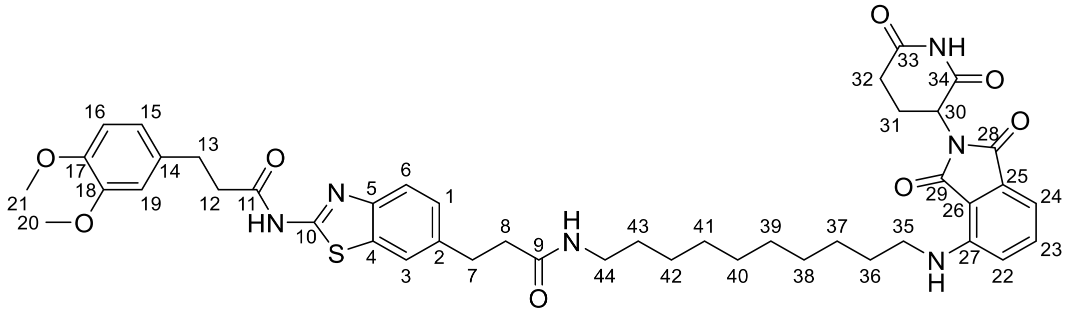

According to general procedure A, SI22 (0.07 mmol) and SI6 (0.08 mmol) provided the title compound as a yellow solid (45.00 mg, 74%). **Rf** = 0.63 (DCM/MeOH 9:1); $^{1}$**H NMR** (400 MHz, DMSO): δ 12.26 (s, 1H, C10-**NH**-C11), 11.09 (s, 1H, C33-**NH**-C34), 7.76 – 7.71 (m, 2H, C9-**NH**-C44, **H3**), 7.62 – 7.53 (m, 2H, **H6, H23**), 7.25 (dd, *J* = 8.3, 1.7 Hz, 1H, **H1**), 7.07 (d, *J* = 8.6 Hz, 1H, **H22**), 7.01 (dd, *J* = 6.9, 2.0 Hz, 1H, **H24**), 6.88 – 6.78 (m, 2H, **H16, H19**), 6.73 (dd, *J* = 8.2, 2.0 Hz, 1H, **H15**), 6.51 (t, *J* = 5.8 Hz, 1H, C27-**NH**-C35), 5.05 (dd, *J* = 12.9, 5.3 Hz, 1H, **H30**), 3.71 (s, 3H, **H20**), 3.69 (s, 3H, **H21**), 3.27 (q, *J* = 6.6 Hz, 2H, **H35**), 2.98 (q, *J* = 6.6 Hz, 2H, **H44**), 2.93 – 2.81 (m, 5H, **H7, H13, H32**), 2.80 – 2.72 (m, 2H, **H12**), 2.63 – 2.51 (m, 2H, **H31, H32**), 2.39 (t, *J* = 7.5 Hz, 2H, **H8**), 2.06 – 1.97 (m, 1H, **H31**), 1.60 – 1.50 (m, 2H, **H36**), 1.35 – 1.05 (m, 14H, **H37-H43**) ppm. $^{13}$**C NMR** (101 MHz, DMSO): δ 172.8 (**C33**), 171.4 (**C11**), 170.9 (**C9**), 170.1 (**C34**), 169.0 (**C29**), 167.3 (**C28**), 157.1 (**C10**), 148.6 (**C17**), 147.2 (**C18**), 146.8 (**C5**), 146.4 (**C27**), 136.9 (**C2**), 136.3 (**C23**), 133.1 (**C14**), 132.2 (**C25**), 131.5 (**C4**), 126.7 (**C1**), 120.7 (**C3**), 120.1 (**C6**), 120.0 (**C15**), 117.2 (**C22**), 112.2 (**C19**), 111.9 (**C16**), 110.4 (**C24**), 109.0 (**C26**), 55.5 (**C20**), 55.3 (**C21**), 48.5 (**C30**), 41.8 (**C35**), 38.4 (**C44**), 37.3 (**C8**), 37.1 (**C12**), 31.1 (**C32**), 31.0 (**C7**), 30.0 (**C13**), 29.1 (**C37-C43**), 29.0 (**C37-C43**), 28.9 (**C37-C43**), 28.8 (**C37-C43**), 28.7 (**C37-C43**), 26.3 (**C37-C43**), 22.2 (**C31**) ppm. LCMS: **HRMS (ESI):** m/z $[M + H]^+$ calcd for $C_{44}H_{53}N_6O_8S^+$ 825.3645; found 825.3646.

(2*S*,4*R*)-1-((*S*)-2-(*tert*-butyl)-15-(2-(3-(3,4-dimethoxyphenyl)propanamido)benzo[*d*]thiazol-6-yl)-4,13-dioxo-6,9-dioxa-3,12-diazapentadecanoyl)-4-hydroxy-*N*-(4-(4-methylthiazol-5-yl)benzyl)pyrrolidine-2-carboxamide **18**

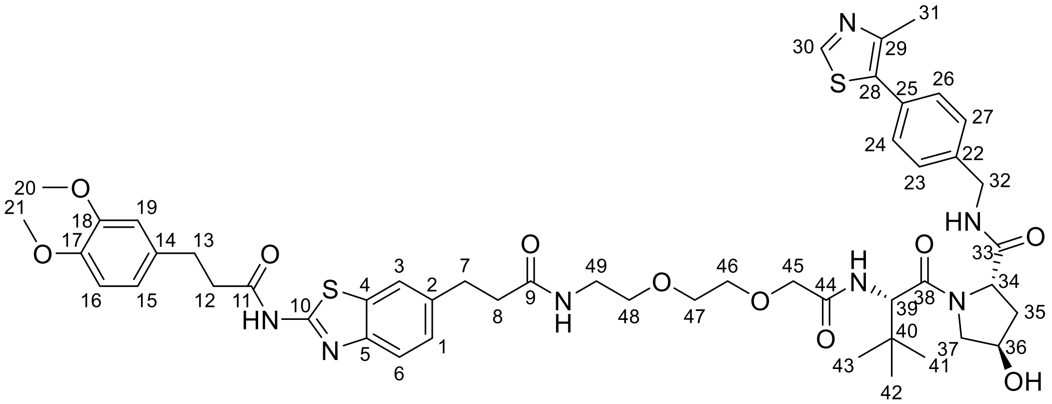

According to general procedure B, SI22 (0.07 mmol) and SI13 (0.08 mmol) provided the title compound as a white solid (59.00 mg, 84%). **Rf** = 0.47 (DCM/MeOH 9:1); **$^1$H NMR** (400 MHz, DMSO): δ 12.25 (s, 1H, C10-**NH**-C11), 8.97 (s, 1H, **H30**), 8.58 (t, *J* = 6.0 Hz, 1H, C32-**NH**-C33), 7.89 (t, *J* = 5.7 Hz, 1H, C9-**NH**-C49), 7.75 (d, *J* = 1.7 Hz, 1H, **H3**), 7.60 (d, *J* = 8.3 Hz, 1H, **H6**), 7.44 (d, *J* = 6.4 Hz, 1H, C39-**NH**-C44), 7.38 (s, 4H, **H23, H24, H26, H27**), 7.28 – 7.20 (m, 1H, **H1**), 6.88 – 6.81 (m, 2H, **H16, H19**), 6.74 (dd, *J* = 8.2, 2.0 Hz, 1H, **H15**), 5.16 (d, *J* = 3.5 Hz, 1H, C36-**OH**), 4.58 (d, *J* = 9.6 Hz, 1H, **H39**), 4.45 (t, *J* = 8.2 Hz, 1H, **H34**), 4.41 – 4.31 (m, 2H, **H32, H36**), 4.26 (dd, *J* = 15.7, 5.8 Hz, 1H, **H32**), 3.96 (s, 2H, **H45**), 3.71 (s, 3H **H20**), 3.69 (s, 3H, **H21**), 3.68 – 3.54 (m, 4H, **H37, H46**), 3.50 (q, *J* = 4.8, 4.2 Hz, 2H, **H47**), 3.39 (t, *J* = 5.8 Hz, 2H, **H48**), 3.25 – 3.15 (m, 2H, **H49**), 2.95 – 2.81 (m, 4H, **H7, H13**), 2.80 – 2.72 (m, 2H, **H12**), 2.47 – 2.37 (m, 5H, **H8, H31**), 2.06 (dd, *J* = 13.0, 7.7 Hz, 1H, **H35**), 1.90 (ddd, *J* = 13.0, 8.8, 4.5 Hz, 1H, **H35**), 0.94 (s, 9H, **H41-H43**) ppm. **$^{13}$C NMR** (101 MHz, DMSO): δ 171.7 (**C33**), 171.5 (**C11**), 171.4 (**C9**), 169.2 (**C38**), 168.6 (**C44**), 157.2 (**C10**), 151.4 (**C30**), 148.6 (**C17**), 147.8 (**C29**), 147.2 (**C18**), 146.8 (**C5**), 139.4 (**C22**), 136.9 (**C2**), 133.1 (**C14**), 131.5 (**C4**), 131.1 (**C28**), 129.7 (**C25**), 128.7 (**C23/24/26/27**), 127.4 (**C23/24/26/27**), 126.7 (**C1**), 120.7 (**C3**), 120.1 (**C6**), 120.0 (**C15**), 112.2 (**C19**), 111.9 (**C16**), 70.4 (**C46**), 69.5 (**C45**), 69.4 (**C47**), 69.3 (**C48**), 68.9 (**C36**), 58.7 (**C34**), 56.6 (**C37**), 55.7 (**C39**), 55.5 (**C20**), 55.4 (**C21**), 41.7 (**C32**), 38.5 (**C49**), 38.0 (**C35**), 37.2 (**C8**), 37.2 (**C12**), 35.7 (**C40**), 31.1 (**C7**), 30.0 (**C13**), 26.2 (**C41-C43**), 15.9 (**C31**) ppm. **HRMS (ESI):** m/z $[M + H]^+$ calcd for $C_{49}H_{62}N_7O_{10}S_2{}^+$ 972.3999; found 972.3996. **Optical Rotation:** $[\alpha]_D^{20}$ = -26.61 (c = 0.68, EtOH).

(2*S*,4*R*)-1-((*S*)-2-(*tert*-butyl)-22-(2-(3-(3,4-dimethoxyphenyl)propanamido)benzo[*d*]thiazol-6-yl)-4,20-dioxo-7,10,13,16-tetraoxa-3,19-diazadocosanoyl)-4-hydroxy-*N*-(4-(4-methylthiazol-5-yl)benzyl)pyrrolidine-2-carboxamide **19**

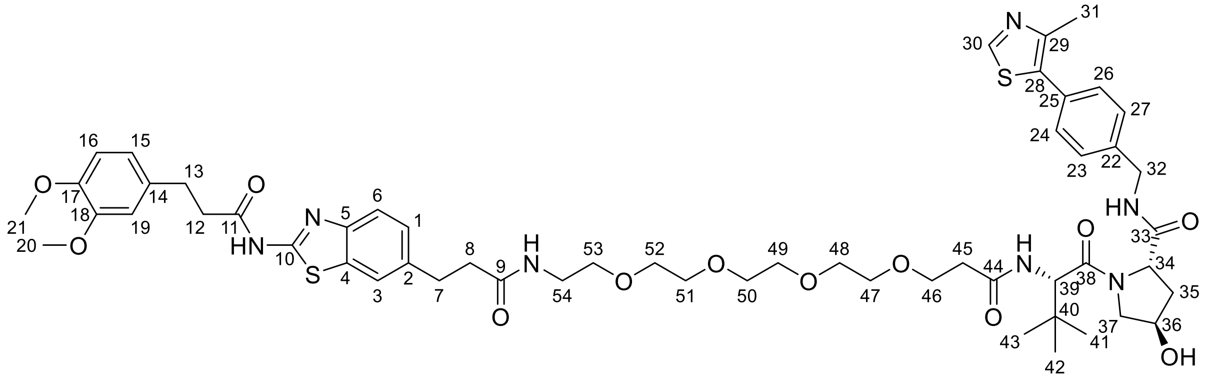

According to general procedure B, SI22 (0.07 mmol) and SI14 (0.08 mmol) provided the title compound as a white solid (71.50 mg, 92%). **Rf** = 0.41 (DCM/MeOH 9:1); **$^1$H NMR** (400 MHz, DMSO): δ 12.26 (s, 1H, C10-**NH**-C11), 8.98 (s, 1H, **H30**), 8.57 (t, *J* = 6.1 Hz, 1H, C32-**NH**-C33), 7.97 – 7.85 (m, 2H, C9-**NH**-C54, C39-**NH**-C44), 7.75 (d, *J* = 1.7 Hz, 1H, **H3**), 7.61 (d, *J* = 8.3 Hz, 1H, **H6**), 7.40 (q, *J* = 8.4 Hz, 4H, **H23, H24, H26, H27**), 7.25 (dd, *J* = 8.3, 1.8 Hz, 1H, **H1**), 6.88 – 6.81 (m, 2H, **H16, H19**), 6.74 (dd, *J* = 8.2, 2.0 Hz, 1H, **H15**), 5.13 (d, *J* = 3.5 Hz, 1H, C36-**OH**), 4.55 (d, *J* = 9.4 Hz, 1H, **H39**), 4.50 – 4.38 (m, 2H, **H32, H34**), 4.35 (s, 1H, **H36**), 4.21 (dd, *J* = 15.9, 5.4 Hz, 1H, **H32**), 3.71 (s, 3H, **H20**), 3.69 (s, 3H, **H21**), 3.68 – 3.53 (m, 4H, **H37, H46**), 3.50 – 3.42 (m, 12H, **H47-H52**), 3.35 – 3.31 (m, 2H **H53**), 3.17 (q, *J* = 5.8 Hz, 2H, **H54**), 2.94 – 2.83 (m, 4H, **H7, H13**), 2.80 – 2.72 (m, 2H, **H12**), 2.59 – 2.52 (m, 1H, **H45**), 2.47 – 2.29 (m, 6H, **H8, H31, H45**), 2.03 (ddd, *J* = 11.0, 7.6, 2.5 Hz, 1H, **H35**), 1.90 (ddd, *J* = 12.9, 8.6, 4.6 Hz, 1H, **H35**), 0.93 (s, 9H, **H41-H43**) ppm. **$^{13}$C NMR** (101 MHz, DMSO): δ 171.9 (**C33**), 171.5 (**C11**), 171.3 (**C44**), 169.9 (**C38**), 169.5 (**C9**), 157.2 (**C10**), 151.5 (**C30**), 148.6 (**C17**), 147.7 (**C29**), 147.2 (**C18**), 146.9 (**C5**), 139.5 (**C22**), 136.9 (**C2**), 133.1 (**C14**), 131.5 (**C4**), 131.2 (**C28**), 129.6 (**C25**), 128.6 (**C23/24/26/27**), 127.4 (**C23/24/26/27**), 126.7 (**C1**), 120.7 (**C3**), 120.1 (**C6**), 120.0 (**C15**), 112.2 (**C19**), 111.9 (**C16**), 69.8 (**C47-C52**), 69.7 (**C47-C52**), 69.6 (**C47-C52**), 69.5 (**C47-C52**), 69.2 (**C53**), 68.9 (**C36**), 66.9 (**C46**), 58.7 (**C34**), 56.4 (**C37**), 56.3 (**C39**), 55.5 (**C20**), 55.4 (**C21**), 41.6 (**C32**), 38.5 (**C54**), 37.9 (**C35**), 37.3 (**C8**), 37.2 (**C12**), 35.6 (**C45**), 35.4 (**C40**), 31.1 (**C7**), 30.0 (**C13**), 26.3 (**C41-C43**), 15.9 (**C31**) ppm. LCMS: **HRMS (ESI):** m/z $[M + Na]^+$ calcd for $C_{54}H_{72}N_7O_{12}S_2Na^+$ 1096.4494; found 1096.4501. **Optical Rotation:** $[\alpha]_D^{20}$ = -11.04 (c = 1.03, EtOH).

(2*S*,4*R*)-1-((*S*)-2-(6-(3-(2-(3-(3,4-dimethoxyphenyl)propanamido)benzo[*d*]thiazol-6-yl)propanamido)hexanamido)-3,3-dimethylbutanoyl)-4-hydroxy-*N*-(4-(4-methylthiazol-5-yl)benzyl)pyrrolidine-2-carboxamide **20**

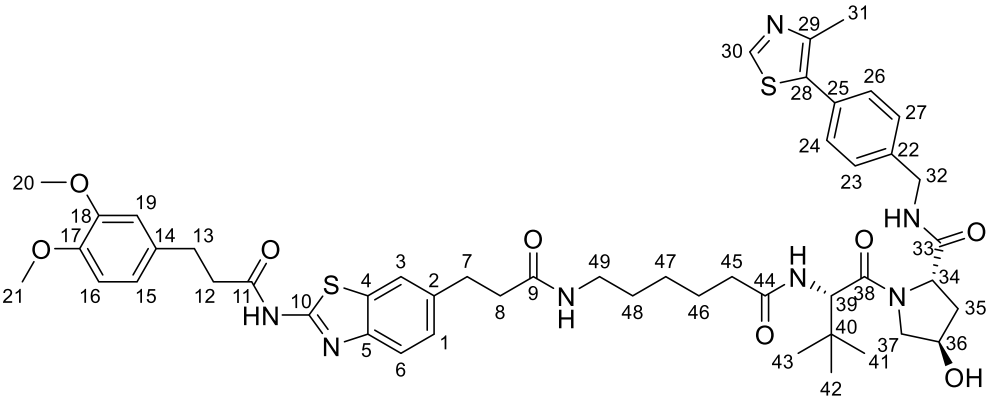

According to general procedure B, SI22 (0.06 mmol) and SI14 (0.07 mmol) provided the title compound as a white solid (49.00 mg, 72%). **Rf** = 0.40 (DCM/MeOH 9:1); **$^{1}$H NMR** (400 MHz, DMSO): δ 12.25 (s, 1H, C10-**NH**-C11), 8.98 (s, 1H, **H30**), 8.56 (t, *J* = 6.1 Hz, 1H C32-**NH**-C33), 7.85 (d, *J* = 9.4 Hz, 1H, C9-**NH**-C49), 7.80 – 7.73 (m, 2H, **H3**, C39-**NH**-C44), 7.61 (d, *J* = 8.3 Hz, 1H, **H6**), 7.40 (q, *J* = 8.5 Hz, 4H, **H23, H24, H26, H27**), 7.25 (dd, *J* = 8.3, 1.7 Hz, 1H, **H1**), 6.88 – 6.79 (m, 2H, **H16, H19**), 6.74 (dd, *J* = 8.2, 2.0 Hz, 1H, **H15**), 5.13 (d, *J* = 3.5 Hz, 1H, C36-**OH**), 4.54 (d, *J* = 9.4 Hz, 1H, **H39**), 4.48 – 4.38 (m, 2H, **H32, H34**), 4.35 (br s, 1H, **H36**), 4.21 (dd, *J* = 15.9, 5.5 Hz, 1H, **H32**), 3.71 (s, 3H, **H20**), 3.69 (s, 3H, **H21**), 3.70 – 3.60 (m, 2H, **H37**), 2.99 (q, *J* = 6.6 Hz, 2H, **H49**), 2.94 – 2.83 (m, 4H, **H7, H13**), 2.81 – 2.72 (m, 2H, **H12**), 2.44 (s, 3H, **H31**), 2.39 (t, *J* = 7.8 Hz, 2H, **H8**), 2.25 – 2.16 (m, **H45**), 2.14 – 1.98 (m, 2H, **H35, H45**), 1.90 (ddd, *J* = 12.9, 8.7, 4.6 Hz, 1H, **H35**), 1.47 (dq, *J* = 14.6, 7.0 Hz, 2H, **H46**), 1.40 – 1.28 (m, 2H, **H48**), 1.25 – 1.12 (m, 2H, **H47**), 0.93 (s, 9H, **H41-43**) ppm. **$^{13}$C NMR** (101 MHz, DMSO): δ 172.0 (**C33**), 171.9 (**C44**), 171.5 (**C11**), 171.0 (**C9**), 169.8 (**C38**), 157.2 (**C10**), 151.5 (**C30**), 148.6 (**C17**), 147.7 (**C29**), 147.2 (**C18**), 146.8 (**C5**), 139.5 (**C22**), 137.0 (**C2**), 133.1 (**C14**), 131.5 (**C4**), 131.2 (**C28**), 129.6 (**C25**), 128.6 (**C23/24/26/27**), 127.4 (**C23/24/26/27**), 126.7 (**C1**), 120.7 (**C3**), 120.1 (**C6**), 120.0 (**C15**), 112.2 (**C19**), 111.9 (**C16**), 68.9 (**C36**), 58.7 (**C34**), 56.4 (**C37**), 56.3 (**C39**), 55.5 (**C20**), 55.4 (**C21**), 41.6 (**C32**), 38.4 (**C49**), 38.0 (**C35**), 37.4 (**C8**) , 37.1 (**C12**), 35.2 (**C40**), 34.8 (**C45**), 31.2 (**C7**), 30.0 (**C13**), 28.9 (**C48**), 26.4 (**C41-43**), 26.1 (**C47**), 25.2 (**C46**), 15.9 (**C31**) ppm. **HRMS (ESI):** m/z $[M + H]^+$ calcd for $C_{49}H_{62}N_7O_8S_2^+$ 940.4101; found 940.4096. **Optical Rotation:** $[\alpha]_D^{20}$ = -91.90 (c = 0.65, EtOH).

(2*S*,4*R*)-1-((*S*)-2-(10-(3-(2-(3-(3,4-dimethoxyphenyl)propanamido)benzo[*d*]thiazol-6-yl)propanamido)decanamido)-3,3-dimethylbutanoyl)-4-hydroxy-*N*-(4-(4-methylthiazol-5-yl)benzyl)pyrrolidine-2-carboxamide **21**

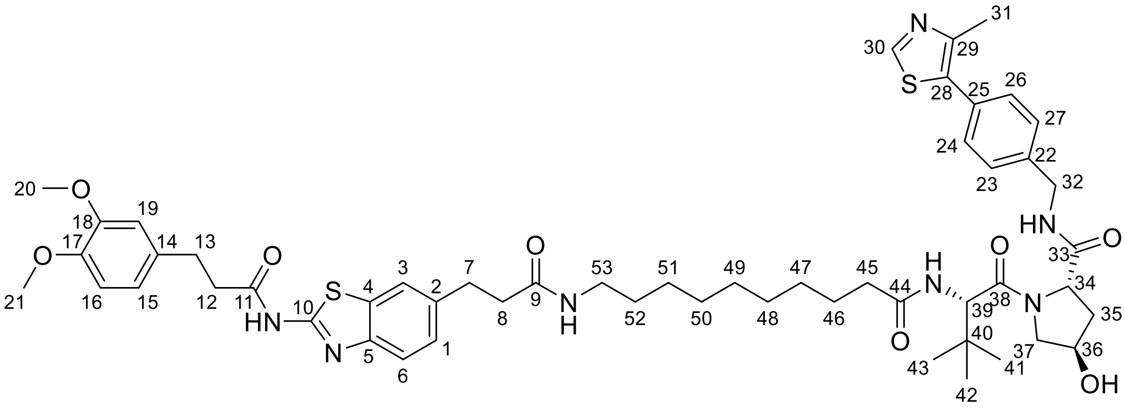

According to general procedure B, SI22 (0.07 mmol) and SI12 (0.08 mmol) provided the title compound as a white solid (70.00 mg, 97%). **Rf** = 0.48 (DCM/MeOH 9:1); **$^1$H NMR** (400 MHz, DMSO): δ 12.26 (s, 1H, C10-**NH**-C11), 8.98 (s, 1H, **H30**), 8.56 (t, *J* = 6.0 Hz, 1H, C32-**NH**-C33), 7.83 (d, *J* = 9.3 Hz, 1H, C9-**NH**-C53), 7.78 – 7.71 (m, 2H, **H3**, C39-**NH**-C44), 7.60 (d, *J* = 8.3 Hz, 1H, **H6**), 7.40 (q, *J* = 8.4 Hz, 4H, **H23, H24, H26, H27**), 7.25 (dd, *J* = 8.4, 1.8 Hz, 1H, **H1**), 6.88 – 6.81 (m, 2H, **H16, H19**), 6.74 (dd, *J* = 8.2, 2.0 Hz, 1H, **H15**), 5.12 (d, *J* = 3.5 Hz, 1H, C36-**OH**), 4.54 (d, *J* = 9.4 Hz, 1H, **H39**), 4.48 – 4.39 (m, 2H, **H32, H34**), 4.39 – 4.31 (m, 1H, **H36**), 4.21 (dd, *J* = 15.9, 5.4 Hz, 1H, **H32**), 3.71 (s, 3H, **H20**), 3.69 (s, 3H, **H21**), 3.70 – 3.60 (m, 2H, **H37**), 2.99 (q, *J* = 6.6 Hz, 2H, **H53**), 2.89 (q, *J* = 8.1 Hz, 4H, **H7, H13**), 2.81 – 2.72 (m, 2H, **H12**), 2.44 (s, 3H, **H31**), 2.39 (t, *J* = 7.6 Hz, 2H, **H8**), 2.31 – 2.19 (m, 1H, **H45**), 2.16 – 1.98 (m, 2H, **H35, H45**), 1.90 (ddd, *J* = 12.9, 8.6, 4.6 Hz, 1H, **H35**), 1.45 – 1.37 (m, 2H, **H46**), 1.33 – 1.07 (m, 12H, **H47-H52**), 0.93 (s, 9H, **H41-H43**) ppm. **$^{13}$C NMR** (101 MHz, DMSO): δ 172.1 (**C33**), 171.9 (**C44**), 171.5 (**C11**), 171.0 (**C9**), 169.7 (**C38**), 157.2 (**C10**), 151.5 (**C30**), 148.6 (**C17**), 147.7 (**C29**), 147.2 (**C18**), 146.8 (**C5**), 139.5 (**C22**), 136.9 (**C2**), 133.1 (**C14**), 131.5 (**C4**), 131.2 (**C28**), 129.6 (**C25**), 128.6 (**C23/24/26/27**), 127.4 (**C23/24/26/27**), 126.7 (**C1**), 120.7 (**C3**), 120.1 (**C6**), 120.0 (**C15**), 112.2 (**C19**), 111.9 (**C16**), 68.9 (**C36**), 58.7 (**C34**), 56.3 (**C37**), 56.3 (**C39**), 55.5 (**C20**), 55.3 (**C21**), 41.6 (**C32**), 38.4 (**C53**), 38.0 (**C35**), 37.3 (**C8**), 37.2 (**C12**), 35.2 (**C40**), 34.9 (**C45**), 31.1 (**C7**), 30.0 (**C13**), 29.1 (**C52**), 28.9 (**C47-C52**), 28.7 (**C47-C52**), 28.7 (**C47-C52**), 26.4 (**C41-C43**), 25.5 (**C46**), 15.9 (**C31**) ppm. **HRMS (ESI):** m/z $[M + H]^+$ calcd for $C_{53}H_{70}N_7O_8S_2{}^+$ 996.4727; found 996.4720. **Optical Rotation:** $[\alpha]_D^{20}$ = -18.41 (c = 1.03, EtOH).

*N*-(6-(1-((3*r*,5*r*,7*r*)-adamantan-1-yl)-2,13-dioxo-6,9-dioxa-3,12-diazapentadecan-15-yl)benzo[*d*]thiazol-2-yl)-3-(3,4-dimethoxyphenyl)propanamide **22**

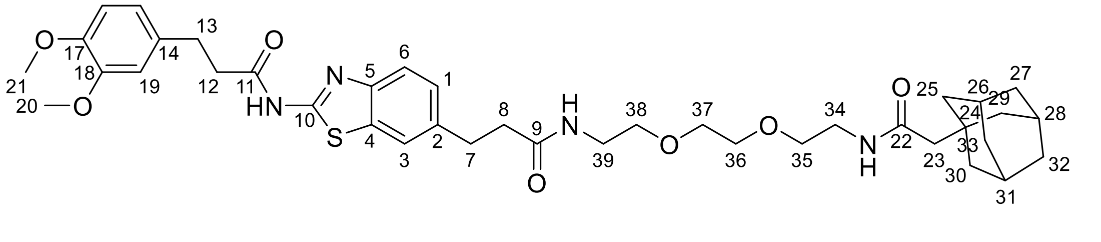

According to general procedure B, SI22 (0.07 mmol) and SI7 (0.08 mmol) provided the title compound as a white solid (41.00 mg, 79%). **Rf** = 0.48 (DCM/MeOH 9:1); **$^1$H NMR** (400 MHz,

DMSO): δ 12.27 (s, 1H, C10-**NH**-C11), 7.89 (t, *J* = 5.7 Hz, 1H C9-**NH**-C39), 7.76 (d, *J* = 1.7 Hz, 1H, **H3**), 7.69 (t, *J* = 5.7 Hz, 1H, C22-**NH**-C34), 7.61 (d, *J* = 8.3 Hz, 1H, **H6**), 7.26 (dd, *J* = 8.3, 1.8 Hz, 1H, **H1**), 6.88 – 6.81 (m, 2H **H16, H19**), 6.74 (dd, *J* = 8.2, 2.0 Hz, 1H, **H15**), 3.72 (s, 3H, **H20**), 3.70 (s, 3H, **H21**), 3.45 (s, 4H, **H35, H38**), 3.40 – 3.34 (m, 4H, **H36, H37**), 3.17 (p, *J* = 5.7 Hz, 4H, **H34, H39**), 2.94 – 2.84 (m, 4H, **H7, H13**), 2.81 – 2.72 (m, 2H, **H12**), 2.42 (dd, *J* = 8.5, 6.9 Hz, 2H, **H8**), 1.91 – 1.84 (m, 3H, **H26, H28, H31**), 1.80 (s, 2H, **H23**), 1.67 – 1.50 (m, 12H, **H25, H27, H29, H30, H32, H33**) ppm. $^{13}$**C NMR** (101 MHz, DMSO): δ 171.5 (**C11**), 171.3 (**C9**) , 170.0 (**C22**), 157.2 (**C10**), 148.6 (**C17**), 147.2 (**C18**), 146.9 (**C5**), 136.9 (**C2**), 133.1 (**C14**), 131.5 (**C4**), 126.7 (**C1**), 120.7 (**C3**), 120.2 (**C6**), 120.0 (**C15**), 112.2 (**C19**), 111.9 (**C16**), 69.5, 69.5, 69.2, 69.2, 55.5 (**C20**), 55.4 (**C21**), 50.0 (**C23**), 42.1 (**C25, C27, C29, C30, C32, C33**), 38.5 (**C39**), 38.3 (**C34**), 37.3 (**C8**), 37.2 (**C12**), 36.5 (**C25, C27, C29, C30, C32, C33**), 32.1 (**C24**), 31.1 (**C7**), 30.0 (**C13**), 28.0 (**C26, C28, C31**) ppm. **HRMS (ESI):** m/z $[M + H]^+$ calcd for $C_{39}H_{53}N_4O_7S^+$ 721.3635; found 721.3638.

*N*-(6-(1-((3*r*,5*r*,7*r*)-adamantan-1-yl)-2,17-dioxo-7,12-dioxa-3,16-diazanonadecan-19-yl)benzo[*d*]thiazol-2-yl)-3-(3,4-dimethoxyphenyl)propanamide **23**

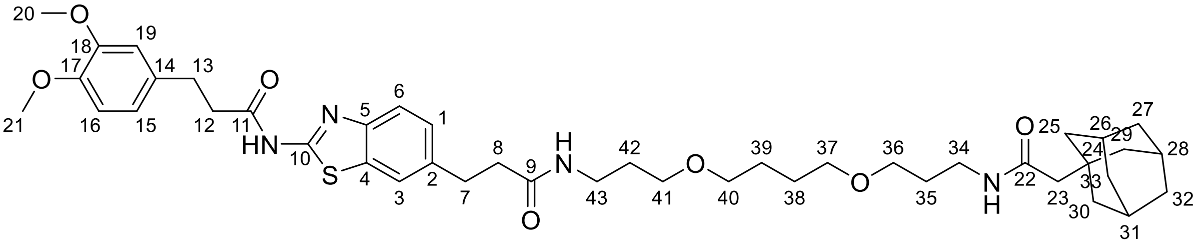

According to general procedure B, SI22 (0.07 mmol) and SI8 (0.08 mmol) provided the title compound as a white solid (40.00 mg, 71%). **Rf** = 0.41 (DCM/MeOH 9:1); $^{1}$**H NMR** (400 MHz, DMSO): δ 12.27 (s, 1H, C10-**NH**-C11), 7.81 – 7.73 (m, 2H, C9-**NH**-C43, **H3**), 7.72 – 7.56 (m, 2H, C22-**NH**-C34, **H6**), 7.25 (dd, *J* = 8.3, 1.7 Hz, 1H, **H1**), 6.88 – 6.81 (m, 2H, **H16, H19**), 6.74 (dd, *J* = 8.2, 2.0 Hz, 1H, **H15**), 3.71 (s, 3H, **H20**), 3.70 (s, 3H, **H21**), 3.38 – 3.28 (m, 4H, **H40, H41**), 3.29 – 3.19 (m, 4H, **H36, H37**), 3.05 (q, *J* = 6.5 Hz, 4H, **H34, H43**), 2.89 (dt, *J* = 10.6, 7.5 Hz, 4H, **H7, H13**), 2.81 – 2.72 (m, 2H, **H12**), 2.40 (dd, *J* = 8.3, 6.9 Hz, 2H, **H8**), 1.89 (q, *J* = 3.2 Hz, 3H, **H26, H28, H31**), 1.79 (s, 2H, **H23**), 1.67 – 1.44 (m, 20H, **H25, H27, H29, H30, H32, H33, H35, H38, H39, H42**) ppm. $^{13}$**C NMR** (101 MHz, DMSO): δ 171.5 (**C11**), 171.1 (**C9**), 169.8 (**C22**), 157.2 (**C10**), 148.6 (**C17**), 147.2 (**C18**), 146.9 (**C5**), 136.9 (**C2**), 133.1 (**C14**), 131.5 (**C4**), 126.7 (**C1**), 120.7 (**C3**), 120.1 (**C6**), 120.0 (**C15**), 112.2 (**C19**), 111.9 (**C16**), 69.9 (**C40**), 69.8 (**C37**), 67.7 (**C41**), 67.5 (**C36**), 55.5 (**C20**), 55.4 (**C21**), 50.1 (**C23**), 42.1 (**C25, C27, C29, C30, C32, C33**), 37.3 (**C8**), 37.1 (**C12**), 36.5 (**C25, C27, C29, C30, C32, C33**), 35.7 (**C43**), 35.6 (**C34**), 32.1 (**C24**), 31.1 (**C7**), 30.0 (**C13**), 29.5 (**C35/C38/C39/C42**), 29.4 (**C35/C38/C39/C42**), 28.0 (**C26, C28, C31)**, 26.0 (**C35/C38/C39/C42**), 26.0 (**C35/C38/C39/C42**) ppm. LCMS: **HRMS (ESI):** m/z $[M + H]^+$ calcd for $C_{43}H_{61}N_4O_7S^+$ 777.4261; found 777.4265.

*N*-(6-(1-((3*r*,5*r*,7*r*)-adamantan-1-yl)-2,18-dioxo-7,10,13-trioxa-3,17-diazaicosan-20-yl)benzo[*d*]thiazol-2-yl)-3-(3,4-dimethoxyphenyl)propanamide **24**

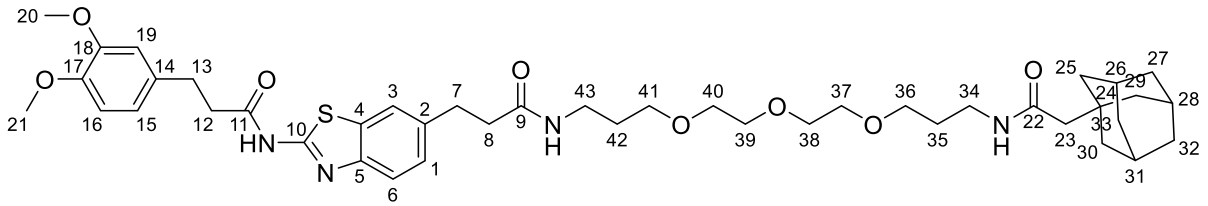

According to general procedure B, SI22 (0.07 mmol) and SI9 (0.08 mmol) provided the title compound as a white solid (44.00 mg, 77%). **Rf** = 0.37 (DCM/MeOH 9:1); **$^1$H NMR** (400 MHz, DMSO): δ 12.27 (s, 1H, C10-**NH**-C11), 7.82 – 7.73 (m, 2H, C9-**NH**-C43, **H3**), 7.70 – 7.58 (m, 2H, C22-**NH**-C34, **H6**), 7.25 (dd, *J* = 8.3, 1.7 Hz, 1H, **H1**), 6.88 – 6.81 (m, 2H, **H16, H19**), 6.74 (dd, *J* = 8.2, 2.0 Hz, 1H, **H15**), 3.71 (s, 3H, **H20**), 3.70 (s, 3H, **H21**), 3.52 – 3.43 (m, 6H, **H38-H40**), 3.43 – 3.36 (m, 4H, **H37, H41**), 3.29 (t, *J* = 6.4 Hz, 2H, **H36**), 3.05 (qd, *J* = 6.7, 2.5 Hz, 4H, **H34, H43**), 2.89 (dt, *J* = 10.5, 7.4 Hz, 4H, **H7, H13**), 2.81 – 2.72 (m, 2H, **H12**), 2.40 (dd, *J* = 8.4, 6.9 Hz, 2H, **H8**), 1.89 (dd, *J* = 5.8, 3.0 Hz, 3H, **H26, H28, H31**), 1.79 (s, 2H, **H23**), 1.67 – 1.51 (m, 16H, **H25, H27, H29, H30, H32, H33, H35, H42**) ppm. **$^{13}$C NMR** (101 MHz, DMSO): δ 171.5 (**C11**), 171.1 (**C9**), 169.8 (**C22**), 157.2 (**C10**), 148.6 (**C17**), 147.2 (**C18**), 146.9 (**C5**), 136.9 (**C2**), 133.1 (**C14**), 131.5 (**C4**), 126.7 (**C1**), 120.7 (**C3**), 120.1 (**C6**), 120.0 (**C15**), 112.2 (**C19**), 111.9 (**C16**), 69.7 (**C37-C40**), 69.5 (**C37-C40**), 69.5 (**C37-C40**), 68.1 (**C41**), 68.0 (**C36**), 55.5 (**C20**), 55.4 (**C21**), 50.1 (**C23**), 42.1 (**C25, C27, C29, C30, C32, C33**), 37.3 (**C8**), 37.2 (**C12**), 36.5 (**C25, C27, C29, C30, C32, C33**), 35.7 (**C43**), 35.6 (**C34**), 32.2 (**C24**), 31.1 (**C7**), 30.0 (**C13**), 29.5 (**C35/C42**), 29.3(**C35/C42**), 28.0 (**C26, C28, C31)** ppm. **HRMS (ESI):** m/z [M + H]$^+$ calcd for $C_{43}H_{61}N_4O_8S^+$ 793.4210; found 793.4211.

*N*-(6-(2-((3*r*,5*r*,7*r*)-adamantan-1-yl)acetamido)hexyl)-3-(2-(3-(3,4-dimethoxyphenyl)propanamido)benzo[*d*]thiazol-6-yl)propanamide **25**

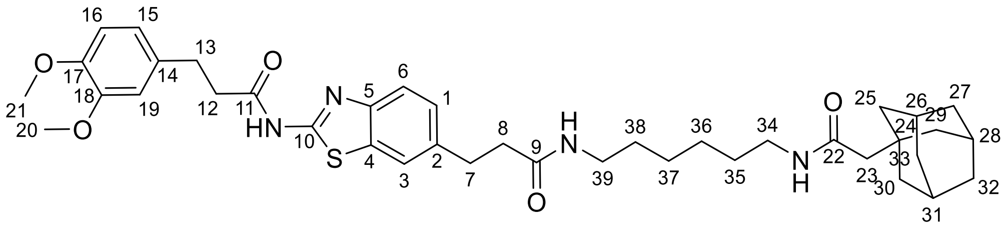

According to general procedure B, SI22 (0.07 mmol) and SI10 (0.08 mmol) provided the title compound as a white solid (40.00 mg, 80%). **Rf** = 0.67 (DCM/MeOH 9:1); **$^1$H NMR** (400 MHz, DMSO): δ 12.27 (s, 1H, C10-**NH**-C11), 7.80 – 7.72 (m, 2H, C9-**NH**-C43, **H3**), 7.64 – 7.57 (m, 2H, C22-**NH**-C34, **H6**), 7.25 (dd, *J* = 8.3, 1.8 Hz, 1H, **H1**), 6.88 – 6.81 (m, 2H, **H16, H19**), 6.74 (dd, *J* = 8.2, 2.0 Hz, 1H, **H15**), 3.71 (s, 3H, **H20**), 3.70 (s, 3H, **H21**), 2.98 (p, *J* = 6.6 Hz, 4H, **H34, H39**), 2.89 (td, *J* = 8.8, 6.7 Hz, 4H, **H7, H13**), 2.81 – 2.72 (m, 2H, **H12**), 2.39 (dd, *J* = 8.4, 6.9 Hz, 2H, **H8**), 1.92 – 1.85 (m, 3H, **H26, H28, H31**), 1.79 (s, 2H, **H23**), 1.67 – 1.51 (m, 12H, **H25, H27, H29, H30, H32, H33**), 1.37 – 1.25 (m, 4H, **H35, H38**), 1.25 – 1.11 (m, 4H, **H36, H37**) ppm.**$^{13}$C NMR** (101 MHz, DMSO): δ 171.5 (**C11**), 171.1 (**C9**), 169.7 (**C22**), 157.2 (**C10**), 148.6 (**C17**), 147.2 (**C18**), 146.9 (**C5**), 137.0 (**C2**), 133.1 (**C14**), 131.5 (**C4**), 126.7 (**C1**), 120.7 (**C3**), 120.1 (**C6**), 120.0 (**C15**), 112.2 (**C19**), 111.9 (**C16**), 55.5 (**C20**), 55.4 (**C21**), 50.1 (**C23**), 42.1 (**C25, C27, C29, C30, C32, C33**), 38.3 (**C34/C39**), 38.2 (**C34/C39**), 37.4 (**C8**), 37.2 (**C12**), 36.5 (**C25, C27, C29, C30, C32, C33**), 32.1 (**C24**), 31.1 (**C7**), 30.0 (**C13**), 29.1 (**C37, C38**),

28.0 (**C26, C28, C31**), 26.1 (**C36/C37**), 26.0 (**C36/C37**) ppm. **HRMS (ESI):** m/z $[M + H]^+$ calcd for $C_{39}H_{53}N_4O_5S^+$ 689.3736; found 689.3742.

5-((6-(3-(2-(3-(3,4-dimethoxyphenyl)propanamido)benzo[d]thiazol-6-yl)propanamido)hexyl)carbamoyl)-2-(6-(dimethylamino)-3-(dimethyliminio)-3*H*-xanthen-9-yl)benzoate **27**

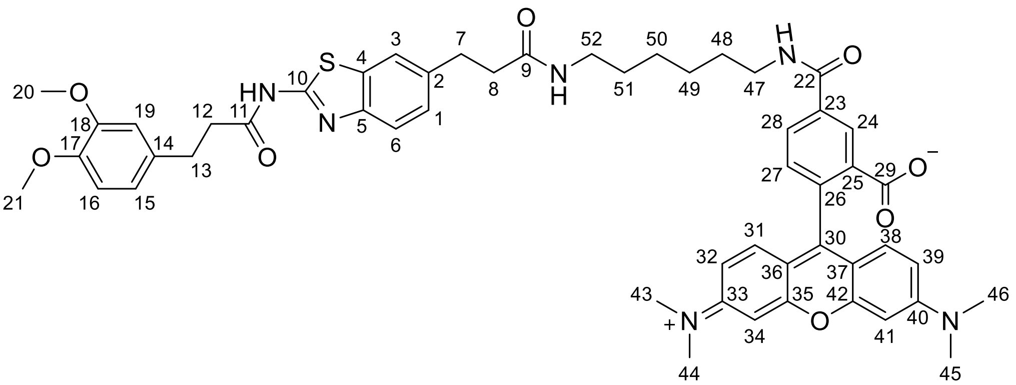

**Purple solid (50 mg, 42%); Rf = 0.45 (DCM/MeOH/$Et_3N$ 90/8/2); $^1$H NMR** (400 MHz, MeOD): δ 8.69 (t, *J* = 5.7 Hz, 1H, NH, C22-**NH**-C47), 8.65 (d, *J* = 1.8 Hz, 1H, **H24**), 8.14 (dd, *J* = 7.9, 1.9 Hz, 1H, **H28**), 7.88 (t, *J* = 5.8 Hz, 1H, NH, C9-**NH**-C52), 7.65 (d, *J* = 1.7 Hz, 1H, **H3**), 7.59 (d, *J* = 8.3 Hz, 1H, **H6**), 7.39 (d, *J* = 7.9 Hz, 1H, **H27**), 7.26 (dd, *J* = 8.3, 1.8 Hz, 1H, **H1**), 7.14 (d, *J* = 9.5 Hz, 2H, **H31, H38**), 6.92 (dd, *J* = 9.5, 2.5 Hz, 2H, **H32, H39**), 6.86 (d, *J* = 2.4 Hz, 2H, **H34, H41**), 6.82 – 6.78 (m, 2H, **H16, H19**), 6.73 (dd, *J* = 8.2, 2.0 Hz, 1H, **H15**), 3.74 (s, 3H, **H20**), 3.73 (s, 3H, **H21**), 3.36 (t, *J* = 7.1 Hz, 2H, **H47**), 3.24 (s, 12H, **H43-H46**), 3.15 – 3.04 (m, 2H, **H52**), 3.01 (t, *J* = 7.2 Hz, 2H, **H7**), 2.92 (dd, *J* = 14.4, 6.8 Hz, 2H, **H13**), 2.79 – 2.71 (m, 2H, **H12**), 2.51 (t, *J* = 7.2 Hz, 2H, **H8**), 1.53 (p, *J* = 7.2 Hz, 2H, **H48**), 1.39 – 1.32 (m, 2H, **H51**), 1.29 – 1.23 (m, 2H), 1.20 – 1.10 (m, 2H) ppm. One NH missing due to deuterium exchange (C10-**NH**-C11). **$^{13}$C NMR** (101 MHz, MeOD): δ 174.9 (**C9**), 173.6 (**C11**), 170.4 (**C29**), 168.6 (**C22**), 161.5 (**C30**), 159.2 (**C33/C40**), 158.9 (**C33/C40**), 158.7, 150.4 (**C17/C18**), 148.4 (**C17/C18**), 148.1 (**C5**), 138.1 (**C2**), 137.5, 137.3, 134.7 (**C14**), 133.4 (**C4**), 132.3 (**C31 + C38**), 131.3 (**C27**), 130.6 (**C28**), 130.3 (**C24**), 128.2 (**C1**), 121.8 (**C3**), 121.6 (**C15**), 121.6 (**C6**), 115.2 (**C32/C39**), 114.7 (**C32/C39**), 113.4 (**C16/C19**), 113.1(**C16/C19**), 97.4 (**C34 + C41**), 56.5 (**C20**), 56.4 (**C21**), 41.1 (**C47**), 40.9 (**C43-C46**), 40.2 (**C52**), 39.0 (**C8**), 38.8 (**C12**), 32.8 (**C7**), 31.7 (**C13**), 30.4 (**C51**), 30.3 (**C48**), 27.7 (**C49/C50**), 27.5 (**C49/C50**) ppm. **HRMS (ESI):** m/z $[M + 2H]^+$ calcd for $C_{52}H_{58}N_6O_8S^+$ 463.2013; found 463.2015.

**NMR data**

1-(4-((4-chloro-2-((2-chlorophenyl)(hydroxy)methyl)phenyl)(neopentyl)amino)-4-oxobutanoyl)-N-(2-(2-(2-((2-(2,6-dioxopiperidin-3-yl)-1,3-dioxoisoindolin-4-yl)amino)ethoxy)ethoxy)ethyl)piperidine-4-carboxamide **1**

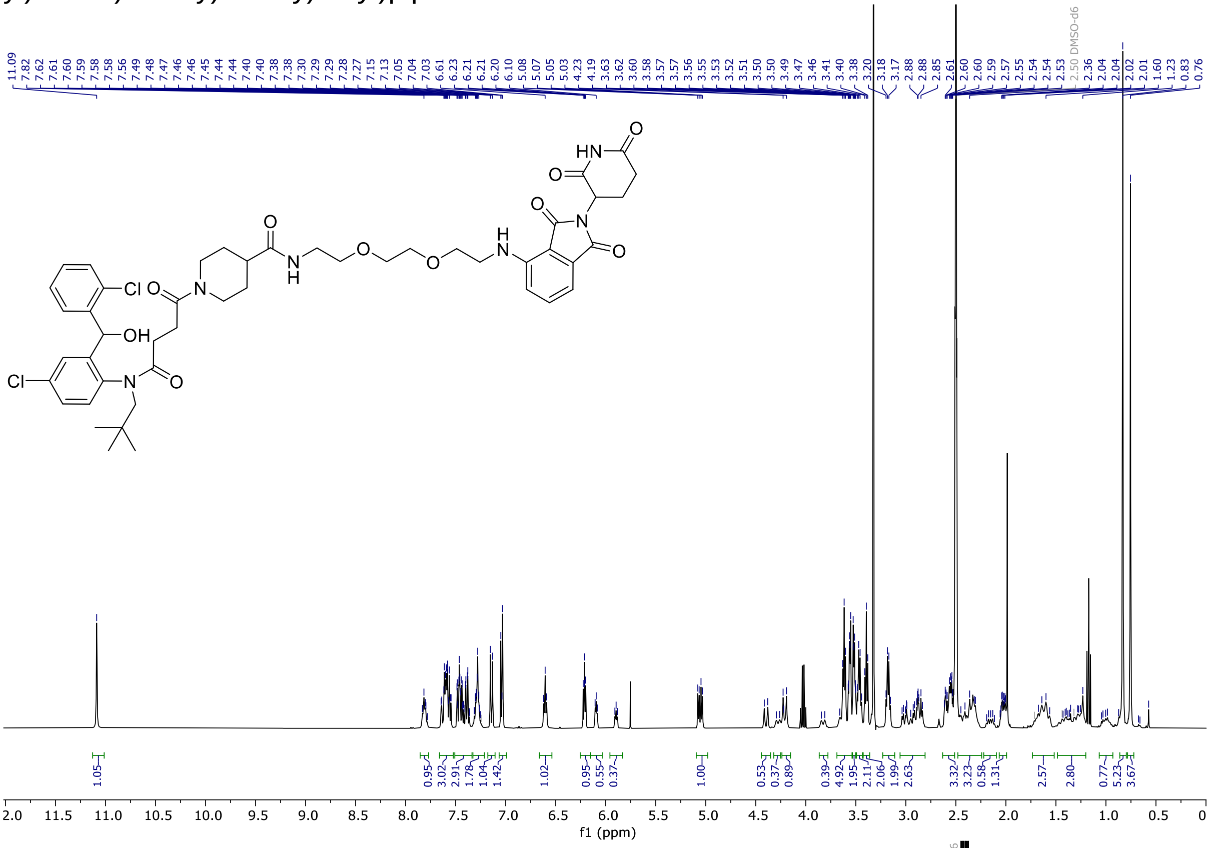


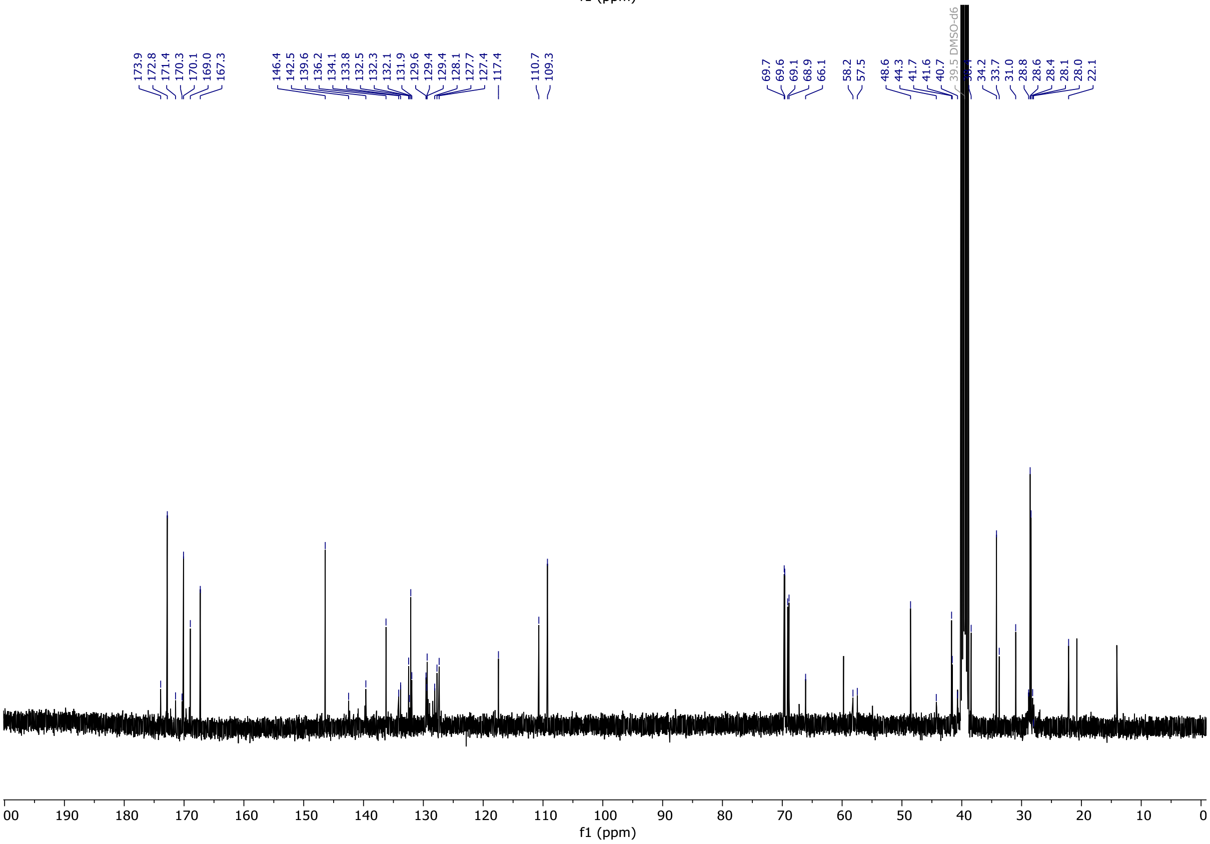

1-(4-((4-chloro-2-((2-chlorophenyl)(hydroxy)methyl)phenyl)(neopentyl)amino)-4-oxobutanoyl)-N-(3-(4-(3-((2-(2,6-dioxopiperidin-3-yl)-1,3-dioxoisoindolin-4-yl)amino)propoxy)butoxy)propyl)piperidine-4-carboxamide **2**

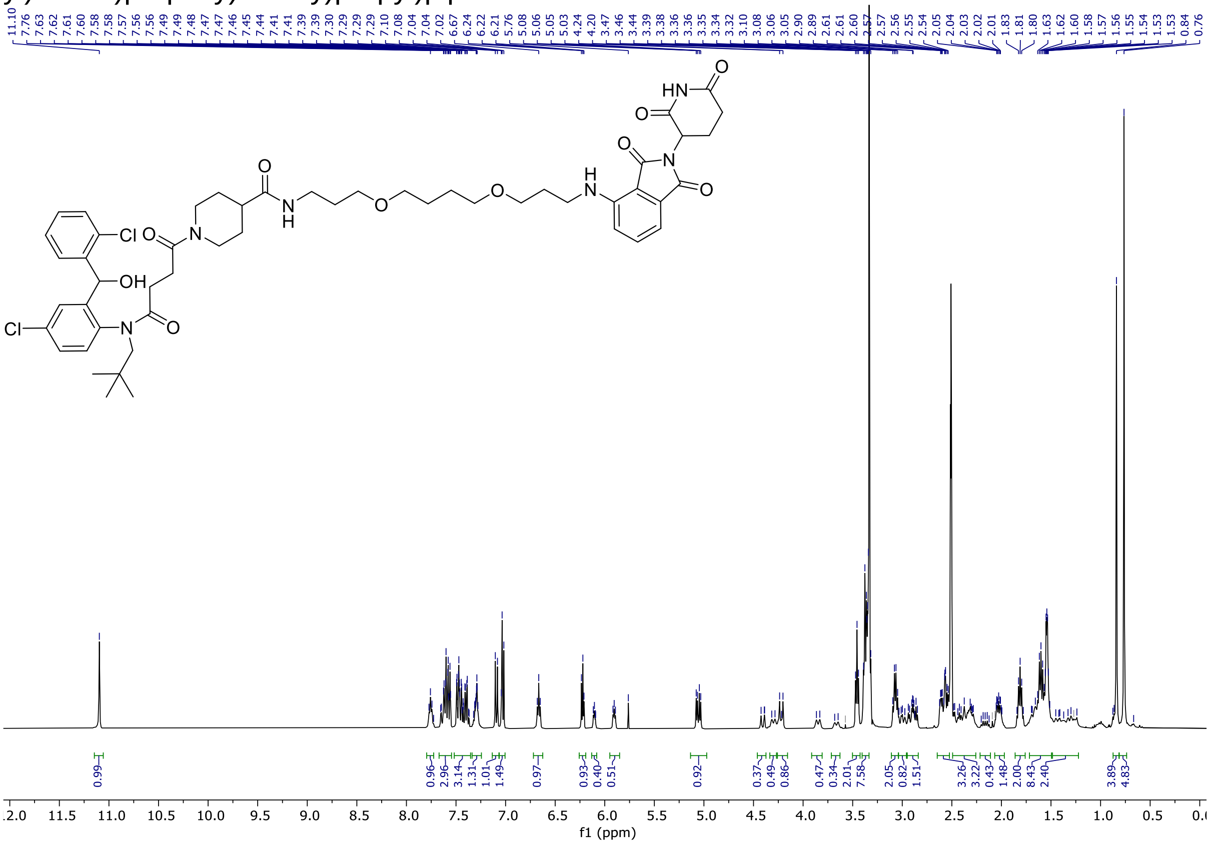

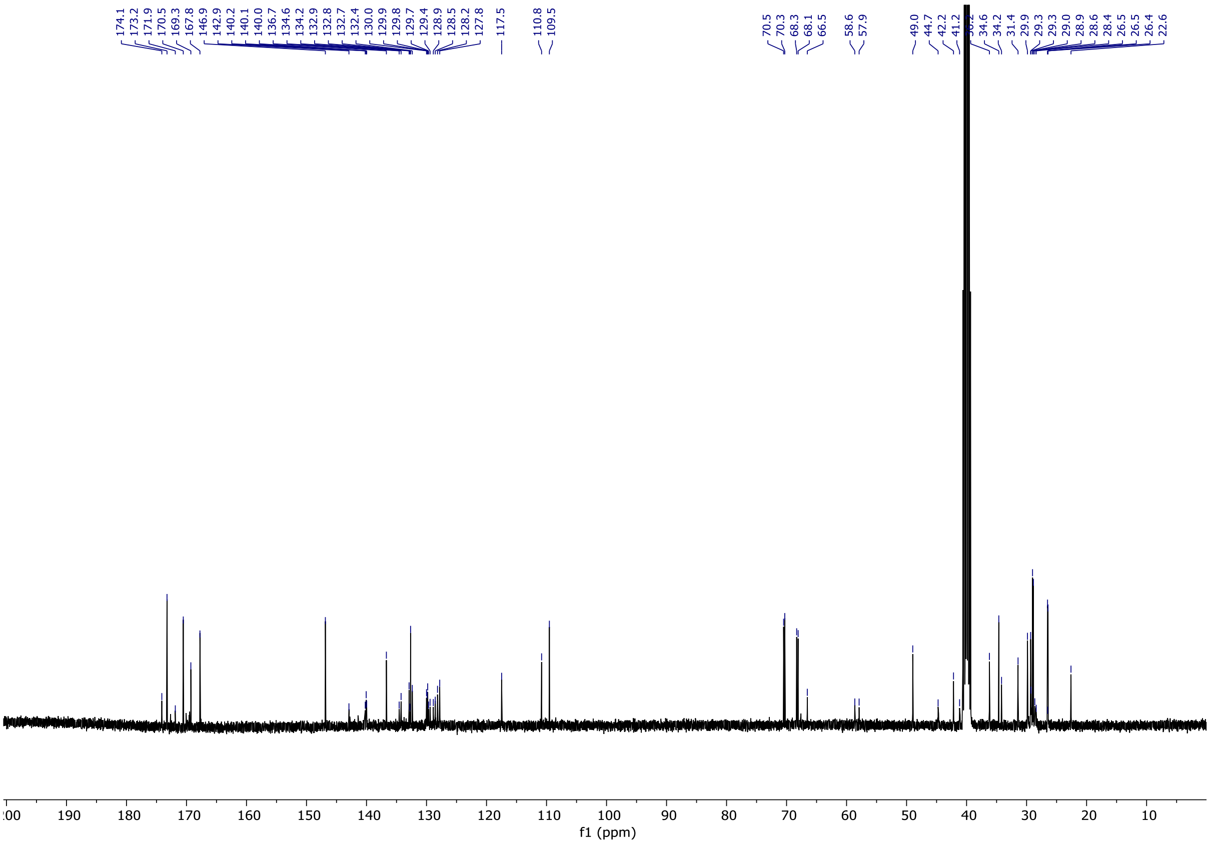

1-(4-((4-chloro-2-((2-chlorophenyl)(hydroxy)methyl)phenyl)(neopentyl)amino)-4-oxobutanoyl)-N-(3(2-(2-(3-((2-(2,6-dioxopiperidin-3-yl)-1,3-dioxoisoindolin-4-yl)amino)propoxy)ethoxy)ethoxy)propyl)piperidine-4-carboxamide **3**

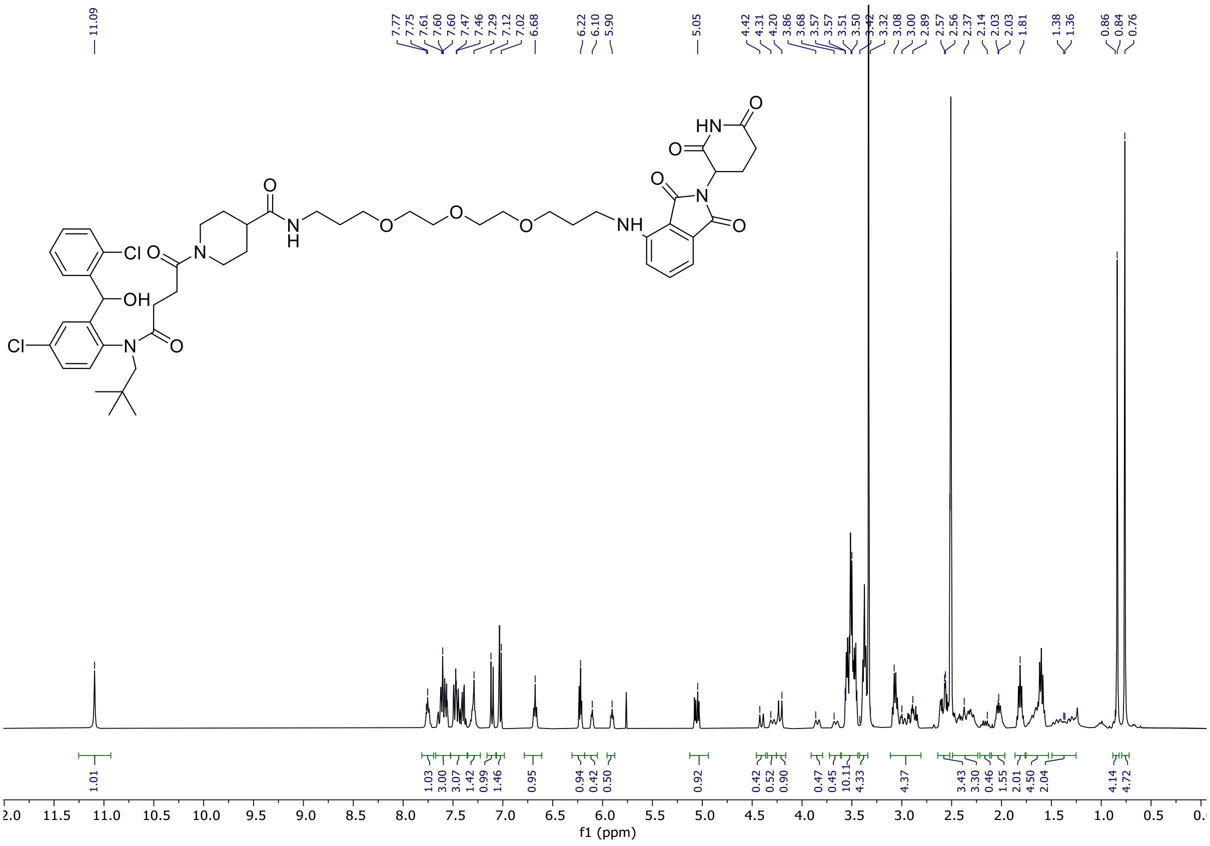


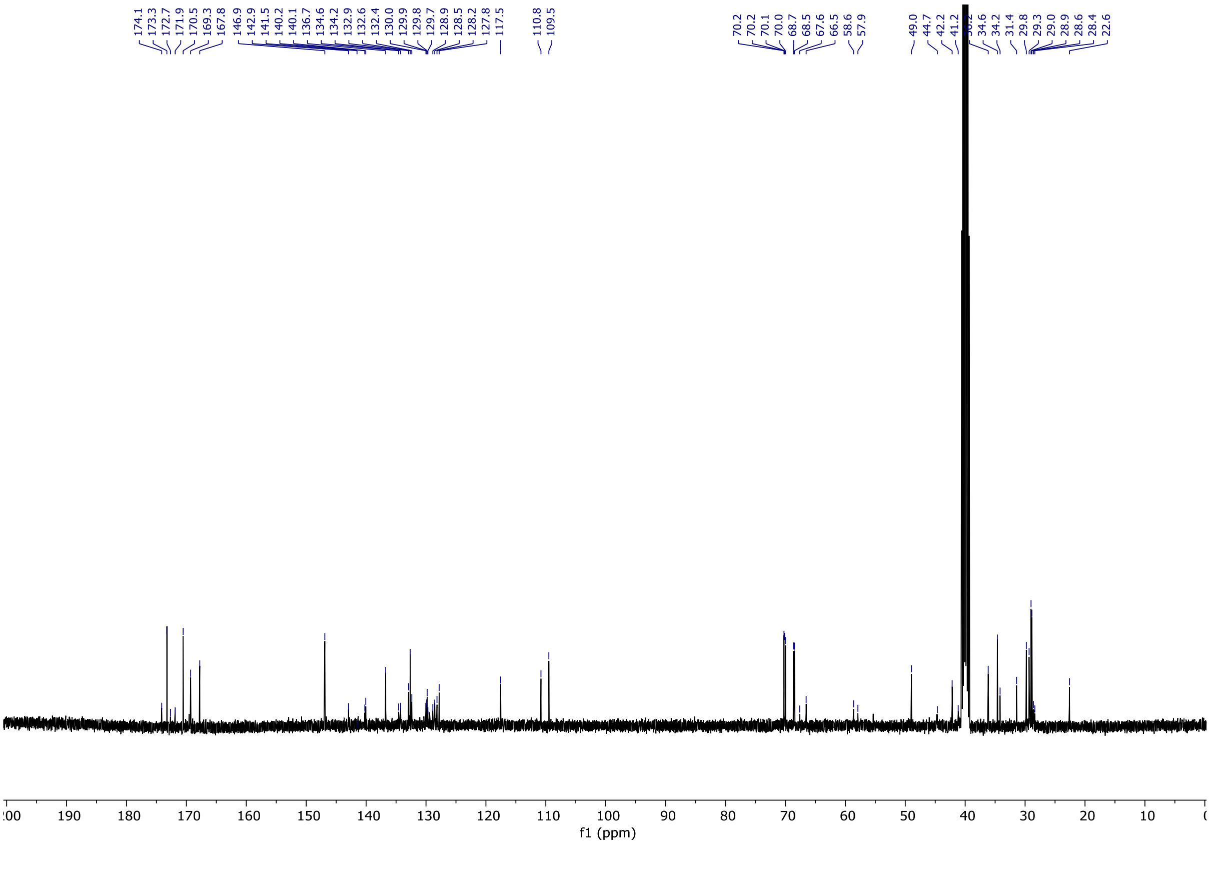

1-(4-((4-chloro-2-((2-chlorophenyl)(hydroxy)methyl)phenyl)(neopentyl)amino)-4-oxobutanoyl)-N-(6-((2-(2,6-dioxopiperidin-3-yl)-1,3-dioxoisoindolin-4-yl)amino)hexyl)piperidine-4-carboxamide **4**

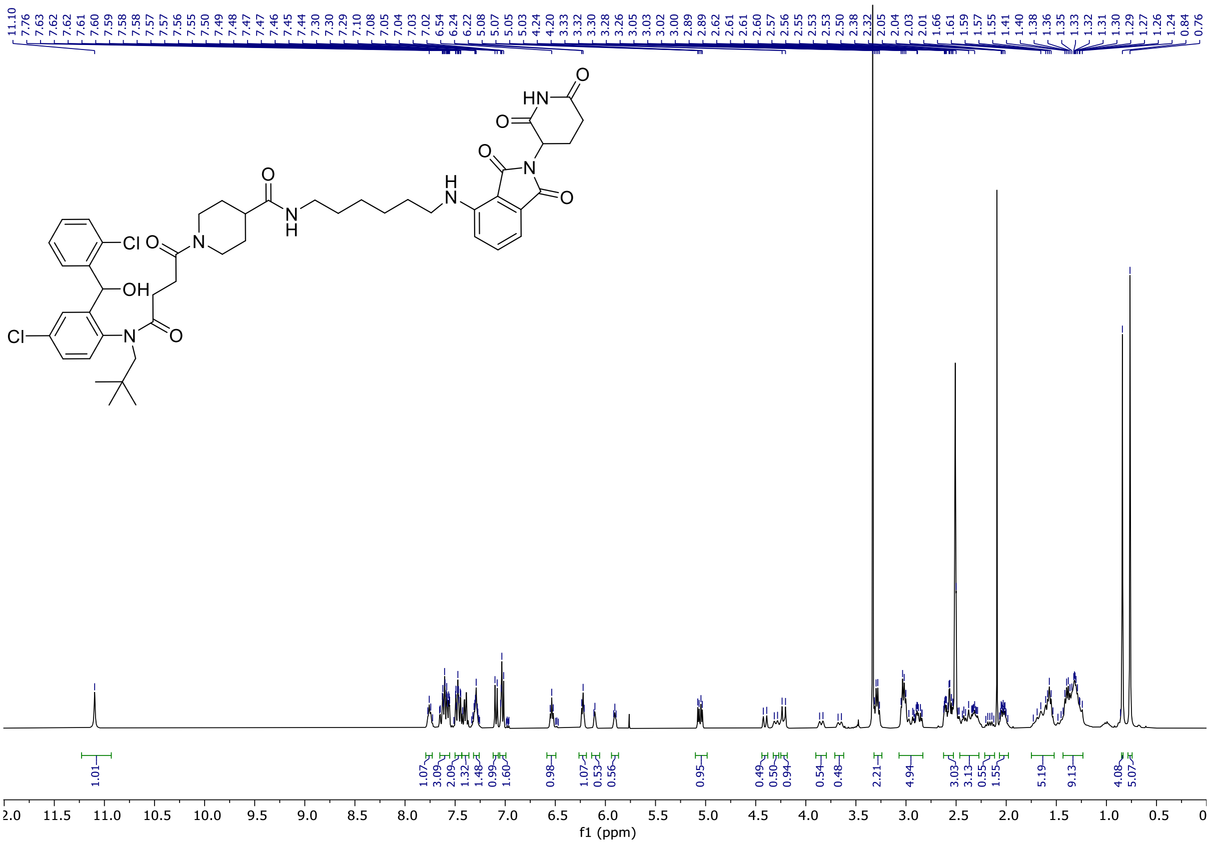


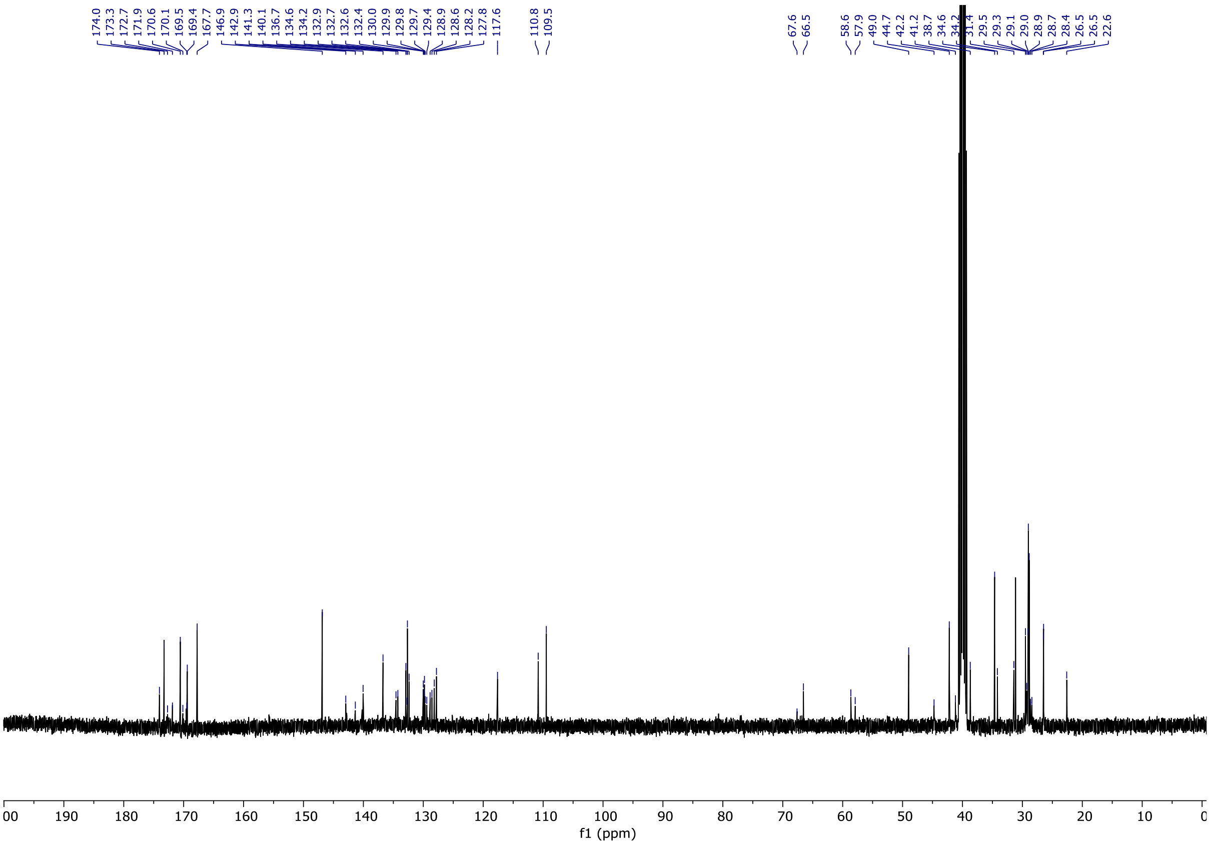

1-(4-((4-chloro-2-((2-chlorophenyl)(hydroxy)methyl)phenyl)(neopentyl)amino)-4-oxobutanoyl)-N-(10-((2-(2,6-dioxopiperidin-3-yl)-1,3-dioxoisoindolin-4-yl)amino)decyl)piperidine-4-carboxamide **5**

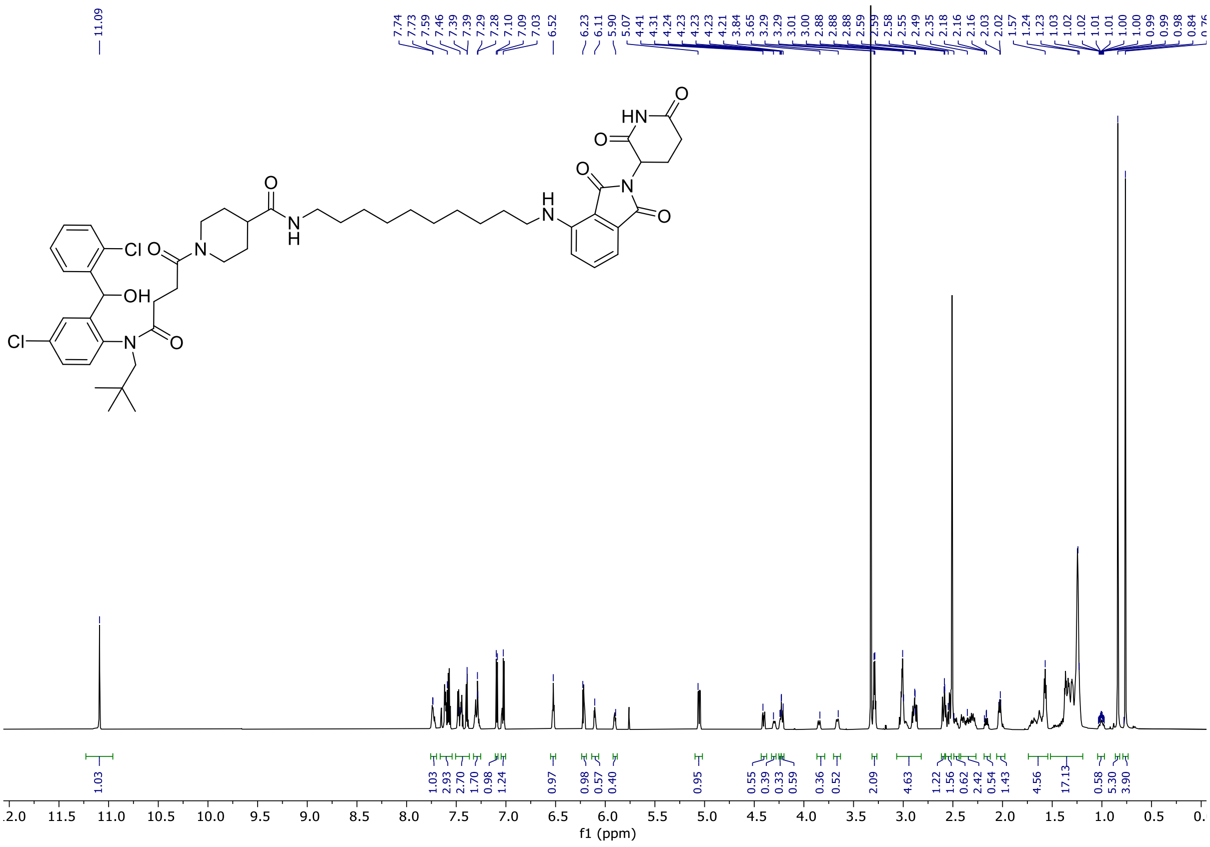


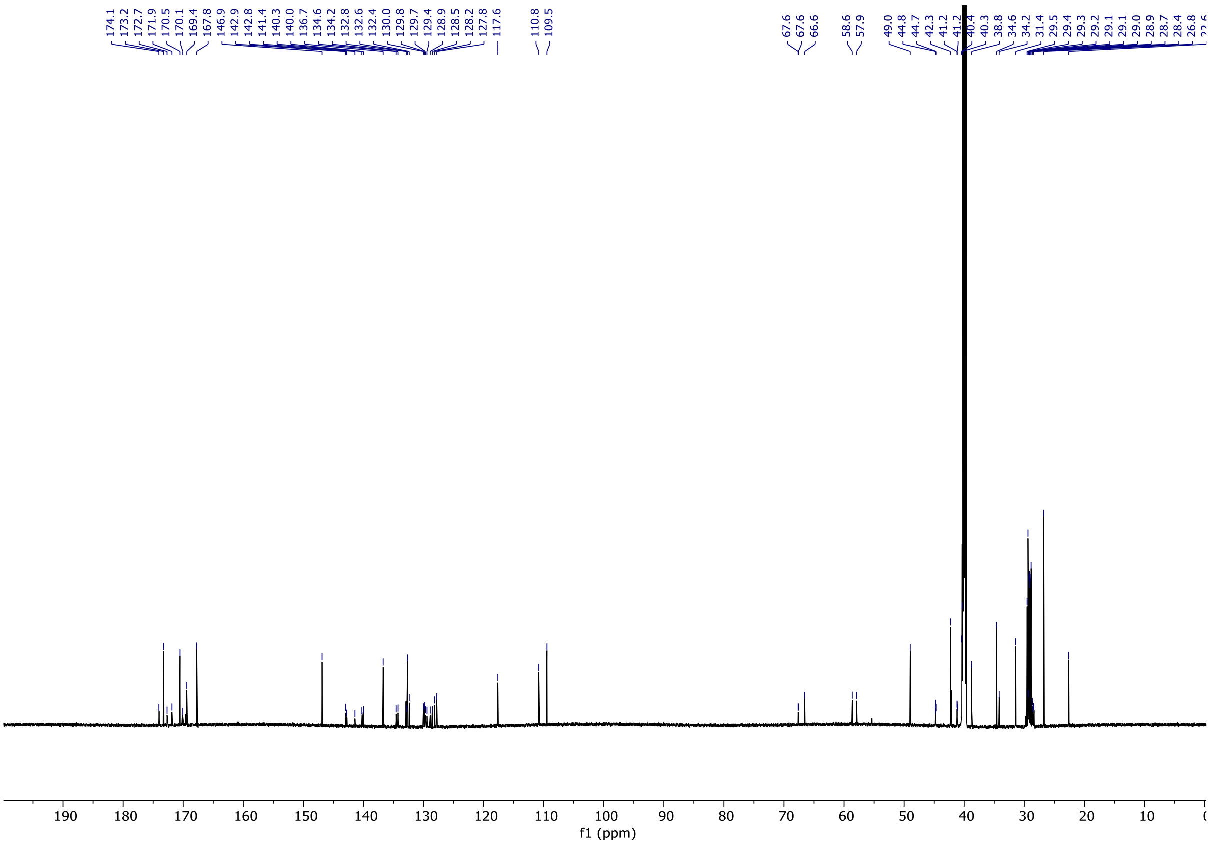

1-(4-((4-chloro-2-((2-chlorophenyl)(hydroxy)methyl)phenyl)(neopentyl)amino)-4-oxobutanoyl)-N-(2-(2-(2-(((S)-1-((2S,4R)-4-hydroxy-2-((4-(4-methylthiazol-5-yl)benzyl)carbamoyl)-pyrrolidin-1-yl)-3,3-dimethyl-1-oxobutan-2-yl)amino)-2-oxoethoxy)ethoxy)ethyl)piperidine-4-carboxamide **6**

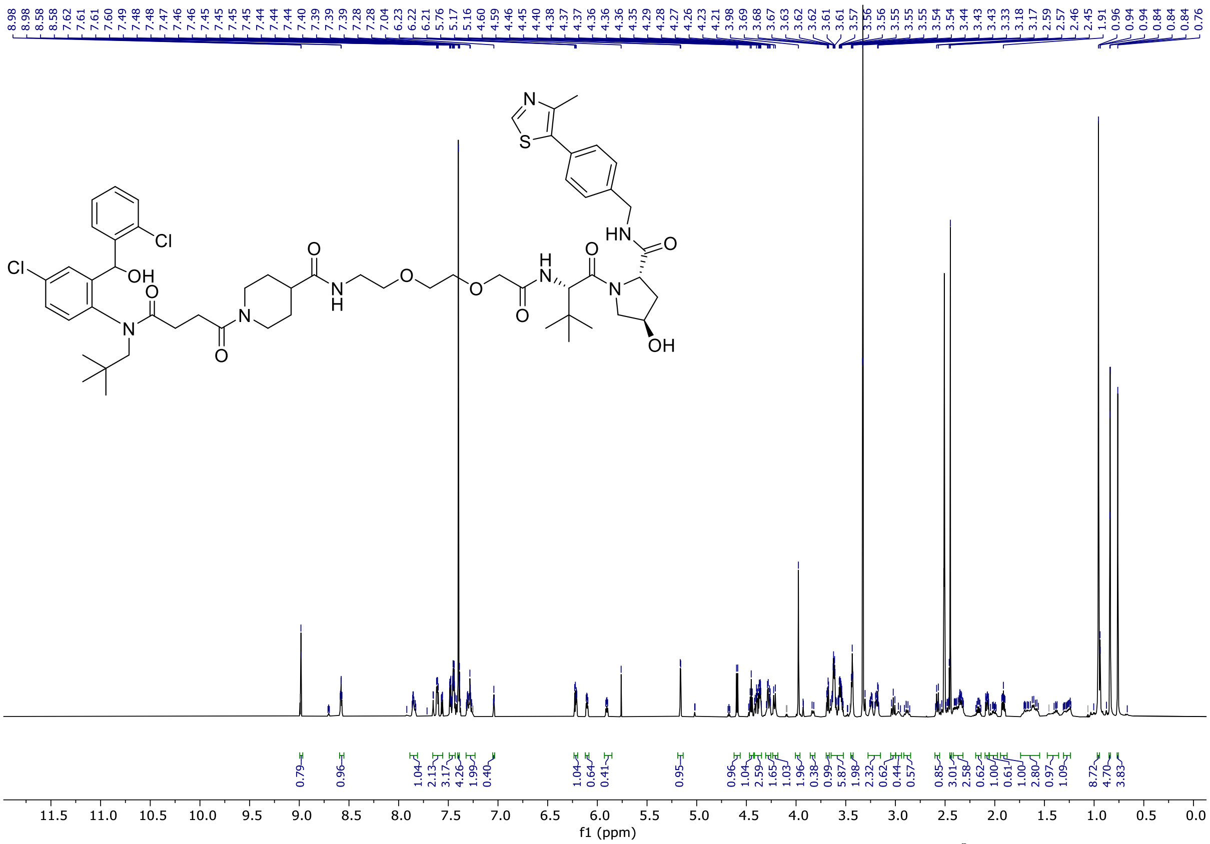


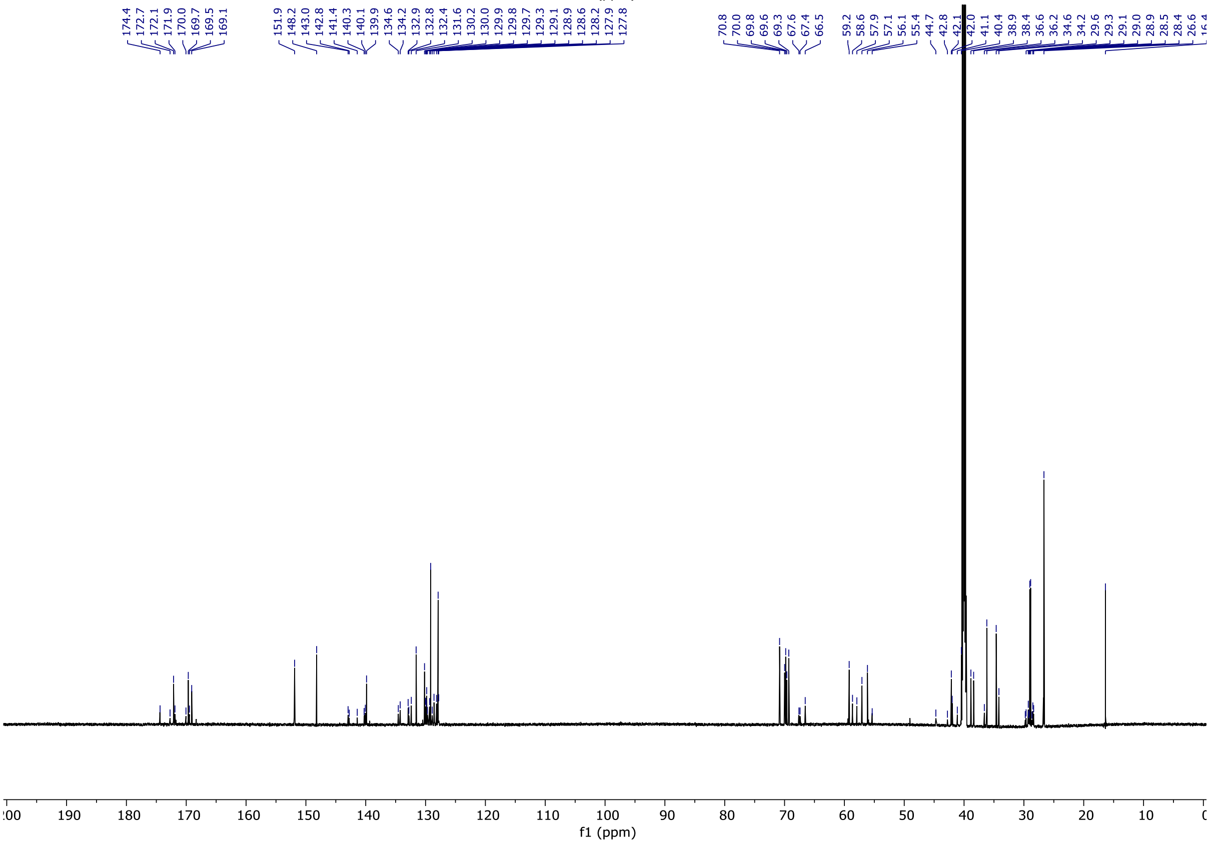

1-(4-((4-chloro-2-((2-chlorophenyl)(hydroxy)methyl)phenyl)(neopentyl)amino)-4-oxobutanoyl)-N-(6-(((S)-1-((2S,4R)-4-hydroxy-2-((4-(4-methylthiazol-5-yl)benzyl)carbamoyl)pyrrolidin-1-yl)-3,3-dimethyl-1-oxobutan-2-yl)amino)-6-oxohexyl)piperidine-4-carboxamide **7**

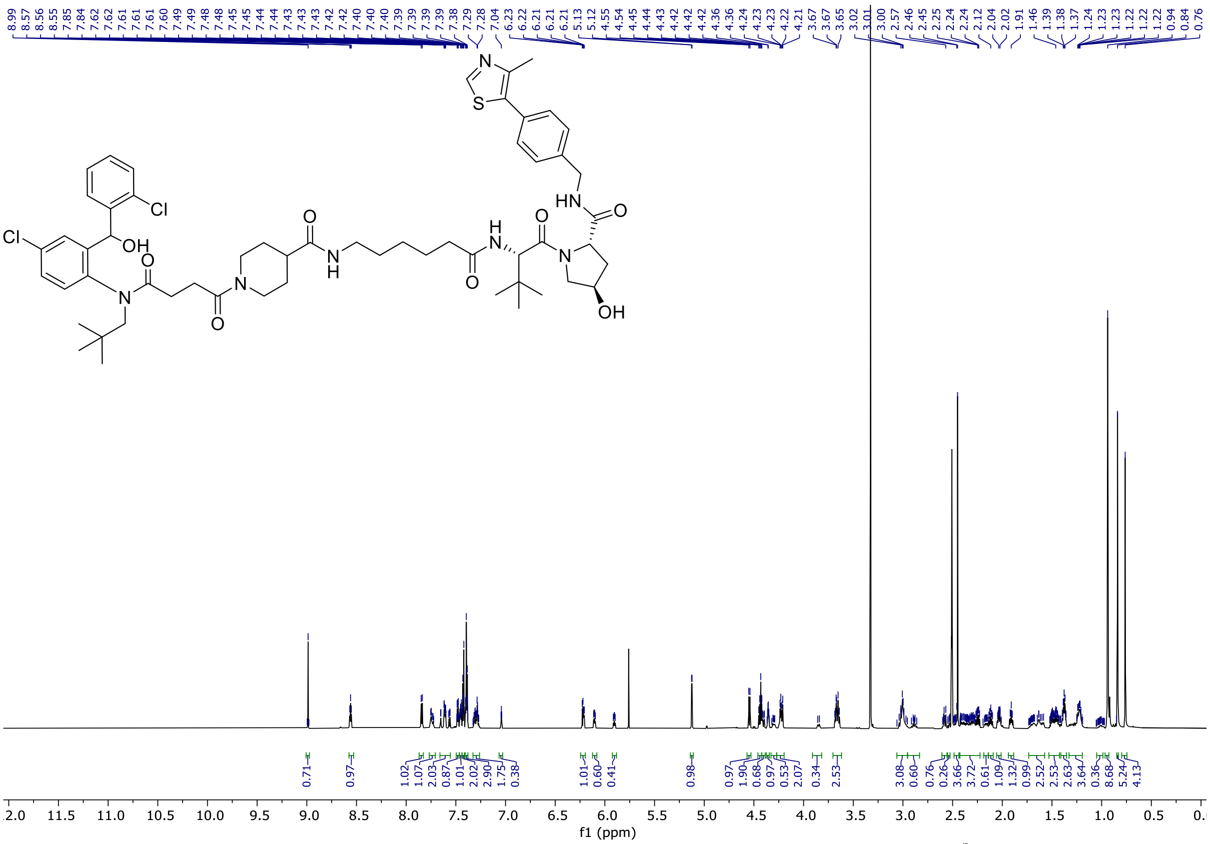

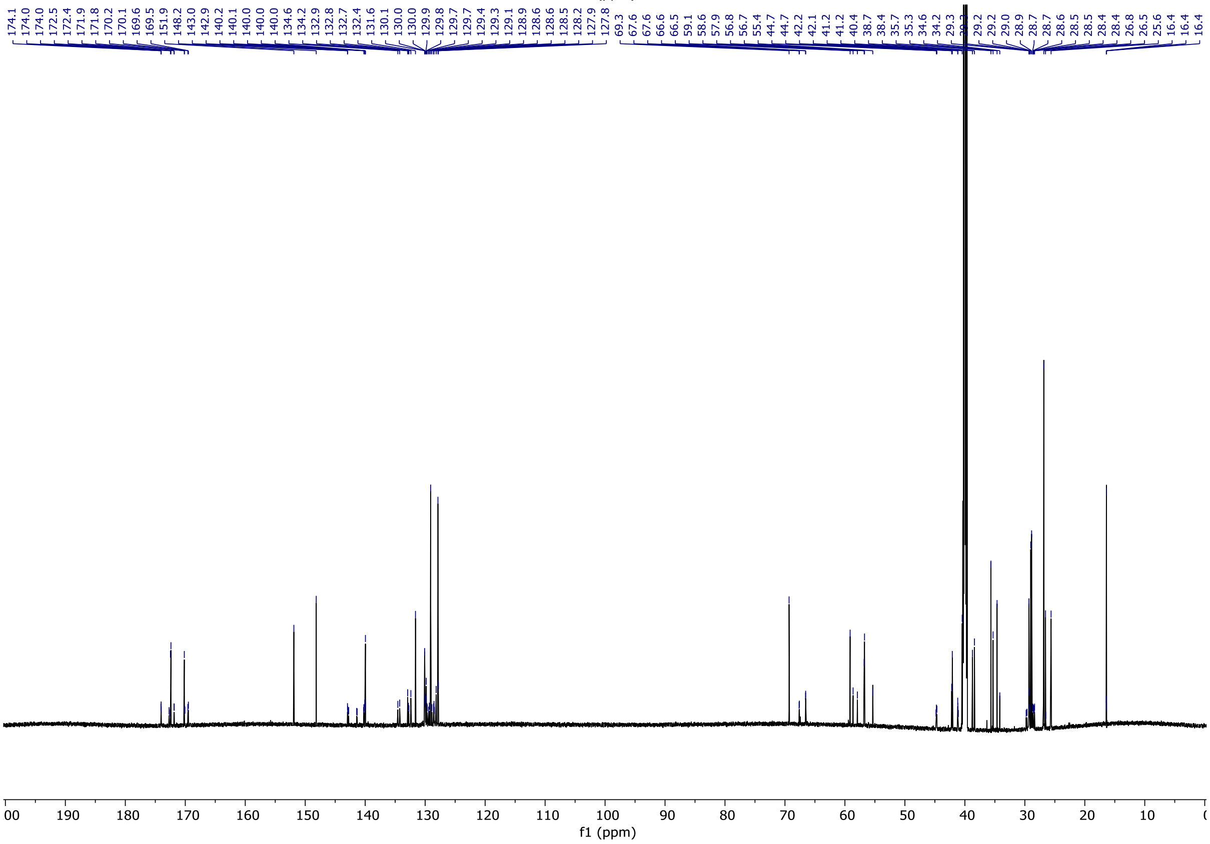

1-(4-((4-chloro-2-((2-chlorophenyl)(hydroxy)methyl)phenyl)(neopentyl)amino)-4-oxobutanoyl)-N-(10-(((S)-1-((2S,4R)-4-hydroxy-2-((4-(4-methylthiazol-5-yl)benzyl)carbamoyl)-pyrrolidin-1-yl)-3,3-dimethyl-1-oxobutan-2-yl)amino)-10-oxodecyl)piperidine-4-carboxamide **8**

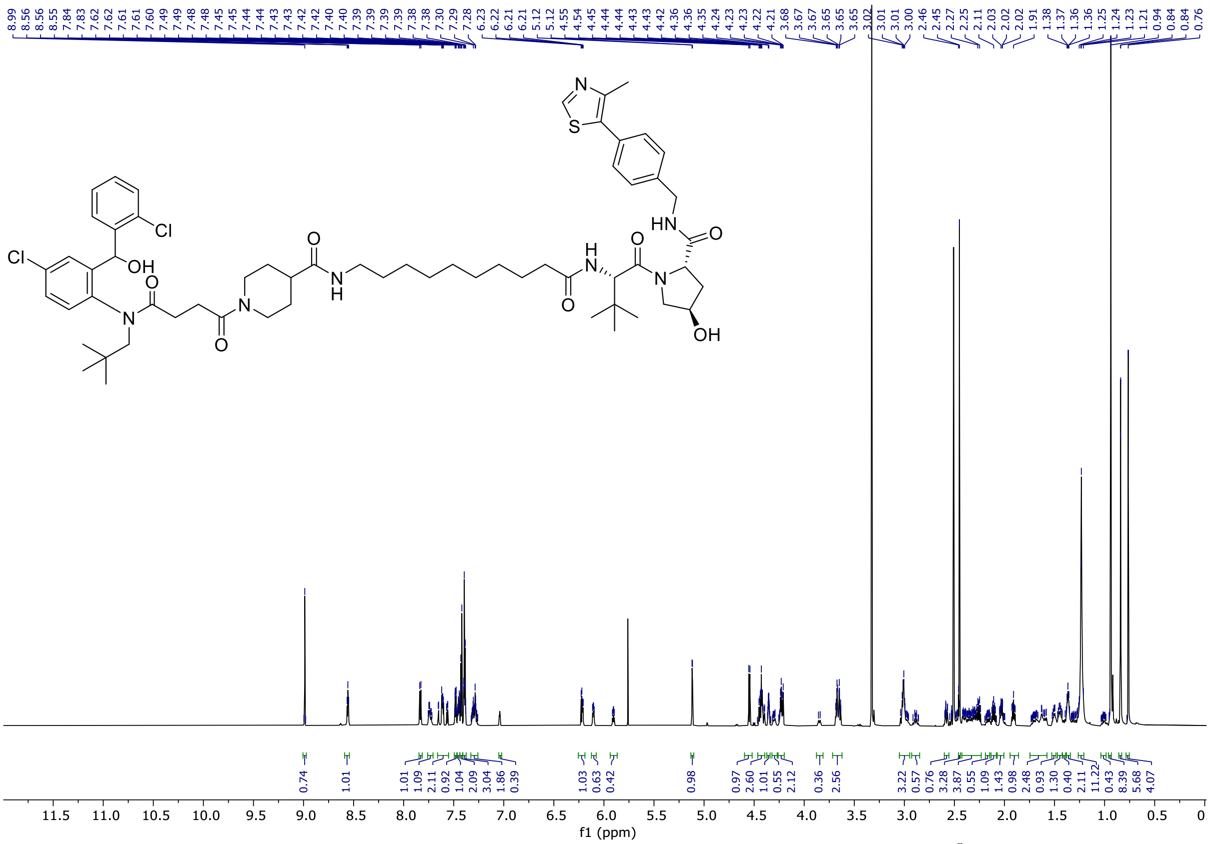


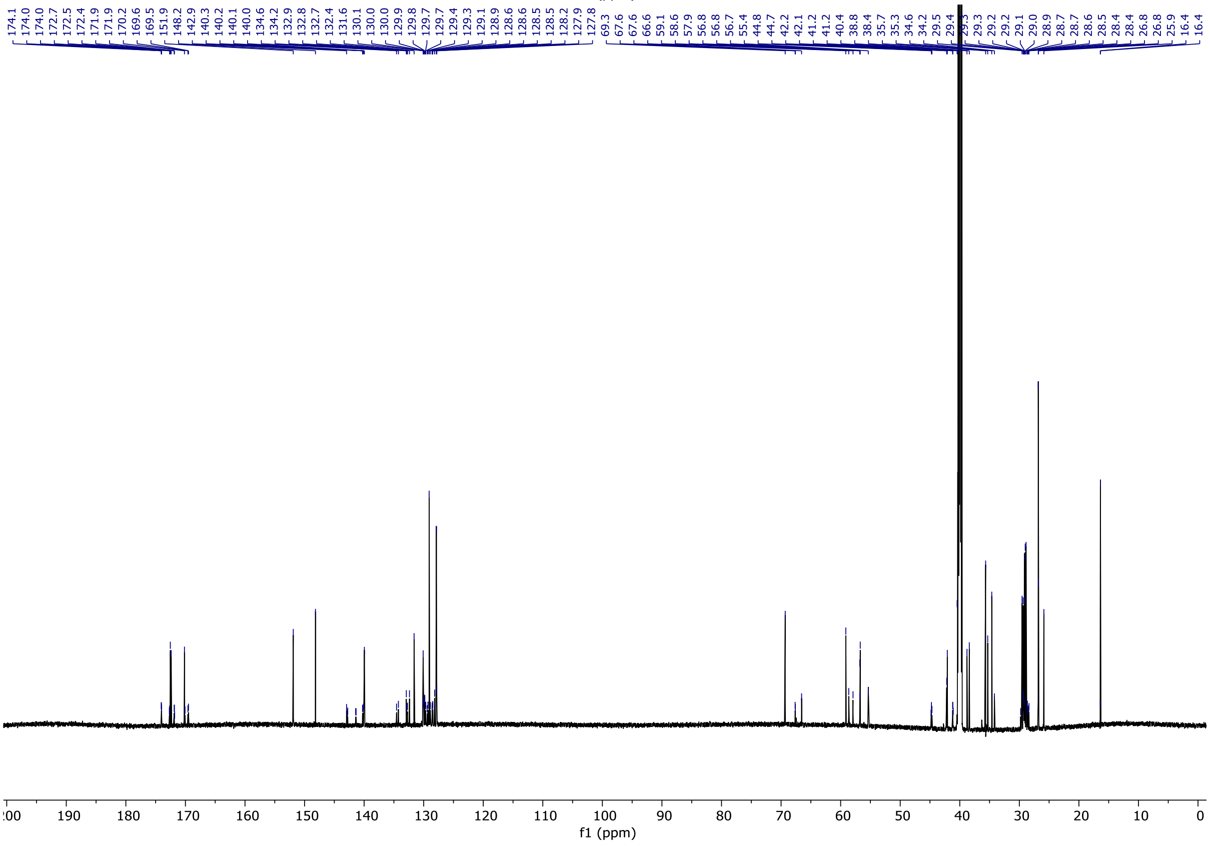

N-(2-(2-(2-(2-((1s,3s)-adamantan-1-yl)acetamido)ethoxy)ethoxy)ethyl)-1-(4-((4-chloro-2-((2-chlorophenyl)(hydroxy)methyl)phenyl)(neopentyl)amino)-4-oxobutanoyl)piperidine-4-carboxamide **9**

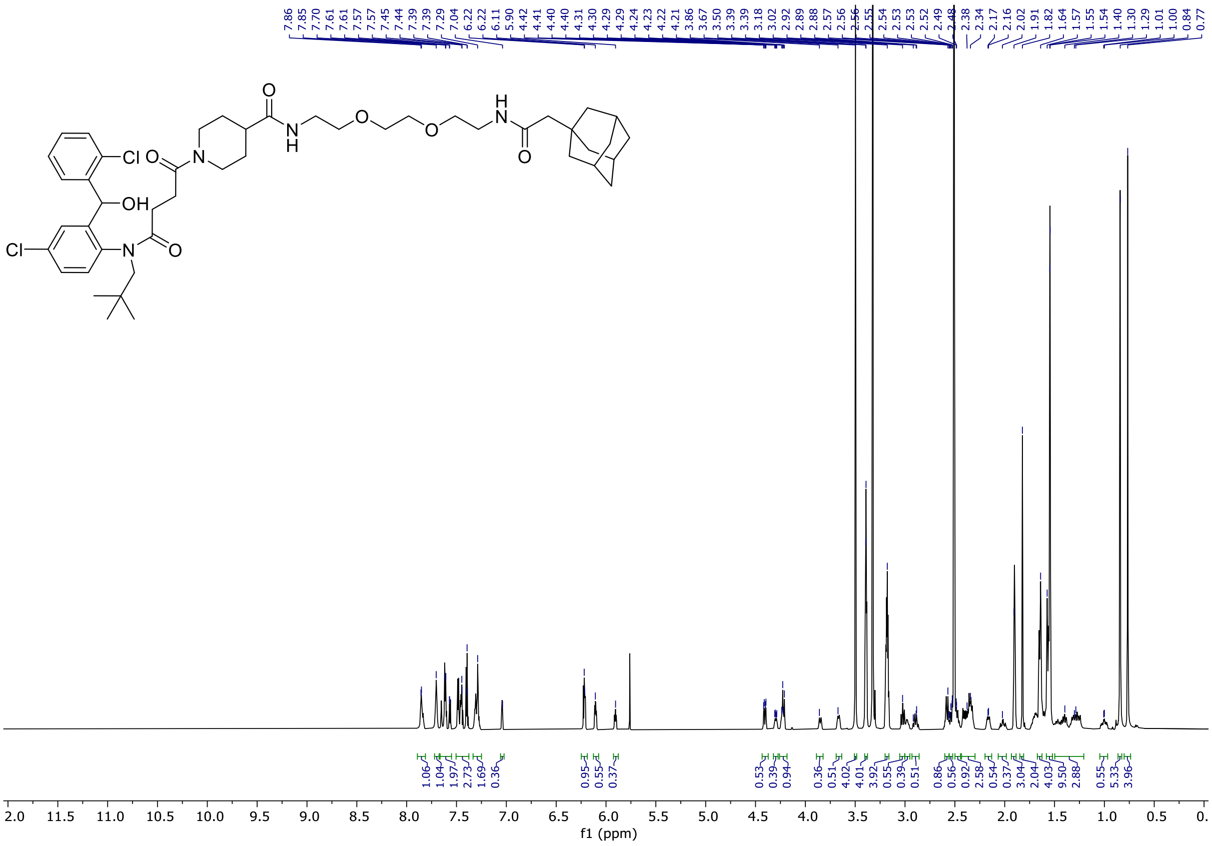


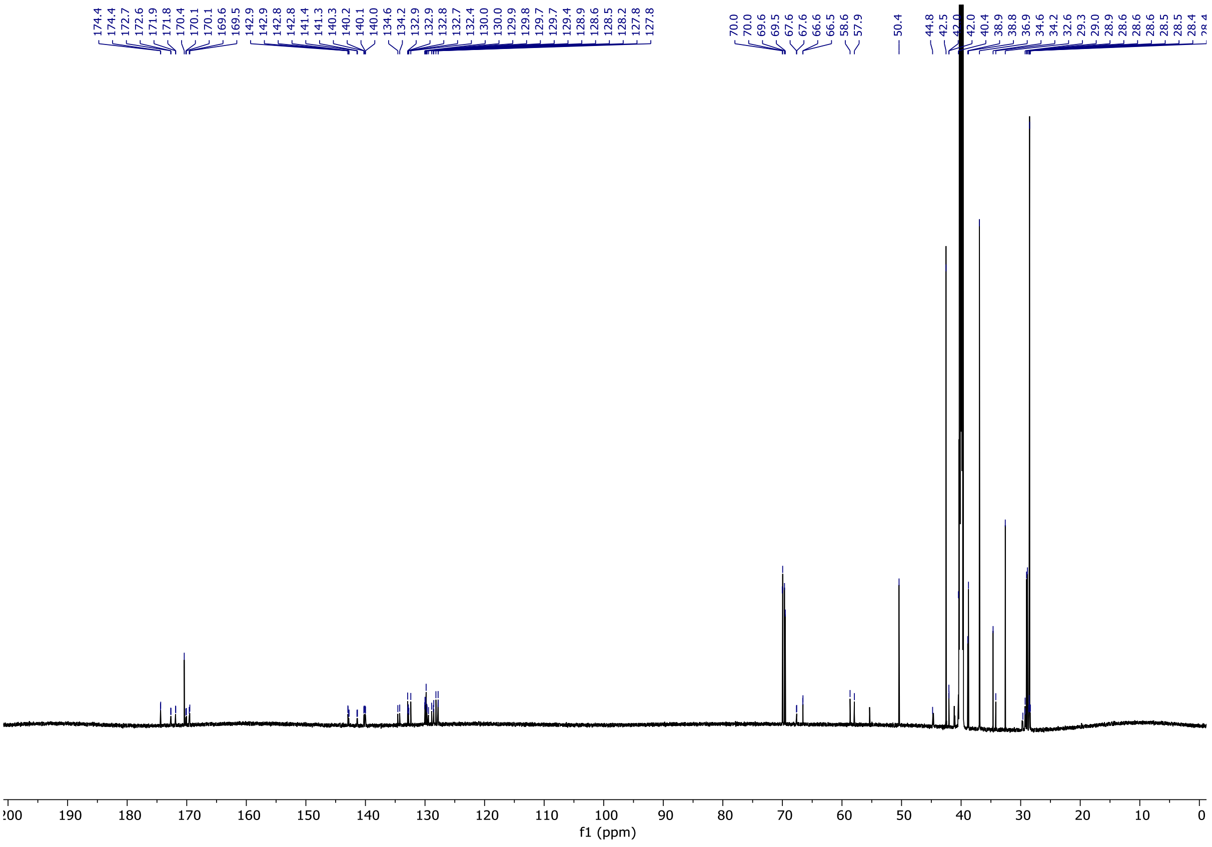

N-(3-(4-(3-(2-((1s,3s)-adamantan-1-yl)acetamido)propoxy)butoxy)propyl)-1-(4-((4-chloro-2-((2-chlorophenyl)(hydroxy)methyl)phenyl)(neopentyl)amino)-4-oxobutanoyl)piperidine-4-carboxamide **10**

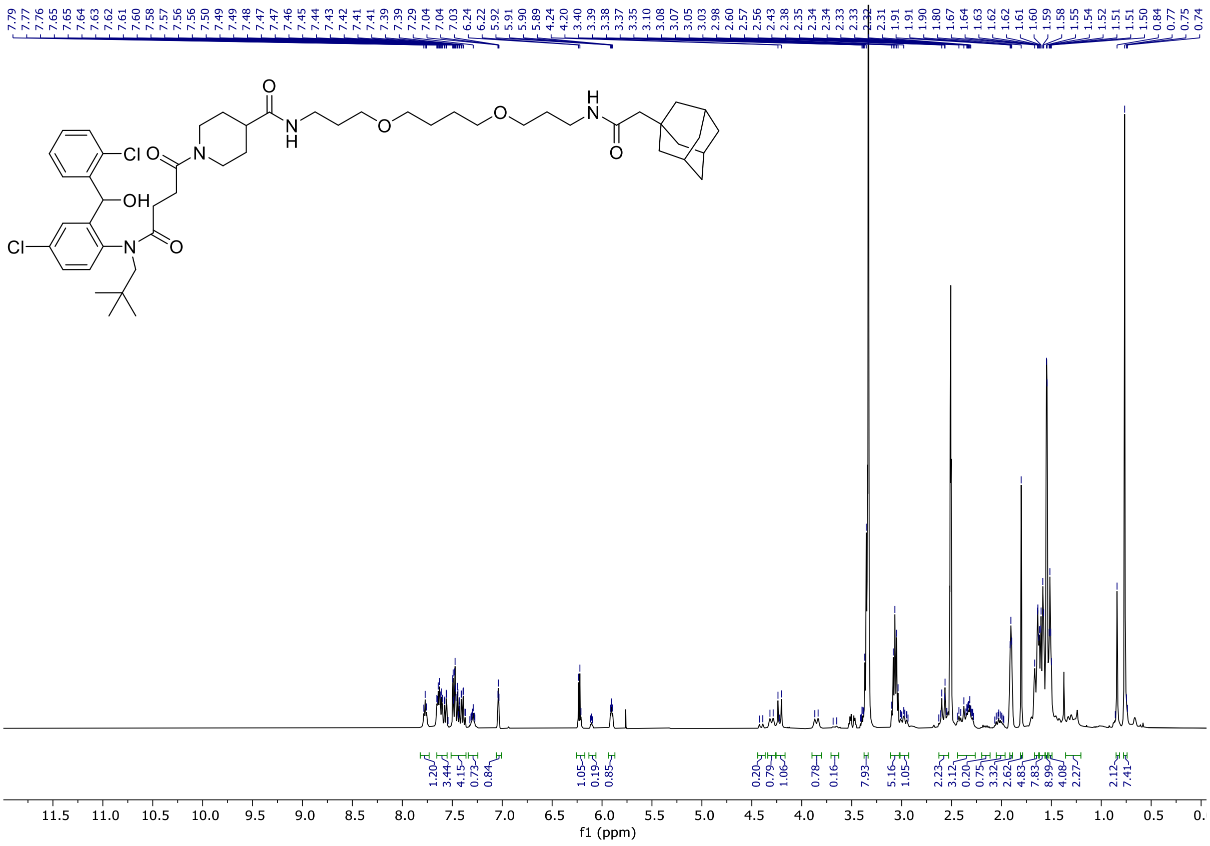

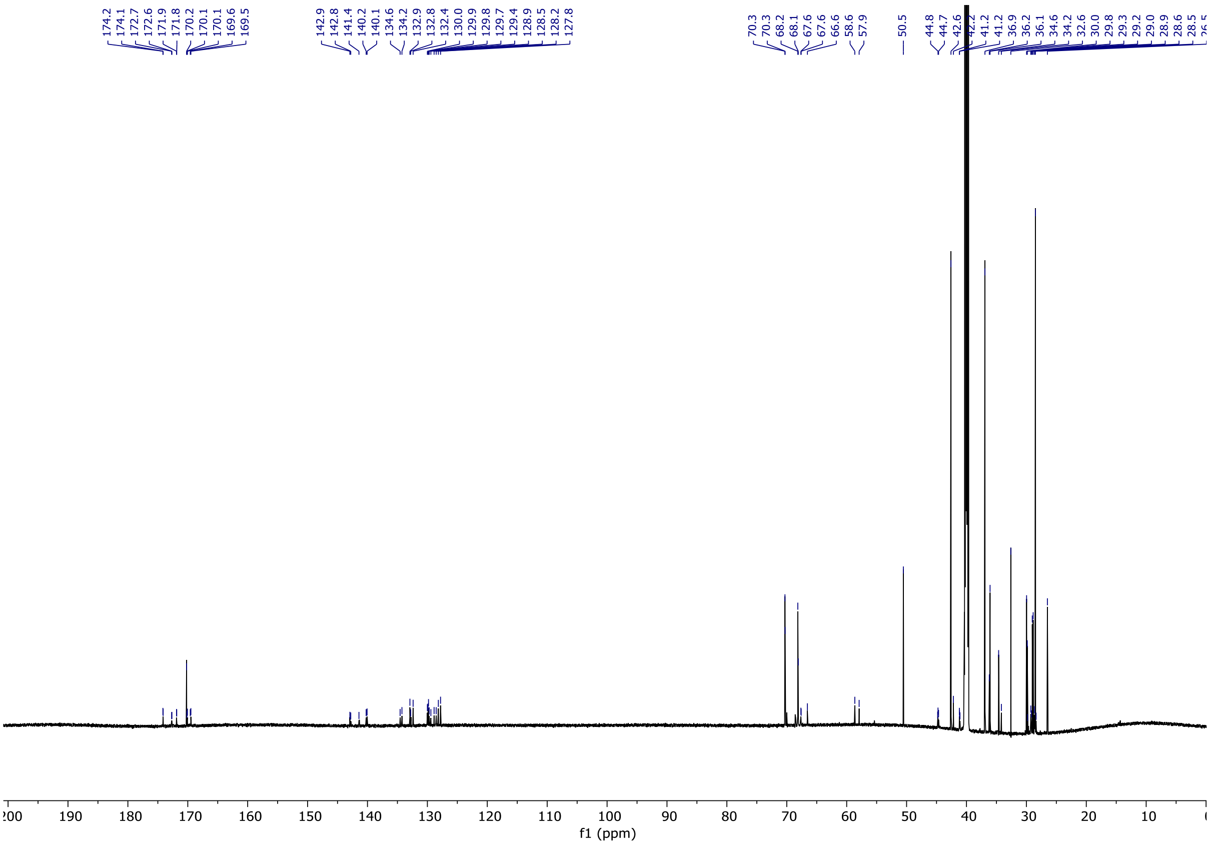

N-(1-((3r,5r,7r)-adamantan-1-yl)-2-oxo-7,10,13-trioxa-3-azahexadecan-16-yl)-1-(4-((4-chloro-2-((2-chlorophenyl)(hydroxy)methyl)phenyl)(neopentyl)amino)-4-oxobutanoyl)piperidine-4-carboxamide **11**

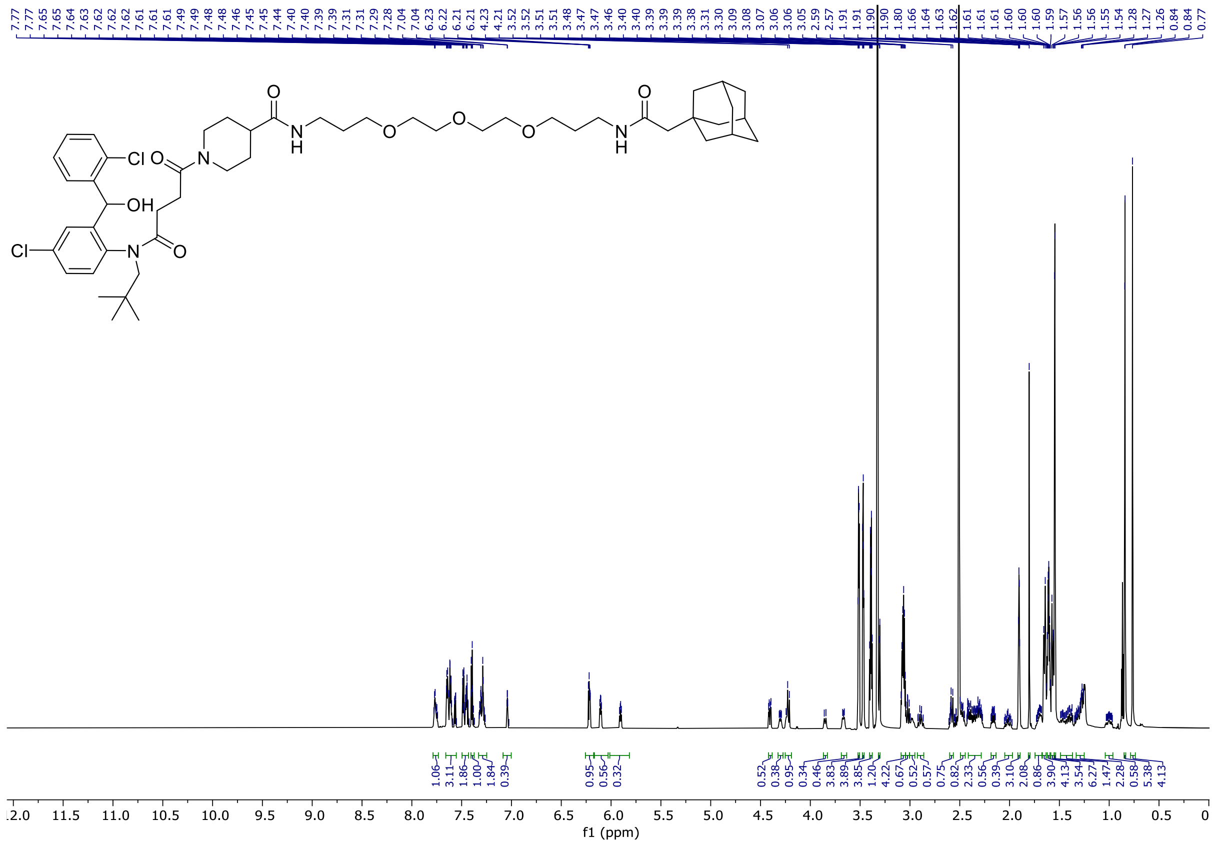


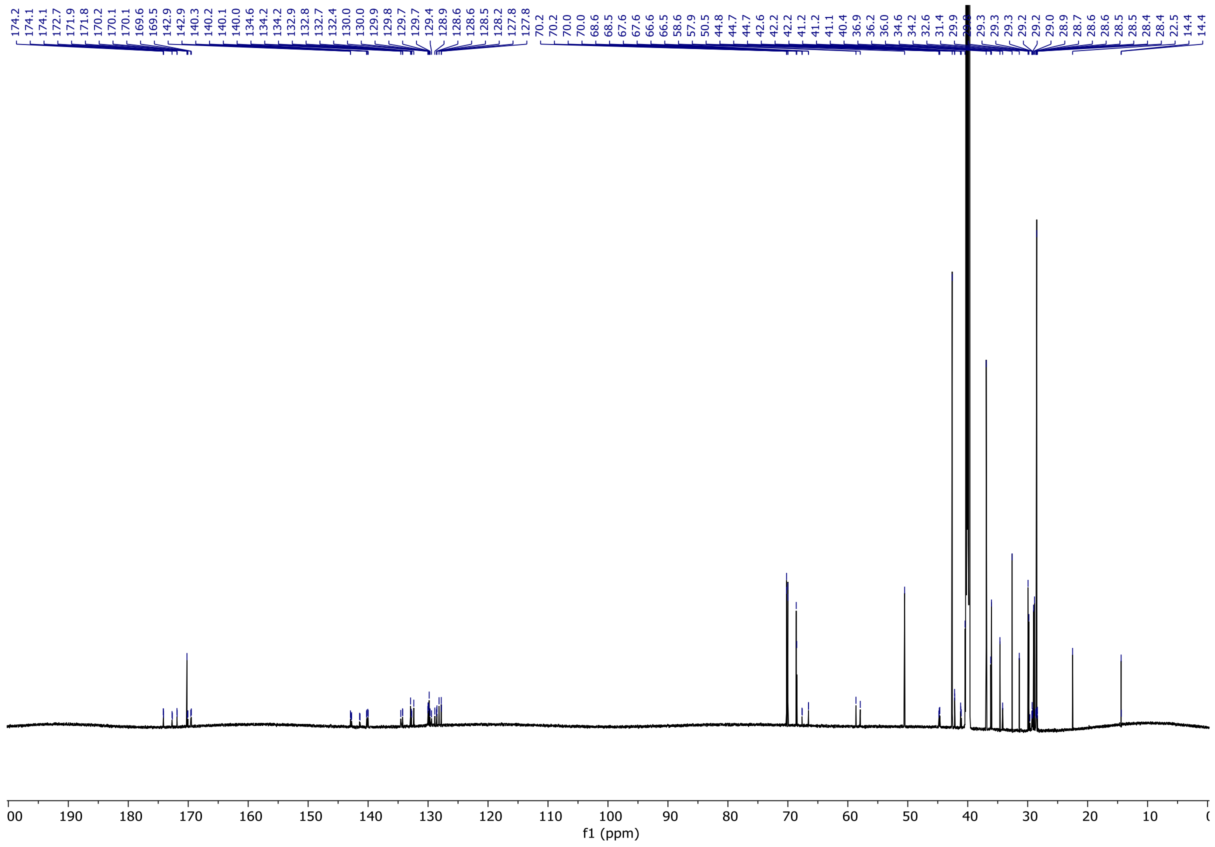

N-(6-(2-((1s,3s)-adamantan-1-yl)acetamido)hexyl)-1-(4-((4-chloro-2-((2-chlorophenyl)(hydroxy)methyl)phenyl)(neopentyl)amino)-4-oxobutanoyl)piperidine-4-carboxamide **12**

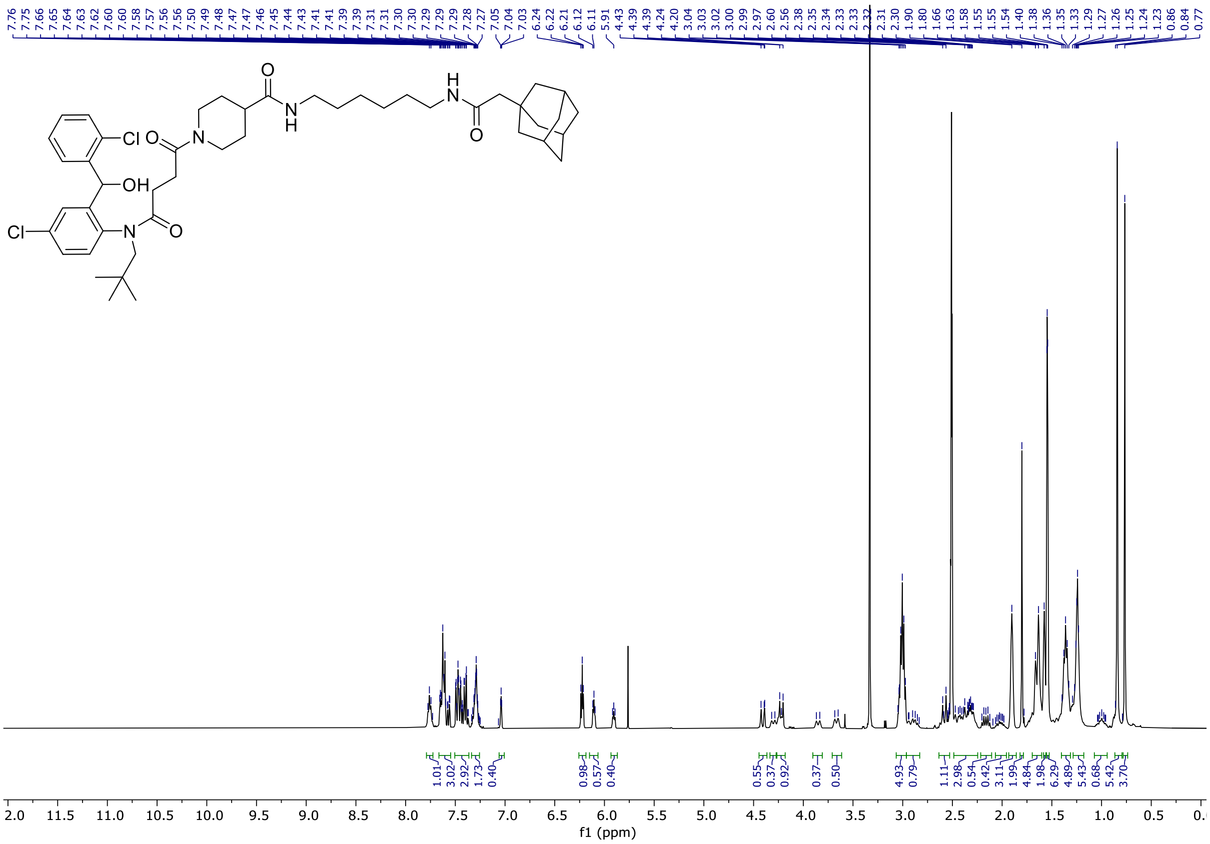

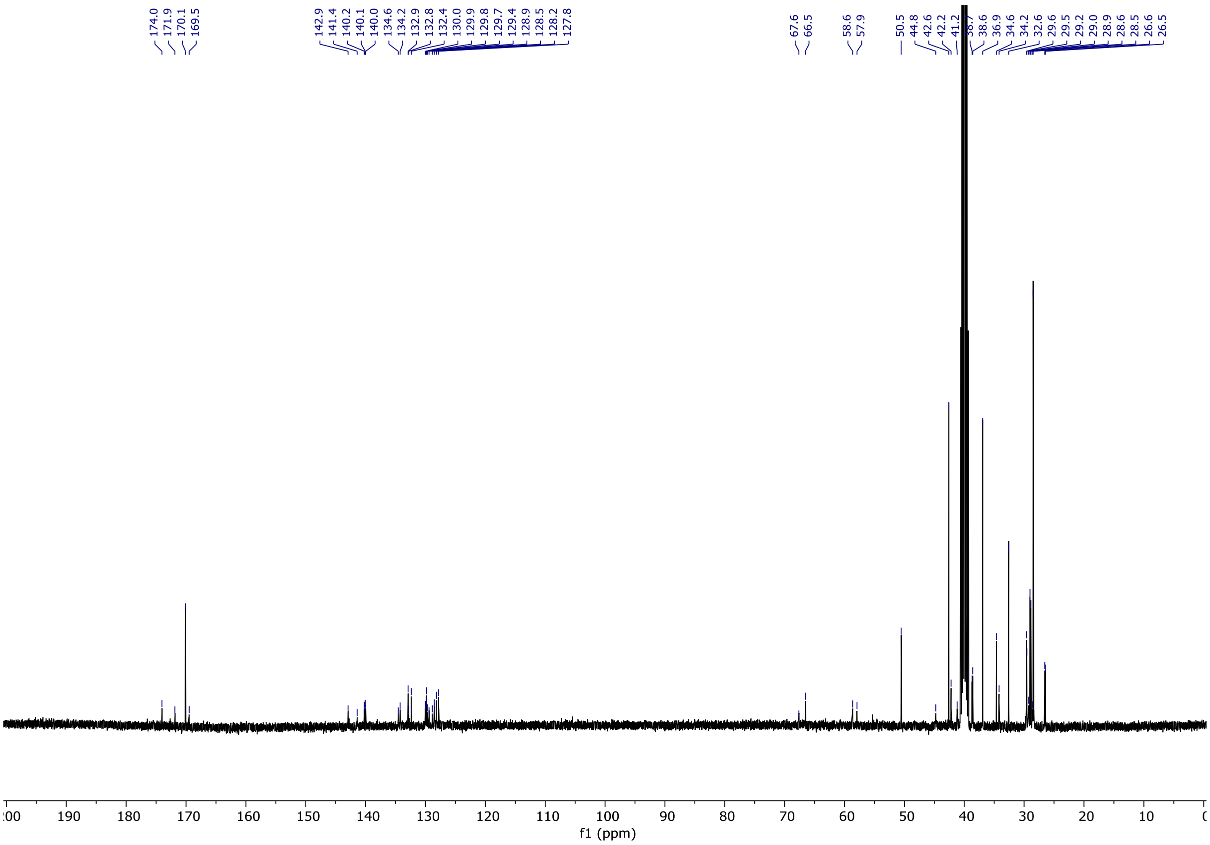

5-((2-(2-(2-(1-(4-((4-chloro-2-((2-chlorophenyl)(hydroxy)methyl)phenyl)(neopentyl)amino)-4-oxobutanoyl)piperidine-4-carboxamido)ethoxy)ethoxy)ethyl)carbamoyl)-2-(6-(dimethylamino)-3-(dimethyliminio)-3H-xanthen-9-yl)benzoate **26**

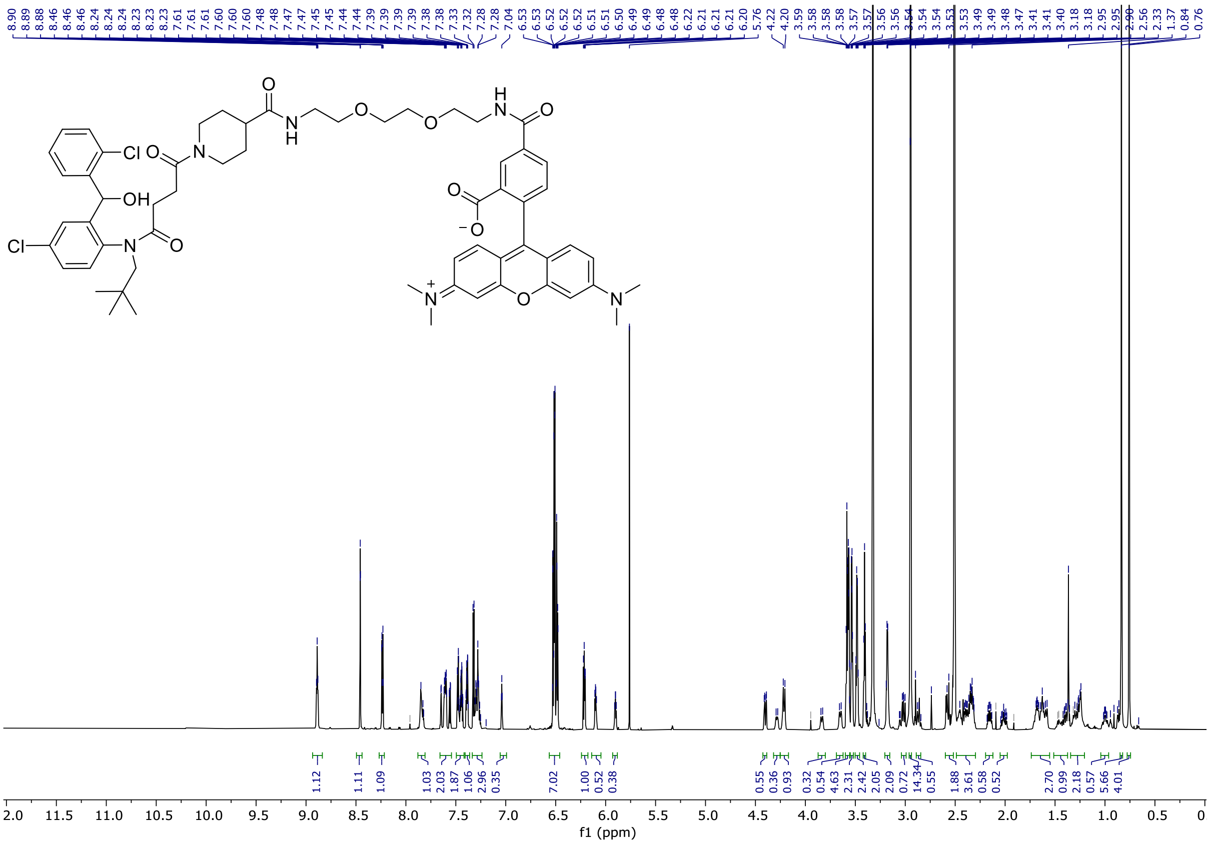

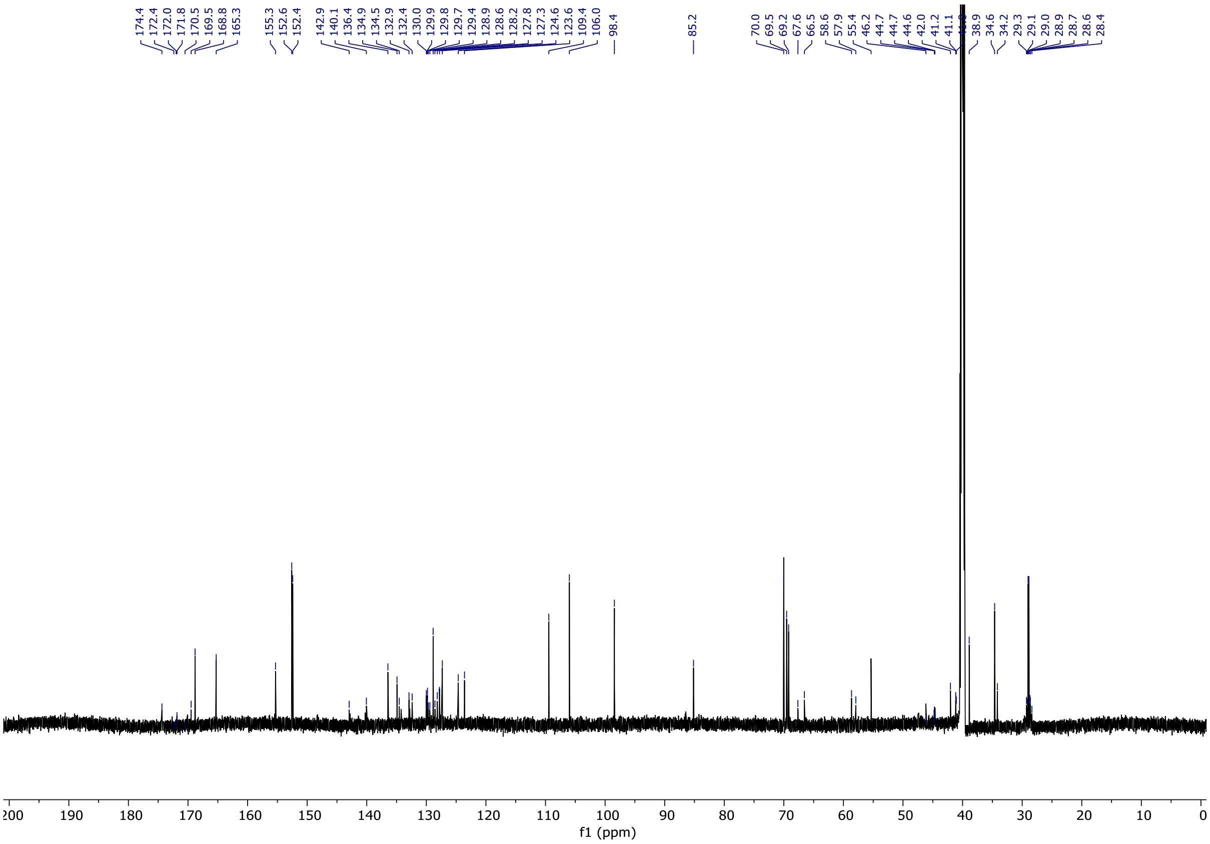

3-(3,4-dimethoxyphenyl)-*N*-(6-(3-((2-(2-(2-((2-(2,6-dioxopiperidin-3-yl)-1,3-dioxoisoindolin-4-yl)amino)ethoxy)ethoxy)ethyl)amino)-3-oxopropyl)benzo[d]thiazol-2-yl)propanamide **13**

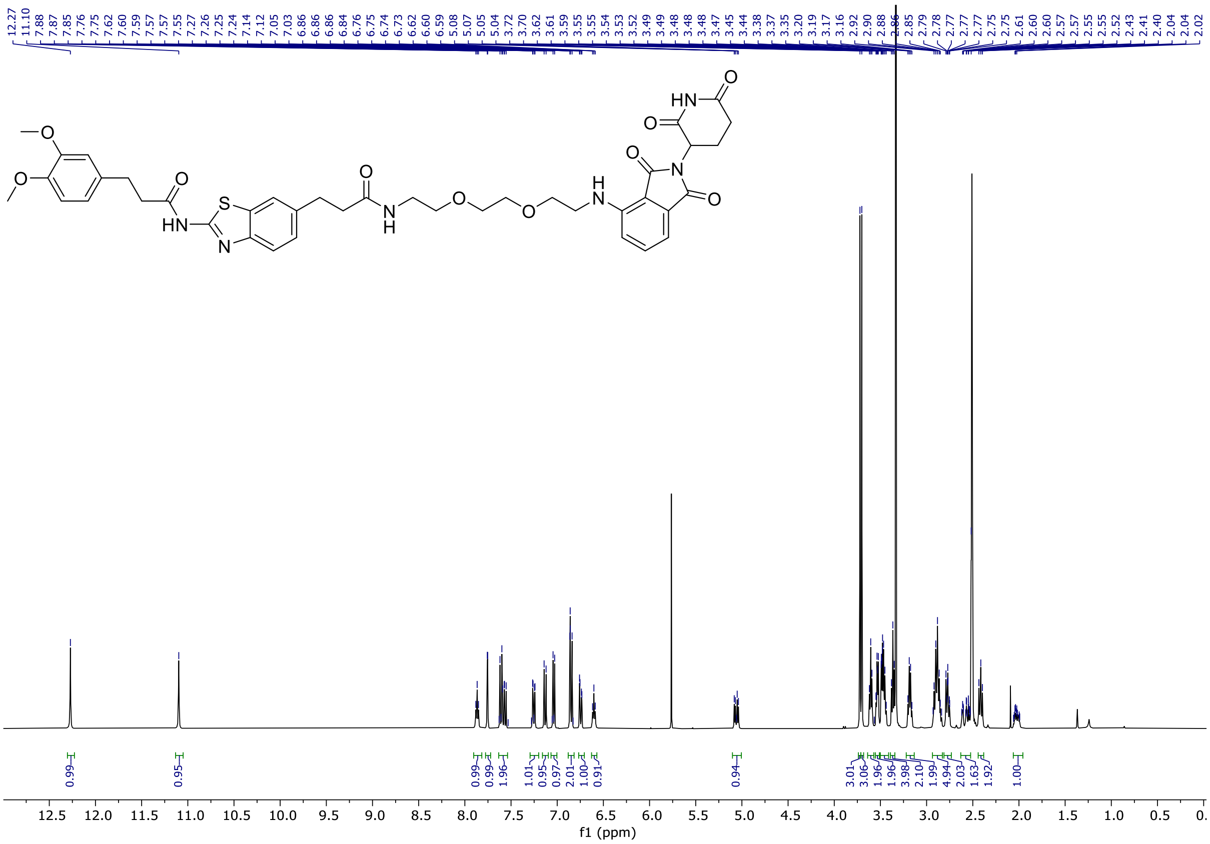


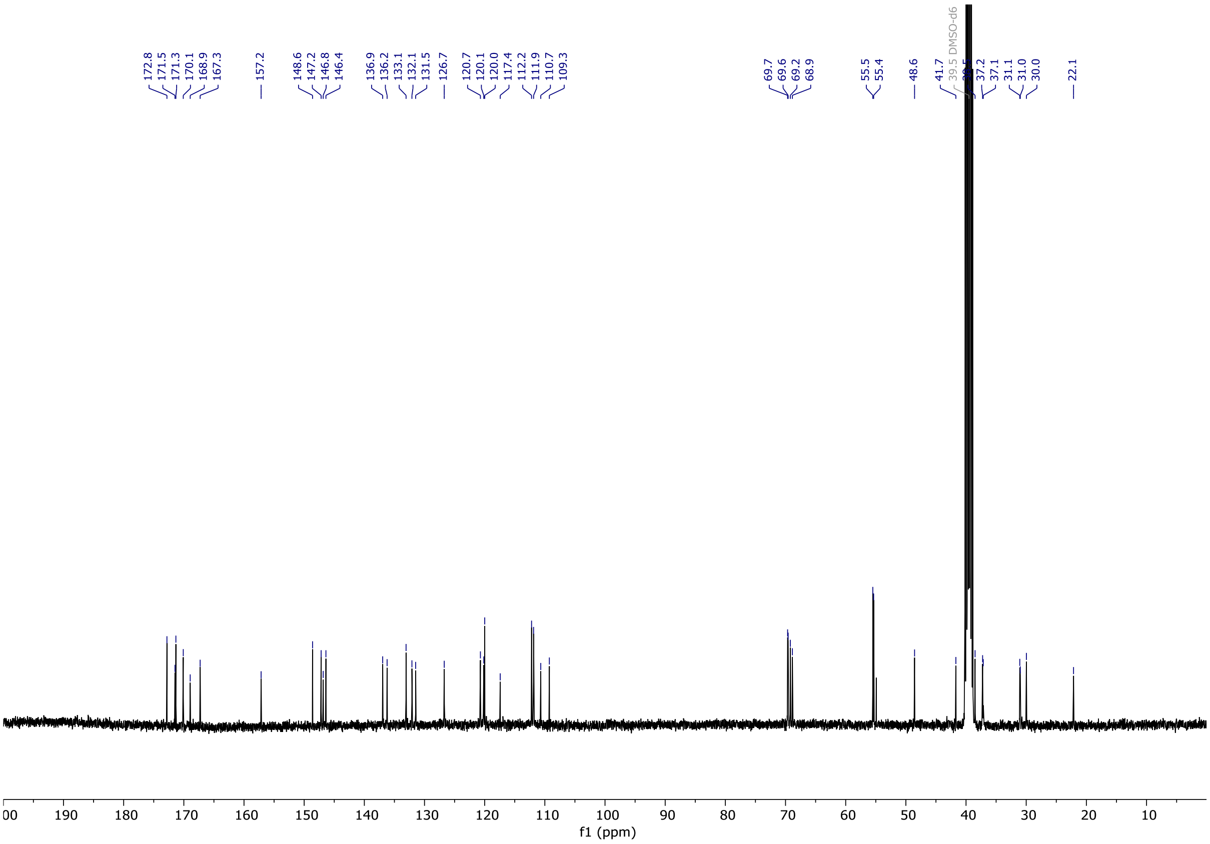

3-(3,4-dimethoxyphenyl)-*N*-(6-(3-((3-(4-(3-((2-(2,6-dioxopiperidin-3-yl)-1,3-dioxoisoindolin-4-yl)amino)propoxy)butoxy)propyl)amino)-3-oxopropyl)benzo[d]thiazol-2-yl)propanamide **14**

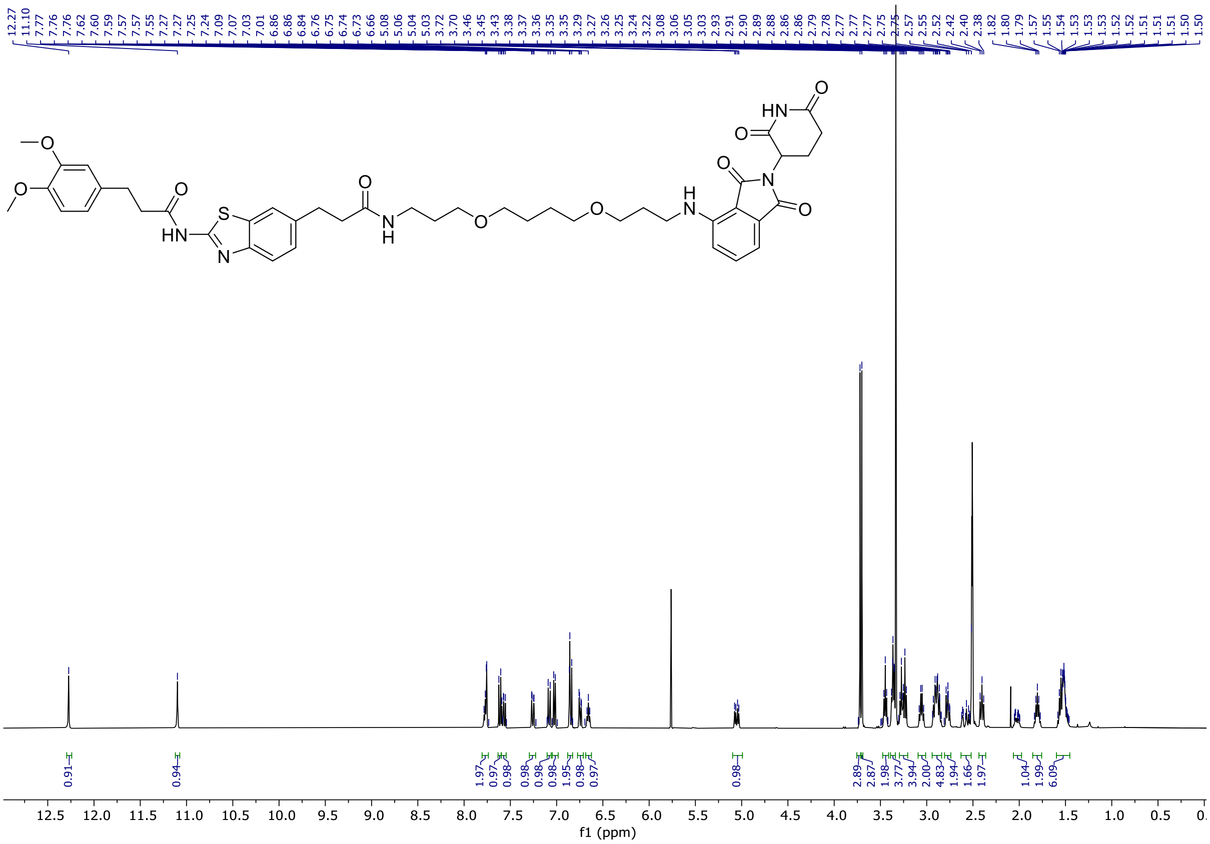


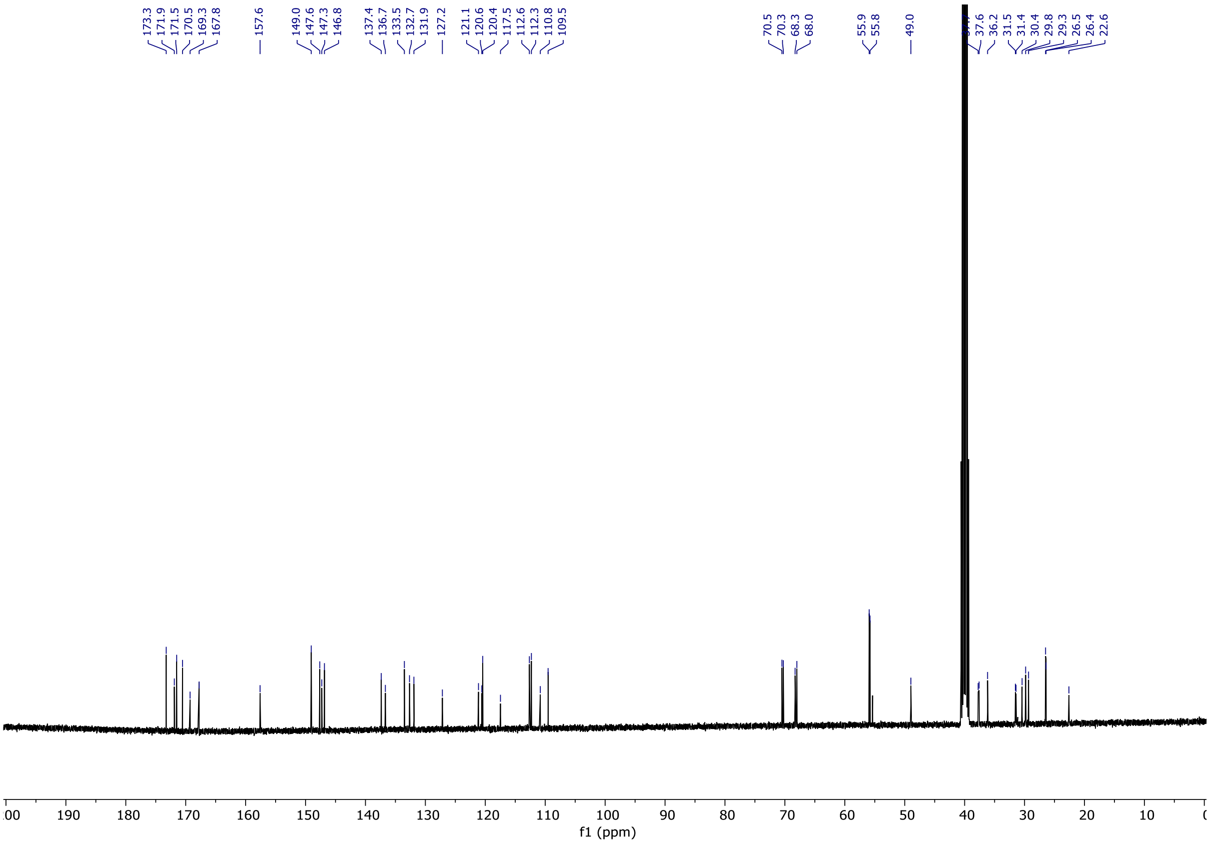

3-(3,4-dimethoxyphenyl)-*N*-(6-(1-((2-(2,6-dioxopiperidin-3-yl)-1,3-dioxoisoindolin-4-yl)amino)-15-oxo-4,7,10-trioxa-14-azaheptadecan-17-yl)benzo[d]thiazol-2-yl)propanamide **15**

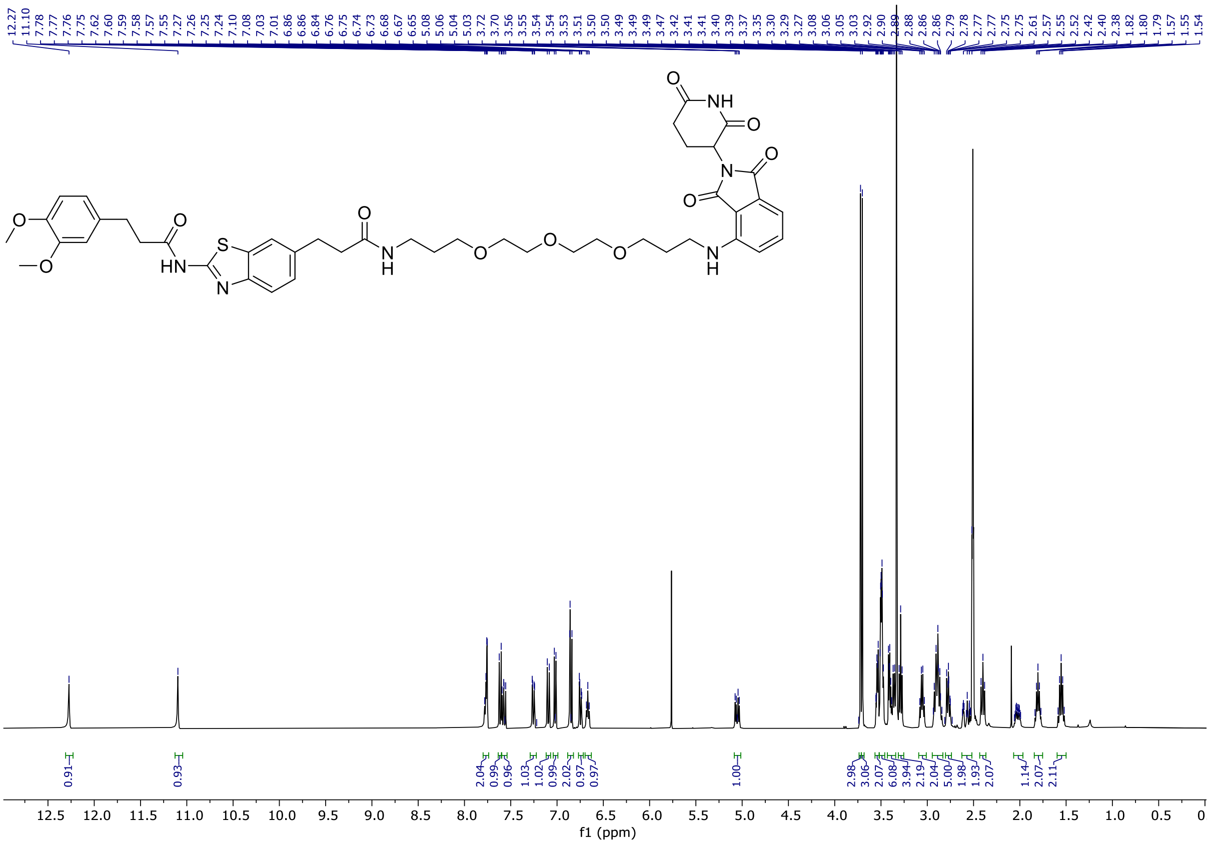


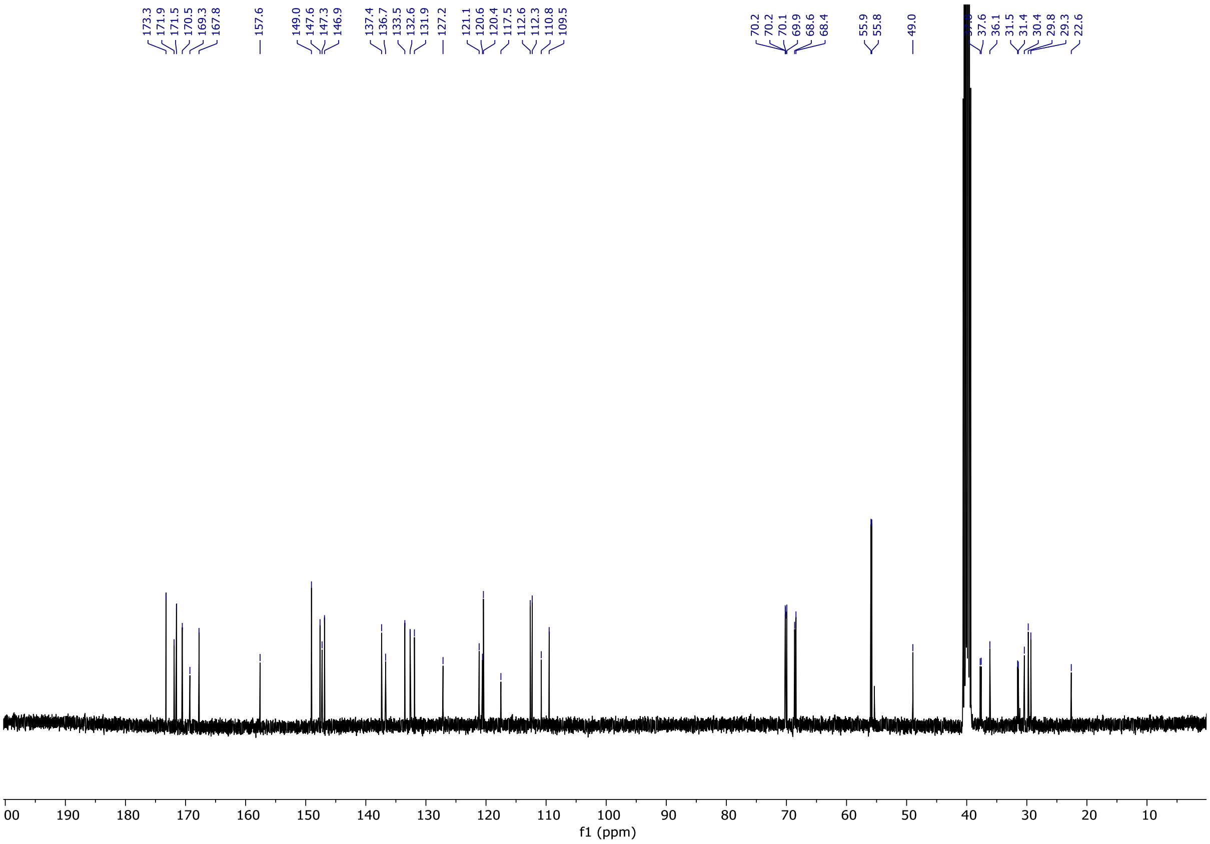

3-(3,4-dimethoxyphenyl)-*N*-(6-(3-((6-((2-(2,6-dioxopiperidin-3-yl)-1,3-dioxoisoindolin-4-yl)amino)hexyl)amino)-3-oxopropyl)benzo[d]thiazol-2-yl)propanamide **16**

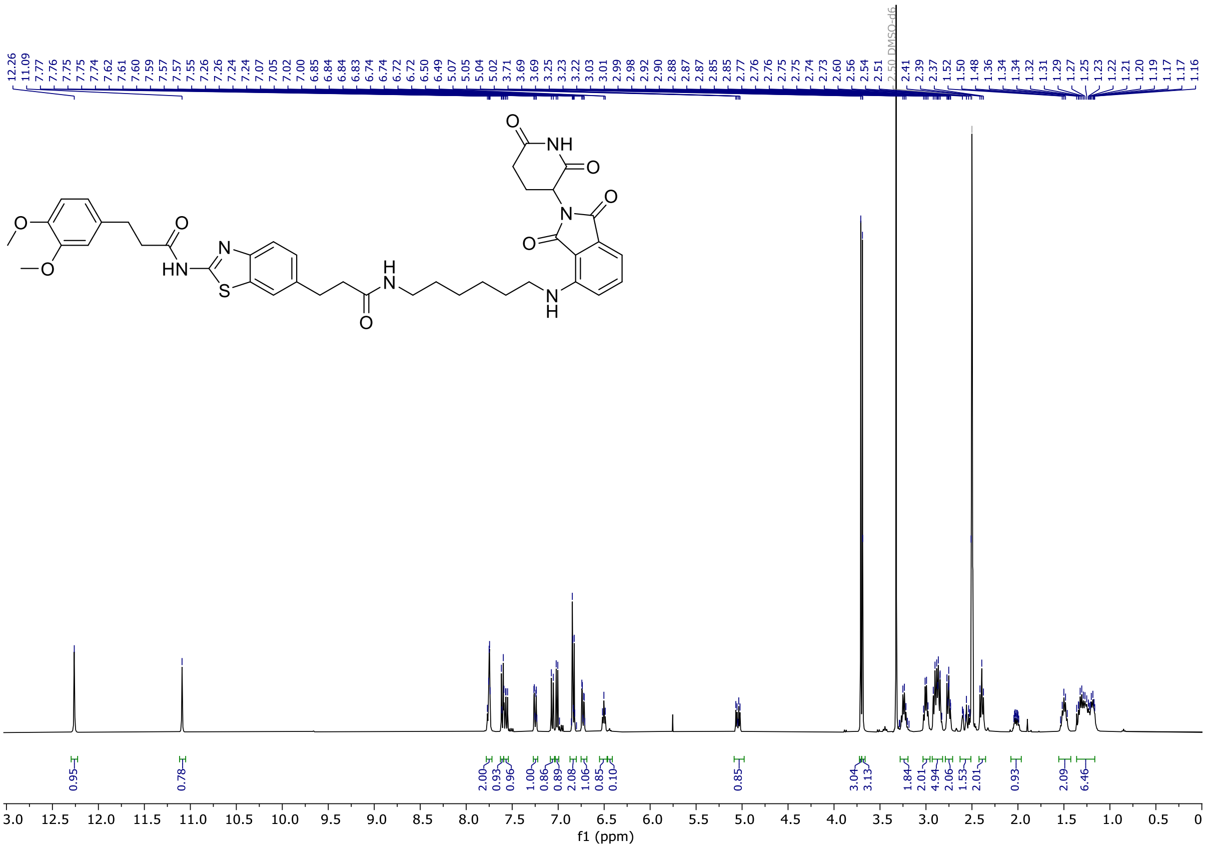


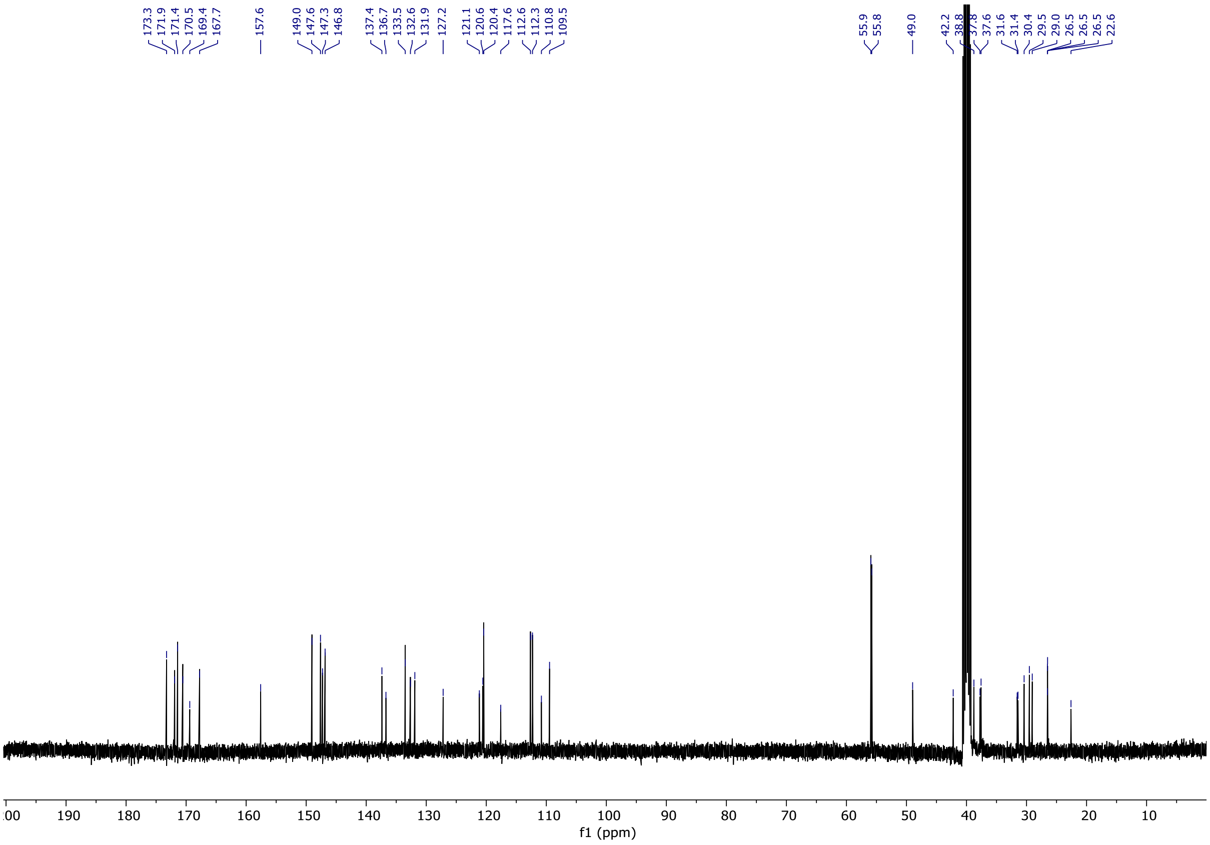

3-(3,4-dimethoxyphenyl)-*N*-(6-(3-((10-((2-(2,6-dioxopiperidin-3-yl)-1,3-dioxoisoindolin-4-yl)amino)decyl)amino)-3-oxopropyl)benzo[d]thiazol-2-yl)propanamide **17**

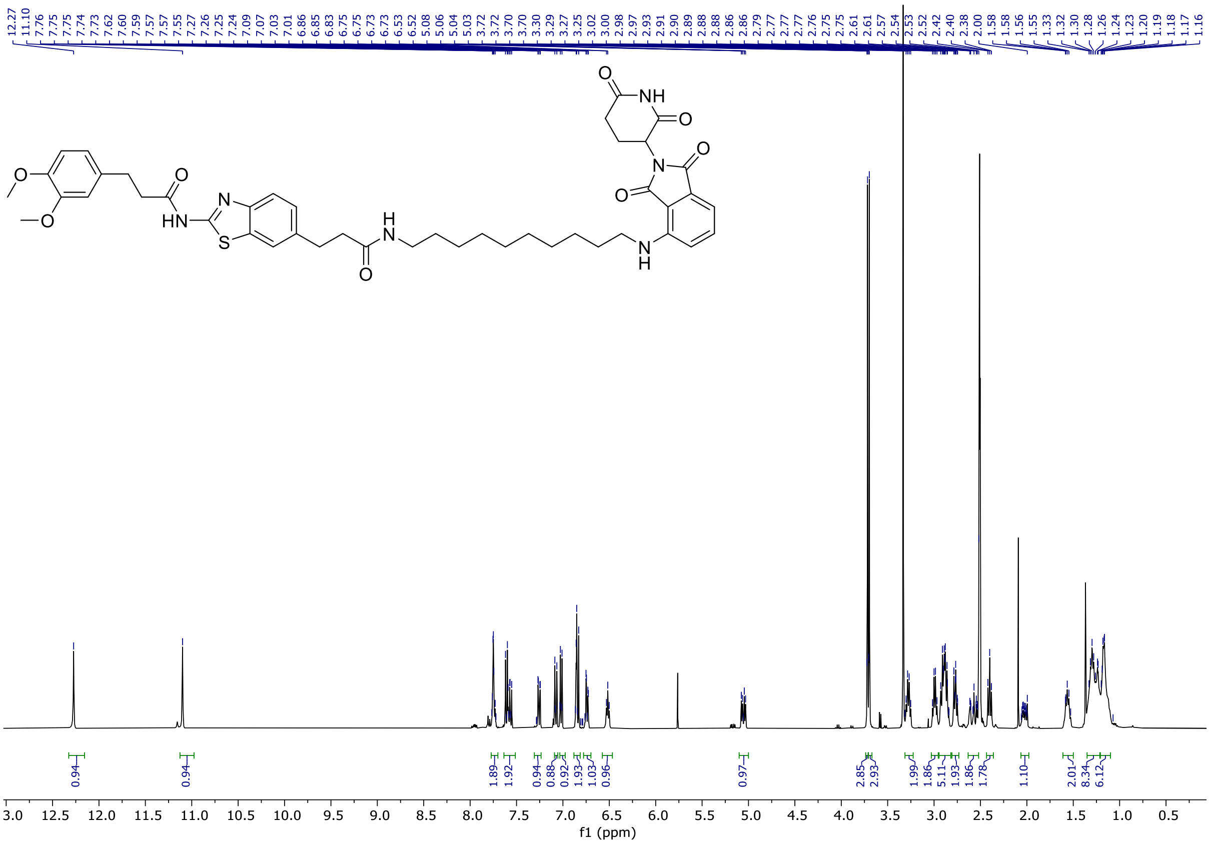


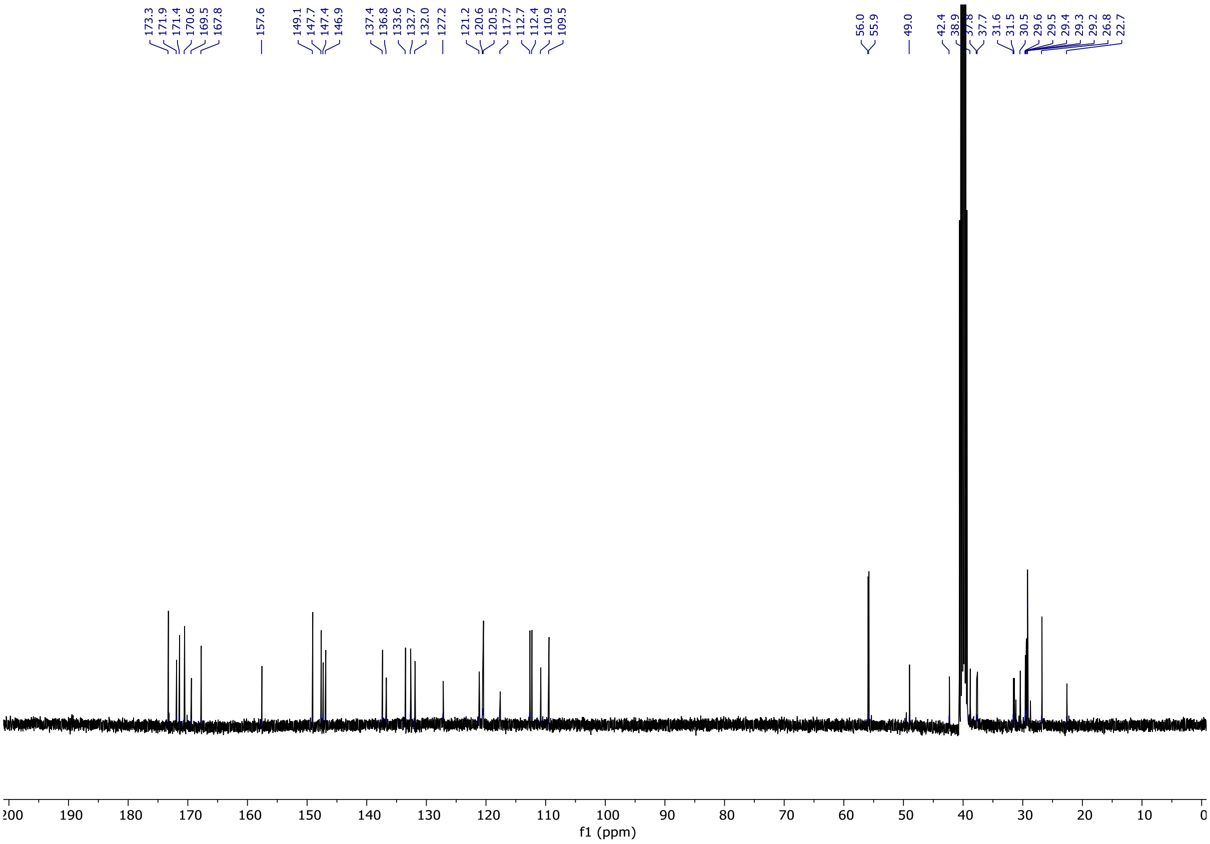

(2*S*,4*R*)-1-((*S*)-2-(*tert*-butyl)-15-(2-(3-(3,4-dimethoxyphenyl)propanamido)benzo[*d*]thiazol-6-yl)-4,13-dioxo-6,9-dioxa-3,12-diazapentadecanoyl)-4-hydroxy-*N*-(4-(4-methylthiazol-5-yl)benzyl) pyrrolidine-2-carboxamide **18**

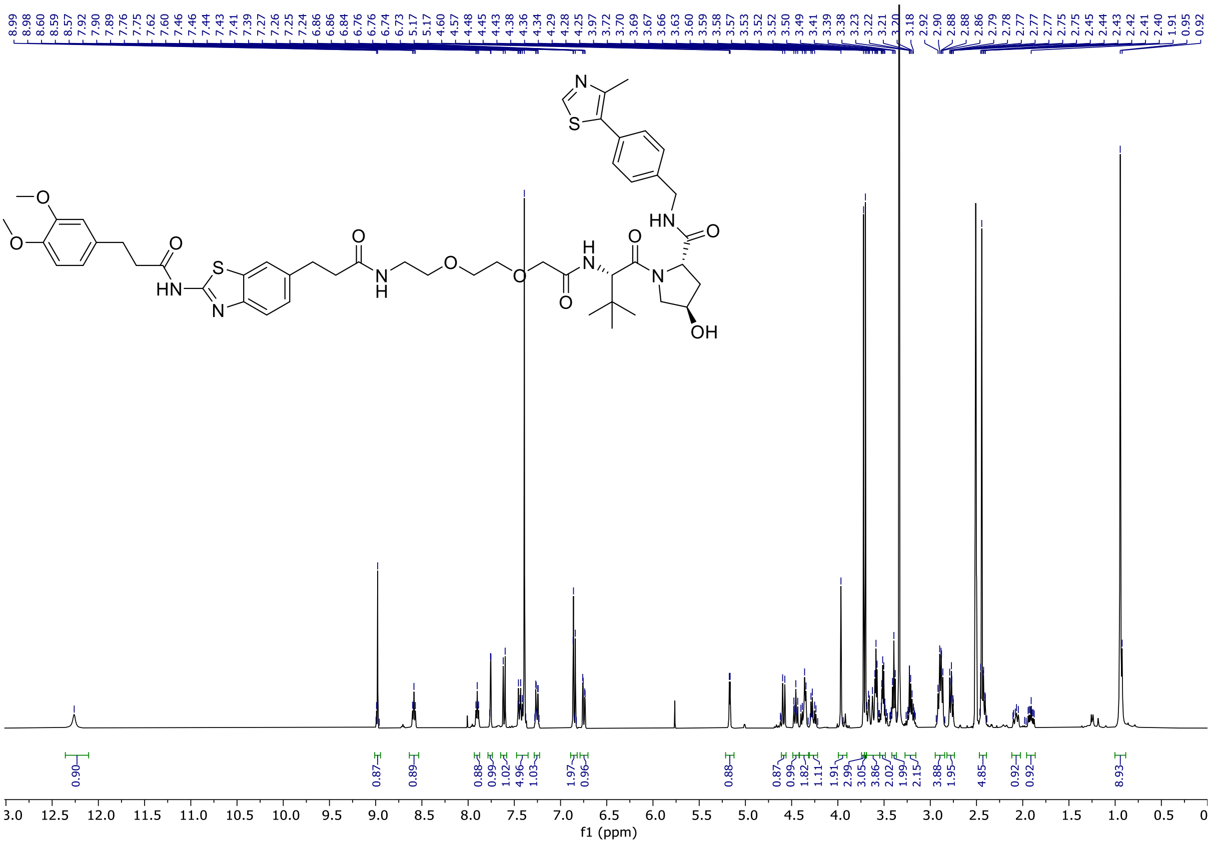


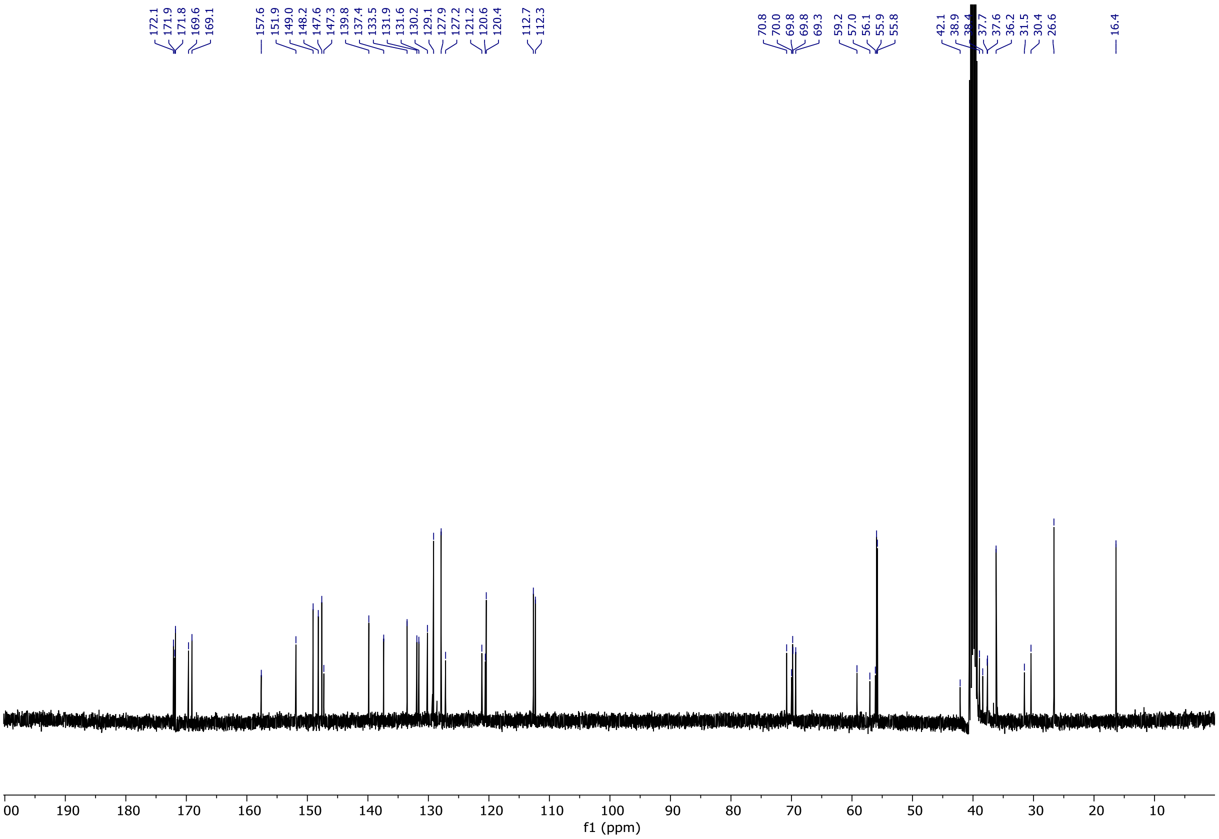

(2*S*,4*R*)-1-((*S*)-2-(*tert*-butyl)-22-(2-(3-(3,4-dimethoxyphenyl)propanamido)benzo[*d*]thiazol-6-yl)-4,20-dioxo-7,10,13,16-tetraoxa-3,19-diazadocosanoyl)-4-hydroxy-*N*-(4-(4-methylthiazol-5-yl)benzyl)pyrrolidine-2-carboxamide **19**

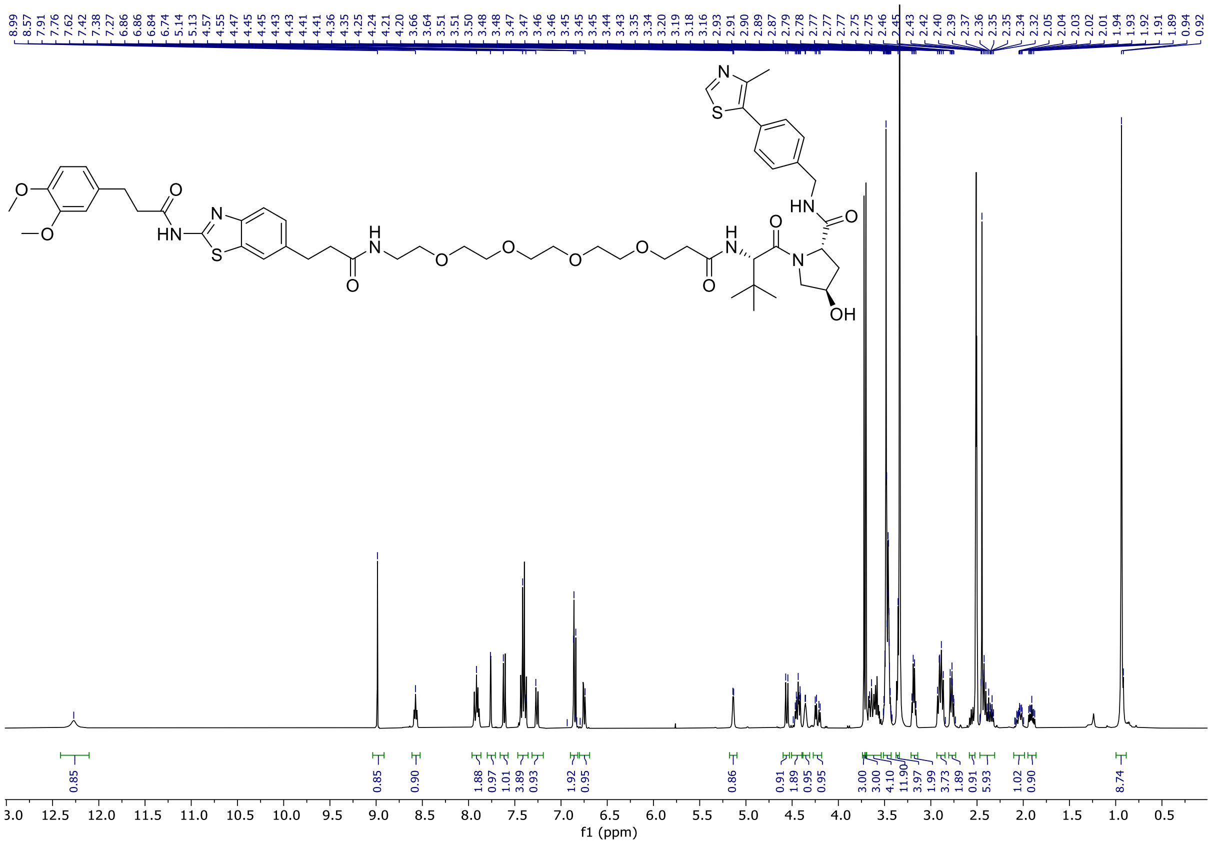


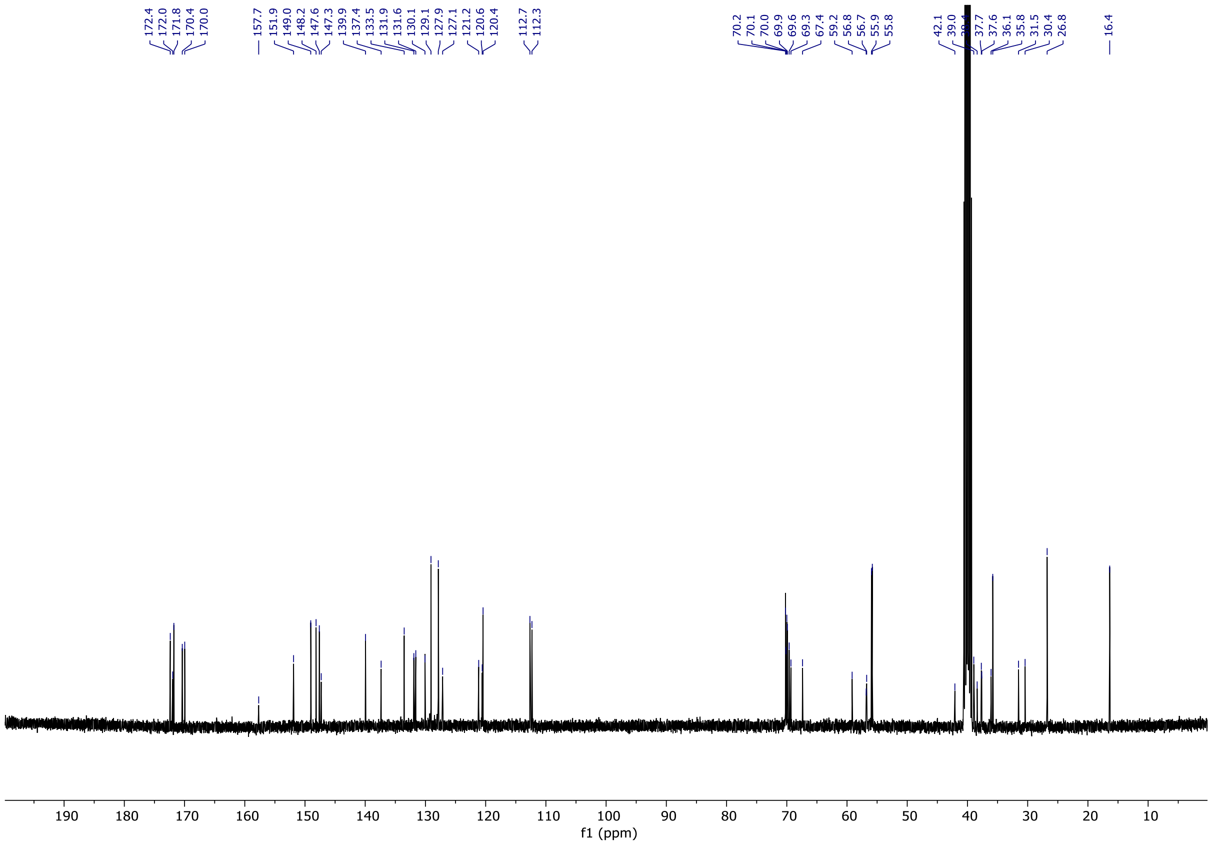

(2*S*,4*R*)-1-((*S*)-2-(6-(3-(2-(3-(3,4-dimethoxyphenyl)propanamido)benzo[*d*]thiazol-6-yl)propanamido)hexanamido)-3,3-dimethylbutanoyl)-4-hydroxy-*N*-(4-(4-methylthiazol-5-yl)benzyl)pyrrolidine-2-carboxamide **20**

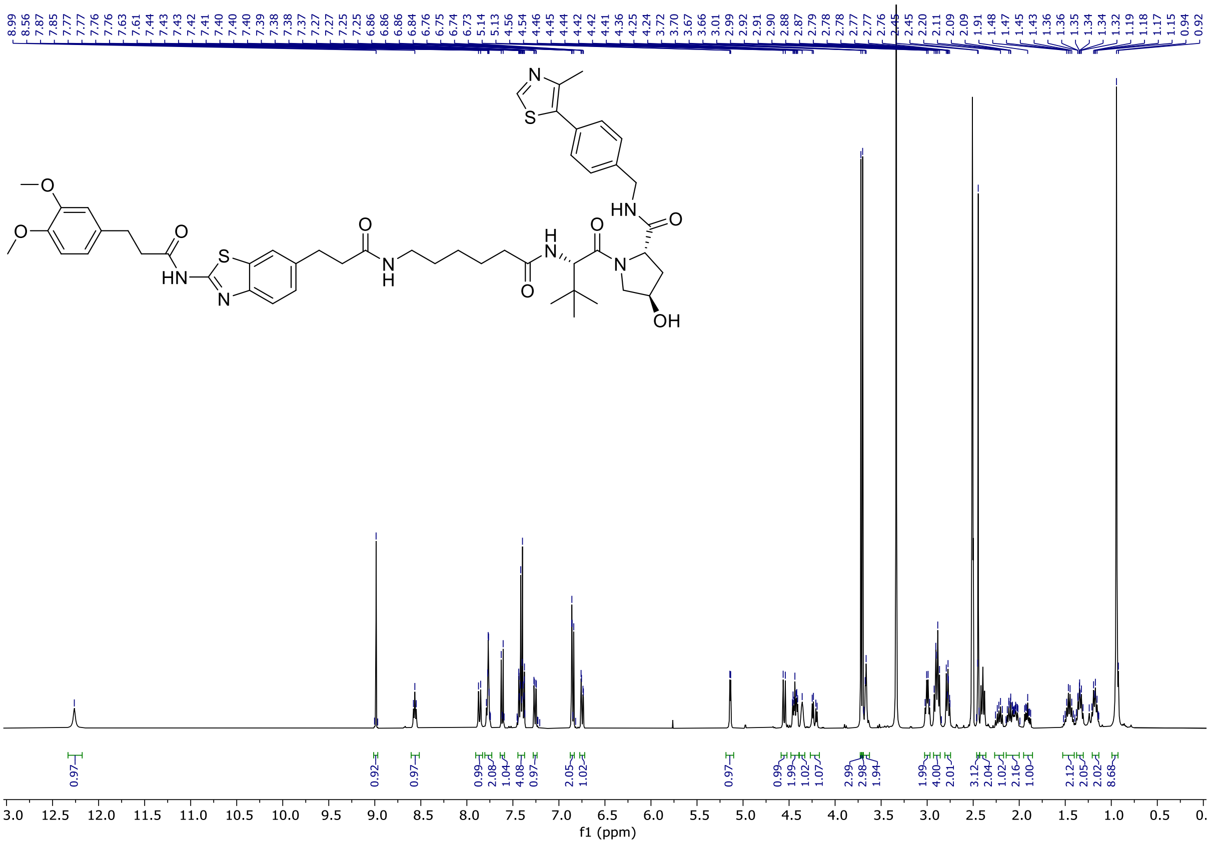


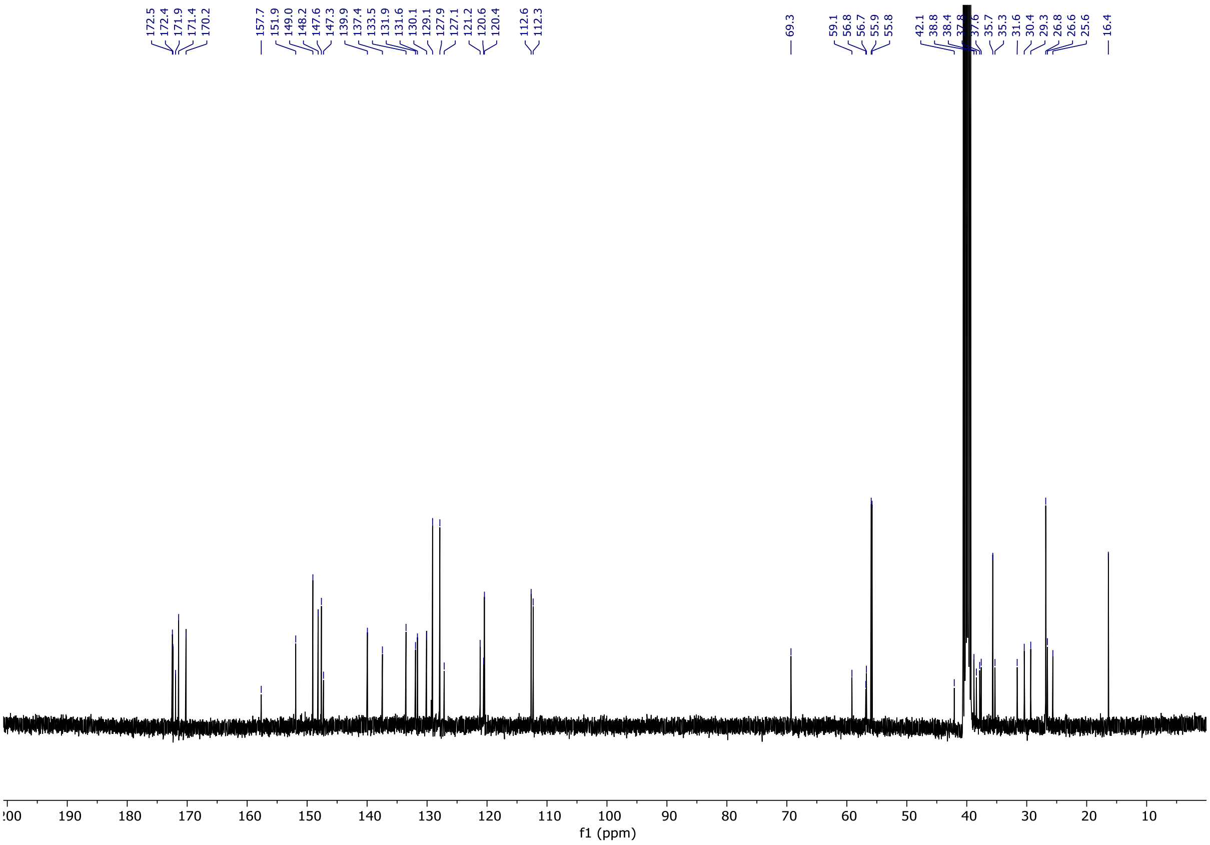

(2*S*,4*R*)-1-((*S*)-2-(10-(3-(2-(3-(3,4-dimethoxyphenyl)propanamido)benzo[*d*]thiazol-6-yl)propanamido)decanamido)-3,3-dimethylbutanoyl)-4-hydroxy-*N*-(4-(4-methylthiazol-5-yl)benzyl)pyrrolidine-2-carboxamide **21**

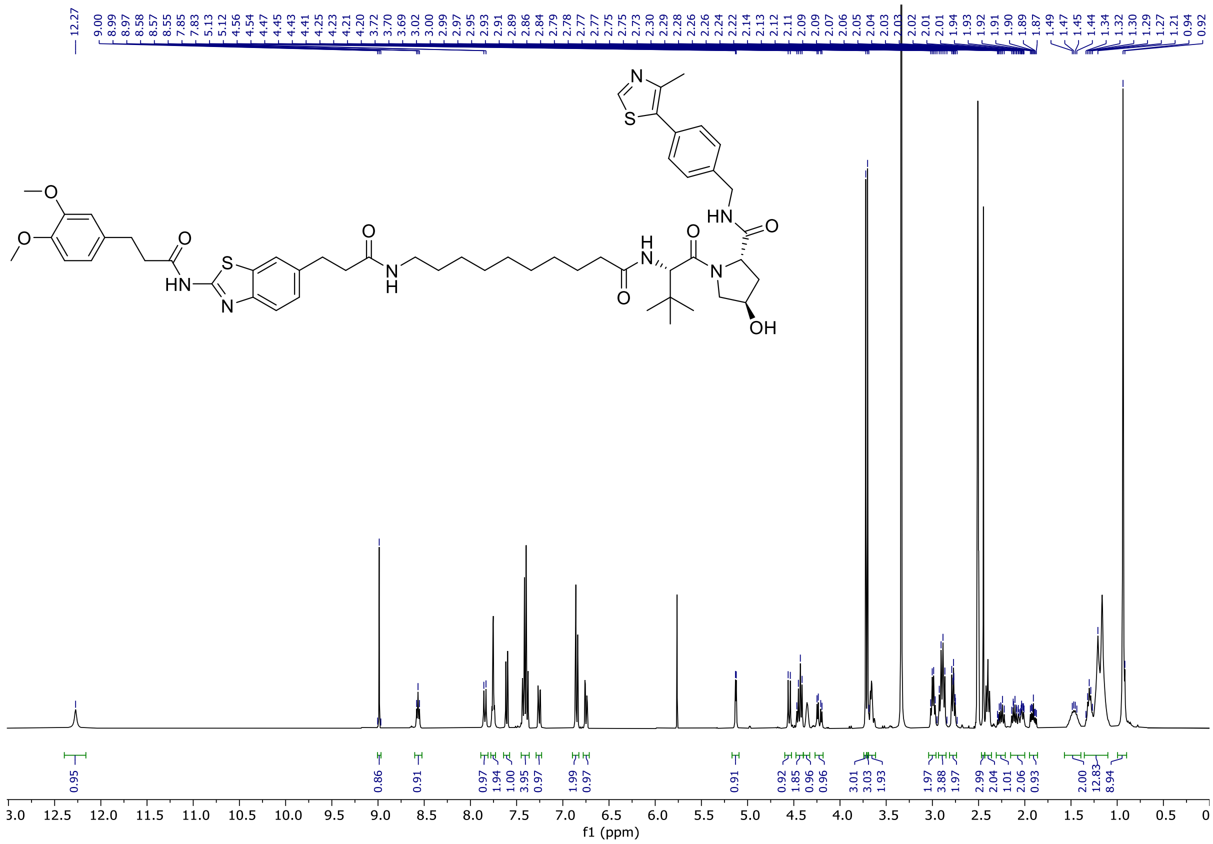


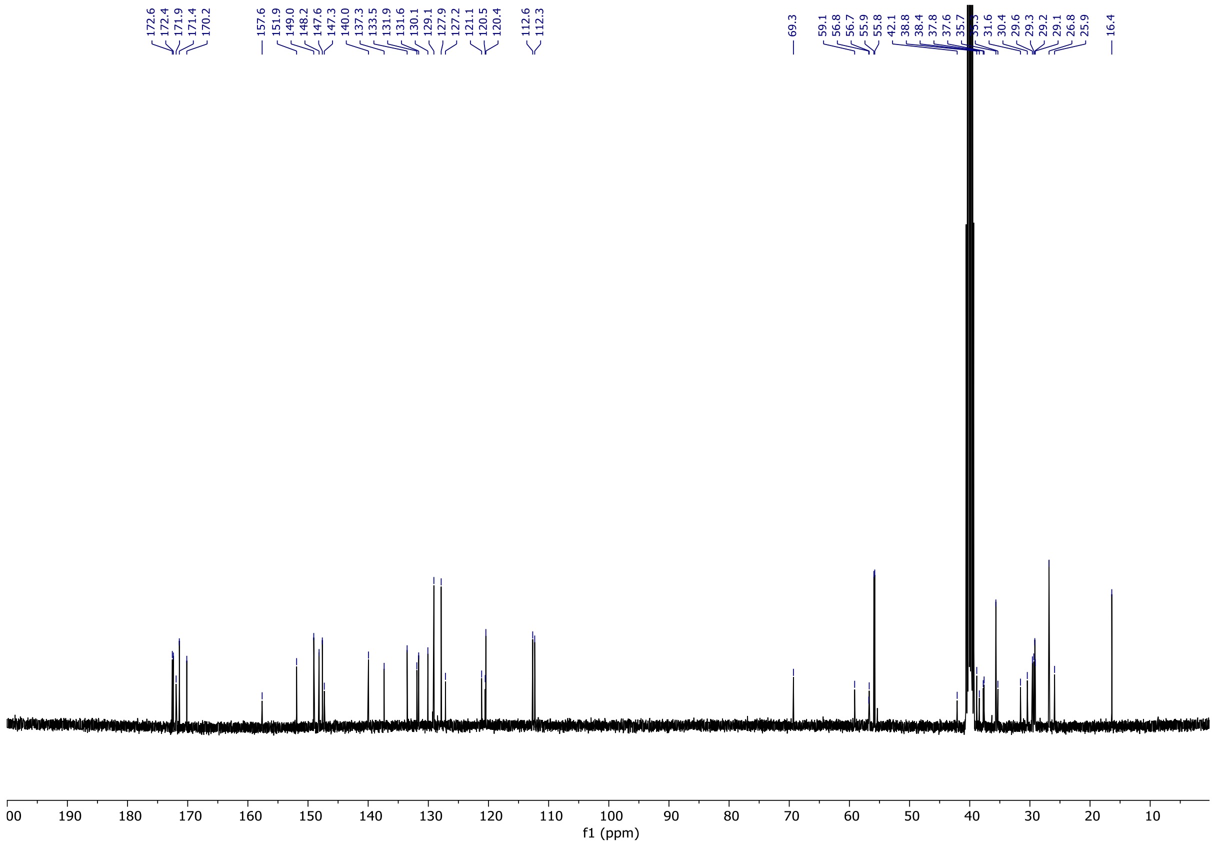

*N*-(6-(1-((3*r*,5*r*,7*r*)-adamantan-1-yl)-2,13-dioxo-6,9-dioxa-3,12-diazapentadecan-15-yl)benzo[*d*]thiazol-2-yl)-3-(3,4-dimethoxyphenyl)propanamide **22**

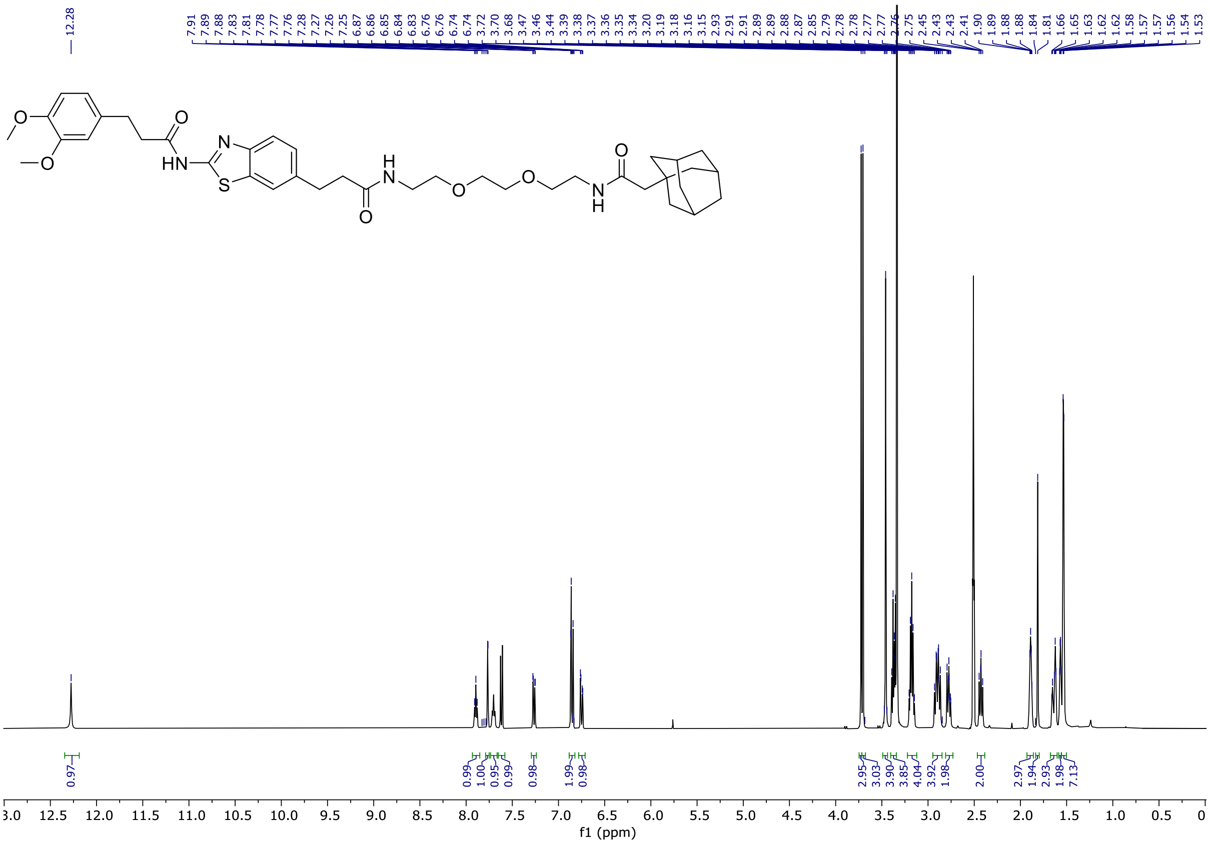


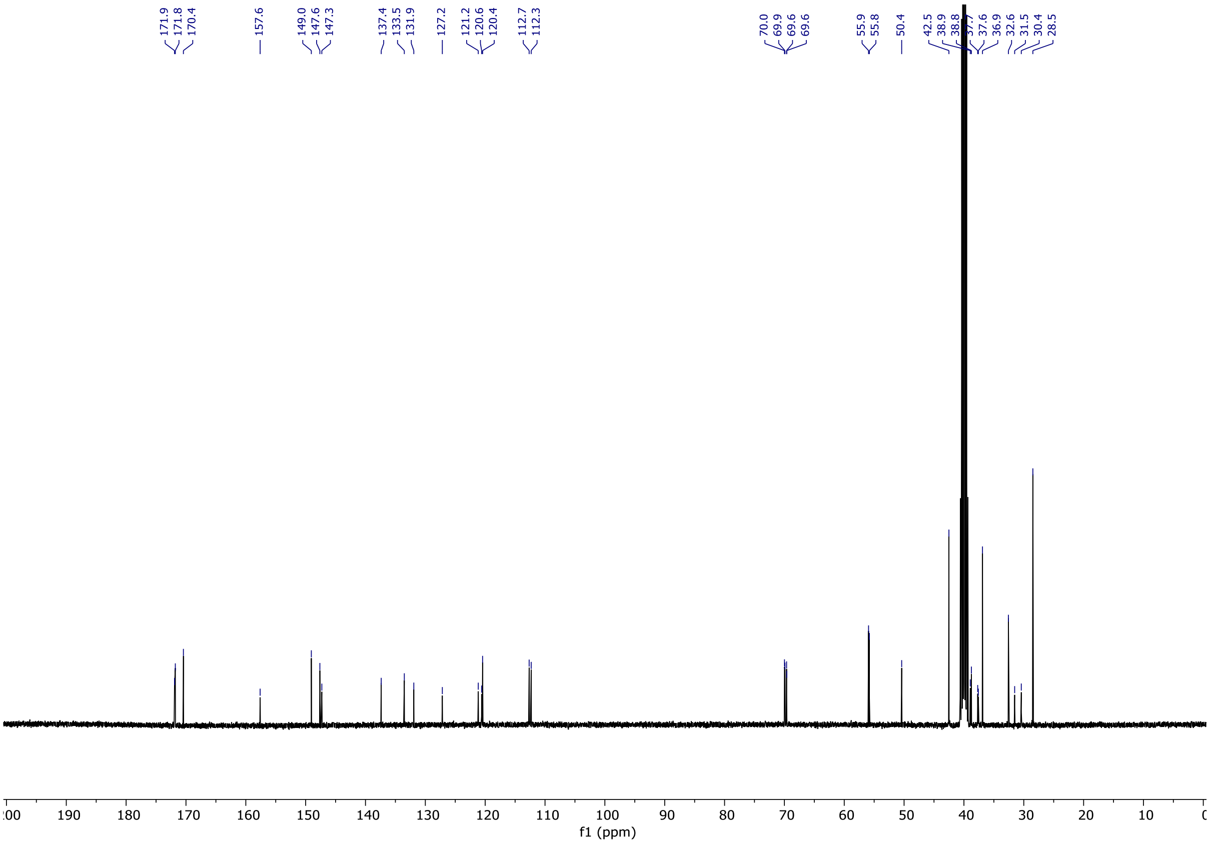

*N*-(6-(1-((3*r*,5*r*,7*r*)-adamantan-1-yl)-2,17-dioxo-7,12-dioxa-3,16-diazanonadecan-19-yl)benzo[*d*]thiazol-2-yl)-3-(3,4-dimethoxyphenyl)propanamide **23**

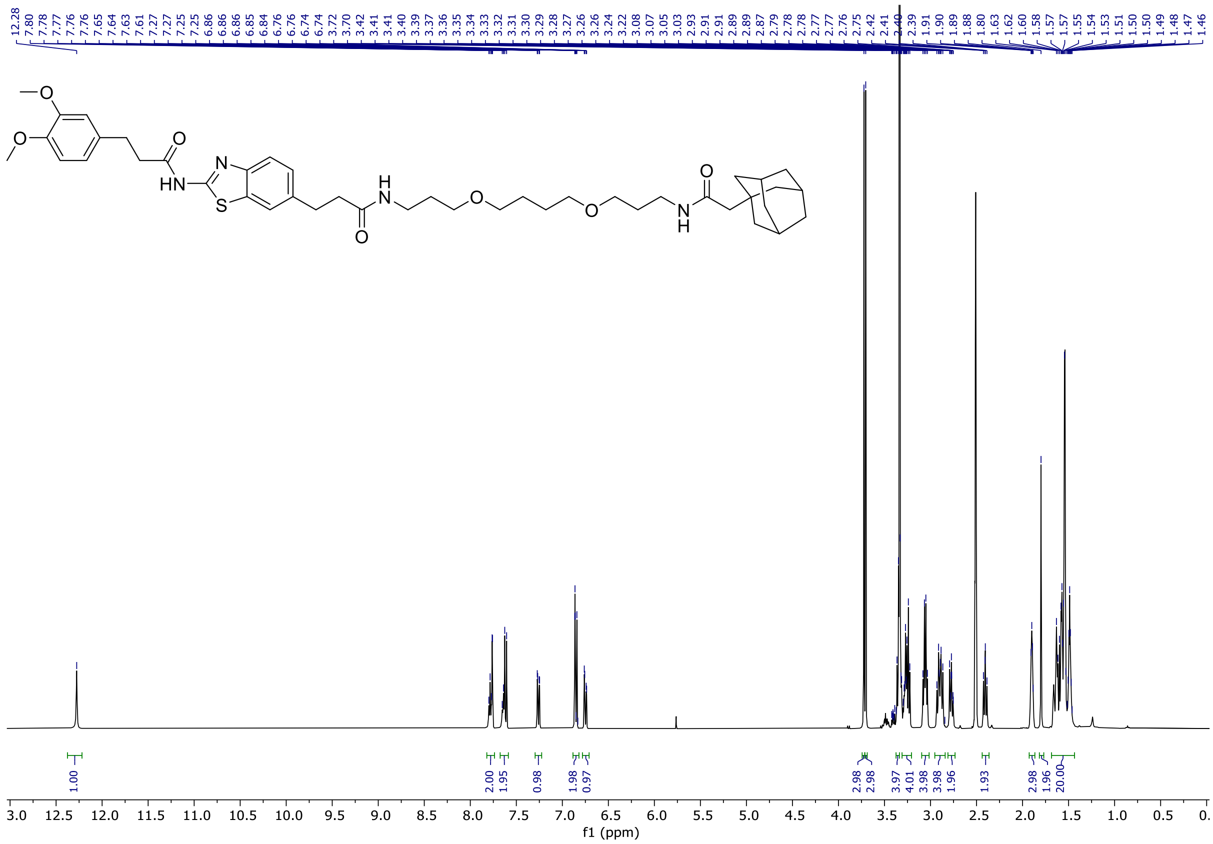


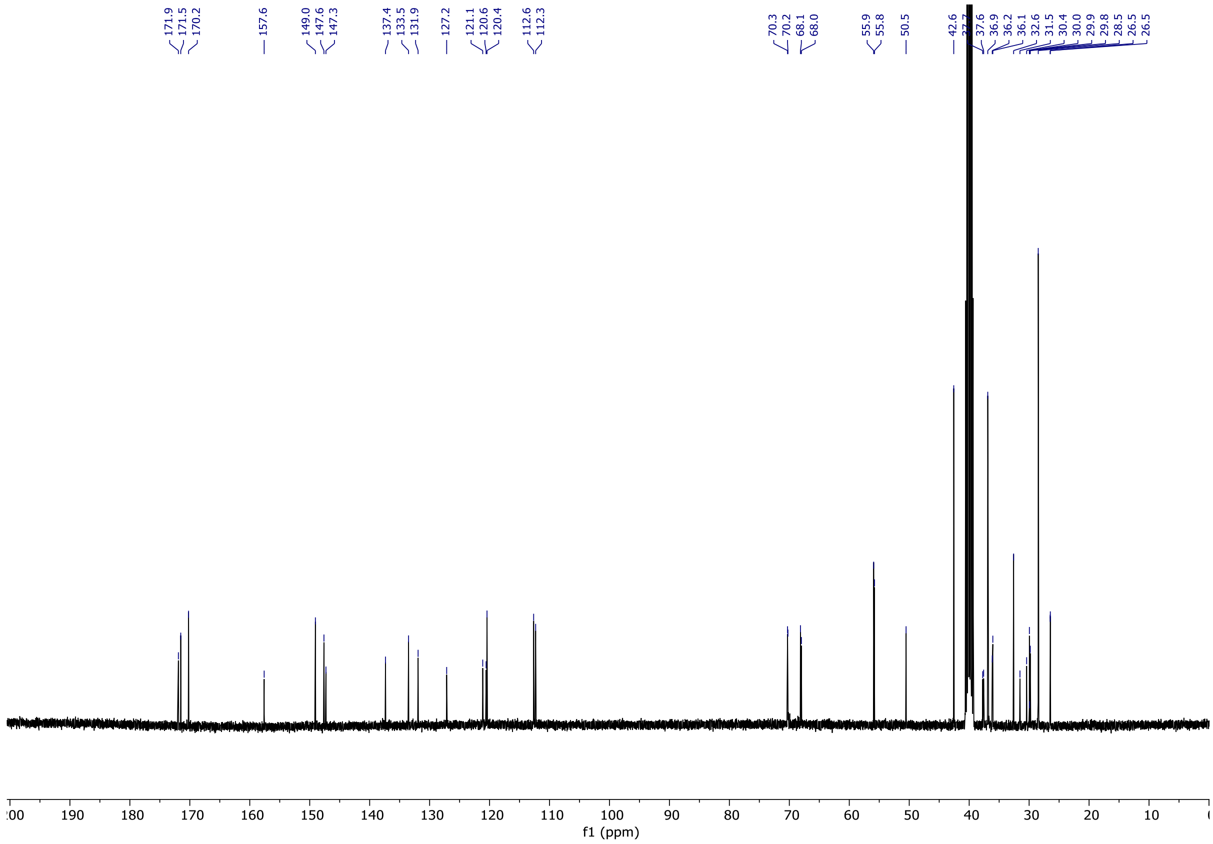

*N*-(6-(1-((3*r*,5*r*,7*r*)-adamantan-1-yl)-2,18-dioxo-7,10,13-trioxa-3,17-diazaicosan-20-yl)benzo[*d*]thiazol-2-yl)-3-(3,4-dimethoxyphenyl)propanamide **24**

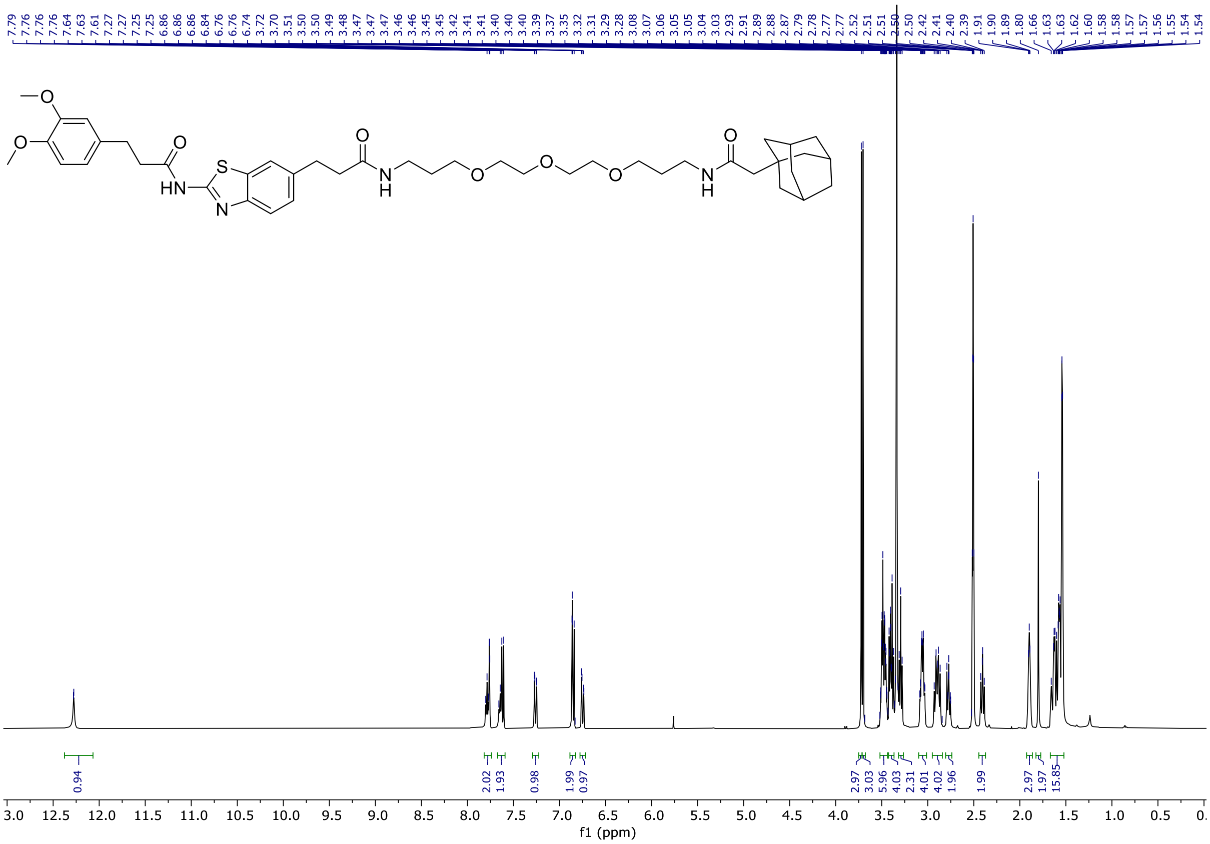


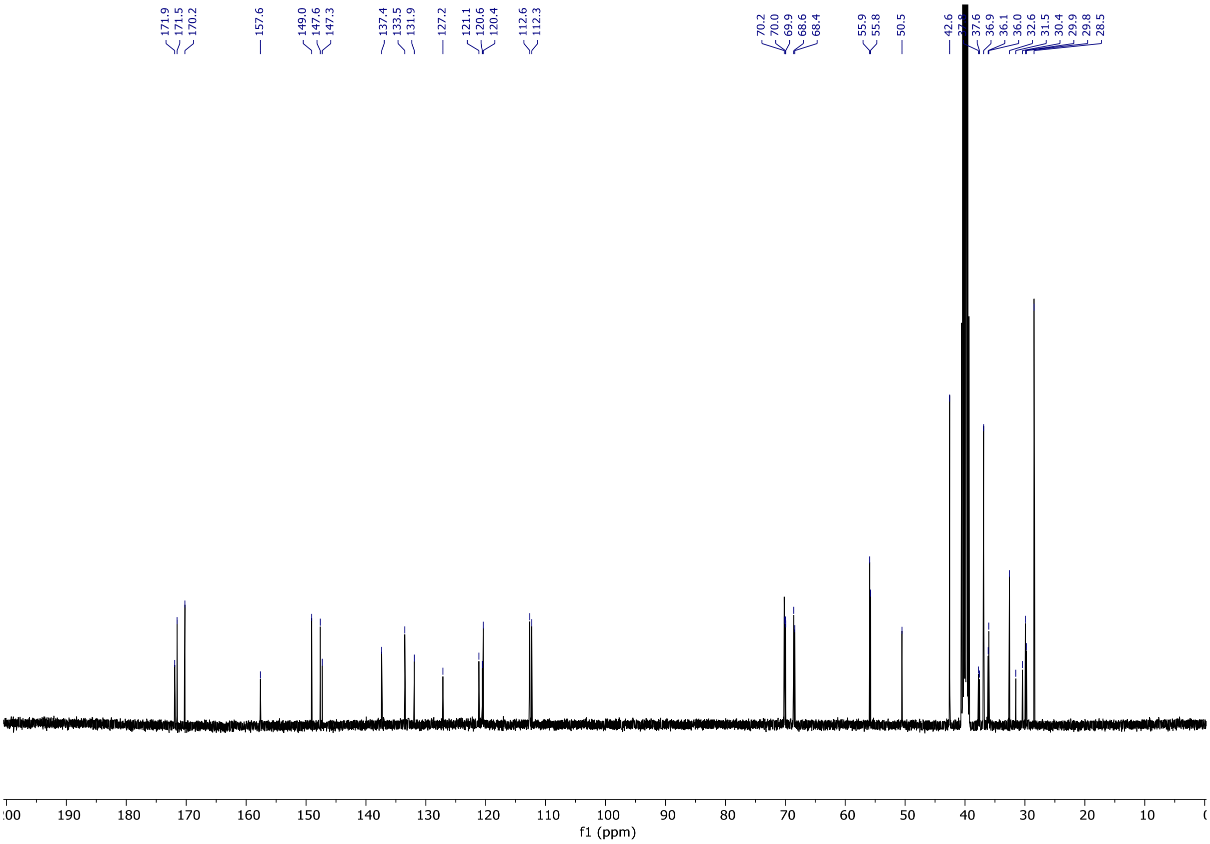

*N*-(6-(2-((3*r*,5*r*,7*r*)-adamantan-1-yl)acetamido)hexyl)-3-(2-(3-(3,4-dimethoxyphenyl)propanamido)benzo[*d*]thiazol-6-yl)propanamide **25**

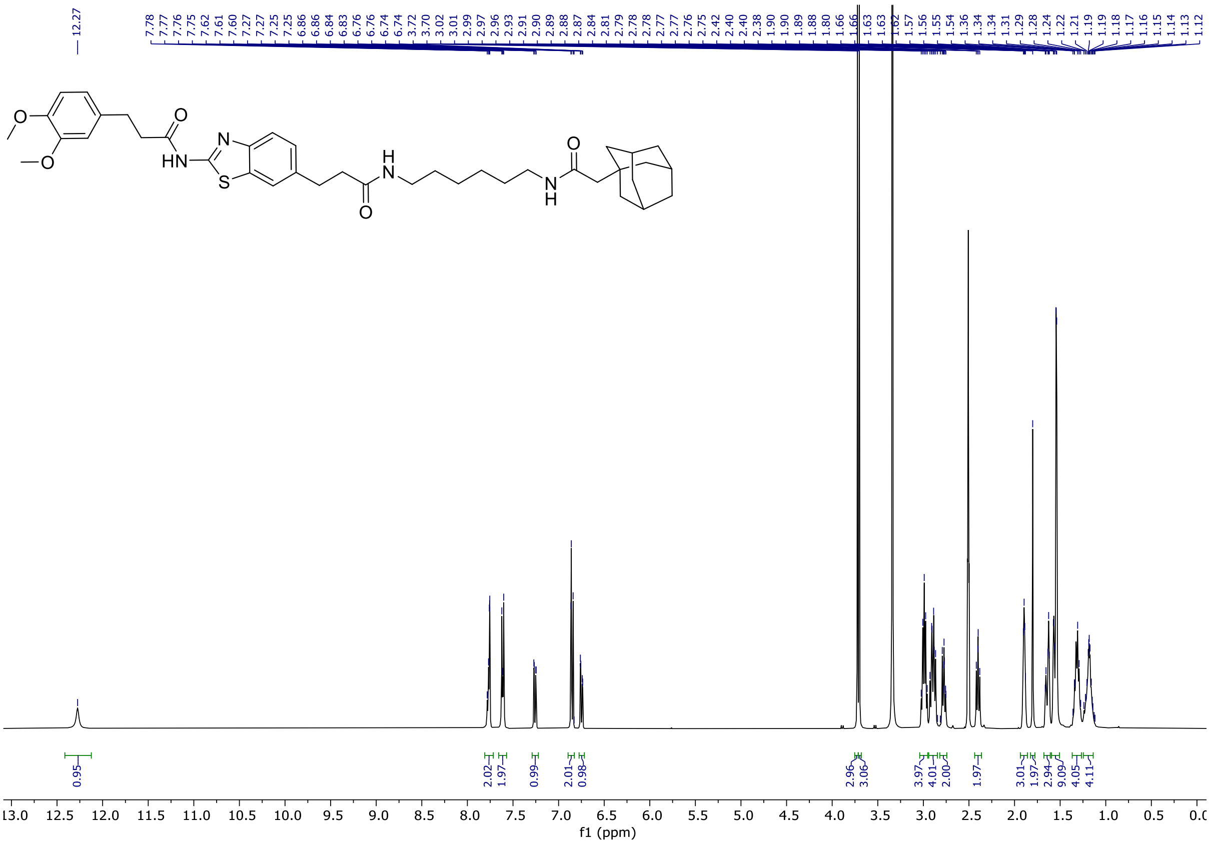


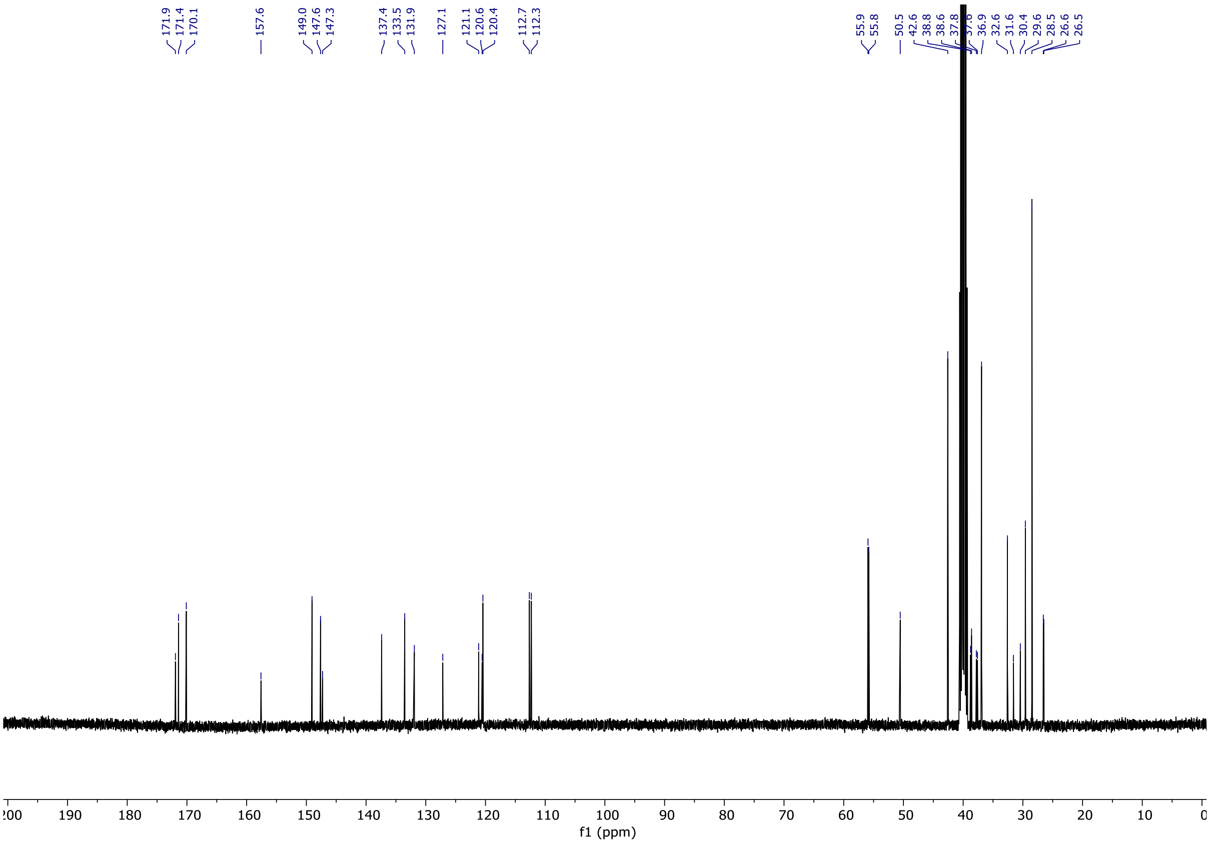

5-((6-(3-(2-(3-(3,4-dimethoxyphenyl)propanamido)benzo[d]thiazol-6-yl)propanamido)hexyl)carbamoyl)-2-(6-(dimethylamino)-3-(dimethyliminio)-3H-xanthen-9-yl)benzoate **27**

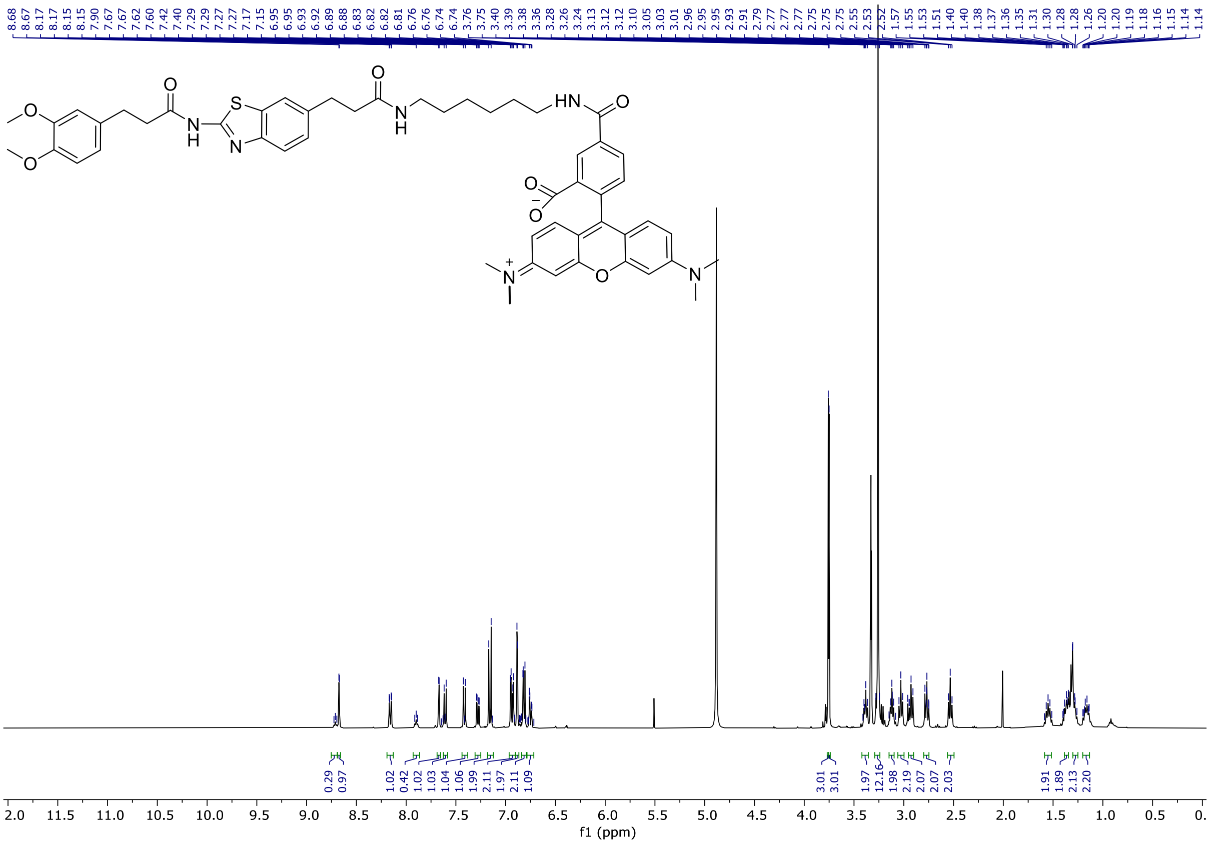


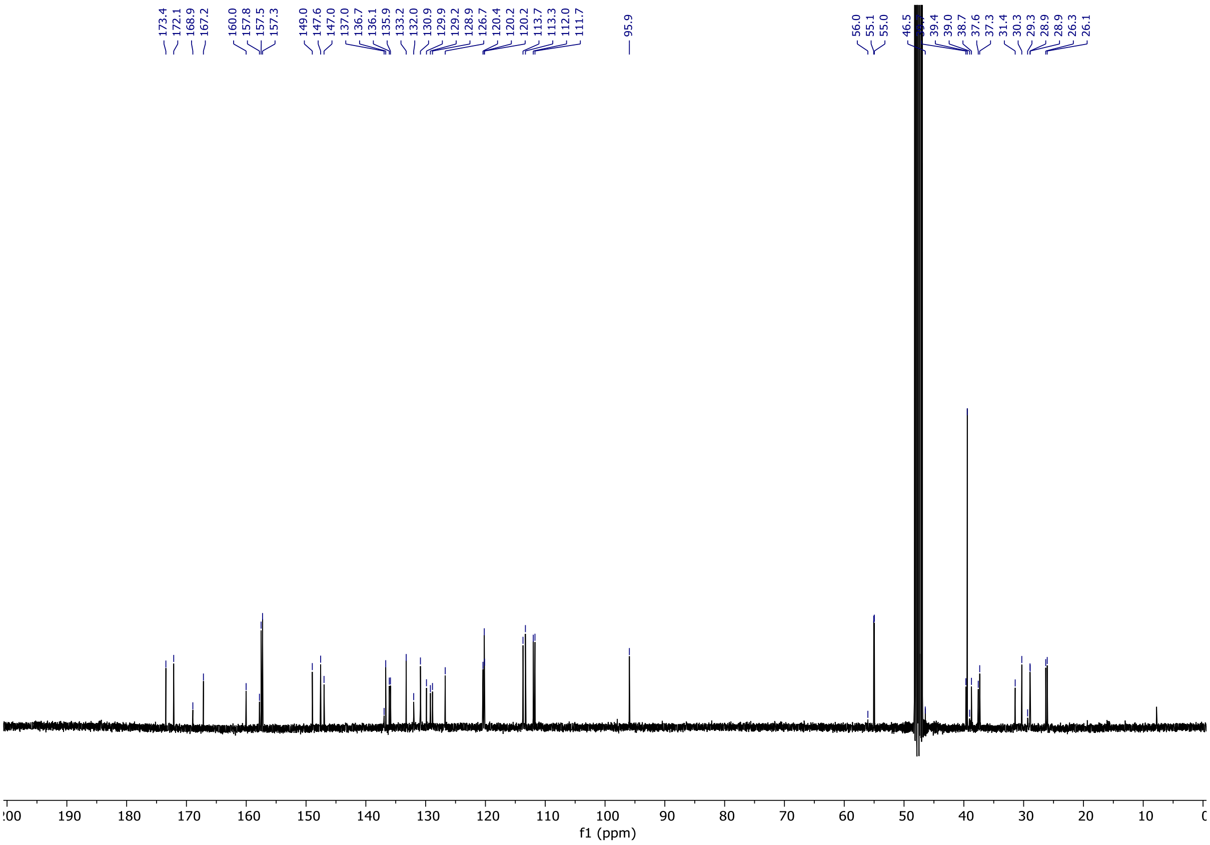

### LCMS purity

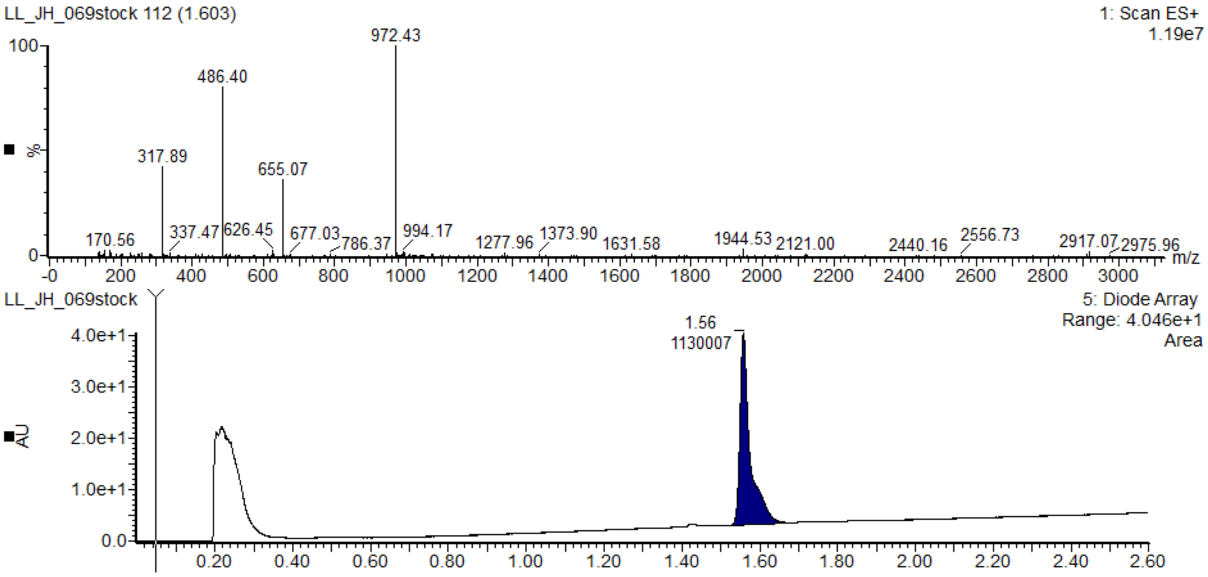


**SI Figure 9:** LCMS data for a sample taken from a DMSO stock for PROTAC **18**.

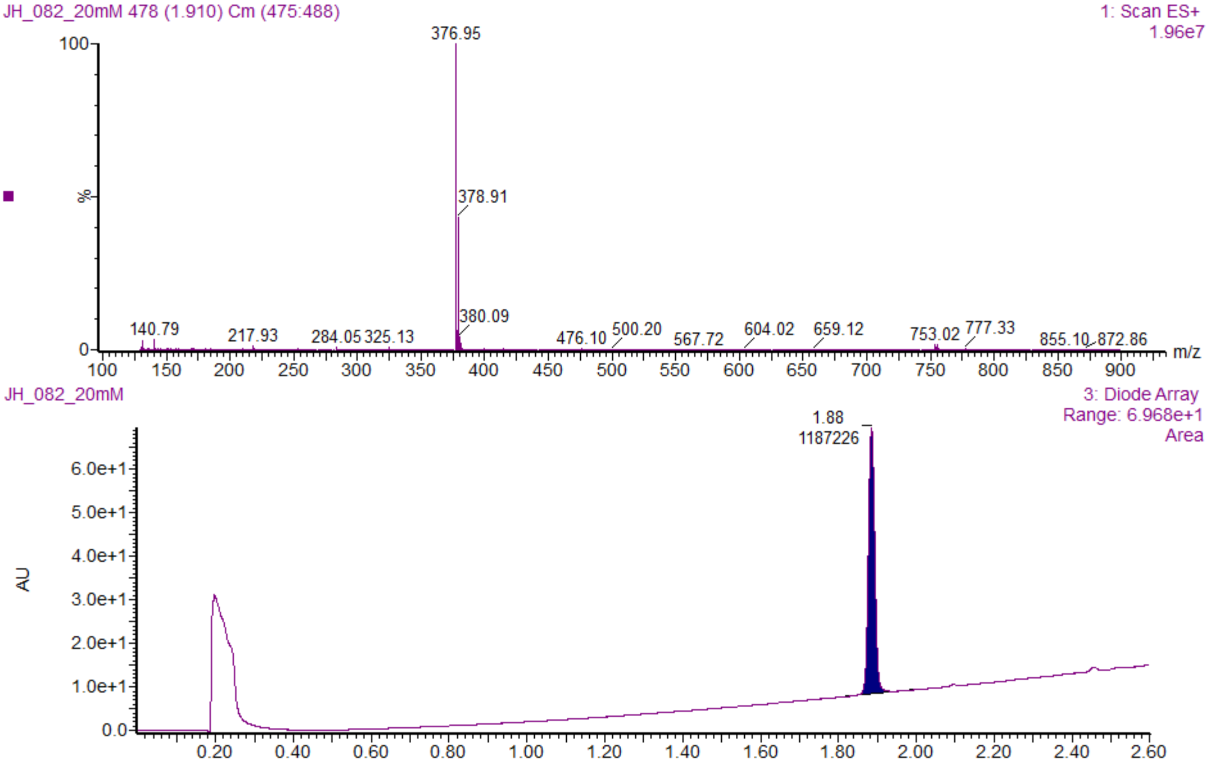


**SI Figure 10:** LCMS data for a sample taken from a DMSO stock for KY02111.

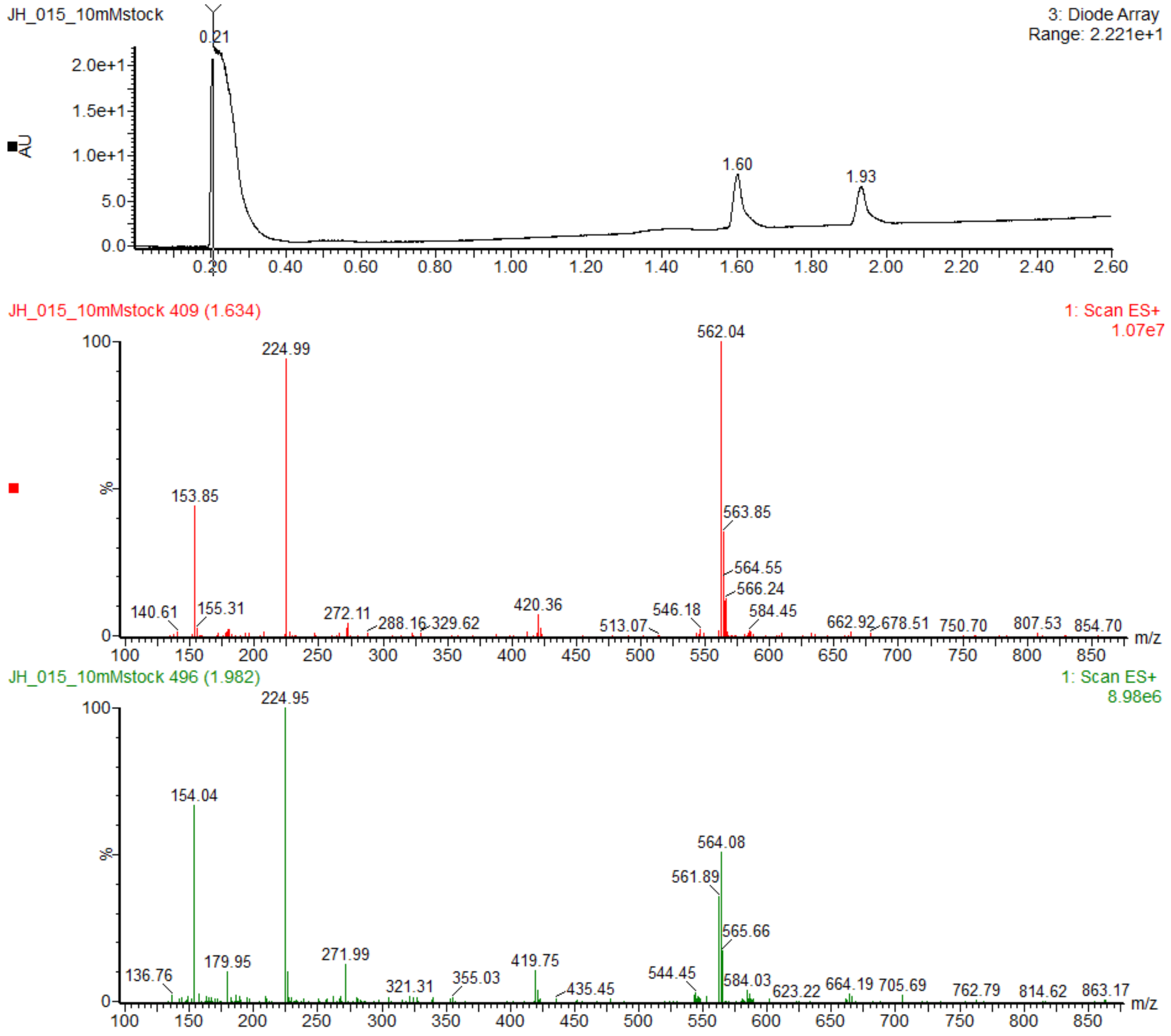


**SI Figure 11:** LCMS Data for a sample from a SQSI stock in DMSO. Two atropisomers are visible.
